# ARTIC RSV amplicon sequencing reveals global RSV genotype dynamics

**DOI:** 10.1101/2024.08.23.609324

**Authors:** Daniel M Maloney, Goncalo Fernandes, Seema Jasim, Tom Williams, Sarah Namugenyi, Michael Carr, Jennifer R Meyer, Aryan Sharma, Louise Marshal, B Ethan Nunley, Hong Xie, Mateo Carvajal, Henry D. Kunerth, Arif Tanmoy, Áine O’Toole, Amelia Weixler, Kelsey R Lanter, Gabrial Gonzalez, Jaydee Sereewit, Rommel Guevara, Matt Loose, Padraic M. Fanning, Denisse Benítez, Juan Carlos Fernandez, Paúl Cárdenas, Daniel Hare, Alex Greninger, Thomas Williams, Gaia Nebbia, Anthony C. Fries, Patrick McClure, Pavitra Roychoudhury, Xiong Wang, Senjuti Saha, Rebecca Dewar, Kate Templeton, Andrew Rambaut

## Abstract

Respiratory syncytial virus (RSV) is a leading cause of lower respiratory tract infections (LTRIs) in young children and adults over 65, contributing significantly to global healthcare burdens. With the recent approval of multiple pharmacological interventions for RSV, there is an increased demand for efficient, high-throughput sequencing methods to monitor RSV genetic diversity and any potential impact these interventions may have. Here we introduce two novel amplicon-based sequencing schemes designed for RSV A and B, optimised for integration with widespread existing ARTIC sequencing workflows. We demonstrate that these primer schemes can produce high quality genomes from RSV samples across the globe, with eight laboratories in five countries generating complete genomes on both Nanopore and Illumina sequencing platforms. The ability to effectively multiplex these RSV A and B primer schemes, enables streamlined, high-throughput sequencing without prior subtyping. Furthermore, these results provide a snapshot of the circulating diversity of RSV. Phylogenetic analysis of the 882 samples sequenced for this study suggests only minimal geographic clustering of RSV sequences, underscoring the global nature of RSV spread. It also highlights the distinct lineage dynamics seen between RSV A and B. This study represents an advancement in RSV genomics, providing robust tools for global sequencing efforts aimed at tracking RSV evolution and assessing the efficacy of new therapeutic interventions both rapidly and at scale.

## Introduction

Respiratory syncytial virus (RSV) remains one of the leading causes of lower respiratory tract infections (LTRIs) in young children and represents a significant seasonal burden on healthcare systems around the globe. Annually, there are in the region of 33 million RSV-associated acute LTRI cases in children under the age of 5 years, with around 10% requiring hospital admission (Y. Li et al. 2022). Despite a significant perturbation to the usual seasonal waves of RSV infections during the SARS-CoV-2 pandemic (Chuang et al. 2023), levels of RSV infection have returned to pre-pandemic levels (Meslé et al. 2023). In England, some 33,000 cases of RSV in children under 5 results in hospitalisation annually (Reeves et al. 2017). The burden of disease is highest in infants, with an estimated 28,561 admissions in infants <1 years of age in England and Scotland (Williams et al. 2023). Pre-pandemic bronchiolitis case levels in children under 2 in Scotland were estimated to be ∼37.7 per 1000 children, with RSV infection suspected to account for ∼78% of those cases (Chung et al. 2020).

Now that multiple novel pharmacological interventions for RSV have been approved there is a greater need than ever to sequence RSV in a reliable, high throughput and affordable manner. The Food and Drug Administration in the USA have recently approved two vaccines and one monoclonal antibody for the treatment of RSV. Abrysvo (RSVpreF, Pfizer (Walsh et al. 2023)) and Arexvy (RSVpreF3, GSK) are both vaccines targeting the F protein of RSV, while Beyfortus (nirsevimab, AstraZeneca and Sanofi (Hammitt et al. 2022) is a monoclonal antibody also targeting the F protein. In the UK, the Joint Committee for Vaccination and Immunisation (JCVI) have chosen Abrysvo as the intervention of choice for both older adults and pregnant mothers (Introduction of new NHS vaccination programmes against respiratory syncytial virus (RSV), 24^th^ of June, 2024). At the time of writing only limited data on the efficacy of these treatments outside of clinical trials is available, however early adopters of nirsevimab such as Spain have shown it to be 70% effective in preventing hospitalisations for RSV positive children with LTRIs (López-Lacort et al. 2024) and 90% effective in an American study (Moline et al. 2024), although a small sample size limits the latter study.

RSV is a single stranded RNA virus with a genome ∼15 kilobases (kb) in length and has two subgroups, A and B (Mufson et al. 1985). The two subgroups maintain genome synteny but only show a sequence similarity of ∼85% at the nucleotide level. They regularly show differences in the seasonality of cases, frequently with either A or B dominating infections in a given year (Obando-Pacheco et al. 2018).

The overall levels of genetic diversity within subgroups during a typical RSV season has not been thoroughly investigated, but recent efforts to develop an updated lineage system for RSV will aid in answering this question (Goya et al. 2024). Another key factor in characterising this diversity is the availability of reliable high-throughput RSV sequencing approaches. Currently widely used RSV sequencing methods work well but typically rely on longer amplicons up to 4.3 kb in length in multiple separate PCR reactions per sample (Dong et al. 2023; Schobel et al. 2016). Such methods often then require the separate quantification of each PCR reaction prior to pooling, which can cause considerable added labour and limit high-throughput sequencing efforts, and any amplicon dropout results in significant loss of data. To circumvent some of these issues we sought to develop a novel high-throughput amplicon-based RSV sequencing approach.

The widespread adoption of the ARTIC methodologies for SARS-CoV-2 sequencing means there is a readily available virus genomics infrastructure in much of the globe that can be leveraged simply by switching the primer scheme to target a different pathogen. With this in mind, we focused efforts on designing a 400bp fragment size amplicon scheme to directly slot into existing sequencing workflows. Current SARS-CoV-2 workflows routinely multiplex hundreds of samples and would allow a large-scale increase in RSV sequence generation.

Here we detail the development and validation of novel RSV A and RSV B tiled amplicon primer schemes to sequence clinical samples here in Scotland and describe the efficacy of these primers in a range of different labs around the world. We then use this information to investigate a subset of the diversity of currently circulating RSV A and B viral genomes.

## Results

### Internal pilot results

The RSV A and B amplicon schemes achieved high levels of genome completeness using either Nanopore or Illumina sequencing platforms. Figure 1 A and B show sequencing results on both Nanopore and Illumina platforms for RSV A and B samples. For Nanopore, median genome completeness was 94.9% and 95.8% for RSV A and B respectively (RSVA, n = 107; RSVB, n = 99). Illumina sequencing showed higher median genome completeness at 97% and 97.8% for RSV A and B respectively (RSVA, n = 189; RSVB, n = 170).

**Figure 1.**
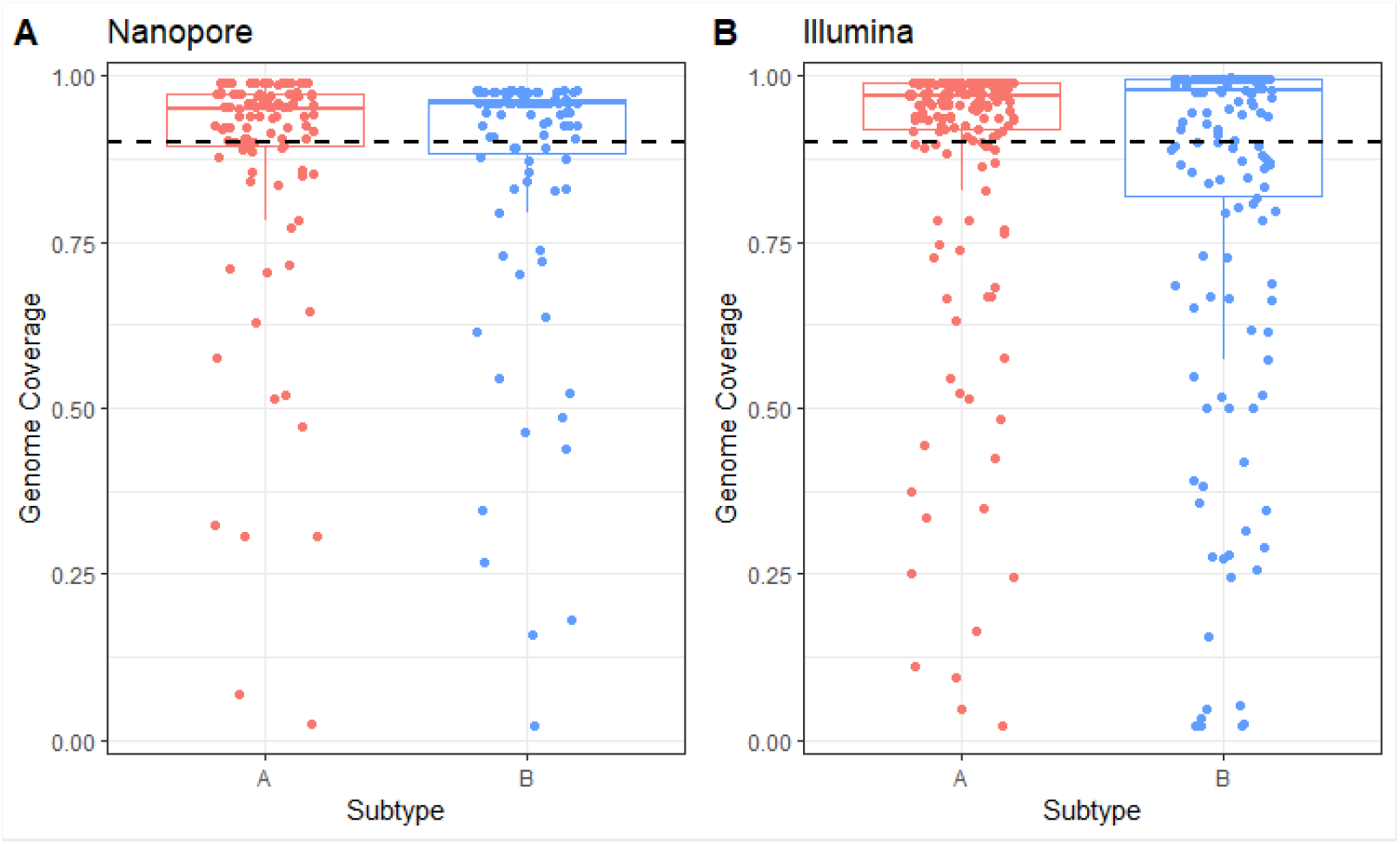
Genome Coverage of RSV A (red) and B (blue) samples sequenced using either **A)** Nanopore sequencing (RSVA n = 107; RSVB n = 99) or **B)** Illumina sequencing (RSVA n = 189; RSVB n = 170). Dashed line represents 90% genome coverage.

Sample selection was not based on a Ct cut off as not all samples had Ct values available, but most samples that had recorded Ct values were less than 30 (Illumina n = 186 of 208, 359 samples sequenced in total; Nanopore n = 194 of 206, 206 samples sequenced in total). Some samples up to Ct 35 achieved >90% genome completeness, whilst others with Cts as low as 20 achieving ∼25% coverage. This suggests that Ct alone is not predictive of genome completeness. It is possible the low Ct samples with poor genome coverage had low quality extractions and thus low starting input, samples could have been stored at unideal conditions prior to extraction, or there could have been inhibitors present that prevented downstream amplification. Only 35 samples were sequenced on both platforms and all but five of these sequenced better using Illumina technology (86% of samples; Supplemental Figure 1). Throughout this work a threshold of 90% genome completeness has been used for further analysis, and it is worth noting that despite 30 samples having higher quality sequencing outputs using Illumina than Nanopore, only three of these samples crossed the 90% genome completeness threshold.

### External sequencing results

To establish and validate a new primer scheme for a given pathogen, it must be shown to perform well beyond the developer’s own lab. In addition to variation in methodologies particular to a given lab, the circulating diversity of the virus in question may be distinct enough to impact the performance of the primer scheme in other geographic locations. To investigate this, we established a network of 8 labs from 5 different countries (Table 1) who sequenced RSV A and B samples using these primer sets. Samples were sequenced using both Illumina and Nanopore sequencing platforms, see methods section for details.

**Table 1.** Details of all external labs submitting RSV sequences using the primer schemes outlined in this study.

| External Lab | Country | Platform |
| --- | --- | --- |
| Child Health Research Foundation (CHRF) | Bangladesh | Illumina |
| Guy's and St. Thomas Hospital (GSTT) | England | Illumina |
| Minnesota Department of Health (MDH) | USA | Nanopore |
| National Virus Reference Laboratory (NVRL) | Ireland | Nanopore |
| University of Nottingham (UoN) | England | Nanopore |
| U.S. Air Force School of Aerospace Medicine (USAF) | USA | Illumina |
| Universidad San Francisco de Quito (USFQ) | Ecuador | Nanopore |
| University of Washington (UW) | USA | Illumina |

The primer schemes performed well for all testing labs, with a median sequencing coverage of 96.14% and 96.12% across all external RSV A and RSV B samples respectively (RSV A n = 177; RSV B n = 186; See Figure 2 A and B). Of the eight external sites four used Illumina platforms (CHRF, GSTT, USAF and UW) and four used Nanopore (MDH, NVRL, UoN and USFQ). In general, samples sequenced on Illumina platforms had more complete genomes than those on Nanopore platforms.

**Figure 2.**
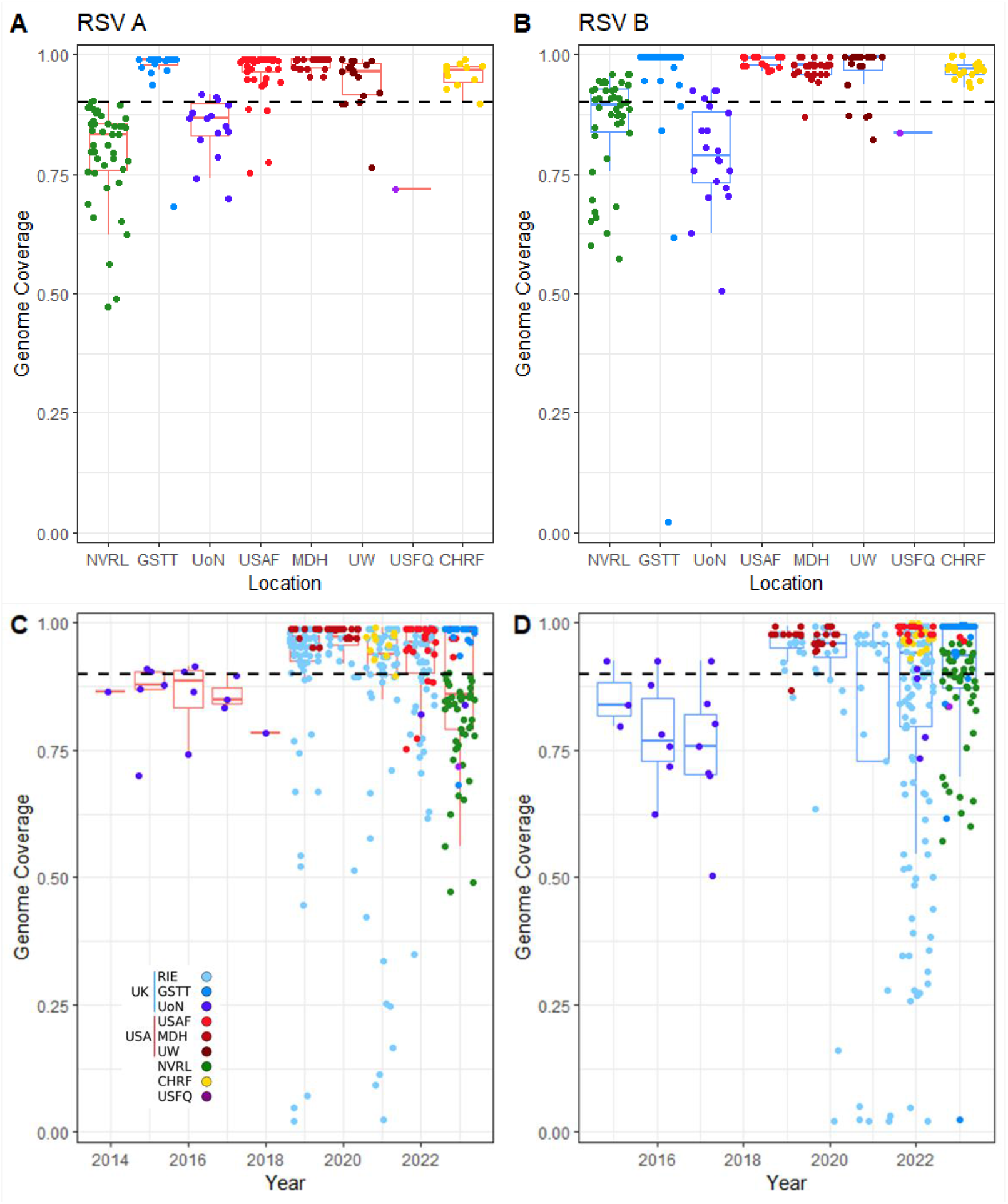
Genome coverage of RSV A (red boxplots) and RSV B (blue boxplots) f rom 8 external laboratories. **A)** and **B)** show plots of genome coverage f rom each external lab. **C)** and **D)** show sequences separated by collection year with dots coloured by external lab. Dashed line represents 90% genome coverage. Note that NVRL and UoN used Nanopore whereas all others used Illumina based sequencing approaches. CHRF - Child Health Research Foundation (Bangladesh); GSTT - Guy’s and St. Thomas Hospital (England); MDH - Minnesota Department of Health (USA); NVRL - National Virus Reference Laboratory (Ireland); RIE Royal Inf irmary of Edinburgh (Scotland); UoN - University of Nottingham (England); USAF - U.S. Air Force School of Aerospace Medicine (USA); USFQ - Universidad San Francisco de Quito (Ecuador); UW - University of Washington (USA).

The samples sequenced externally spanned a range of collection dates from 2014 to the start of 2023. Sequences above 90% genome completeness were obtained for 7 out of the 10 previous years.

### Global lineage diversity

The recent development of a novel lineage classification system for RSV has facilitated finer resolution of circulating RSV diversity (Goya et al. 2024). To assess this, we built maximum-likelihood phylogenetic trees of both RSV A and B using all newly acquired sequences with >90% genome coverage (n = 322 RSVA and n = 298 RSVB samples), using a high-quality set of sequences as the background (See methods for full details). We then assigned RSV lineages to these sequences using Nextclade (Figure 3 A and B). Both the RSV A and B trees demonstrate that the newly sequenced RSV samples for this study are broadly representative of the observed recent diversity of RSV. As these sequences were all at least 90% complete, this suggests the primer schemes developed here can successfully capture a wide range of the circulating diversity of both RSV A and B. As may be anticipated for a respiratory virus, there appears to be minimal geographic clustering of samples on the phylogenetic tree, however we do observe a temporal signal in samples as they cluster by year of sampling (Supplementary Figure 2).

**Figure 3.**
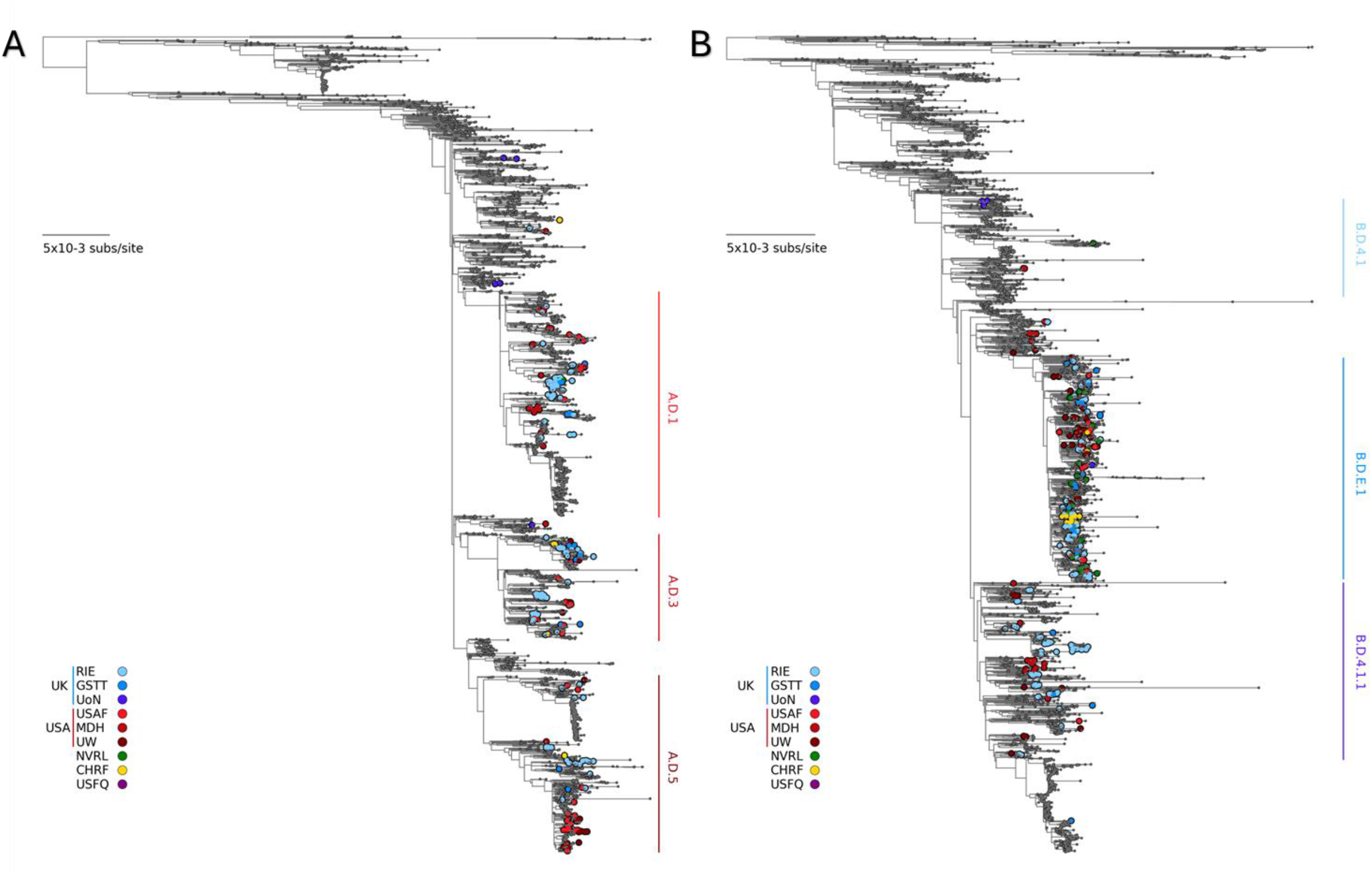
Midpoint rooted maximum likelihood tree of all internal and external RSV sequences with at least 90%genome coverage, with all “high quality”‘ RSV sequences available on GISAID as background. **A)** showsRSV A samples and **B)** showsRSV B. Samplesare coloured by the Location of the submittingLab, highlighting widespread dispersal of sequences acrossthe tree. Scale bar=5×1o-3 substitutions per site.

Of note, RSV A and B show distinct lineage dynamics. When assigning lineage to the >90% complete RSV genomes, RSV A samples were predominantly one of three distinct lineage groups, A.D.1, A.D.3 and A.D.5, whereas most RSV B samples belonged to a single lineage, B.D.E.1, which dominates the recent observed diversity of RSV B (Figure 4, Supplementary Figure 2).

**Figure 4.**
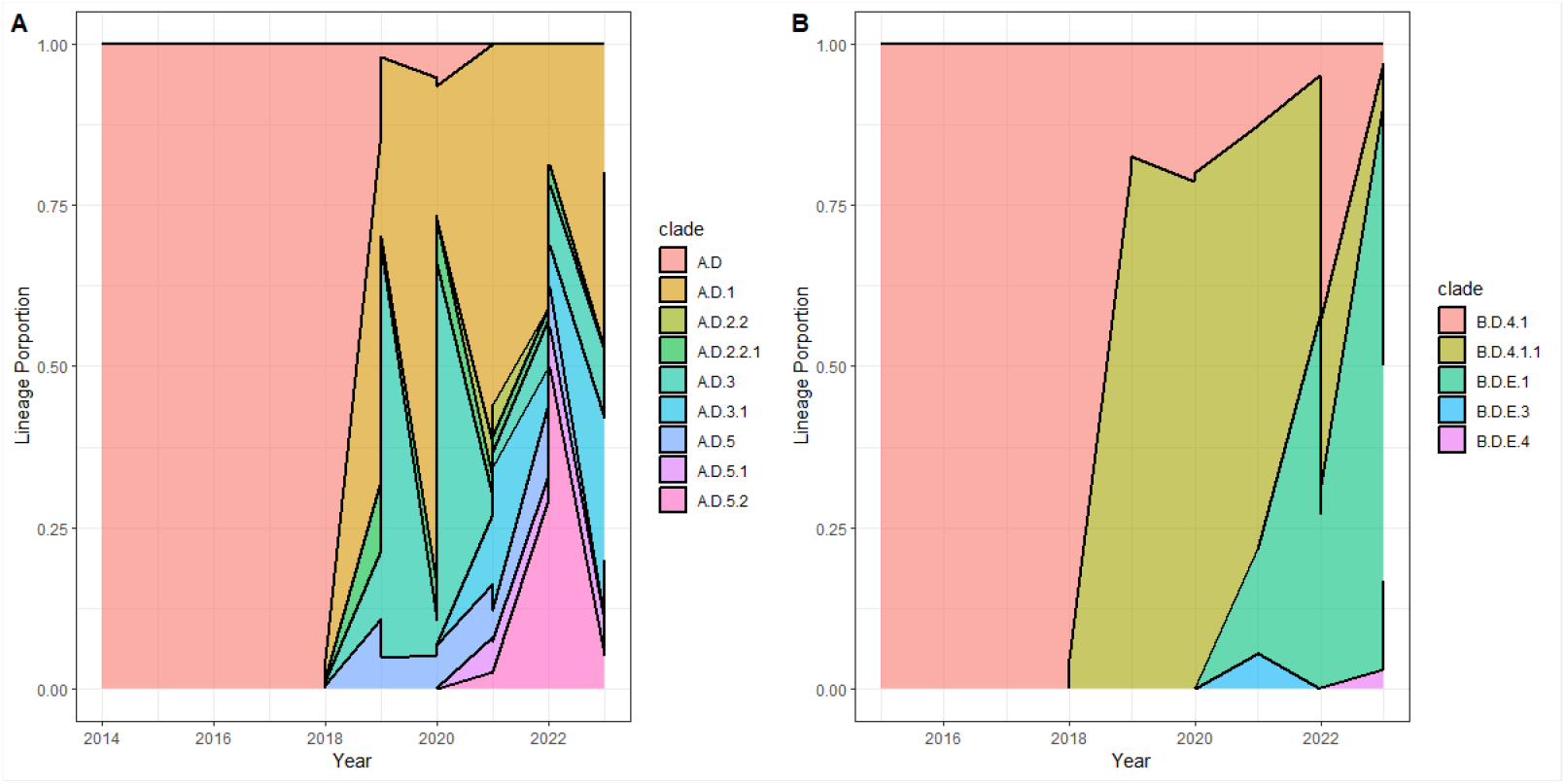
Breakdown of the proportion of RSV A and B lineages across the samples sequenced within this study that could have a lineage assigned by Nextclade (n = 448 and 434 for RSV A and RSV B respectively).

### F and G gene coverage

The F and G genes of RSV A and B represent the two major neutralising antibody targets on the surface of RSV viral particles (McLellan, Ray, and Peeples 2013). As such they are of keen interest to researchers analysing any potential immunological changes due to the introduction of the new pharmaceutical interventions targeting the F and G proteins and it is important that any viable primer schemes show robust coverage of the F and G genes. Across this study a total of 882 unique RSV samples were sequenced. The median genome completeness for the F and G proteins for both RSV subtypes was 100% (results summarised in Table 2; Figure 5).

**Table 2.**
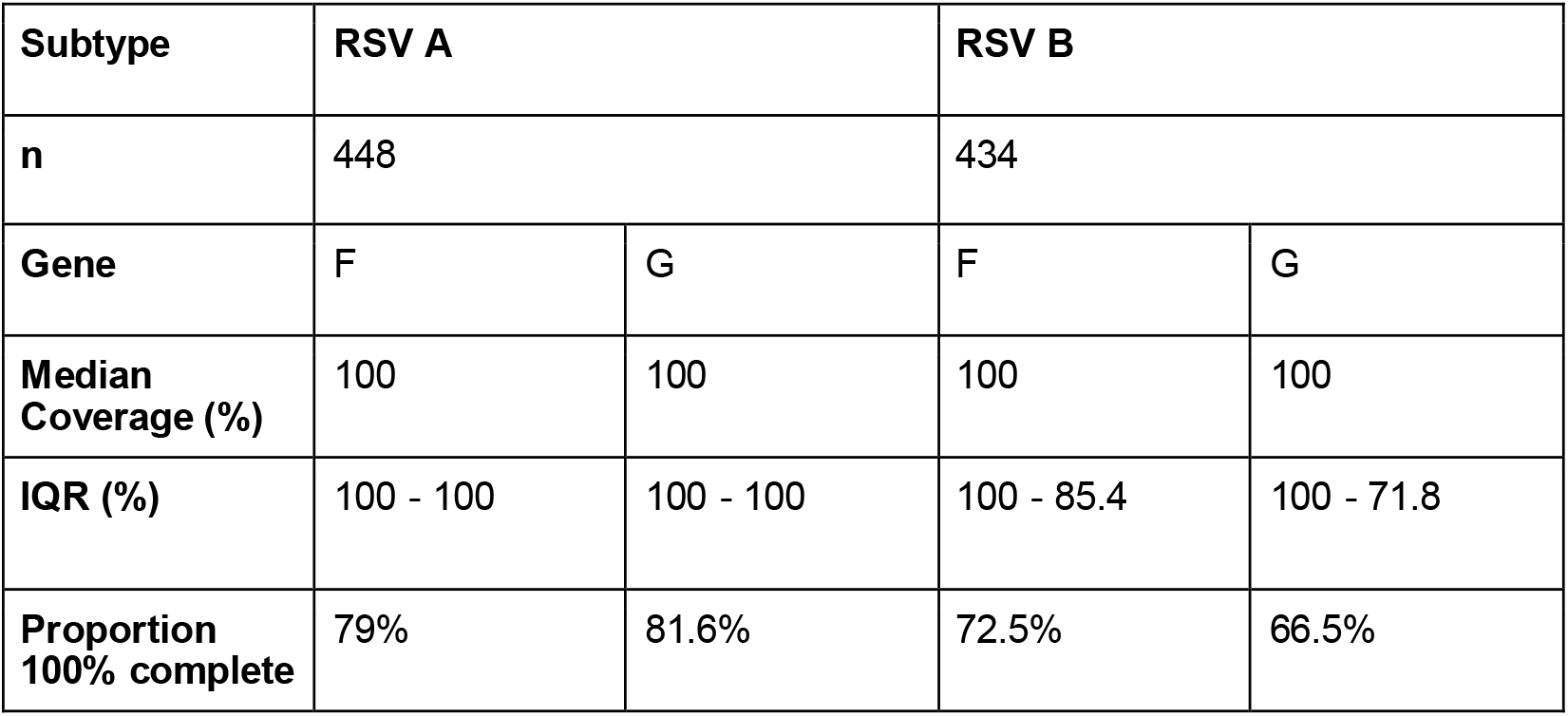
Details of the F and G gene coverage of the RSV A and B samples sequenced in this study.

**Figure 5.**
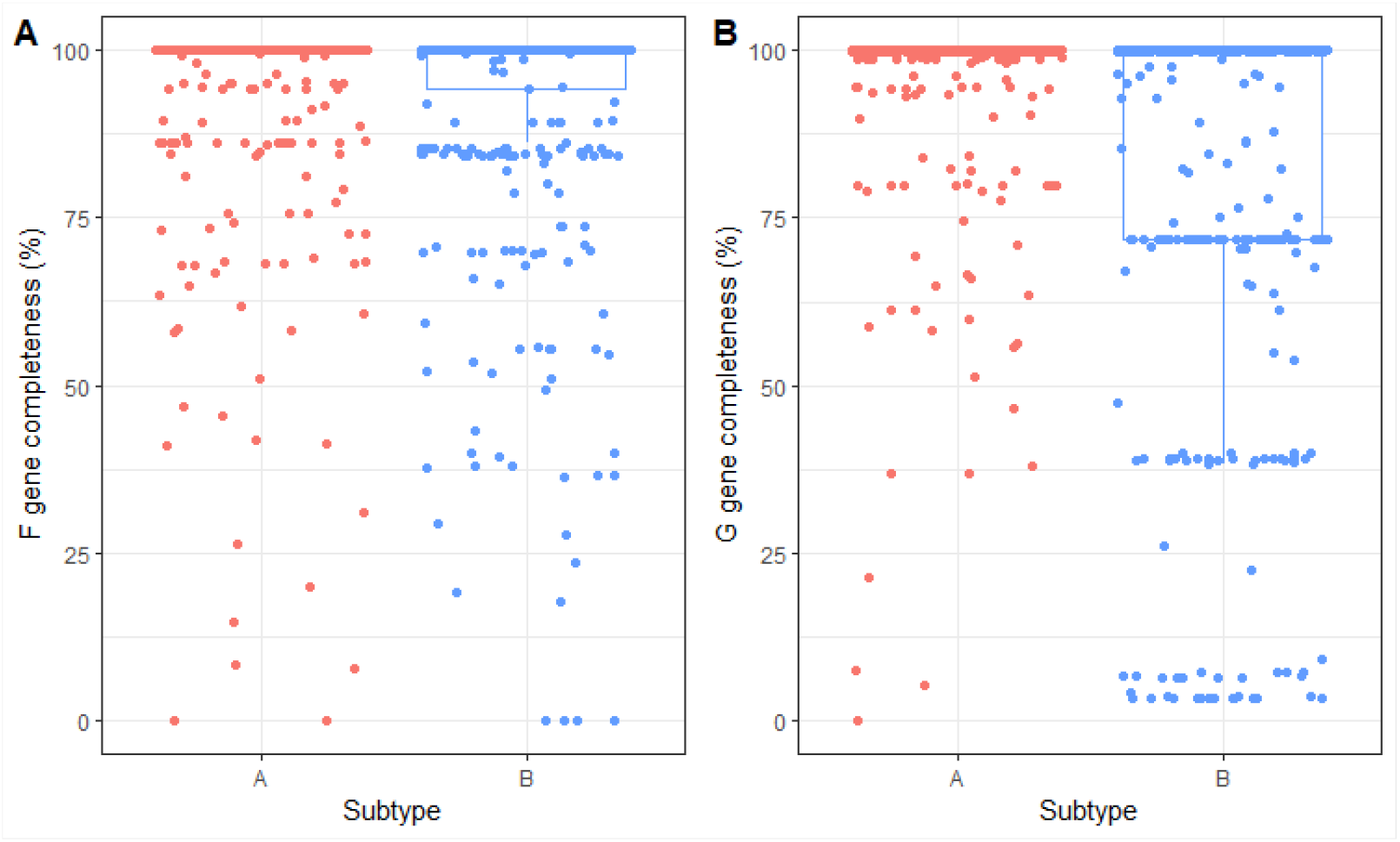
Boxplots of **A)** F gene and **B)** G gene completeness. Graphs are separated by RSV subtype with RSV A in red and RSV B in blue (n = 448 and 434 for RSV A and RSV B respectively), representing all sequenced samples f rom both Illumina and Nanopore platforms.

### Primer Multiplexing

A current limitation of large amplicon scheme sequencing for RSV is the need to distinguish between RSV subtypes prior to commencing sequencing. Many routinely used diagnostic PCRs only test for presence or absence of RSV, rather than the subtype itself, which then leads to delays and additional expense due to conducting an additional RT-PCR for subtyping prior to sequencing. A concern with mixing two primer schemes for closely related viruses is that primer mis-binding between the two schemes cannot be easily ruled out. For instance, a primer from the RSV B scheme could bind to RSV A to produce a product not expected when using the RSV A scheme alone. This interaction can have serious consequences and impact the quality of the final genome sequence, due to primer sequences not being completely removed prior to consensus sequence generation potentially leading to erroneous SNP or reference calls at these primer mis-binding sites.

To address this, we have developed a bioinformatic solution to this primer mis-binding problem for Nanopore sequencing approaches. After mapping, our pipeline checks that the start and end positions of a given sequencing read are located within an expected primer binding site according to the bed file for the appropriate primer scheme and excludes sequencing reads that lie outside these expected boundaries (see methods for further details). To validate our approach, we sequenced 69 samples each of RSV A and B both with the A and B primer schemes both multiplexed and individually. Samples with a wide range of Cts (min: 13.8, max: 41.1, mean: 24.7) were chosen. The multiplexed primers performed well and were able to generate consensus genomes at similar levels of completeness as samples sequenced with the single correct primer scheme (Figure 6). There was a small but statistically significant decrease in coverage when using the primer schemes multiplexed together compared to individually (Paired Wilcoxon signed rank test: RSV A, n = 69, p = 2.2×10^−9^; RSV B, n = 69, p = 1.8×10^−7^). When sequencing an RSV A sample, there was a mean decrease in the difference of genome coverage of 7.07% when using the multiplexed primers, and for RSV B samples there was a mean difference of 5.17%.

**Figure 6.**
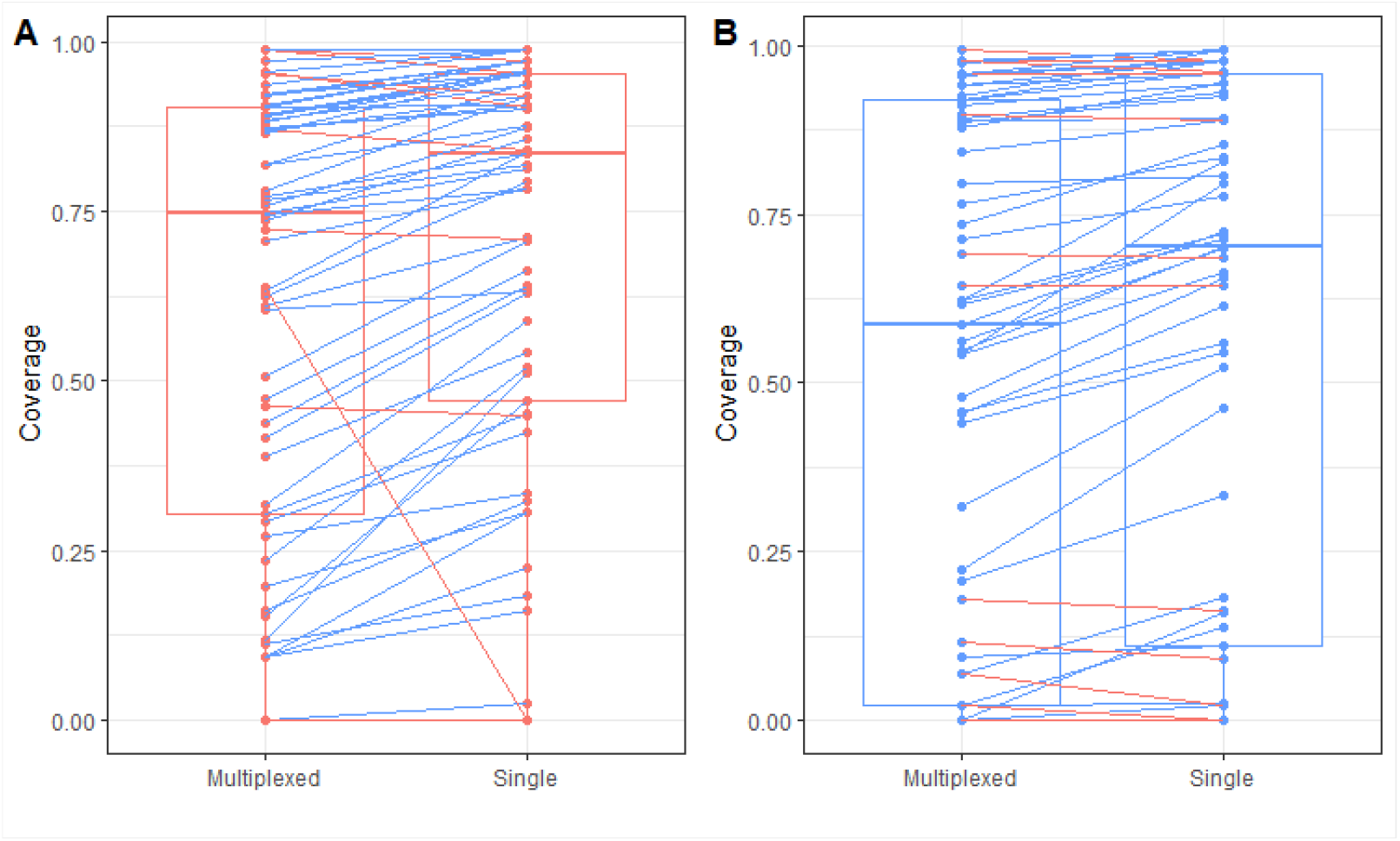
Genome coverage of samples using either the multiplexed A and B primer schemes or just the appropriate primer scheme singly. A) RSV A samples and B) RSV B samples. Lines between points connect the same samples sequenced with either method, with blue lines indicating the single primer scheme resulted in higher genome coverage and red indicating the multiplexed primer scheme did. A modest but statistically significant decrease in genome coverage when using the multiplexed primers was observed.

Despite a decrease in mean coverage when using the multiplexed primers there were no differences in lineage assignment caused by using the different sequencing methods. This demonstrates that RSV genomic insights can be gained quicker and cheaper using the primer multiplex approach compared to using the singleplex primers schemes.

## Discussion

Here we introduce two novel ARTIC amplicon schemes for sequencing RSV A and B. We have demonstrated that both primer schemes perform well and capture the circulating diversity of on RSV A and B samples in an array of labs from a range of sites across the globe and can also produce informative, near complete genomes on samples as old as 2015. The ability for a diverse range of labs, from healthcare to research settings, to quickly use these primer schemes to generate high quality RSV genomes highlights the benefits of the ARTIC style of amplicon-based primer schemes for large scale genome sequencing efforts.

Initial sequencing results were strong, with high median genome coverage for both RSV A and B samples in singleplex assays (Figure 1). Furthermore, almost all samples sequenced had complete coverage of both the F and G genes which represent a key target for current and future vaccine development efforts (Figure 5). This study presents an important step forward by assessing the multiplexing of both RSV A and B primer schemes and developing the bioinformatics package that allows streamlined, robust analysis from the command line or ONT’s EPI2ME platform (labs.epi2me.io). The ability to multiplex these schemes and get accurate lineage classification of RSV lineages removes the need to type samples before moving on to sequencing and represents a step forward f rom large amplicon scheme sequencing efforts for RSV. However, the multiplexing currently comes at a cost to coverage of some samples and future work will focus on streamlining this process to minimise any coverage losses due to incorrect primer bindings (Figure 6).

The effective sequencing of historic samples suggest that these primer schemes can handle the changes in sequence diversity that can arise over ∼8 years of natural selection, providing an indication of how these primers could perform going forward. There was a noticeable dip in coverage of the oldest samples in this study, however this could be due to the choice of Nanopore as the sequencing platform. Genomes with >90% coverage were obtained for the older samples and included in the phylogenetic analysis (Figure 3).

We also show that the observed circulating diversity of RSV is shared across the geographic regions within this study, despite considerable distances between sites (Figure 3). A caveat to this is the lack of sequences from the Southern hemisphere in this study, where differences in climate alter RSV seasonality. However, there is work ongoing to assess the performance of these primers in KEMRI, Kenya and these primers have performed well on Australian samples (Doherty Institute for infection and immunity, private correspondence). It is interesting to note that the sequences within this study did not tend to strongly cluster with sequences from similar geographic regions, but showed more dispersal across the tree, suggesting large scale import and export of RSV between the regions surveyed.

Our phylogenetic analysis also highlighted distinct RSV A and B dynamics, such that RSV A has three distinct clades co-circulating at roughly equal proportions (assigned lineages A.D.1, A.D.3 and A.D.5 and respective sub-lineages) and the majority of RSV B samples belong to a single clade (lineage B.D.E.1), that has almost entirely replaced the previously dominant RSV B lineage (B.D.4). Further work needs to be done to establish if this holds true on a larger scale, and we believe our primer schemes outlined here will help in this effort.

The novel primer schemes and bioinformatics tools for consensus generation outlined here represent a constructive step towards high-throughput, robust sequencing of RSV. The ease of use of these primer schemes, coupled with the straightforward integration into existing ARTIC amplicon-based sequencing methodologies, will help bolster the global efforts to sequence RSV on the large scale needed to capture the ever-changing landscape of RSV genomics and interventions.

## Methods

### Primer Scheme Design

Primer schemes were designed using Primal Scheme (Joshua Quick et al. 2017). In an effort to capture global diversity 6 RSV A and 6 RSV B sequences available from GISAID chosen from geographically distinct sites and different epidemiological years were used as input. This resulted in two primer sets of 50 amplicons with a target amplicon size of 400bp. Primer sequences can be found on the arctic-rsv GitHub repository: https://github.com/artic-network/artic-rsv. Table 3 details the GISAID ID as well as the country or origin and year of collection of each sequence used.

**Table 3.**
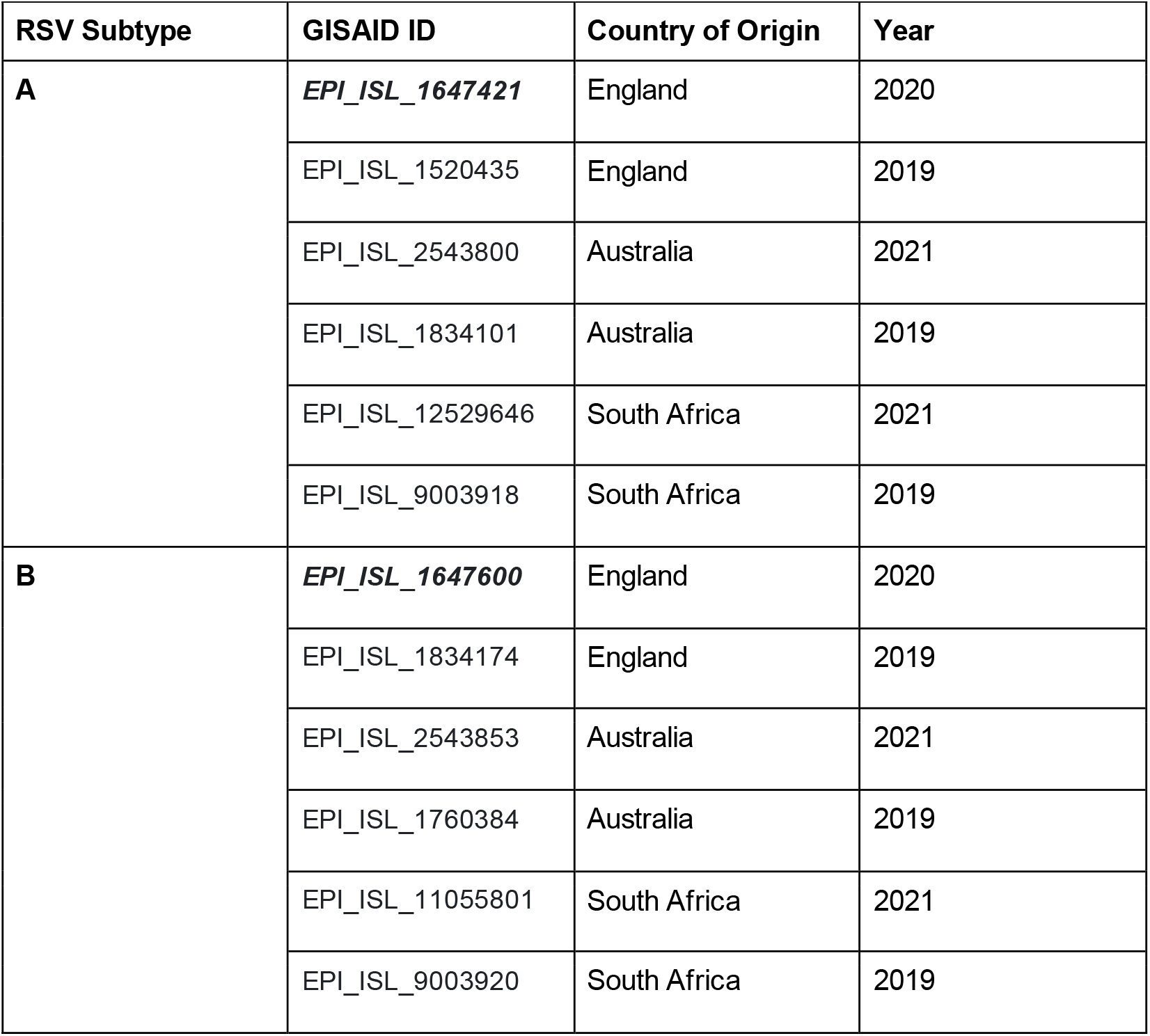
Details of the sequences used in designing the RSV A and B primer schemes. The ID italicised and in bold is the reference to which the primer bed files are indexed to.

| RSV Subtype | GISAID ID | Country of Origin | Year |
| --- | --- | --- | --- |
| **A** | ***EPI_ISL_1647421*** | England | 2020 |
|  | EPI_ISL_1520435 | England | 2019 |
|  | EPI_ISL_2543800 | Australia | 2021 |
|  | EPI_ISL_1834101 | Australia | 2019 |
|  | EPI_ISL_12529646 | South Africa | 2021 |
|  | EPI_ISL_9003918 | South Africa | 2019 |
| **B** | ***EPI_ISL_1647600*** | England | 2020 |
|  | EPI_ISL_1834174 | England | 2019 |
|  | EPI_ISL_2543853 | Australia | 2021 |
|  | EPI_ISL_1760384 | Australia | 2019 |
|  | EPI_ISL_11055801 | South Africa | 2021 |
|  | EPI_ISL_9003920 | South Africa | 2019 |

### Sample Selection – RIE

Patients attending hospital with respiratory symptoms are tested for the presence of SARS-CoV-2, Flu or RSV using the cepheid platform. Any positive nasopharyngeal and throat swab samples were referred to the virology diagnostic laboratory, NHS Lothian, in tubes containing 3 mL of Viral Transport Medium (VTM) or aspirate. After a positive RSV diagnosis, an aliquot of around 250μl were kept and stored at -80°C until use. A random selection of samples was chosen from these stored samples for sequencing in this study. Extraction Method - RIE: Viral RNA was extracted from 200 μl of Nasal and Throat Swab (NTS) sample in Molecular Sample Solution (MSS) using the NucliSENS EMAG, EasyMAG or the KingFisher Flex System and reagents (Biomerieux or MagMAX™ Nucleic Acid Isolation Kit), following the manufacturer’s instructions.

### RT-PCR - RIE

RSV positive samples were identified via Luminex NxTag Respiratory Pathogen Panel testing and CT values were established using the protocol outlined in (Templeton et al. 2004) on the Applied Biosystems 7500 Real-Time PCR System.

### Nanopore sequencing - RIE

Nanopore sequencing libraries were generated using a modified version of the SARS-CoV-2 ARTIC V3 loCost sequencing method (Josh Quick 2020) which can be found on the arctic-rsv GitHub repository: https://github.com/artic-network/artic-rsv. Briefly, for the synthesis of complementary DNA (cDNA) from the extracted RNA, genome amplification and library preparation, the NEBNext® ARTIC SARS-CoV-2 Companion Kit (Oxford Nanopore Technologies) was used, with two different PCR reactions being set-up for RSV A (pool 1 and pool 2) and a separate set used for RSV B (pool 1 and pool 2), and then pooled together at the start of library preparation. When using the RSV A and B primers in a multiplex reaction, primer pool 1 for both the RSV A and RSV B schemes were combined in equal concentrations (and the same for pool 2), with the remainder of the library preparation proceeding as normal. A positive control, a negative control, the negative extraction controls and one sample at random were quantified using the Qubit dsDNA HS kit.

Volume for loading on flow cell was calculated noting that 20 ng are required for sequencing in volume of 12 µl. For priming the flow cell and preparing/loading the sequencing library, Sequencing buffer, Loading beads, Flush buffer and Flush tether were used. Final libraries were loaded on R9.4.1 flow cells and run for 16 hours.

Detailed protocols can be found at: https://www.protocols.io/view/rsv-nanopore-whole-genome-sequencing-dhp835rw

### Illumina sequencing – RIE

Illumina Sequencing libraries were generated using the Illumina COVIDseq Test (RUO) kit as per the manufacturer’s recommendations but with the RSV primers exchanged for the ARTIC SARS-CoV-2 primers included in the kit. IDT for Illumina Nextera DNA Unique Dual Indexes Sets A–D allow the multiplexing of up to 384 samples per run. Libraries were sequenced the NextSeq 2000 platform using the P2 (200 cycle) flow cell.

Detailed protocols can be found at: https://www.protocols.io/view/rsv-illumina-whole-genome-sequencing-dhp935r6

### Genome Assembly - RIE

The sequence data was handled differently depending on which sequencing platform was utilised. For Nanopore data, an in house nextflow pipeline was used to assemble genomes, which can be found at https://github.com/artic-network/artic-rsv. This pipeline can be used either as a standalone tool or through ONT’s epi2me platform.

Briefly, ampli_clean (https://github.com/Desperate-Dan/ampli_clean) a custom python-based tool was first used to map reads to either the RSV A or B reference using minimap2 (H. Li 2018). These binned reads are then filtered using the provided primer bed files to remove any reads that do not map to a given pair of primer coordinates, thus removing some reads produced due to mispriming, before being passed to the “field bioinformatics” pipeline (v1.2.1) for consensus sequence generation (GitHub - artic-network/fieldbioinformatics: The ARTIC field bioinformatics pipeline).

For Illumina data, reads are first trimmed with Trim Galore (Krueger, n.d.), followed by mapping using BWA-MEM (H. Li 2013). Primer sequences were then trimmed, variants called, and consensus sequences generated using iVar (Grubaugh et al. 2019).

All genomes required a minimum depth of 20 reads for consensus generation. Coverage stats were acquired using Nextclade, as well as all lineage calls (Aksamentov et al. 2021). Maximum-Likelihood trees using the Jukes-Cantor model of substitution (Jukes and Cantor, 1969) were generated using IQTree v1.6.12 (Nguyen et al. 2015) including only sequences with at least 90% genome completeness. Resulting trees were midpoint rooted and rendered using baltic (https://github.com/evogytis/baltic).

Background datasets for RSV A and B were generated from available RSV sequences on GISAID by the 08/12/2023. Sequences were filtered using the “complete” and “high coverage” flags, then curated after download to remove any misidentified or highly divergent sequences. The full acknowledgement table can be found in the supplemental materials.

### Ethics Approval – RIE

Samples were tested using appropriate tissue bank approval for Scotland Southeast, with Lothian NRS BioResource RTB approval (REC ref – 20/ES/0061).

### Sequencing Strategy – CHRF

CHRF conducted studies to understand the RSV burden at the Bangladesh Shishu Hospital and Institute (BSHI) in Dhaka, Bangladesh. Nasopharyngeal swab samples were collected from children aged 0–59 months who were admitted at BSHI with suspected viral respiratory tract infections (following the WHO’s RSV hospital-based surveillance case definition). These samples were tested for RSV, Influenza A (FluA), and Influenza B (FluB) infections. The protocols were approved by the ethical review board of BSHI, and written informed consent was obtained from the guardians of children admitted to the hospital before sample collection.

Viral RNA was extracted from the swab samples using the Zymo Quick-DNA/RNA Viral MagBead Kit (Zymo Research, Orange, CA, USA), and qPCR was performed using the AgPath-ID One Step RT-PCR Kit (Thermo Fisher Scientific, Waltham, MA, USA) with previously reported primers and probes specific to RSV and its subtypes (Fry et al. 2010, Aamir et al. 2013)

For sequencing, 23 RSV-positive samples were randomly selected based on a Ct value of <25 from the years 2021-2022. RNA extraction for sequencing was carried out using the Zymo Quick-RNA Viral Kit (Zymo Research, Orange, CA, USA). cDNA synthesis and sequencing library preparation were performed following the ARTIC v4 protocol, with SARS-CoV-2 primers replaced by primers for RSV-A or RSV-B designed in this study. Sequencing was conducted on the Illumina NextSeq 2000 platform or the iseq100 platform (151-bp PE). The resulting Fastq files were processed using the CZID Viral Consensus Genome Pipeline. All generated RSV sequences had at least 90% of their genomes covered. All sequences achieved >1000x coverage except for one (792x).

### Sequencing Strategy – GSTT

Patients with suspected viral respiratory tract infection had routine nasopharyngeal swabs collected for analysis using the AusDiagnostics Universal 24-well Respiratory Pathogen Panel. A selection of RSV-positive samples from October 2022 to March 2023 retrieved for sequencing. Collection of surplus samples and linked clinical data was approved by South Central—Hampshire B REC (20/SC/0310). Amplified cDNA was generated using a modified version Artic Lo-Cost V3 protocol for SARS-CoV-2 sequencing (https://www.protocols.io/view/ncov-2019-sequencing-protocol-v3-locost-bp2l6n26rgqe/v3), replacing primers for SARS-CoV-2 with primers for RSV-A or RSV-B. The sequencing library was prepared with a rapid library preparation kit (SQK-RBK110.96) using a × 1 pooled barcoded sample volume to Ampure XP bead volume ratio and sequenced on an R9.4.1 flow cell using a GridION Mk1. A no template control was included on each sequencing run. A further non-RSV in-run control was introduced at the library preparation stage, with lambda phage DNA used to provide a measure of barcode misassignment.

Consensus sequences were generated using artic field-bioinformatics v1.2.1 (https://github.com/artic-network/fieldbioinformatics) with a 20x depth threshold and using medaka for variant calling.

Extracted samples were sent to the Royal Infirmary of Edinburgh to be resequenced using the Illumina nextseq 2000 platform as described above. The Illumina sequences are reported here as those from GSTT.

### Sequencing Strategy – MDH

RNA was extracted from samples that had been collected in 2019 and 2020 for primer testing and genome assembly. Separate amplification reactions using RSV-A and RSV-B primer sets were done on all samples before library preparation and sequencing on two separate platforms: the Oxford Nanopore GridION using R9.4.1 flow cells and the Illumina MiSeq using 500 cycle v2 kit (Illumina). Quality control, assembly, and consensus genome sequence generation were conducted using the nf-core/viralrecon v2.6.0 pipeline (https://nf-co.re/viralrecon/2.6.0/). We provided the custom primer schemes, Nextclade lineage designations, and reference sequences (hRSV/A/England/RS20000581/2020|EPI_ISL_1647421|2020-02-04 and hRSV/B/England/RE20000104/2020|EPI_ISL_1647600|2020-01-30). Genome assembly was performed within this pipeline using Medaka 1.6.1 and the r941_min_sup_g507 Medaka model (for ONT reads) using a minimum depth of 20x coverage and no greater than 10% missing data. This resulted in a dataset of 50 sequences, of which 25 were RSV-A and 25 were RSV-B.

We compared the Oxford Nanopore and Illumina assemblies and other than minor differences in missing sites, no differences such as SNP or indel calls were detected. As such the data we are reporting here are all ONT generated.

### Sequencing Strategy – NVRL

The considered Irish samples were collected between 2022 and 2023. Viral RNA was extracted using the Roche MagNA Pure 96 DNA and Viral NA Small Volume kit according to the manufacturer’s instructions and eluates stored at –80C prior to use. Samples were tested using the NxTAG Respiratory Pathogen Panel + SARS-CoV-2 assay (Luminex), which identifies both RSV-A and –B subgroups. Specimens with a viral titre yielding results of MDD > 300 (NxTAG) were considered eligible for sequence analysis. Samples for RSV-A (n=43) and RSV-B (n=46) were chosen to be run in a single 96-plate including negative samples for each group in the amplification stage and a negative sample for the sequencing stage. The genomes were amplified following the ARTIC protocol and primers provided in this study, and amplicons were deep sequenced by using the GridION nanopore device (Oxford Nanopore Technologies, https://nanoporetech.com) on genomic DNA by using the Ligation Sequencing Kit (Oxford Nanopore Technologies) and Native Barcodes (Oxford Nanopore Technologies).

### Sequencing Strategy – UoN

Clinical specimens were originally submitted for routine diagnosis from sputum, nasopharyngeal aspirate, or throat swab samples from patients with suspected respiratory virus infections in primary and secondary care units in the Nottingham University Hospitals Trust catchment area. Specimens were extracted in the microbiology lab in Queen’s Medical Centre (QMC), Nottingham, UK, following the procedure described by Chellapuri et. al, 2017 and nucleic acids screened using in house methods prior to February 2016 and the commercial AusDiagnostics screening panel from March 2016 to date. RSV-positive specimens selected for whole genome sequencing were submitted between 5th December 2014 and 18th May 2023 and had Ct values of between 16.4 and 30.6 (in house outputs) or between 1,300 and 14,006,853 RSV genome copies/10 l total RNA (AusDiagnostics outputs). cDNA was synthesised by random hexamer priming as previously described (Chellapuri et. al, 2017) and PCR-amplified using previously described RSV genotype-specific primers and thermocycling conditions (https://virological.org/t/preliminary-results-from-two-novel-artic-style-amplicon-based-sequencing-approaches-for-rsv-a-and-rsv-b/918), except with 45 cycles of amplification used as standard. PCR products from each sample and primer pool were visualised using gel electrophoresis prior to pools being combined for each sample, quantified using the QubitTM dsDNA Quantification Assay Kit and diluted to 100 ng in 7.5 l with nuclease-free H2O.

Sequencing was performed on Oxford Nanopore PromethION sequencers and processed using the ARTIC amplicon sequencing protocol. Sequences were aligned using MAFFT (Katoh et al, 2017).

Use of residual diagnostic nucleic acids and associated anonymized patient information was covered by ethical approval granted to the Nottingham Health Science Biobank Research Tissue Bank, by the Northwest - Greater Manchester Central Research Ethics Committee, UK, reference 15/NW/0685.

### Sequencing Strategy – USAF

Viral RNA was extracted from 200 μl of Nasopharyngeal (NP) swabs using the NucliSENS EMAG, EasyMAG, or the KingFisher Flex System and reagents (Biomerieux or MagMAX™ Nucleic Acid Isolation Kit), following the manufacturer’s instructions. Specimens were tested on the Luminex NxTAG Respiratory Pathogen Panel and remnant clinical specimens were subsequently deidentified and chosen for sequencing to test the amplicon panel. Specimen extract was also tested using RTPCR to establish Ct values according to Todd et al. 2021 (Todd et al, 2021). Post ARTIC amplification for RSV, sequencing libraries were then prepared using the Illumina Nextera XT DNA Library Preparation Kit. Libraries were then sequenced on an Illumina NextSeq 550 device using a mid-output 300 cycle flow cell. Raw reads were processed using a variation of the Iterative Refinement Meta Assembler (Shepard et al, 2016) and the RSV0 module of IRMA-RSV (https://github.com/JianiC/IRMA-RSV). These specimens were selected and processed under a Not Human Subjects Research determination from the USAF School of Aerospace Medicine IRB (FWR20200110N).

### Sequencing Strategy – USFQ

RSV A and B positive samples using the Respiratory qPCR panels for GeneExpert were included in the study USFQ-CEISH 2020-022M after informed consent was signed by patients. Viral RNA from samples was extracted using Qiagen RNeasy Mini Kit (QIAGEN, Germany) following the manufacturer’s protocols. We used a modified version of the Artic Lo-Cost V3 protocol for SARS-CoV-2 sequencing (https://www.protocols.io/view/ncov-2019-sequencing-protocol-v3-locost-bp2l6n26rgqe/v3) to generate amplified cDNA. Instead of SARS-CoV-2 primers, we used primers for RSV-A or RSV-B. The sequencing library was prepared using the ligation sequencing kit (SQK-LSK109 with EXP-NBD196) with a 1x pooled barcoded sample volume to Ampure XP bead volume ratio. Each sequencing run included a template control. We conducted the sequencing on an R9.4.1 flow cell using a GridION Mk1 with a High accuracy (HAC) basecalling model. We generated consensus sequences using artic field-bioinformatics v1.2.1 (https://github.com/artic-network/fieldbioinformatics) with a 20x depth threshold. Consensus sequences were uploaded to GISAID with the accession numbers EPI_ISL_18386604 and EPI_ISL_18386605.

### Sequencing Strategy – UW

Remnant clinical specimens tested at the University of Washington Virology Lab were chosen for sequencing. After extraction, sscDNA was prepared from samples using the COVIDseq kit (Illumina), according to manufacturer’s instructions. Amplicon PCR was performed using COVIDseq PCR mix reagents and the ARTIC RSV amplicon panel primer pool 1 and pool 2 sets. Thermocycler conditions were as follows: 98°C for 3 minutes, 34 cycles of 98°C for 15 seconds and 63°C for 5 minutes, with a 4°C hold. The lid temperature was set to 105°C. Following amplicon PCR, pool 1 and pool 2 were combined and library preparation was performed using the COVIDseq kit according to manufacturer’s instructions. Samples were pooled in equal volumes and a 0.8x cleanup was performed using Illumina Tune Beads. Sequencing of library pools was performed on Illumina NextSeq 2000 instruments with a 2×150 read format with on-board denaturation and targeting 1 million reads per sample.

Raw reads were assembled into consensus genomes using a custom bioinformatic pipeline (https://github.com/greninger-lab/rsv_ampseq). Briefly, raw reads underwent adapter and quality trimming using fastp v0.23.2. Trimmed reads were then aligned to a multi-sequence fasta file containing references for both RSV-A (GISAID accession EPI_ISL_1647421) and RSV-B (GISAID accession EPI_ISL_1647600) using BBMap v39.01. Subsequently, the trimmed reads were remapped to either the RSV-A or the RSV-B reference, based on whichever exhibited more mapped reads from the initial mapping. Amplicon primers were trimmed from the resulting BAM file using iVar trim v1.4. Variant calling was performed used iVAR variants v1.4 and bcftools v1.17 at a minimum threshold of 0.01, minimum base quality of 20, and minimum depth of 10. A consensus sequence was generated using iVar consensus v1.4 at a minimum threshold of 0.6, a minimum base quality of 20, and a minimum depth of 10, below which ‘N’ bases were called.

The use of remnant clinical specimens was approved by the University of Washington Institutional Review Board (protocol STUDY00000408).

## Supporting information

GISAID acknowledgement table

## Data Availability

Each sequencing site retains primary control over the sequences produced by their team in this study. The sequences produced by the RIE team that met the >90% genome completeness threshold to be included in the phylogenetic analysis have been deposited to GenBank (BioProject ID: PRJNA1153476). Primer scheme sequences are publicly available at https://github.com/artic-network/artic-rsv. All other data can be made available upon request.

## Acknowledgements

The authors would like to thank all the members of the Viral Sequencing Service for their tireless work at the Royal Infirmary of Edinburgh. We would like to thank Nick Loman for initially developing “fieldbioinformatics” and Joshua Quick for “primal scheme”. We also extend our thanks to members of the wider scientific community that have used these primers since the instigation of this study. In particular, thank you to those who hoped to be a part of this study but due to logistical issues could not get primers in time. Furthermore, we would like to thank Dr. Barnabas King for their contribution to this manuscript. Finally, we gratefully acknowledge all data contributors, i.e., the Authors and their Originating laboratories responsible for obtaining the specimens, and their Submitting laboratories for generating the genetic sequence and metadata and sharing via the GISAID Initiative, on which the background RSV data set for the phylogenetic analysis was based.

