## Supplementary Figure 1 for "ARTIC RSV amplicon sequencing reveals global RSV genotype dynamics"

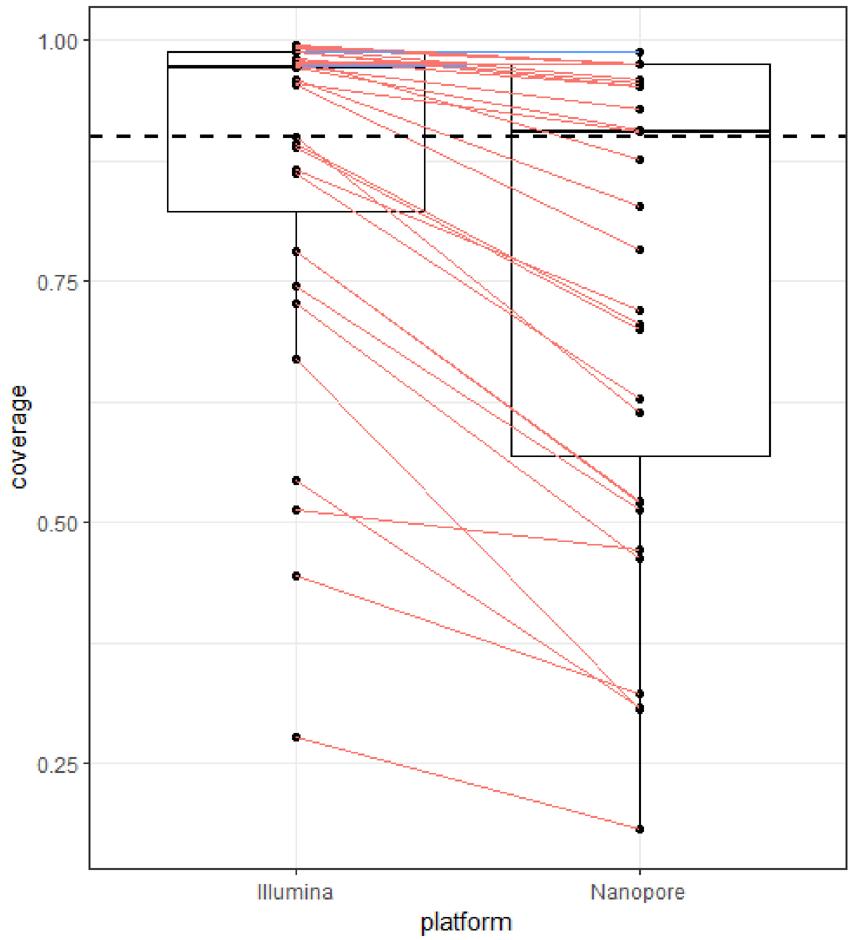


**Supplementary Figure 1:** Comparison of genome coverage of samples that were sequenced on both Illumina and Nanopore platforms with lines connecting the same sample on each platform (n = 35). Red lines connecting dots shows samples that had lower genome completeness when sequenced with Nanopore and blue lines denote samples with higher genome completeness using Nanopore. Dashed line indicates 90% genome completeness.
