## Supplementary Figure 2 for "ARTIC RSV amplicon sequencing reveals global RSV genotype dynamics"

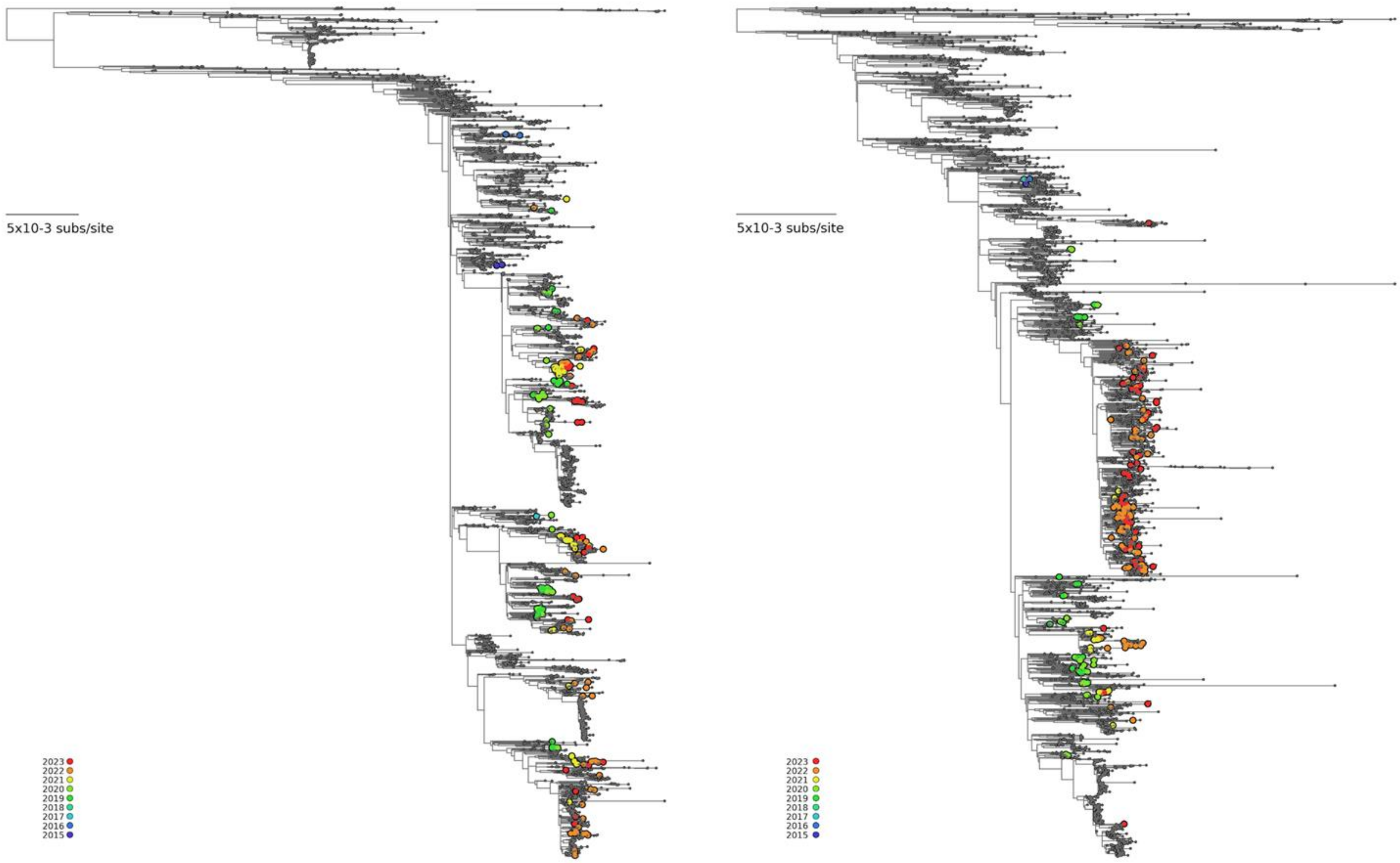


**Supplementary Figure 1:** Mid-point rooted maximum likelihood trees of all **A)** RSV A and **B)** RSV B samples sequenced for this study, coloured by the year of collection.
