## Supplementary material for "ARTIC RSV amplicon sequencing reveals global RSV genotype dynamics": GISAID acknowledgement table: gisaid_rsv_acknowledgement_table_2023_12_08_10_A.pdf

We gratefully acknowledge the following Authors from the Originating laboratories responsible for obtaining the specimens, as well as the Submitting laboratories where the genome data were generated and shared via GISAID, on which this research is based.

All Submitters of data may be contacted directly via www.gisaid.org

Authors are sorted alphabetically.

| Accession ID | Originating Laboratory | Submitting Laboratory | Authors |
|---|---|---|---|
| EPI_ISL_1074028, EPI_ISL_1074029, EPI_ISL_1074031, EPI_ISL_1074034, EPI_ISL_1074035, EPI_ISL_1074036, EPI_ISL_1074037, EPI_ISL_1074038, EPI_ISL_1074039, EPI_ISL_1074040, EPI_ISL_1074041, EPI_ISL_1074042, EPI_ISL_1074043, EPI_ISL_1074044, EPI_ISL_1074045, EPI_ISL_1074046, EPI_ISL_1074047, EPI_ISL_1074048, EPI_ISL_1074049, EPI_ISL_1074050, EPI_ISL_1074051, EPI_ISL_1074052, EPI_ISL_1074053, EPI_ISL_1074054, EPI_ISL_1074055, EPI_ISL_1074056, EPI_ISL_1074057, EPI_ISL_1074058, EPI_ISL_1074059, EPI_ISL_1074060, EPI_ISL_1074061, EPI_ISL_1074062, EPI_ISL_1074063, EPI_ISL_1074064, EPI_ISL_1074065, EPI_ISL_1074066, EPI_ISL_1074067, EPI_ISL_1074068, EPI_ISL_1074088, EPI_ISL_1074089, EPI_ISL_1074090, EPI_ISL_1074094, EPI_ISL_1074098, EPI_ISL_1074099, EPI_ISL_1074100, EPI_ISL_1074103, EPI_ISL_1074104, EPI_ISL_1074105, EPI_ISL_1074106, EPI_ISL_1074108, EPI_ISL_1074111, EPI_ISL_1074112, EPI_ISL_1074113, EPI_ISL_1074114, EPI_ISL_1074115, EPI_ISL_1074118, EPI_ISL_1074120, EPI_ISL_1074124, EPI_ISL_1074125, EPI_ISL_1074126, EPI_ISL_1074127, EPI_ISL_1074135, EPI_ISL_1074136, EPI_ISL_1074137, EPI_ISL_1074138, EPI_ISL_1074139, EPI_ISL_1074140, EPI_ISL_1074141, EPI_ISL_1074145, EPI_ISL_1074146, EPI_ISL_1074149, EPI_ISL_1074150, EPI_ISL_1074151, EPI_ISL_1074152, EPI_ISL_1074153, EPI_ISL_1074154, EPI_ISL_1074156, EPI_ISL_1074157, EPI_ISL_1074158, EPI_ISL_1074159, EPI_ISL_1074160, EPI_ISL_1074161, EPI_ISL_1074164, EPI_ISL_1074165, EPI_ISL_1074166, EPI_ISL_1074167, EPI_ISL_1074168, EPI_ISL_1074169, EPI_ISL_1074170, EPI_ISL_1074171, EPI_ISL_1074172, EPI_ISL_1074173, EPI_ISL_1074183, EPI_ISL_1074184, EPI_ISL_1074187, EPI_ISL_1074188, EPI_ISL_1074190, EPI_ISL_1074191, EPI_ISL_1074194, EPI_ISL_1074195, EPI_ISL_1074208, EPI_ISL_1074209, EPI_ISL_1074210, EPI_ISL_1074211, EPI_ISL_1074212, EPI_ISL_1074213, EPI_ISL_1074214, EPI_ISL_1074227, EPI_ISL_1074228, EPI_ISL_1074233, EPI_ISL_1074234, EPI_ISL_1074240, EPI_ISL_1074243, EPI_ISL_1074244, EPI_ISL_1074245, EPI_ISL_1074246, EPI_ISL_1074248, EPI_ISL_1074249, EPI_ISL_1074250, EPI_ISL_1074253, EPI_ISL_1074255, EPI_ISL_1074256, EPI_ISL_1074257, EPI_ISL_1074259, EPI_ISL_1074260, EPI_ISL_1074262, EPI_ISL_1074267, EPI_ISL_1074270, EPI_ISL_1074271, EPI_ISL_1074272, EPI_ISL_1074273, EPI_ISL_1074274, EPI_ISL_1074275, EPI_ISL_1074276, EPI_ISL_1074277, EPI_ISL_1074278, EPI_ISL_1074279, EPI_ISL_1074280, EPI_ISL_1074281, EPI_ISL_1074282 | | | |
| see above | Virology Laboratory, Ricardo Gutiérrez Children's Hospital | Vanderbilt University Medical Center / Virology Laboratory, Ricardo Gutiérrez Children's Hospital | Goya, Stephanie; Lucion, Maria Florencia; Juarez, Maria del Valle; Shilts, Meghan; Gentile, Angela; Mistchenko, Alicia S.; Das, Suman# & Viegas, Mariana# (#contributed equally to this study) |
| EPI_ISL_10914989 | VIC, Victorian Infectious Diseases Reference Laboratory | WHO Collaborating Centre for Reference and Research on Influenza | Xiaomin Dong, Annette Alafaci,Yi-Mo Deng, Naomi Komadina, Ammar Aziz |
| EPI_ISL_10954007, EPI_ISL_10954008 | Respiratory Virus Unit, Specialised Microbiology and Laboratories Directorate, UK Health Security Agency | Respiratory Virus Unit, Specialised Microbiology and Laboratories Directorate, UK Health Security Agency | Zambon M, Talts T, Ellis J, Miah S, Platt S |
| EPI_ISL_11019782, EPI_ISL_11019880 | NAKORNPING HOSPITAL | National Institute of Health, Department of Medical Sciences, Ministry of Public Health, Thailand | Siripaporn Phuygun; Pakorn Piromtong; Thanutsapa Thanadachakul; Natchaya Khiadsang; Sunthareeya Waicharoen; Malinee Chittaganpitch; Pilailuk Akkapaiboon Okada |
| EPI_ISL_11055715 | Rahima Moosa Mother and child Hospital | National Institute for Communicable Diseases of the National Health Laboratory Service | Amoako DG, Everatt J, Kekana D, Scheepers C, Mohale T, Ntuli N, Mahlangu B, Mnguni A, Ismail A, Bhiman JN, Wolter N |
| EPI_ISL_11055716, EPI_ISL_11055717 | Agincourt clinic | National Institute for Communicable Diseases of the National Health Laboratory Service | Amoako DG, Everatt J, Kekana D, Scheepers C, Mohale T, Ntuli N, Mahlangu B, Mnguni A, Ismail A, Bhiman JN, Wolter N |
| EPI_ISL_11055718 | Edendale Gateway clinic | National Institute for Communicable Diseases of the National Health Laboratory Service | Amoako DG, Everatt J, Kekana D, Scheepers C, Mohale T, Ntuli N, Mahlangu B, Mnguni A, Ismail A, Bhiman JN, Wolter N |
| EPI_ISL_11055719, EPI_ISL_11055720 | Edendale hospital | National Institute for Communicable Diseases of the National Health Laboratory Service | Amoako DG, Everatt J, Kekana D, Scheepers C, Mohale T, Ntuli N, Mahlangu B, Mnguni A, Ismail A, Bhiman JN, Wolter N |
| EPI_ISL_11055721 | Edendale Gateway clinic | National Institute for Communicable Diseases of the National Health Laboratory Service | Amoako DG, Everatt J, Kekana D, Scheepers C, Mohale T, Ntuli N, Mahlangu B, Mnguni A, Ismail A, Bhiman JN, Wolter N |
| EPI_ISL_11055722, EPI_ISL_11055723 | Edendale hospital | National Institute for Communicable Diseases of the National Health Laboratory Service | Amoako DG, Everatt J, Kekana D, Scheepers C, Mohale T, Ntuli N, Mahlangu B, Mnguni A, Ismail A, Bhiman JN, Wolter N |
| EPI_ISL_11055724, EPI_ISL_11055725, EPI_ISL_11055726 | Edendale Gateway clinic | National Institute for Communicable Diseases of the National Health Laboratory Service | Amoako DG, Everatt J, Kekana D, Scheepers C, Mohale T, Ntuli N, Mahlangu B, Mnguni A, Ismail A, Bhiman JN, Wolter N |
| EPI_ISL_11055727, EPI_ISL_11055728, EPI_ISL_11055729, EPI_ISL_11055730, EPI_ISL_11055731 | Edendale hospital | National Institute for Communicable Diseases of the National Health Laboratory Service | Amoako DG, Everatt J, Kekana D, Scheepers C, Mohale T, Ntuli N, Mahlangu B, Mnguni A, Ismail A, Bhiman JN, Wolter N |
| EPI_ISL_11055732, EPI_ISL_11055734 | Edendale Gateway clinic | National Institute for Communicable Diseases of the National Health Laboratory Service | Amoako DG, Everatt J, Kekana D, Scheepers C, Mohale T, Ntuli N, Mahlangu B, Mnguni A, Ismail A, Bhiman JN, Wolter N |
| EPI_ISL_11055735 | Matikwana hospital | National Institute for Communicable Diseases of the National Health Laboratory Service | Amoako DG, Everatt J, Kekana D, Scheepers C, Mohale T, Ntuli N, Mahlangu B, Mnguni A, Ismail A, Bhiman JN, Wolter N |
| EPI_ISL_11055736 | Eastridge clinic | National Institute for Communicable Diseases of the National Health Laboratory Service | Amoako DG, Everatt J, Kekana D, Scheepers C, Mohale T, Ntuli N, Mahlangu B, Mnguni A, Ismail A, Bhiman JN, Wolter N |
| EPI_ISL_11055737 | Agincourt clinic | National Institute for Communicable Diseases of the National Health Laboratory Service | Amoako DG, Everatt J, Kekana D, Scheepers C, Mohale T, Ntuli N, Mahlangu B, Mnguni A, Ismail A, Bhiman JN, Wolter N |
| EPI_ISL_11055738 | Mapulaneng hospital | National Institute for Communicable Diseases of the National Health Laboratory Service | Amoako DG, Everatt J, Kekana D, Scheepers C, Mohale T, Ntuli N, Mahlangu B, Mnguni A, Ismail A, Bhiman JN, Wolter N |
| EPI_ISL_11055740, EPI_ISL_11055741, EPI_ISL_11055742, EPI_ISL_11055743 | Red cross childrens hospital | National Institute for Communicable Diseases of the National Health Laboratory Service | Amoako DG, Everatt J, Kekana D, Scheepers C, Mohale T, Ntuli N, Mahlangu B, Mnguni A, Ismail A, Bhiman JN, Wolter N |
| EPI_ISL_11055744 | Mitchell's plain distric hospital | National Institute for Communicable Diseases of the National Health Laboratory Service | Amoako DG, Everatt J, Kekana D, Scheepers C, Mohale T, Ntuli N, Mahlangu B, Mnguni A, Ismail A, Bhiman JN, Wolter N |
| EPI_ISL_11055745, EPI_ISL_11055746, EPI_ISL_11055747 | Red cross childrens hospital | National Institute for Communicable Diseases of the National Health Laboratory Service | Amoako DG, Everatt J, Kekana D, Scheepers C, Mohale T, Ntuli N, Mahlangu B, Mnguni A, Ismail A, Bhiman JN, Wolter N |
| EPI_ISL_11055748 | Eastridge clinic | National Institute for Communicable Diseases of the National Health Laboratory Service | Amoako DG, Everatt J, Kekana D, Scheepers C, Mohale T, Ntuli N, Mahlangu B, Mnguni A, Ismail A, Bhiman JN, Wolter N |
| EPI_ISL_11055749 | Red cross childrens hospital | National Institute for Communicable Diseases of the National Health Laboratory Service | Amoako DG, Everatt J, Kekana D, Scheepers C, Mohale T, Ntuli N, Mahlangu B, Mnguni A, Ismail A, Bhiman JN, Wolter N |
| EPI_ISL_11055750 | Eastridge clinic | National Institute for Communicable Diseases of the National Health Laboratory Service | Amoako DG, Everatt J, Kekana D, Scheepers C, Mohale T, Ntuli N, Mahlangu B, Mnguni A, Ismail A, Bhiman JN, Wolter N |
| EPI_ISL_11055751 | Red cross childrens hospital | National Institute for Communicable Diseases of the National Health Laboratory Service | Amoako DG, Everatt J, Kekana D, Scheepers C, Mohale T, Ntuli N, Mahlangu B, Mnguni A, Ismail A, Bhiman JN, Wolter N |
| EPI_ISL_11055752, EPI_ISL_11055753, EPI_ISL_11055754, EPI_ISL_11055755, EPI_ISL_11055756, EPI_ISL_11055757, EPI_ISL_11055759, EPI_ISL_11055760 | Mitchell's plain distric hospital | National Institute for Communicable Diseases of the National Health Laboratory Service | Amoako DG, Everatt J, Kekana D, Scheepers C, Mohale T, Ntuli N, Mahlangu B, Mnguni A, Ismail A, Bhiman JN, Wolter N |
| EPI_ISL_11055762, EPI_ISL_11055763, EPI_ISL_11055765, EPI_ISL_11055766 | Red cross childrens hospital | National Institute for Communicable Diseases of the National Health Laboratory Service | Amoako DG, Everatt J, Kekana D, Scheepers C, Mohale T, Ntuli N, Mahlangu B, Mnguni A, Ismail A, Bhiman JN, Wolter N |
| EPI_ISL_11055768 | Edendale Gateway clinic | National Institute for Communicable Diseases of the National Health Laboratory Service | Amoako DG, Everatt J, Kekana D, Scheepers C, Mohale T, Ntuli N, Mahlangu B, Mnguni A, Ismail A, Bhiman JN, Wolter N |
| EPI_ISL_11428295, EPI_ISL_11428299, EPI_ISL_11428301, EPI_ISL_11428309, EPI_ISL_11428318, EPI_ISL_11428328 | Respiratory Virus Unit, Reference Services, UK Health Security Agency | Reference Services, UK Health Security Agency | Zambon M, Talts T, Ellis J, Miah S, Platt S |
| EPI_ISL_11817019, EPI_ISL_11817020, EPI_ISL_11817021, EPI_ISL_11817023, EPI_ISL_11817024, EPI_ISL_11817025, EPI_ISL_11817026, EPI_ISL_11817027, EPI_ISL_11817028, EPI_ISL_11817030, EPI_ISL_11817032, EPI_ISL_11817036, EPI_ISL_11817038, EPI_ISL_11817039, EPI_ISL_11817041, EPI_ISL_11817042, EPI_ISL_11817043, EPI_ISL_11817045, EPI_ISL_11817046, EPI_ISL_11817049, EPI_ISL_11817051, EPI_ISL_11817053, EPI_ISL_11817056, EPI_ISL_11817061, EPI_ISL_11817062, EPI_ISL_11817065, EPI_ISL_11817066, EPI_ISL_11817069, EPI_ISL_11817070, EPI_ISL_11817071, EPI_ISL_11817072, EPI_ISL_11817075, EPI_ISL_11817076, EPI_ISL_11817078, EPI_ISL_11817080, EPI_ISL_11817082, EPI_ISL_11817083, EPI_ISL_11817086 | | | |
| see above | QLD, Royal Brisbane and Woman's Hospital | WHO Collaborating Centre for Reference and Research on Influenza | Xiaomin Dong, Yi-Mo Deng, Ammar Aziz, Naomi Komadina |
| EPI_ISL_11817088, EPI_ISL_11817090, EPI_ISL_11817091, EPI_ISL_11817092, EPI_ISL_11817094, EPI_ISL_11817095, EPI_ISL_11817096, EPI_ISL_11817097, EPI_ISL_11817099, EPI_ISL_11817100, EPI_ISL_11817101, EPI_ISL_11817102, EPI_ISL_11817103, EPI_ISL_11817104, EPI_ISL_11817105, EPI_ISL_11817106, EPI_ISL_11817107, EPI_ISL_11817108, EPI_ISL_11817109, EPI_ISL_11823065 | | | |
| see above | VIC, Royal Children's Hospital | WHO Collaborating Centre for Reference and Research on Influenza | Xiaomin Dong, Annette Alafaci,Yi-Mo Deng, Ammar Aziz, Naomi Komadina |
| EPI_ISL_12529634, EPI_ISL_12529635, EPI_ISL_12529637, EPI_ISL_12529638 | Rahima Moosa Mother and child Hospital | National Institute for Communicable Diseases of the National Health Laboratory Service | Amoako DG, Everatt J, Kekana D, Scheepers C, Mohale T, Ntuli N, Mahlangu B, Mnguni A, Ismail A, Bhiman JN, Wolter N |
| EPI_ISL_12529639, EPI_ISL_12529640 | Mapulaneng hospital | National Institute for Communicable Diseases of the National Health Laboratory Service | Amoako DG, Everatt J, Kekana D, Scheepers C, Mohale T, Ntuli N, Mahlangu B, Mnguni A, Ismail A, Bhiman JN, Wolter N |
| EPI_ISL_12529641 | Agincourt clinic | National Institute for Communicable Diseases of the National Health Laboratory Service | Amoako DG, Everatt J, Kekana D, Scheepers C, Mohale T, Ntuli N, Mahlangu B, Mnguni A, Ismail A, Bhiman JN, Wolter N |
| EPI_ISL_12529642 | Rahima Moosa Mother and child Hospital | National Institute for Communicable Diseases of the National Health Laboratory Service | Amoako DG, Everatt J, Kekana D, Scheepers C, Mohale T, Ntuli N, Mahlangu B, Mnguni A, Ismail A, Bhiman JN, Wolter N |
| EPI_ISL_12529643 | Red cross childrens hospital | National Institute for Communicable Diseases of the National Health Laboratory Service | Amoako DG, Everatt J, Kekana D, Scheepers C, Mohale T, Ntuli N, Mahlangu B, Mnguni A, Ismail A, Bhiman JN, Wolter N |

| | | | |
|---|---|---|---|
| EPI_ISL_12529644 | Rahima Moosa Mother and child Hospital | National Institute for Communicable Diseases of the National Health Laboratory Service | Amoako DG, Everatt J, Kekana D, Scheepers C, Mohale T, Ntuli N, Mahlangu B, Mnguni A, Ismail A, Bhiman JN, Wolter N |
| EPI_ISL_12529645, EPI_ISL_12529646 | Tintswalo Hospital | National Institute for Communicable Diseases of the National Health Laboratory Service | Amoako DG, Everatt J, Kekana D, Scheepers C, Mohale T, Ntuli N, Mahlangu B, Mnguni A, Ismail A, Bhiman JN, Wolter N |
| EPI_ISL_12529647 | Matikwana hospital | National Institute for Communicable Diseases of the National Health Laboratory Service | Amoako DG, Everatt J, Kekana D, Scheepers C, Mohale T, Ntuli N, Mahlangu B, Mnguni A, Ismail A, Bhiman JN, Wolter N |
| EPI_ISL_12529648 | Mitchell's plain distric hospital | National Institute for Communicable Diseases of the National Health Laboratory Service | Amoako DG, Everatt J, Kekana D, Scheepers C, Mohale T, Ntuli N, Mahlangu B, Mnguni A, Ismail A, Bhiman JN, Wolter N |
| EPI_ISL_12544918, EPI_ISL_12544919 | National Influenza Center, National Institute of Hygiene and Epidemiology (NIHE) | National Institute of Hygiene and Epidemiology (NIHE) National Influenza Centre, Virology Department | Ung Thi Hong Trang, Hoang Vu Mai Phuong, Nguyen Huy Hoang, Nguyen Le Khanh Hang, Le Thi Thanh, Nguyen Vu Son, Vuong Duc Cuong, Pham Thi Hien, Tran Thu Huong, Nguyen Co Thach, Nguyen Phuong Anh, Le Quynh Mai |
| EPI_ISL_12870543, EPI_ISL_12870544, EPI_ISL_12870545 | Agincourt clinic | National Institute for Communicable Diseases of the National Health Laboratory Service | Amoako DG, Everatt J, Kekana D, Scheepers C, Mohale T, Ntuli N, Mahlangu B, Mnguni A, Ismail A, Bhiman JN, Wolter N |
| EPI_ISL_12870547 | Rahima Moosa Mother and child Hospital | National Institute for Communicable Diseases of the National Health Laboratory Service | Amoako DG, Everatt J, Kekana D, Scheepers C, Mohale T, Ntuli N, Mahlangu B, Mnguni A, Ismail A, Bhiman JN, Wolter N |
| EPI_ISL_12870552, EPI_ISL_12870553, EPI_ISL_12870554 | Edendale hospital | National Institute for Communicable Diseases of the National Health Laboratory Service | Amoako DG, Everatt J, Kekana D, Scheepers C, Mohale T, Ntuli N, Mahlangu B, Mnguni A, Ismail A, Bhiman JN, Wolter N |
| EPI_ISL_12970401, EPI_ISL_12970403, EPI_ISL_12970405, EPI_ISL_12970407, EPI_ISL_12970408, EPI_ISL_12970409, EPI_ISL_12970411, EPI_ISL_12970412, EPI_ISL_12970413, EPI_ISL_12970414, EPI_ISL_12970415, EPI_ISL_12970416, EPI_ISL_12970417, EPI_ISL_12970418, EPI_ISL_12970419, EPI_ISL_12970420, EPI_ISL_12970421, EPI_ISL_12970422, EPI_ISL_12970423, EPI_ISL_12970425, EPI_ISL_12970426, EPI_ISL_12970427, EPI_ISL_12970428 | | | |
| see above | Department of Virology, Research Institute for Tropical Medicine | RVDT/RVD/DVD/CDC | Lijuan Wang, Mayan U. Lumandas, Vina Lea Arguelles, Jonjee Morin, Roman Tatusov and Everardo Vega |
| EPI_ISL_13297850, EPI_ISL_13297987 | Sciensano, Department of infectious diseases, Laboratory of viral diseases | Sciensano, Department of infectious diseases, Laboratory of viral diseases | François E. Dufrasne, Sarah Denayer, Steven Van Gucht, Cyril Barbezange |
| EPI_ISL_14018007, EPI_ISL_14018008, EPI_ISL_14039044, EPI_ISL_14039045, EPI_ISL_14039046, EPI_ISL_14039047, EPI_ISL_14039048, EPI_ISL_14039049, EPI_ISL_14039050 | Infectious Diseases Research Collaboration (IDRC), Uganda | Chan Zuckerberg Biohub, USA | Greenhouse,Bryan; Kamya,Moses; Mwakibete,Lusajo; Rek,John; Rodriguez-Barraquer,Isabel; Ssewanyana,Isaac; Takahashi,Saki; Tato,Cristina |
| EPI_ISL_14084089, EPI_ISL_14084090, EPI_ISL_14084091 | Complexo Hospitalario Universitario de Vigo. Microbiology Department | Complexo Hospitalario Universitario de Vigo. Microbiology Department | Daviña, Carlos; Pizcueta, Juan; Perez, Sonia |
| EPI_ISL_14769848 | Rahima Moosa | National Institute for Communicable Diseases of the National Health Laboratory Service | Amoako DG, Everatt J, Kekana D, Mahlangu B, Stock N, Ntuli N, Mnguni A, Motsatsi G, Nzimande A, Ismail A, Bhiman JN, Wolter N |
| EPI_ISL_14769850 | Red Cross | National Institute for Communicable Diseases of the National Health Laboratory Service | Amoako DG, Everatt J, Kekana D, Mahlangu B, Stock N, Ntuli N, Mnguni A, Motsatsi G, Nzimande A, Ismail A, Bhiman JN, Wolter N |
| EPI_ISL_15004433, EPI_ISL_15004435 | National Institute of Hygiene, virology departement, National Influenza Center | Centers for Disease Control and Prevention, Coronavirus and Other Respiratory Viruses Division | Megha Aggarwal, Lijuan Wang, Ji In Park, Amanda Smith and Everardo Vega |
| EPI_ISL_15055310, EPI_ISL_15055311, EPI_ISL_15055312, EPI_ISL_15055313, EPI_ISL_15055314, EPI_ISL_15055315, EPI_ISL_15055316, EPI_ISL_15055317, EPI_ISL_15055318, EPI_ISL_15055319, EPI_ISL_15055320, EPI_ISL_15055321, EPI_ISL_15055322, EPI_ISL_15055323, EPI_ISL_15055324, EPI_ISL_15055325, EPI_ISL_15055326, EPI_ISL_15055327, EPI_ISL_15055328, EPI_ISL_15067675, EPI_ISL_15067676, EPI_ISL_15067677, EPI_ISL_15067678, EPI_ISL_15067679, EPI_ISL_15067680, EPI_ISL_15067681, EPI_ISL_15067682, EPI_ISL_15067683, EPI_ISL_15067684, EPI_ISL_15067685, EPI_ISL_15067686, EPI_ISL_15067687, EPI_ISL_15067688, EPI_ISL_15067689, EPI_ISL_15067690, EPI_ISL_15067691, EPI_ISL_15067692, EPI_ISL_15067693 | | | |
| see above | Virology Laboratory, Ricardo Gutiérrez Children's Hospital | Virology Laboratory, Ricardo Gutiérrez Children's Hospital | Acuña, Dolores; Goya, Stephanie; Nabaes Jodar, Mercedes S.; Lucion, M. Florencia; Juárez, María del Valle; Gentile, Ángela; Mistchenko, Alicia S.; Viegas, Mariana |
| EPI_ISL_15120665, EPI_ISL_15120666, EPI_ISL_15120667, EPI_ISL_15120668, EPI_ISL_15120669, EPI_ISL_15120670, EPI_ISL_15120671, EPI_ISL_15120672, EPI_ISL_15120673, EPI_ISL_15120674, EPI_ISL_15120675, EPI_ISL_15120676, EPI_ISL_15120677, EPI_ISL_15120678, EPI_ISL_15120679, EPI_ISL_15120680, EPI_ISL_15120681, EPI_ISL_15120682, EPI_ISL_15120683, EPI_ISL_15120688, EPI_ISL_15120689, EPI_ISL_15120690, EPI_ISL_15120692, EPI_ISL_15120694, EPI_ISL_15120703, EPI_ISL_15120706, EPI_ISL_15120713, EPI_ISL_15120715, EPI_ISL_15120716, EPI_ISL_15120717, EPI_ISL_15120719, EPI_ISL_15120721, EPI_ISL_15120723, EPI_ISL_15120726, EPI_ISL_15120727, EPI_ISL_15120728, EPI_ISL_15120729, EPI_ISL_15120730, EPI_ISL_15120738, EPI_ISL_15120739, EPI_ISL_15120743, EPI_ISL_15120745, EPI_ISL_15120770, EPI_ISL_15120771, EPI_ISL_15120772, EPI_ISL_15120773, EPI_ISL_15120774, EPI_ISL_15120775, EPI_ISL_15120776, EPI_ISL_15120777, EPI_ISL_15120778, EPI_ISL_15120780, EPI_ISL_15120781, EPI_ISL_15120782, EPI_ISL_15120783, EPI_ISL_15120785, EPI_ISL_15120786 | | | |
| see above | National Institute of Hygiene, virology departement, National Influenza Center | Centers for Disease Control and Prevention, Coronavirus and Other Respiratory Viruses Division | Megha Aggarwal, Lijuan Wang, Ji In Park, Amanda Smith and Everardo Vega |
| EPI_ISL_1520381, EPI_ISL_1520382, EPI_ISL_1520383, EPI_ISL_1520384, EPI_ISL_1520385, EPI_ISL_1520386, EPI_ISL_1520387, EPI_ISL_1520388, EPI_ISL_1520389, EPI_ISL_1520390, EPI_ISL_1520391, EPI_ISL_1520392, EPI_ISL_1520393, EPI_ISL_1520394, EPI_ISL_1520395, EPI_ISL_1520396, EPI_ISL_1520397, EPI_ISL_1520398, EPI_ISL_1520399, EPI_ISL_1520400, EPI_ISL_1520401, EPI_ISL_1520402, EPI_ISL_1520403, EPI_ISL_1520404, EPI_ISL_1520405, EPI_ISL_1520406, EPI_ISL_1520407, EPI_ISL_1520408, EPI_ISL_1520409, EPI_ISL_1520410, EPI_ISL_1520411, EPI_ISL_1520412, EPI_ISL_1520413, EPI_ISL_1520414, EPI_ISL_1520415, EPI_ISL_1520416, EPI_ISL_1520417, EPI_ISL_1520418, EPI_ISL_1520419, EPI_ISL_1520420, EPI_ISL_1520421, EPI_ISL_1520422, EPI_ISL_1520423, EPI_ISL_1520424, EPI_ISL_1520425, EPI_ISL_1520426, EPI_ISL_1520427, EPI_ISL_1520428, EPI_ISL_1520429, EPI_ISL_1520430, EPI_ISL_1520431, EPI_ISL_1520432, EPI_ISL_1520433, EPI_ISL_1520434, EPI_ISL_1520435, EPI_ISL_1520436, EPI_ISL_1520437, EPI_ISL_1520438, EPI_ISL_1520439, EPI_ISL_1520440 | | | |
| see above | Respiratory Virus Unit, National Infection Service, Public Health England | National Infection Service, Public Health England | Zambon M, Talts T, Ellis J, Miah S, Platt S |
| EPI_ISL_15421344 | Department of Infectious Diseases in Humans, Viral Diseases Laboratory, Sciensano, Brussels | Department of Infectious Diseases in Humans, Viral Diseases Laboratory, Sciensano, Brussels | François Dufrasne, Sarh Denayer, Steven Van Gucht, Cyril Barbezange |
| EPI_ISL_15728615 | Edendale | National Institute for Communicable Diseases of the National Health Laboratory Service | Everatt J, Kekana D, Mahlangu B, Stock N, Ntozini B, Ntuli N, Mnguni A, Nzimande A, Ismail A, Bhiman JN, Wolter N |
| EPI_ISL_15728616 | Klerksdorp | National Institute for Communicable Diseases of the National Health Laboratory Service | Everatt J, Kekana D, Mahlangu B, Stock N, Ntozini B, Ntuli N, Mnguni A, Nzimande A, Ismail A, Bhiman JN, Wolter N |
| EPI_ISL_15728617 | Mapulaneng | National Institute for Communicable Diseases of the National Health Laboratory Service | Everatt J, Kekana D, Mahlangu B, Stock N, Ntozini B, Ntuli N, Mnguni A, Nzimande A, Ismail A, Bhiman JN, Wolter N |
| EPI_ISL_15728618 | Matikwane | National Institute for Communicable Diseases of the National Health Laboratory Service | Everatt J, Kekana D, Mahlangu B, Stock N, Ntozini B, Ntuli N, Mnguni A, Nzimande A, Ismail A, Bhiman JN, Wolter N |
| EPI_ISL_15750126 | Eurasian Institute of Zoonotic Infections, The Federal Research Center of Fundamental and Translational Medicine | Eurasian Institute of Zoonotic Infections, The Federal Research Center of Fundamental and Translational Medicine | Solomatina,M.V., Kurskaya,O.G., Kabilov,M.R., Tupikin,A.E., Saroyan,T.A., Derko,A.A., Dubovitskiy,N.A., Sobolev,I.A., Shestopalov,A.M., Sharshov,K.A. |
| EPI_ISL_15750187, EPI_ISL_15750188, EPI_ISL_15750189, EPI_ISL_15750190, EPI_ISL_15750191, EPI_ISL_15750196, EPI_ISL_15750197, EPI_ISL_15750198, EPI_ISL_15750199, EPI_ISL_15750200 | Inserm UMR1137, Universite de Paris | Inserm UMR1137, Universite de Paris | Coppee,R., Chenane,H.R., Bridier-Nahmias,A., Tcherakian,C., Catherinot,E., Collin,G., Lebourgeois,S., Visseaux,B., Descamps,D., Vasse,M., Farfour,E., Coppee,R., Chenane,H.R., Bridier-Nahmias,A., Collin,G., Lebourgeois,S., Visseaux,B., Descamps,D. |
| EPI_ISL_15750840, EPI_ISL_15750841, EPI_ISL_15750842, EPI_ISL_15750843, EPI_ISL_15750844 | State Key Laboratory of Pathogen and Biosecurity, Beijing Institute of Microbiology and Epidemiology | State Key Laboratory of Pathogen and Biosecurity, Beijing Institute of Microbiology and Epidemiology | Jia,N., Cao,W.-C., Hu,Y.-L., Ye,R.-Z., Que,T.-C., Xia,L.-Y., Cui,X.-M., Zhang,Y.-W., Jiang,J.-F., Wang,Q.-H., Wang,Q., Jia,N., Cao,W.-C., Hu,Y.-L., Ye,R.-Z., Que,T.-C., Xia,L.-Y., Cui,X.-M., Zhang,Y.-W., Jiang,J.-F., Wang,Q.-H., Wang,Q. |
| EPI_ISL_15752005, EPI_ISL_15752006, EPI_ISL_15752007, EPI_ISL_15752008, EPI_ISL_15752009, EPI_ISL_15752010, EPI_ISL_15752011, EPI_ISL_15752012, EPI_ISL_15752013, EPI_ISL_15752014, EPI_ISL_15752015, EPI_ISL_15752016, EPI_ISL_15752017, EPI_ISL_15752018, EPI_ISL_15752019, EPI_ISL_15752020, EPI_ISL_15752021, EPI_ISL_15752022, EPI_ISL_15752023, EPI_ISL_15752024, EPI_ISL_15752025, EPI_ISL_15752026, EPI_ISL_15752027, EPI_ISL_15752028, EPI_ISL_15752029, EPI_ISL_15752030, EPI_ISL_15752031, EPI_ISL_15752032, EPI_ISL_15752033, EPI_ISL_15752034, EPI_ISL_15752035, EPI_ISL_15752036, EPI_ISL_15752037, EPI_ISL_15752038, EPI_ISL_15752039, EPI_ISL_15752040, EPI_ISL_15752041, EPI_ISL_15752042, EPI_ISL_15752043, EPI_ISL_15752044, EPI_ISL_15752045, EPI_ISL_15752046, EPI_ISL_15752047, EPI_ISL_15752048, EPI_ISL_15752049, EPI_ISL_15752050, EPI_ISL_15752051, EPI_ISL_15752052, EPI_ISL_15752053, EPI_ISL_15752054, EPI_ISL_15752055, EPI_ISL_15752056, EPI_ISL_15752057, EPI_ISL_15752058, EPI_ISL_15752059, EPI_ISL_15752060, EPI_ISL_15752061, EPI_ISL_15752062, EPI_ISL_15752063, EPI_ISL_15752064, EPI_ISL_15752065, EPI_ISL_15752066, EPI_ISL_15752067, EPI_ISL_15752068, EPI_ISL_15752069, EPI_ISL_15752070, EPI_ISL_15752071, EPI_ISL_15752072, EPI_ISL_15752073, EPI_ISL_15752074, EPI_ISL_15752075, EPI_ISL_15752076, EPI_ISL_15752077, EPI_ISL_15752078, EPI_ISL_15752079, EPI_ISL_15752080, EPI_ISL_15752081, EPI_ISL_15752082 | | | |
| see above | Respiratory Viruses Branch, Division of Viral Diseases | Respiratory Viruses Branch, Division of Viral Diseases | Wang,L., Ng,T.F.F., Castro,C.J., Marine,R.L., Magana,L.C., Esona,M., Peret,T.C.T., Thornburg,N.J., Wang,L., Ng,T.F.F., Castro,C.J., Marine,R.L., Magana,L.C., Esona,M., Peret,T.C.T., Thornburg,N.J. |
| EPI_ISL_15753254, EPI_ISL_15753255, EPI_ISL_15753256, EPI_ISL_15753257, EPI_ISL_15753258, EPI_ISL_15753259, EPI_ISL_15753260, EPI_ISL_15753263, EPI_ISL_15753264, EPI_ISL_15753265, EPI_ISL_15753266, EPI_ISL_15753267, EPI_ISL_15753268, EPI_ISL_15753269, EPI_ISL_15753270, EPI_ISL_15753271, EPI_ISL_15753272, EPI_ISL_15753273, EPI_ISL_15753274, EPI_ISL_15753275, EPI_ISL_15753276, EPI_ISL_15753277, EPI_ISL_15753279, EPI_ISL_15753280, EPI_ISL_15753281, EPI_ISL_15753282, EPI_ISL_15753283, EPI_ISL_15753284, EPI_ISL_15753285, EPI_ISL_15753286, EPI_ISL_15753287, EPI_ISL_15753288, EPI_ISL_15753289, EPI_ISL_15753290, EPI_ISL_15753291, EPI_ISL_15753292, EPI_ISL_15753293, EPI_ISL_15753295, EPI_ISL_15753296, EPI_ISL_15753297, EPI_ISL_15753298, EPI_ISL_15753299, EPI_ISL_15753300, EPI_ISL_15753301, EPI_ISL_15753302, EPI_ISL_15753303, EPI_ISL_15753305, EPI_ISL_15753306, EPI_ISL_15753307, EPI_ISL_15753308, EPI_ISL_15753309, EPI_ISL_15753310, EPI_ISL_15753311, EPI_ISL_15753314, EPI_ISL_15753315, EPI_ISL_15753316, EPI_ISL_15753317, EPI_ISL_15753318, EPI_ISL_15753319, EPI_ISL_15753320, EPI_ISL_15753321, EPI_ISL_15753322, EPI_ISL_15753323, EPI_ISL_15753324, EPI_ISL_15753325, EPI_ISL_15753326, EPI_ISL_15753327, EPI_ISL_15753328, EPI_ISL_15753330, EPI_ISL_15753331, EPI_ISL_15753332, EPI_ISL_15753333, EPI_ISL_15753334, EPI_ISL_15753336, EPI_ISL_15753337, EPI_ISL_15753338, EPI_ISL_15753340, EPI_ISL_15753341, EPI_ISL_15753342, EPI_ISL_15753343, EPI_ISL_15753344, EPI_ISL_15753345, EPI_ISL_15753346, EPI_ISL_15753348, EPI_ISL_15753349, EPI_ISL_15753350, EPI_ISL_15753351, EPI_ISL_15753353, EPI_ISL_15753354, EPI_ISL_15753355, EPI_ISL_15753356, EPI_ISL_15753357, EPI_ISL_15753358, EPI_ISL_15753360, EPI_ISL_15753361, EPI_ISL_15753362, EPI_ISL_15753363, EPI_ISL_15753364, EPI_ISL_15753365, EPI_ISL_15753367, EPI_ISL_15753368, EPI_ISL_15753369, EPI_ISL_15753370, EPI_ISL_15753371, EPI_ISL_15753372, EPI_ISL_15753373, EPI_ISL_15753374, EPI_ISL_15753375, EPI_ISL_15753376, EPI_ISL_15753377, EPI_ISL_15753378, EPI_ISL_15753379, EPI_ISL_15753380, EPI_ISL_15753381, EPI_ISL_15753383, EPI_ISL_15753384, EPI_ISL_15753386, EPI_ISL_15753387, EPI_ISL_15753388, EPI_ISL_15753389, EPI_ISL_15753390, EPI_ISL_15753392, EPI_ISL_15753393, EPI_ISL_15753394, EPI_ISL_15753395, EPI_ISL_15753396, EPI_ISL_15753397, EPI_ISL_15753398, EPI_ISL_15753399, EPI_ISL_15753400, EPI_ISL_15753402, EPI_ISL_15753403, EPI_ISL_15753405, EPI_ISL_15753407, EPI_ISL_15753408, EPI_ISL_15753412, EPI_ISL_15753413, EPI_ISL_15753414, EPI_ISL_15753415, EPI_ISL_15753416, EPI_ISL_15753417, EPI_ISL_15753418, EPI_ISL_15753419, EPI_ISL_15753420, EPI_ISL_15753422, EPI_ISL_15753423, EPI_ISL_15753424, EPI_ISL_15753425, EPI_ISL_15753426, EPI_ISL_15753427, EPI_ISL_15753428, EPI_ISL_15753429, EPI_ISL_15753430, EPI_ISL_15753431, EPI_ISL_15753432, EPI_ISL_15753433, EPI_ISL_15753434, EPI_ISL_15753435, EPI_ISL_15753437, EPI_ISL_15753439, EPI_ISL_15753440, EPI_ISL_15753442, EPI_ISL_15753443, EPI_ISL_15753444, EPI_ISL_15753446, EPI_ISL_15753447, EPI_ISL_15753448, EPI_ISL_15753449, EPI_ISL_15753450, EPI_ISL_15753451, EPI_ISL_15753452, EPI_ISL_15753453, EPI_ISL_15753454, EPI_ISL_15753455, EPI_ISL_15753457, EPI_ISL_15753458, EPI_ISL_15753459, EPI_ISL_15753460, EPI_ISL_15753461, EPI_ISL_15753462, EPI_ISL_15753463, EPI_ISL_15753464, EPI_ISL_15753465, EPI_ISL_15753466, EPI_ISL_15753467, EPI_ISL_15753468, EPI_ISL_15753469, EPI_ISL_15753470, EPI_ISL_15753471, EPI_ISL_15753472, EPI_ISL_15753473, EPI_ISL_15753474, EPI_ISL_15753476, EPI_ISL_15753477, EPI_ISL_15753478, EPI_ISL_15753479, EPI_ISL_15753480, EPI_ISL_15753481, EPI_ISL_15753482, EPI_ISL_15753483, EPI_ISL_15753484, EPI_ISL_15753485, EPI_ISL_15753486, EPI_ISL_15753488, EPI_ISL_15753490, EPI_ISL_15753491, EPI_ISL_15753492, EPI_ISL_15753493, EPI_ISL_15753494, EPI_ISL_15753495, EPI_ISL_15753496, EPI_ISL_15753497, EPI_ISL_15753498, EPI_ISL_15753499, EPI_ISL_15753500, EPI_ISL_15753501, EPI_ISL_15753502, EPI_ISL_15753503, EPI_ISL_15753504, EPI_ISL_15753505, EPI_ISL_15753507, EPI_ISL_15753508, EPI_ISL_15753509, EPI_ISL_15753510, EPI_ISL_15753511, EPI_ISL_15753512, EPI_ISL_15753514, EPI_ISL_15753515, EPI_ISL_15753516, EPI_ISL_15753517, EPI_ISL_15753518, EPI_ISL_15753519, EPI_ISL_15753520, EPI_ISL_15753521, EPI_ISL_15753523, EPI_ISL_15753524, EPI_ISL_15753525, EPI_ISL_15753526, EPI_ISL_15753527, EPI_ISL_15753528, EPI_ISL_15753529, EPI_ISL_15753530, EPI_ISL_15753531, EPI_ISL_15753532, EPI_ISL_15753534, EPI_ISL_15753535, EPI_ISL_15753537, EPI_ISL_15753538, EPI_ISL_15753539, EPI_ISL_15753540, EPI_ISL_15753541, EPI_ISL_15753542, EPI_ISL_15753544, EPI_ISL_15753545, EPI_ISL_15753546, EPI_ISL_15753547, EPI_ISL_15753548, EPI_ISL_15753549, EPI_ISL_15753550, EPI_ISL_15753551, EPI_ISL_15753552, EPI_ISL_15753553, EPI_ISL_15753554, EPI_ISL_15753555, EPI_ISL_15753556, EPI_ISL_15753557, EPI_ISL_15753558, EPI_ISL_15753559, EPI_ISL_15753560, EPI_ISL_15753561, EPI_ISL_15753562, EPI_ISL_15753563, EPI_ISL_15753564, EPI_ISL_15753565, EPI_ISL_15753566, EPI_ISL_15753568, EPI_ISL_15753569 | | | |
| see above | Department of Paediatrics, University of Oxford | Department of Paediatrics, University of Oxford | Lin,G.-L., Drysdale,S.B., Snape,M.D., O'Connor,D., Brown,A., MacIntyre-Cockett,G., Mellado-Gomez,E., de Cesare,M., Bonsall,D., Ansari,M.A., Oner,D., Aerssens,J., Butler,C., Bont,L., Openshaw,P., Martinon-Torres,F., Nair,H., Bowden,R., Golubchik,T., Pollard,A.J., Lin,G.-L. |
| EPI_ISL_15753574 | Eurasian Institute of zoonotic infections, The Federal Research Center of Fundamental and Translational Medicine | Eurasian Institute of zoonotic infections, The Federal Research Center of Fundamental and Translational Medicine | Dubovitskiy,N.A., Sobolev,I.A., Kurskaya,O.G., Anoshina,A.V., Leonova,N.V., Murashkina,T.A., Solomatina,M.V., Derko,A.A., Saroyan,T.A., Kabilov,M.R., Tupikin,A.E., Sharshov,K.A., Shestopalov,A.M. |
| EPI_ISL_15753575, EPI_ISL_15753577 | Eurasian Institute of zoonotic infections, The Federal Research Center of Fundamental and | Eurasian Institute of zoonotic infections, The Federal Research Center of Fundamental and | Dubovitskiy,N.A., Sobolev,I.A., Kurskaya,O.G., Simkina,O.A., Komissarova,T.V., Murashkina,T.A., Solomatina,M.V., Derko,A.A., Saroyan,T.A., Kabilov,M.R., Tupikin,A.E., Sharshov,K.A., Shestopalov,A.M. |

| | | | |
|---|---|---|---|
| | Translational Medicine | Translational Medicine | |
| EPI_ISL_15771587, EPI_ISL_15771589, EPI_ISL_15771591, EPI_ISL_15771593, EPI_ISL_15771595, EPI_ISL_15771598, EPI_ISL_15771600, EPI_ISL_15771601, EPI_ISL_15771603, EPI_ISL_15771604, EPI_ISL_15771606, EPI_ISL_15771609, EPI_ISL_15771611, EPI_ISL_15771613, EPI_ISL_15771615 | | | |
| see above | Respiratory Viruses Branch, Division of Viral Diseases, Centers for Disease Control and Prevention | Respiratory Viruses Branch, Division of Viral Diseases, Centers for Disease Control and Prevention | Wang,L., Ng,T.F.F., Castro,C.J., Marine,R.L., Magana,L.C., Esona,M., Peret,T.C.T. and Thornburg,N.J. |
| EPI_ISL_15772191, EPI_ISL_15772194 | Laboratory Medicine, UW Virology | Laboratory Medicine, UW Virology | Sereewit,J., Xie,H. and Greninger,A. |
| EPI_ISL_15773986, EPI_ISL_15773991, EPI_ISL_15773995, EPI_ISL_15774001, EPI_ISL_15774006, EPI_ISL_15774010, EPI_ISL_15774011, EPI_ISL_15774012, EPI_ISL_15774013, EPI_ISL_15774014, EPI_ISL_15774015, EPI_ISL_15774016, EPI_ISL_15774017, EPI_ISL_15774018, EPI_ISL_15774020, EPI_ISL_15774021, EPI_ISL_15774022, EPI_ISL_15774023, EPI_ISL_15774024, EPI_ISL_15774025, EPI_ISL_15774026, EPI_ISL_15774027, EPI_ISL_15774029, EPI_ISL_15774030, EPI_ISL_15774031, EPI_ISL_15774032, EPI_ISL_15774033, EPI_ISL_15774035, EPI_ISL_15774038, EPI_ISL_15774039, EPI_ISL_15774042, EPI_ISL_15774043, EPI_ISL_15774045, EPI_ISL_15774046, EPI_ISL_15774047, EPI_ISL_15774049, EPI_ISL_15774050, EPI_ISL_15774053, EPI_ISL_15774054, EPI_ISL_15774056, EPI_ISL_15774058, EPI_ISL_15774059, EPI_ISL_15774060, EPI_ISL_15774062, EPI_ISL_15774063, EPI_ISL_15774064, EPI_ISL_15774065, EPI_ISL_15774066, EPI_ISL_15774067, EPI_ISL_15774068, EPI_ISL_15774070, EPI_ISL_15774071 | | | |
| see above | QLD, Royal Brisbane and Woman's Hospital | WHO Collaborating Centre for Reference and Research on Influenza | Xiaomin Dong, Yi-Mo Deng, Ammar Aziz, Naomi Komadina |
| EPI_ISL_15820266, EPI_ISL_15820267, EPI_ISL_15820268 | Respiratory Viruses Branch, Division of Viral Diseases | Respiratory Viruses Branch, Division of Viral Diseases | Wang,L., Ng,T.F.F., Castro,C.J., Marine,R.L., Magana,L.C., Esona,M., Peret,T.C.T., Thornburg,N.J., Wang,L., Ng,T.F.F., Castro,C.J., Marine,R.L., Magana,L.C., Esona,M., Peret,T.C.T., Thornburg,N.J. |
| EPI_ISL_15894968, EPI_ISL_15894969, EPI_ISL_15894972, EPI_ISL_15894973, EPI_ISL_15894974, EPI_ISL_15894976, EPI_ISL_15894979, EPI_ISL_15894981, EPI_ISL_15894983, EPI_ISL_15894985, EPI_ISL_15894988, EPI_ISL_15894989, EPI_ISL_15894990, EPI_ISL_15894991, EPI_ISL_15894994, EPI_ISL_15894996, EPI_ISL_15894997, EPI_ISL_15895000, EPI_ISL_15895003, EPI_ISL_15895004, EPI_ISL_15895005, EPI_ISL_15895007, EPI_ISL_15895008, EPI_ISL_15895010, EPI_ISL_15895011, EPI_ISL_15895012, EPI_ISL_15895013, EPI_ISL_15895015, EPI_ISL_15895017, EPI_ISL_15895018, EPI_ISL_15895019, EPI_ISL_15895021 | | | |
| see above | Laboratorio Central de Saude Publica do Estado do Rio Grande do Sul (LACEN/RS) | Laboratory of Respiratory Viruses and Measles, Oswaldo Cruz Institute, FIOCRUZ | Paola Resende, Fernando Motta, Elisa Cavalcante Pereira, Bruna Mendonça da Silva, Jéssica Graça Macedo de Carvalho, Larissa Macedo Pinto, Victor Guimaraes, Ighor Leonardo Arantes, Leticia Scalioni, Anderson Brandao Leite, Marilda Siqueira on behalf of the Fiocruz COVID-19 Genomic Surveillance Network |
| EPI_ISL_15895037, EPI_ISL_15895040, EPI_ISL_15895043, EPI_ISL_15895044 | Laboratorio Central de Saude Publica do Estado do Espirito Santo (LACEN/ES) | Laboratory of Respiratory Viruses and Measles, Oswaldo Cruz Institute, FIOCRUZ | Paola Resende, Fernando Motta, Elisa Cavalcante Pereira, Ighor Arantes, Bruna Mendonça da Silva, Jéssica Graça Macedo de Carvalho, Larissa Macedo Pinto, Victor Guimaraes, Leticia Scalioni, Rodrigo Ribeiro Rodrigues, Marilda Siqueira on behalf of the Fiocruz COVID-19 Genomic Surveillance Network |
| EPI_ISL_15895046, EPI_ISL_15895050, EPI_ISL_15895051, EPI_ISL_15895053, EPI_ISL_15895054 | Laboratorio Central de Saude Publica do Estado do Rio Grande do Sul (LACEN/RS) | Laboratory of Respiratory Viruses and Measles, Oswaldo Cruz Institute, FIOCRUZ | Paola Resende, Fernando Motta, Elisa Cavalcante Pereira, Bruna Mendonça da Silva, Jéssica Graça Macedo de Carvalho, Larissa Macedo Pinto, Victor Guimaraes, Ighor Leonardo Arantes, Leticia Scalioni, Anderson Brandao Leite, Marilda Siqueira on behalf of the Fiocruz COVID-19 Genomic Surveillance Network |
| EPI_ISL_15895070, EPI_ISL_15895082, EPI_ISL_15895083, EPI_ISL_15895086, EPI_ISL_15895087, EPI_ISL_15895088, EPI_ISL_15895089, EPI_ISL_15895091, EPI_ISL_15895092 | Laboratorio Central de Saude Publica do Estado do Espirito Santo (LACEN/ES) | Laboratory of Respiratory Viruses and Measles, Oswaldo Cruz Institute, FIOCRUZ | Paola Resende, Fernando Motta, Elisa Cavalcante Pereira, Ighor Arantes, Bruna Mendonça da Silva, Jéssica Graça Macedo de Carvalho, Larissa Macedo Pinto, Victor Guimaraes, Leticia Scalioni, Rodrigo Ribeiro Rodrigues, Marilda Siqueira on behalf of the Fiocruz COVID-19 Genomic Surveillance Network |
| EPI_ISL_15895096, EPI_ISL_15895098, EPI_ISL_15895100, EPI_ISL_15895101, EPI_ISL_15895102, EPI_ISL_15895103, EPI_ISL_15895105, EPI_ISL_15895107, EPI_ISL_15895110, EPI_ISL_15895114, EPI_ISL_15895115, EPI_ISL_15895116, EPI_ISL_15895117, EPI_ISL_15895118, EPI_ISL_15895119, EPI_ISL_15895120 | | | |
| see above | Laboratorio Central de Saude Publica do Estado do Parana (LACEN/PR) | Laboratory of Respiratory Viruses and Measles, Oswaldo Cruz Institute, FIOCRUZ | Paola Resende, Fernando Motta, Elisa Cavalcante Pereira, Ighor Arantes, Bruna Mendonça da Silva, Jéssica Graça Macedo de Carvalho, Larissa Macedo Pinto, Victor Guimaraes, Leticia Scalioni, Irina Riediger, Marilda Siqueira on behalf of the Fiocruz COVID-19 Genomic Surveillance Network |
| EPI_ISL_15895124 | Universidade Federal de São Paulo, Departamento de Medicina, Disciplina de Doenças Infecciosas e Parasitárias. Laboratório de Virologia | Laboratory of Respiratory Viruses and Measles, Oswaldo Cruz Institute, FIOCRUZ | Paola Resende, Fernando Motta, Elisa Cavalcante Pereira, Ighor Arantes, Bruna Mendonça da Silva, Jéssica Graça Macedo de Carvalho, Larissa Macedo Pinto, Victor Guimaraes, Leticia Scalioni, Ana Helena Perosa, Gabriela Rodrigues Barbosa, Nancy Bellei e Marilda Siqueira on behalf of the Fiocruz COVID-19 Genomic Surveillance Network |
| EPI_ISL_15895130 | Laboratorio Central de Saude Publica do Estado do Amapa (LACEN/AP) | Laboratory of Respiratory Viruses and Measles, Oswaldo Cruz Institute, FIOCRUZ | Paola Resende, Fernando Motta, Elisa Cavalcante Pereira, Ighor Arantes, Bruna Mendonça da Silva, Jéssica Graça Macedo de Carvalho, Larissa Macedo Pinto, Victor Guimaraes, Leticia Scalioni, Andreia Santos Costa, Marcia Socorro Pereira Cavalcante, Anne Caroline da Silva Soledade, Lindomar dos Anjos Silva, Marilda Siqueira on behalf of the Fiocruz COVID-19 Genomic Surveillance Network |
| EPI_ISL_15895133, EPI_ISL_15895134 | Universidade Federal de São Paulo, Departamento de Medicina, Disciplina de Doenças Infecciosas e Parasitárias. Laboratório de Virologia | Laboratory of Respiratory Viruses and Measles, Oswaldo Cruz Institute, FIOCRUZ | Paola Resende, Fernando Motta, Elisa Cavalcante Pereira, Ighor Arantes, Bruna Mendonça da Silva, Jéssica Graça Macedo de Carvalho, Larissa Macedo Pinto, Victor Guimaraes, Leticia Scalioni, Ana Helena Perosa, Gabriela Rodrigues Barbosa, Nancy Bellei e Marilda Siqueira on behalf of the Fiocruz COVID-19 Genomic Surveillance Network |
| EPI_ISL_15895136, EPI_ISL_15895137 | Laboratorio Central de Saude Publica do Estado do Parana (LACEN/PR) | Laboratory of Respiratory Viruses and Measles, Oswaldo Cruz Institute, FIOCRUZ | Paola Resende, Fernando Motta, Elisa Cavalcante Pereira, Ighor Arantes, Bruna Mendonça da Silva, Jéssica Graça Macedo de Carvalho, Larissa Macedo Pinto, Victor Guimaraes, Leticia Scalioni, Irina Riediger, Marilda Siqueira on behalf of the Fiocruz COVID-19 Genomic Surveillance Network |
| EPI_ISL_15895140, EPI_ISL_15895141 | Laboratory of Respiratory Viruses and Measles, Oswaldo Cruz Institute, FIOCRUZ | Laboratory of Respiratory Viruses and Measles, Oswaldo Cruz Institute, FIOCRUZ | Paola Resende, Luciana Appolinario, Fernando Motta, Elisa Cavalcante Pereira, Ighor Arantes, Alice Sampaio Rocha, Bruna Mendonça da Silva, Jéssica Graça Macedo de Carvalho, Larissa Macedo Pinto, Victor Guimaraes, Leticia Scalioni, Marilda Siqueira on behalf of the Fiocruz COVID-19 Genomic Surveillance Network |
| EPI_ISL_15895142 | Universidade Federal de São Paulo, Departamento de Medicina, Disciplina de Doenças Infecciosas e Parasitárias. Laboratório de Virologia | Laboratory of Respiratory Viruses and Measles, Oswaldo Cruz Institute, FIOCRUZ | Paola Resende, Fernando Motta, Elisa Cavalcante Pereira, Ighor Arantes, Bruna Mendonça da Silva, Jéssica Graça Macedo de Carvalho, Larissa Macedo Pinto, Victor Guimaraes, Leticia Scalioni, Ana Helena Perosa, Gabriela Rodrigues Barbosa, Nancy Bellei e Marilda Siqueira on behalf of the Fiocruz COVID-19 Genomic Surveillance Network |
| EPI_ISL_15896139, EPI_ISL_15896141, EPI_ISL_15896143 | NT, Royal Darwin Hospital | WHO Collaborating Centre for Reference and Research on Influenza | Xiaomin Dong, Steven Edwards, Yi-Mo Deng, Ammar Aziz, Ian Barr |
| EPI_ISL_15896144, EPI_ISL_15896145, EPI_ISL_15896146 | QLD, Queensland Childrens Hospital | WHO Collaborating Centre for Reference and Research on Influenza | Xiaomin Dong, Steven Edwards, Yi-Mo Deng, Ammar Aziz, Ian Barr |
| EPI_ISL_15896148, EPI_ISL_15896149, EPI_ISL_15896150, EPI_ISL_15896151, EPI_ISL_15896152 | VIC, RCH Molecular Microbiology Dept. (Bio21) | WHO Collaborating Centre for Reference and Research on Influenza | Xiaomin Dong, Steven Edwards, Yi-Mo Deng, Ammar Aziz, Ian Barr |
| EPI_ISL_15896155 | VIC, Royal Children's Hospital | WHO Collaborating Centre for Reference and Research on Influenza | Xiaomin Dong, Steven Edwards, Yi-Mo Deng, Ammar Aziz, Ian Barr |
| EPI_ISL_15896156 | NT, Royal Darwin Hospital | WHO Collaborating Centre for Reference and Research on Influenza | Xiaomin Dong, Steven Edwards, Yi-Mo Deng, Ammar Aziz, Ian Barr |
| EPI_ISL_15896158, EPI_ISL_15896159 | VIC, RCH Molecular Microbiology Dept. (Bio21) | WHO Collaborating Centre for Reference and Research on Influenza | Xiaomin Dong, Steven Edwards, Yi-Mo Deng, Ammar Aziz, Ian Barr |
| EPI_ISL_15896160, EPI_ISL_15896161 | NT, Royal Darwin Hospital | WHO Collaborating Centre for Reference and Research on Influenza | Xiaomin Dong, Steven Edwards, Yi-Mo Deng, Ammar Aziz, Ian Barr |
| EPI_ISL_15896162, EPI_ISL_15896164 | QLD, Queensland Childrens Hospital | WHO Collaborating Centre for Reference and Research on Influenza | Xiaomin Dong, Steven Edwards, Yi-Mo Deng, Ammar Aziz, Ian Barr |
| EPI_ISL_15896166, EPI_ISL_15896167 | VIC, RCH Molecular Microbiology Dept. (Bio21) | WHO Collaborating Centre for Reference and Research on Influenza | Xiaomin Dong, Steven Edwards, Yi-Mo Deng, Ammar Aziz, Ian Barr |
| EPI_ISL_15896168, EPI_ISL_15896169, EPI_ISL_15896170, EPI_ISL_15896171 | NT, Royal Darwin Hospital | WHO Collaborating Centre for Reference and Research on Influenza | Xiaomin Dong, Steven Edwards, Yi-Mo Deng, Ammar Aziz, Ian Barr |
| EPI_ISL_15896172 | QLD, Queensland Childrens Hospital | WHO Collaborating Centre for Reference and Research on Influenza | Xiaomin Dong, Steven Edwards, Yi-Mo Deng, Ammar Aziz, Ian Barr |
| EPI_ISL_15896175, EPI_ISL_15896176, EPI_ISL_15896177, EPI_ISL_15896178, EPI_ISL_15896179, EPI_ISL_15896181, EPI_ISL_15896182, EPI_ISL_15896183 | VIC, RCH Molecular Microbiology Dept. (Bio21) | WHO Collaborating Centre for Reference and Research on Influenza | Xiaomin Dong, Steven Edwards, Yi-Mo Deng, Ammar Aziz, Ian Barr |
| EPI_ISL_15896186, EPI_ISL_15896187, EPI_ISL_15896188, EPI_ISL_15896189, EPI_ISL_15896193 | NT, Royal Darwin Hospital | WHO Collaborating Centre for Reference and Research on Influenza | Xiaomin Dong, Steven Edwards, Yi-Mo Deng, Ammar Aziz, Ian Barr |
| EPI_ISL_15896194, EPI_ISL_15896195 | VIC, RCH Molecular Microbiology Dept. (Bio21) | WHO Collaborating Centre for Reference and Research on Influenza | Xiaomin Dong, Steven Edwards, Yi-Mo Deng, Ammar Aziz, Ian Barr |
| EPI_ISL_15896198, EPI_ISL_15896199, EPI_ISL_15896202, EPI_ISL_15896203 | NT, Royal Darwin Hospital | WHO Collaborating Centre for Reference and Research on Influenza | Xiaomin Dong, Steven Edwards, Yi-Mo Deng, Ammar Aziz, Ian Barr |
| EPI_ISL_15896211, EPI_ISL_15896213, EPI_ISL_15896218 | VIC, RCH Molecular Microbiology Dept. (Bio21) | WHO Collaborating Centre for Reference and Research on Influenza | Xiaomin Dong, Steven Edwards, Yi-Mo Deng, Ammar Aziz, Ian Barr |
| EPI_ISL_15896222 | NT, Royal Darwin Hospital | WHO Collaborating Centre for Reference and Research on Influenza | Xiaomin Dong, Steven Edwards, Yi-Mo Deng, Ammar Aziz, Ian Barr |
| EPI_ISL_15896234 | QLD, Queensland Childrens Hospital | WHO Collaborating Centre for Reference and Research on Influenza | Xiaomin Dong, Steven Edwards, Yi-Mo Deng, Ammar Aziz, Ian Barr |
| EPI_ISL_15896257 | NT, Royal Darwin Hospital | WHO Collaborating Centre for Reference and Research on Influenza | Xiaomin Dong, Steven Edwards, Yi-Mo Deng, Ammar Aziz, Ian Barr |
| EPI_ISL_15896294 | VIC, RCH Molecular Microbiology Dept. (Bio21) | WHO Collaborating Centre for Reference and Research on Influenza | Xiaomin Dong, Steven Edwards, Yi-Mo Deng, Ammar Aziz, Ian Barr |
| EPI_ISL_16132328 | University of WashingtonLaboratory Medicine and Pathology, UW Medicine | University of WashingtonLaboratory Medicine and Pathology, UW Medicine | Goya,S., Sereewit,J., Pfalmer,D., Nguyen,T., Sobolik,E.B. and Greninger,A.L. |
| EPI_ISL_16132330, EPI_ISL_16132331, EPI_ISL_16132332, EPI_ISL_16132333, EPI_ISL_16132334, EPI_ISL_16132336, EPI_ISL_16132338, EPI_ISL_16132339, EPI_ISL_16132340, EPI_ISL_16132341, EPI_ISL_16132342, EPI_ISL_16132343, EPI_ISL_16132344, EPI_ISL_16132345, EPI_ISL_16132346, EPI_ISL_16132347, EPI_ISL_16132348, EPI_ISL_16132349, EPI_ISL_16132350, EPI_ISL_16132351, EPI_ISL_16132353, EPI_ISL_16132354, EPI_ISL_16132355, EPI_ISL_16132356, EPI_ISL_16132357, EPI_ISL_16132360, EPI_ISL_16132361, EPI_ISL_16132362, EPI_ISL_16132363, EPI_ISL_16132364, EPI_ISL_16132365, EPI_ISL_16132366, EPI_ISL_16132368, EPI_ISL_16132369, EPI_ISL_16132371, EPI_ISL_16132372, EPI_ISL_16132376, EPI_ISL_16132377, EPI_ISL_16132378 | | | |
| see above | Infectious Disease Program, Broad Institute of Harvard and MIT | Infectious Disease Program, Broad Institute of Harvard and MIT | Adams,G., Uddin,R., Messer,K., Dobbins,S., Kotzen,B., Siddle,K.J., Petros,B., Paull,J., Brock-Fisher,T., Levine,Z., Krasilnikova,L., Negrete-Arenas,F., Tomkins-Tinch,C., Chaluvadi,S., Marmol,C., DeRuff,K., Loreth,C., Birren,B.W., Park,D.J., MacInnis,B.L., Sabeti,P.C., Rosenberg,E., Turbett,S., Lemieux,J.E., Adams,G., Uddin,R., Messer,K., Dobbins,S., Kotzen,B., Siddle,K.J., Petros,B., Paull,J., Brock-Fisher,T., Levine,Z., Krasilnikova,L., Negrete-Arenas,F., Tomkins-Tinch,C., Chaluvadi,S., Marmol,C., DeRuff,K., Loreth,C., Birren,B.W., Park,D.J., MacInnis,B.L., Sabeti,P.C., Rosenberg,E., Turbett,S., Lemieux,J.E. |
| EPI_ISL_16132384, EPI_ISL_16132385, EPI_ISL_16132386, EPI_ISL_16132387, EPI_ISL_16132388, EPI_ISL_16132389, EPI_ISL_16132390, EPI_ISL_16132391, EPI_ISL_16132392, EPI_ISL_16132393, EPI_ISL_16132394, EPI_ISL_16132395, EPI_ISL_16132396, EPI_ISL_16132397, EPI_ISL_16132398, EPI_ISL_16132399, EPI_ISL_16132400, EPI_ISL_16132401, EPI_ISL_16132402, EPI_ISL_16132403, | | | |

| | | | |
|---|---|---|---|
| EPI_ISL_16132404, EPI_ISL_16132405, EPI_ISL_16132406, EPI_ISL_16132407, EPI_ISL_16132408, EPI_ISL_16132409, EPI_ISL_16132410, EPI_ISL_16132411, EPI_ISL_16132412 | | | |
| see above | University of WashingtonLaboratory Medicine and Pathology, UW Medicine | University of WashingtonLaboratory Medicine and Pathology, UW Medicine | Goya,S., Sereewit,J., Pfalmer,D., Nguyen,T., Sobolik,E.B. and Greninger,A.L. |
| EPI_ISL_16251628, EPI_ISL_16251629, EPI_ISL_16251631, EPI_ISL_16251632, EPI_ISL_16251633, EPI_ISL_16251634, EPI_ISL_16251635, EPI_ISL_16251638, EPI_ISL_16251639, EPI_ISL_16251640, EPI_ISL_16251641, EPI_ISL_16251646, EPI_ISL_16251647, EPI_ISL_16251650, EPI_ISL_16251651, EPI_ISL_16251652, EPI_ISL_16251653, EPI_ISL_16251654, EPI_ISL_16251656, EPI_ISL_16251657, EPI_ISL_16251658, EPI_ISL_16251661, EPI_ISL_16251663, EPI_ISL_16251665, EPI_ISL_16251668, EPI_ISL_16251671, EPI_ISL_16251676, EPI_ISL_16251682, EPI_ISL_16251683 | | | |
| see above | Microbiology Department. Complexo Hospitalario Universitario de Vigo | Microbiology Department. Complexo Hospitalario Universitario de Vigo | Daviña C, Martinez L, Perez-Castro S |
| EPI_ISL_16282716, EPI_ISL_16282721, EPI_ISL_16282737, EPI_ISL_16282746, EPI_ISL_16282749 | National Influenza Center, National Institute of Hygiene and Epidemiology (NIHE) | National Influenza Center, National Institute of Hygiene and Epidemiology (NIHE) | Ung Thi Hong Trang, Hoang Vu Mai Phuong, Nguyen Huy Hoang, Nguyen Le Khanh Hang, Le Thi Quynh Mai |
| EPI_ISL_16289021, EPI_ISL_16289023, EPI_ISL_16289025, EPI_ISL_16289027, EPI_ISL_16289028, EPI_ISL_16289030, EPI_ISL_16289031, EPI_ISL_16289032, EPI_ISL_16289033, EPI_ISL_16289034, EPI_ISL_16289039, EPI_ISL_16289041, EPI_ISL_16289042, EPI_ISL_16289044, EPI_ISL_16289047, EPI_ISL_16289048 | | | |
| see above | Microbiology Department. Complexo Hospitalario Universitario de Vigo | Microbiology Department. Complexo Hospitalario Universitario de Vigo | Daviña C, Martinez L, Perez-Castro S |
| EPI_ISL_1647383, EPI_ISL_1647384, EPI_ISL_1647385, EPI_ISL_1647386, EPI_ISL_1647387, EPI_ISL_1647388, EPI_ISL_1647389, EPI_ISL_1647390, EPI_ISL_1647391, EPI_ISL_1647392, EPI_ISL_1647393, EPI_ISL_1647394, EPI_ISL_1647395, EPI_ISL_1647396, EPI_ISL_1647397, EPI_ISL_1647398, EPI_ISL_1647399, EPI_ISL_1647400, EPI_ISL_1647401, EPI_ISL_1647402, EPI_ISL_1647403, EPI_ISL_1647404, EPI_ISL_1647405, EPI_ISL_1647406, EPI_ISL_1647407, EPI_ISL_1647408, EPI_ISL_1647409, EPI_ISL_1647410, EPI_ISL_1647411, EPI_ISL_1647412, EPI_ISL_1647413, EPI_ISL_1647414, EPI_ISL_1647415, EPI_ISL_1647416, EPI_ISL_1647417, EPI_ISL_1647418, EPI_ISL_1647419, EPI_ISL_1647420, EPI_ISL_1647421, EPI_ISL_1647422 | | | |
| see above | Respiratory Virus Unit, National Infection Service, Public Health England | National Infection Service, Public Health England | Zambon M, Talts T, Ellis J, Miah S, Platt S |
| EPI_ISL_1647456, EPI_ISL_1647457, EPI_ISL_1647458, EPI_ISL_1647459, EPI_ISL_1647460, EPI_ISL_1647461, EPI_ISL_1647462, EPI_ISL_1647463, EPI_ISL_1647464, EPI_ISL_1647465, EPI_ISL_1647466, EPI_ISL_1647467, EPI_ISL_1647468, EPI_ISL_1647469 | | | |
| see above | Instituto Nacional de Saúde | WHO Influenza Centre for Reference and Research on Influenza | Angela Todd, Yi-Mo Deng, Almiro Rogerio Tivane, Naomi Komadina |
| EPI_ISL_16533868, EPI_ISL_16533869, EPI_ISL_16533870, EPI_ISL_16533871, EPI_ISL_16533873, EPI_ISL_16533874 | Center for Virology, Medical University of Vienna | Center for Virology, Medical University of Vienna | Jeremy V. Camp, Monika Redlberger-Fritz |
| EPI_ISL_1653937 | Royal Children's Hospital | WHO Influenza Centre for Reference and Research on Influenza | Angela Todd, Yi-Mo Deng, Annette Alafaci, Naomi Komadina |
| EPI_ISL_1653938, EPI_ISL_1653939 | Royal Children's Hospital | WHO Influenza Centre for Reference and Research on Influenza | Jean Moselen, Yi-Mo Deng, Annette Alafaci, Naomi Komadina |
| EPI_ISL_1653940 | Royal Children's Hospital | WHO Influenza Centre for Reference and Research on Influenza | Angela Todd, Yi-Mo Deng, Annette Alafaci, Naomi Komadina |
| EPI_ISL_1653941, EPI_ISL_1653942 | Royal Children's Hospital | WHO Influenza Centre for Reference and Research on Influenza | Jean Moselen, Yi-Mo Deng, Annette Alafaci, Naomi Komadina |
| EPI_ISL_1653943 | Royal Children's Hospital | WHO Influenza Centre for Reference and Research on Influenza | Angela Todd, Yi-Mo Deng, Annette Alafaci, Naomi Komadina |
| EPI_ISL_1653944 | Royal Children's Hospital | WHO Influenza Centre for Reference and Research on Influenza | Jean Moselen, Yi-Mo Deng, Annette Alafaci, Naomi Komadina |
| EPI_ISL_1653945 | Royal Children's Hospital | WHO Influenza Centre for Reference and Research on Influenza | Angela Todd, Yi-Mo Deng, Annette Alafaci, Naomi Komadina |
| EPI_ISL_1653946 | Royal Children's Hospital | WHO Influenza Centre for Reference and Research on Influenza | Jean Moselen, Yi-Mo Deng, Annette Alafaci, Naomi Komadina |
| EPI_ISL_1653947 | Royal Children's Hospital | WHO Influenza Centre for Reference and Research on Influenza | Angela Todd, Yi-Mo Deng, Annette Alafaci, Naomi Komadina |
| EPI_ISL_1653948 | Royal Children's Hospital | WHO Influenza Centre for Reference and Research on Influenza | Jean Moselen, Yi-Mo Deng, Annette Alafaci, Naomi Komadina |
| EPI_ISL_1653949, EPI_ISL_1653950, EPI_ISL_1653951, EPI_ISL_1653952, EPI_ISL_1653953, EPI_ISL_1653954, EPI_ISL_1653955, EPI_ISL_1653956, EPI_ISL_1653957, EPI_ISL_1653958, EPI_ISL_1653959, EPI_ISL_1653960, EPI_ISL_1653961, EPI_ISL_1653962, EPI_ISL_1653963, EPI_ISL_1653964, EPI_ISL_1653965, EPI_ISL_1653966, EPI_ISL_1653967, EPI_ISL_1653968, EPI_ISL_1653969, EPI_ISL_1653970, EPI_ISL_1653971, EPI_ISL_1653972, EPI_ISL_1653973, EPI_ISL_1653974, EPI_ISL_1653975, EPI_ISL_1653976, EPI_ISL_1653977, EPI_ISL_1653978, EPI_ISL_1653979, EPI_ISL_1653980, EPI_ISL_1653981, EPI_ISL_1653982, EPI_ISL_1653983, EPI_ISL_1653984, EPI_ISL_1653985, EPI_ISL_1653986, EPI_ISL_1653987, EPI_ISL_1653988, EPI_ISL_1653989, EPI_ISL_1653990, EPI_ISL_1653991, EPI_ISL_1653992 | | | |
| see above | Royal Children's Hospital | WHO Influenza Centre for Reference and Research on Influenza | Angela Todd, Yi-Mo Deng, Annette Alafaci, Naomi Komadina |
| EPI_ISL_16672827, EPI_ISL_16672828, EPI_ISL_16672829 | Respiratory Virus Unit / Reference Microbiology Services / UK Health Security Agency | Reference Microbiology Services / UK Health Security Agency | Zambon M. Talts T. Mosscrop L. Miah S |
| EPI_ISL_16681182, EPI_ISL_16681183, EPI_ISL_16681201, EPI_ISL_16681239, EPI_ISL_16681240, EPI_ISL_16681247, EPI_ISL_16681257, EPI_ISL_16681258, EPI_ISL_16681259, EPI_ISL_16681261, EPI_ISL_16681270, EPI_ISL_16681271, EPI_ISL_16681273, EPI_ISL_16681283, EPI_ISL_16681284, EPI_ISL_16681285, EPI_ISL_16681296, EPI_ISL_16681297, EPI_ISL_16681298, EPI_ISL_16681300, EPI_ISL_16681301, EPI_ISL_16681308, EPI_ISL_16681309, EPI_ISL_16681310, EPI_ISL_16681311, EPI_ISL_16681323, EPI_ISL_16681325, EPI_ISL_16681326, EPI_ISL_16681336, EPI_ISL_16681337, EPI_ISL_16681339, EPI_ISL_16681340, EPI_ISL_16681341, EPI_ISL_16681349, EPI_ISL_16681360, EPI_ISL_16681408 | | | |
| see above | Broad Institute of Harvard and Massachusetts Institute of Technology | Broad Institute of Harvard and Massachusetts Institute of Technology | Adams,G., Uddin,R., Messer,K., Dobbins,S., Kotzen,B., Siddle,K.J., Petros,B., Paull,J., Brock-Fisher,T., Levine,Z., Krasilnikova,L., Negrete-Arenas,F., Tomkins-Tinch,C., Chaluvadi,S., Marmol,C., DeRuff,K., Loreth,C., Birren,B.W., Park,D.J., MacInnis,B.L., Sabeti,P.C., Rosenberg,E., Turbett,S. and Lemieux,J.E. |
| EPI_ISL_16714485, EPI_ISL_16714508, EPI_ISL_16714559, EPI_ISL_16714565 | Respiratory Virus Unit / Reference Microbiology Services / UK Health Security Agency | Reference Microbiology Services / UK Health Security Agency | Zambon M. Talts T. Mosscrop L. Miah S |
| EPI_ISL_16714745, EPI_ISL_16714747, EPI_ISL_16714749 | VIC, RCH Molecular Microbiology Dept. (Bio21) | WHO Collaborating Centre for Reference and Research on Influenza | Xiaomin Dong, Steven Edwards, Yi-Mo Deng, Ammar Aziz, Ian Barr |
| EPI_ISL_16714760, EPI_ISL_16714770, EPI_ISL_16714774, EPI_ISL_16714780 | NT, Royal Darwin Hospital | WHO Collaborating Centre for Reference and Research on Influenza | Xiaomin Dong, Steven Edwards, Yi-Mo Deng, Ammar Aziz, Ian Barr |
| EPI_ISL_16714781, EPI_ISL_16714786, EPI_ISL_16714789, EPI_ISL_16714791, EPI_ISL_16714795, EPI_ISL_16714797, EPI_ISL_16714800 | VIC, Royal Children's Hospital Victoria Australia | WHO Collaborating Centre for Reference and Research on Influenza | Xiaomin Dong, Steven Edwards, Yi-Mo Deng, Ammar Aziz, Ian Barr |
| EPI_ISL_16714803, EPI_ISL_16714809, EPI_ISL_16714810 | NT, Royal Darwin Hospital | WHO Collaborating Centre for Reference and Research on Influenza | Xiaomin Dong, Steven Edwards, Yi-Mo Deng, Ammar Aziz, Ian Barr |
| EPI_ISL_16839032, EPI_ISL_16839033, EPI_ISL_16839034, EPI_ISL_16839035, EPI_ISL_16839036, EPI_ISL_16839037, EPI_ISL_16839038 | West of Scotland Specialist Virology Centre | West of Scotland Specialist Virology Centre | Lynne Ferguson, Imogen Johnston-Menzies, Rory Gunson |
| EPI_ISL_16905445, EPI_ISL_16905447, EPI_ISL_16905448 | National Institute of Hygiene, virology departement, National Influenza Center | Centers for Disease Control and Prevention, Coronavirus and Other Respiratory Viruses Division | Megha Aggarwal, Lijuan Wang, Ji In Park, Amanda Smith and Everardo Vega |
| EPI_ISL_16959495, EPI_ISL_16959500, EPI_ISL_16959507, EPI_ISL_16959514, EPI_ISL_16959522, EPI_ISL_16959528 | ESR - Wallaceville | Institute of Environmental Science and Research | Klarysse Berquist, Lauren Jelley, Una Ren, Meaghan O'Neill |
| EPI_ISL_16959530, EPI_ISL_16959531, EPI_ISL_16959532, EPI_ISL_16959533, EPI_ISL_16959534, EPI_ISL_16959535, EPI_ISL_16959536, EPI_ISL_16959537, EPI_ISL_16959538, EPI_ISL_16959539, EPI_ISL_16959540, EPI_ISL_16959541, EPI_ISL_16959542, EPI_ISL_16959543, EPI_ISL_16959545, EPI_ISL_16959546, EPI_ISL_16959547, EPI_ISL_16959548, EPI_ISL_16959549, EPI_ISL_16959550, EPI_ISL_16959563, EPI_ISL_16959564, EPI_ISL_16959567, EPI_ISL_16959568, EPI_ISL_16959569, EPI_ISL_16959571, EPI_ISL_16959572, EPI_ISL_16959574, EPI_ISL_16959575, EPI_ISL_16959576, EPI_ISL_16959577, EPI_ISL_16959579, EPI_ISL_16959580, EPI_ISL_16959581, EPI_ISL_16959582, EPI_ISL_16959583, EPI_ISL_16959584, EPI_ISL_16959585, EPI_ISL_16959587, EPI_ISL_16959588, EPI_ISL_16959591, EPI_ISL_16959594 | | | |
| see above | Middlemore Hospital | Institute of Environmental Science and Research | Klarysse Berquist, Lauren Jelley, Una Ren, Meaghan O'Neill |
| EPI_ISL_16959595, EPI_ISL_16959597 | PathLab Bay of Plenty | Institute of Environmental Science and Research | Klarysse Berquist, Lauren Jelley, Una Ren, Meaghan O'Neill |
| EPI_ISL_16959598, EPI_ISL_16959600, EPI_ISL_16959603, EPI_ISL_16959604, EPI_ISL_16959605, EPI_ISL_16959606, EPI_ISL_16959607, EPI_ISL_16959608, EPI_ISL_16959609, EPI_ISL_16959612, EPI_ISL_16959613, EPI_ISL_16959615, EPI_ISL_16959616, EPI_ISL_16959617, EPI_ISL_16959618, EPI_ISL_16959619, EPI_ISL_16959620, EPI_ISL_16959622, EPI_ISL_16959625, EPI_ISL_16959626, EPI_ISL_16959627, EPI_ISL_16959628, EPI_ISL_16959629, EPI_ISL_16959630, EPI_ISL_16959632, EPI_ISL_16959634, EPI_ISL_16959635 | | | |
| see above | Middlemore Hospital | Institute of Environmental Science and Research | Klarysse Berquist, Lauren Jelley, Una Ren, Meaghan O'Neill |
| EPI_ISL_16959636, EPI_ISL_16959637, EPI_ISL_16959638, EPI_ISL_16959639 | MedLab South Nelson | Institute of Environmental Science and Research | Klarysse Berquist, Lauren Jelley, Una Ren, Meaghan O'Neill |
| EPI_ISL_16959641, EPI_ISL_16959642, EPI_ISL_16959643, EPI_ISL_16959644, EPI_ISL_16959645, EPI_ISL_16959646, EPI_ISL_16959647, EPI_ISL_16959648, EPI_ISL_16959649, EPI_ISL_16959651, EPI_ISL_16959652, EPI_ISL_16959653, EPI_ISL_16959655, EPI_ISL_16959656, EPI_ISL_16959659, EPI_ISL_16959660, EPI_ISL_16959661, EPI_ISL_16959663, EPI_ISL_16959664, EPI_ISL_16959665, EPI_ISL_16959666, EPI_ISL_16959667, EPI_ISL_16959668, EPI_ISL_16959670, EPI_ISL_16959671, EPI_ISL_16959673, EPI_ISL_16959674, EPI_ISL_16959676, EPI_ISL_16959677, EPI_ISL_16959679, EPI_ISL_16959680, EPI_ISL_16959681, EPI_ISL_16959682, EPI_ISL_16959683, EPI_ISL_16959685, EPI_ISL_16959686, EPI_ISL_16959688, EPI_ISL_16959690, EPI_ISL_16959691, EPI_ISL_16959692, EPI_ISL_16959693, EPI_ISL_16959694, EPI_ISL_16959695, EPI_ISL_16959697, EPI_ISL_16959698, EPI_ISL_16959699, EPI_ISL_16959700, EPI_ISL_16959702, EPI_ISL_16959706, EPI_ISL_16959708, EPI_ISL_16959709, EPI_ISL_16959710, EPI_ISL_16959711, EPI_ISL_16959712, EPI_ISL_16959713, EPI_ISL_16959714, EPI_ISL_16959715, EPI_ISL_16959718, EPI_ISL_16959719, EPI_ISL_16959720, EPI_ISL_16959721, EPI_ISL_16959722, EPI_ISL_16959724, EPI_ISL_16959725, EPI_ISL_16959727, EPI_ISL_16959728, EPI_ISL_16959730, EPI_ISL_16959731, EPI_ISL_16959732, EPI_ISL_16959733, EPI_ISL_16959734, EPI_ISL_16959735 | | | |
| see above | Middlemore Hospital | Institute of Environmental Science and Research | Klarysse Berquist, Lauren Jelley, Una Ren, Meaghan O'Neill |
| EPI_ISL_16959736, EPI_ISL_16959737, EPI_ISL_16959741, EPI_ISL_16959742 | Southern Community Lab - Dunedin | Institute of Environmental Science and Research | Klarysse Berquist, Lauren Jelley, Una Ren, Meaghan O'Neill |
| EPI_ISL_16959743, EPI_ISL_16959744, EPI_ISL_16959745, EPI_ISL_16959746, EPI_ISL_16959747, EPI_ISL_16959748, EPI_ISL_16959749, EPI_ISL_16959750, EPI_ISL_16959751, EPI_ISL_16959754, EPI_ISL_16959757, EPI_ISL_16959758, EPI_ISL_16959759, EPI_ISL_16959762, EPI_ISL_16959764, EPI_ISL_16959765, EPI_ISL_16959766, EPI_ISL_16959767, EPI_ISL_16959768, EPI_ISL_16959769, EPI_ISL_16959770, EPI_ISL_16959772, EPI_ISL_16959773, EPI_ISL_16959774, EPI_ISL_16959775, EPI_ISL_16959776, EPI_ISL_16959777, EPI_ISL_16959778, EPI_ISL_16959779, EPI_ISL_16959780, EPI_ISL_16959781, EPI_ISL_16959782 | | | |
| see above | Middlemore Hospital | Institute of Environmental Science and Research | Klarysse Berquist, Lauren Jelley, Una Ren, Meaghan O'Neill |

| | | | |
|---|---|---|---|
| EPI_ISL_16959783, EPI_ISL_16959788, EPI_ISL_16959806 | Wellington SCL | Institute of Environmental Science and Research | Klarysse Berquist, Lauren Jelley, Una Ren, Meaghan O'Neill |
| EPI_ISL_16959818, EPI_ISL_16959819, EPI_ISL_16959820, EPI_ISL_16959821, EPI_ISL_16959822, EPI_ISL_16959829, EPI_ISL_16959831, EPI_ISL_16959832, EPI_ISL_16959833, EPI_ISL_16959834, EPI_ISL_16959835, EPI_ISL_16959836 | | | |
| see above | Middlemore Hospital | Institute of Environmental Science and Research | Klarysse Berquist, Lauren Jelley, Una Ren, Meaghan O'Neill |
| EPI_ISL_16959846, EPI_ISL_16959847, EPI_ISL_16959849, EPI_ISL_16959851 | Canterbury Health Laboratory | Institute of Environmental Science and Research | Klarysse Berquist, Lauren Jelley, Una Ren, Meaghan O'Neill |
| EPI_ISL_16959852, EPI_ISL_16959856 | Labplus | Institute of Environmental Science and Research | Klarysse Berquist, Lauren Jelley, Una Ren, Meaghan O'Neill |
| EPI_ISL_16959858, EPI_ISL_16959860, EPI_ISL_16959861, EPI_ISL_16959862, EPI_ISL_16959863, EPI_ISL_16959866 | Southern Community Lab - Dunedin | Institute of Environmental Science and Research | Klarysse Berquist, Lauren Jelley, Una Ren, Meaghan O'Neill |
| EPI_ISL_16959871, EPI_ISL_16959872, EPI_ISL_16959873 | Middlemore Hospital | Institute of Environmental Science and Research | Klarysse Berquist, Lauren Jelley, Una Ren, Meaghan O'Neill |
| EPI_ISL_16959879, EPI_ISL_16959881, EPI_ISL_16959882, EPI_ISL_16959883, EPI_ISL_16959885, EPI_ISL_16959886, EPI_ISL_16959887, EPI_ISL_16959889 | Labplus | Institute of Environmental Science and Research | Klarysse Berquist, Lauren Jelley, Una Ren, Meaghan O'Neill |
| EPI_ISL_16959905 | Wellington SCL | Institute of Environmental Science and Research | Klarysse Berquist, Lauren Jelley, Una Ren, Meaghan O'Neill |
| EPI_ISL_16959918, EPI_ISL_16959919, EPI_ISL_16959922, EPI_ISL_16959923, EPI_ISL_16959924, EPI_ISL_16959926, EPI_ISL_16959927 | PathLab Bay of Plenty | Institute of Environmental Science and Research | Klarysse Berquist, Lauren Jelley, Una Ren, Meaghan O'Neill |
| EPI_ISL_16959928 | Canterbury Health Laboratory | Institute of Environmental Science and Research | Klarysse Berquist, Lauren Jelley, Una Ren, Meaghan O'Neill |
| EPI_ISL_16959942, EPI_ISL_16959943, EPI_ISL_16959945 | Middlemore Hospital | Institute of Environmental Science and Research | Klarysse Berquist, Lauren Jelley, Una Ren, Meaghan O'Neill |
| EPI_ISL_16959946, EPI_ISL_16959948, EPI_ISL_16959949, EPI_ISL_16959951, EPI_ISL_16959953, EPI_ISL_16959954, EPI_ISL_16959957 | LabPlus | Institute of Environmental Science and Research | Klarysse Berquist, Lauren Jelley, Una Ren, Meaghan O'Neill |
| EPI_ISL_16959959, EPI_ISL_16959960, EPI_ISL_16959961, EPI_ISL_16959962, EPI_ISL_16959963, EPI_ISL_16959964, EPI_ISL_16959965, EPI_ISL_16959966, EPI_ISL_16959969, EPI_ISL_16959970, EPI_ISL_16959971, EPI_ISL_16959972, EPI_ISL_16959973, EPI_ISL_16959975, EPI_ISL_16959976, EPI_ISL_16959977, EPI_ISL_16959978, EPI_ISL_16959979, EPI_ISL_16959980, EPI_ISL_16959981, EPI_ISL_16959982, EPI_ISL_16959983, EPI_ISL_16959985 | | | |
| see above | MedLab South Nelson | Institute of Environmental Science and Research | Klarysse Berquist, Lauren Jelley, Una Ren, Meaghan O'Neill |
| EPI_ISL_16959995, EPI_ISL_16959997 | Southern Community Lab - Dunedin | Institute of Environmental Science and Research | Klarysse Berquist, Lauren Jelley, Una Ren, Meaghan O'Neill |
| EPI_ISL_16959999, EPI_ISL_16960002, EPI_ISL_16960003, EPI_ISL_16960005, EPI_ISL_16960008 | Waikato Hospital | Institute of Environmental Science and Research | Klarysse Berquist, Lauren Jelley, Una Ren, Meaghan O'Neill |
| EPI_ISL_16960010, EPI_ISL_16960011, EPI_ISL_16960012, EPI_ISL_16960013 | Middlemore Hospital | Institute of Environmental Science and Research | Klarysse Berquist, Lauren Jelley, Una Ren, Meaghan O'Neill |
| EPI_ISL_16960021, EPI_ISL_16960023, EPI_ISL_16960024, EPI_ISL_16960025, EPI_ISL_16960026, EPI_ISL_16960029, EPI_ISL_16960030, EPI_ISL_16960032, EPI_ISL_16960034, EPI_ISL_16960036, EPI_ISL_16960037, EPI_ISL_16960038, EPI_ISL_16960039, EPI_ISL_16960042, EPI_ISL_16960045, EPI_ISL_16960046, EPI_ISL_16960047, EPI_ISL_16960048, EPI_ISL_16960049, EPI_ISL_16960050, EPI_ISL_16960052, EPI_ISL_16960054, EPI_ISL_16960055, EPI_ISL_16960056, EPI_ISL_16960057, EPI_ISL_16960058, EPI_ISL_16960059, EPI_ISL_16960061, EPI_ISL_16960062, EPI_ISL_16960064, EPI_ISL_16960065, EPI_ISL_16960066, EPI_ISL_16960068, EPI_ISL_16960070, EPI_ISL_16960071, EPI_ISL_16960072, EPI_ISL_16960073, EPI_ISL_16960074, EPI_ISL_16960076, EPI_ISL_16960077, EPI_ISL_16960081, EPI_ISL_16960082, EPI_ISL_16960084, EPI_ISL_16960085, EPI_ISL_16960086, EPI_ISL_16960088, EPI_ISL_16960089 | | | |
| see above | LabPlus | Institute of Environmental Science and Research | Klarysse Berquist, Lauren Jelley, Una Ren, Meaghan O'Neill |
| EPI_ISL_16960099, EPI_ISL_16960100, EPI_ISL_16960101 | Middlemore Hospital | Institute of Environmental Science and Research | Klarysse Berquist, Lauren Jelley, Una Ren, Meaghan O'Neill |
| EPI_ISL_16960103 | Wellington SCL | Institute of Environmental Science and Research | Klarysse Berquist, Lauren Jelley, Una Ren, Meaghan O'Neill |
| EPI_ISL_16960111, EPI_ISL_16960112, EPI_ISL_16960114, EPI_ISL_16960115 | Middlemore Hospital | Institute of Environmental Science and Research | Klarysse Berquist, Lauren Jelley, Una Ren, Meaghan O'Neill |
| EPI_ISL_16960118, EPI_ISL_16960125, EPI_ISL_16960129 | Southern Community Lab - Dunedin | Institute of Environmental Science and Research | Klarysse Berquist, Lauren Jelley, Una Ren, Meaghan O'Neill |
| EPI_ISL_16960130, EPI_ISL_16960131, EPI_ISL_16960132, EPI_ISL_16960136, EPI_ISL_16960139 | LabPlus | Institute of Environmental Science and Research | Klarysse Berquist, Lauren Jelley, Una Ren, Meaghan O'Neill |
| EPI_ISL_16960149 | Canterbury Health Laboratory | Institute of Environmental Science and Research | Klarysse Berquist, Lauren Jelley, Una Ren, Meaghan O'Neill |
| EPI_ISL_16960152 | Middlemore Hospital | Institute of Environmental Science and Research | Klarysse Berquist, Lauren Jelley, Una Ren, Meaghan O'Neill |
| EPI_ISL_16982626, EPI_ISL_16982627, EPI_ISL_16982628, EPI_ISL_16982629, EPI_ISL_16982630, EPI_ISL_16982631, EPI_ISL_16982632, EPI_ISL_16982633 | Chinese Center for Disease Control and Prevention | Chinese Center for Disease Control and Prevention | Jiang,Y. |
| EPI_ISL_17066776, EPI_ISL_17066777 | VIC, Royal Children's Hospital Victoria Australia | WHO Collaborating Centre for Reference and Research on Influenza | Xiaomin Dong, Steven Edwards, Yi-Mo Deng, Ammar Aziz, Ian Barr |
| EPI_ISL_17066784 | NT, Royal Darwin Hospital | WHO Collaborating Centre for Reference and Research on Influenza | Xiaomin Dong, Steven Edwards, Yi-Mo Deng, Ammar Aziz, Ian Barr |
| EPI_ISL_17066788, EPI_ISL_17066793, EPI_ISL_17066795, EPI_ISL_17066802, EPI_ISL_17066813, EPI_ISL_17066817 | QLD, Pathology Queensland - Royal Brisbane and Women's Hospital | WHO Collaborating Centre for Reference and Research on Influenza | Xiaomin Dong, Steven Edwards, Yi-Mo Deng, Ammar Aziz, Ian Barr |
| EPI_ISL_17066819, EPI_ISL_17066821 | NT, Royal Darwin Hospital | WHO Collaborating Centre for Reference and Research on Influenza | Xiaomin Dong, Steven Edwards, Yi-Mo Deng, Ammar Aziz, Ian Barr |
| EPI_ISL_17066824, EPI_ISL_17066827, EPI_ISL_17066828, EPI_ISL_17066831, EPI_ISL_17066833, EPI_ISL_17066834 | VIC, Royal Children's Hospital Victoria Australia | WHO Collaborating Centre for Reference and Research on Influenza | Xiaomin Dong, Steven Edwards, Yi-Mo Deng, Ammar Aziz, Ian Barr |
| EPI_ISL_17089183, EPI_ISL_17089184 | University of Washington - Laboratory Medicine | University of Washington - Laboratory Medicine | Sereewit,J., Xie,H., Roychoudhury,P. and Greninger,A.L. |
| EPI_ISL_17127009, EPI_ISL_17127010 | Servicio de Microbiologia, Hospital Universitario Virgen del Rocio | Plataforma de Medicina Computacional, Fundación Progreso y Salud | José Antonio Lepe, Javier Pérez-Florido, María Lara Jiménez |
| EPI_ISL_17221671, EPI_ISL_17221674 | University of Washington - Virology | University of Washington - Virology | Sereewit,J., Xie,H., Roychoudhury,P. and Greninger,A.L. |
| EPI_ISL_17253616, EPI_ISL_17253618, EPI_ISL_17253625 | National Public Health Institute of Slovakia | Laboratory of Genomics and Bioinformatics, Comenius University Science Park | Szemes,Tomáš;Kaliňáková,Anna;Kotvasová,Barbora;Ševčíková,Lucia;Vrabľová,Terézia;Sedláčková,Tatiana;Rusňáková,Diana;Böhmer,Miroslav;Budiš,Jaroslav;Styk,Jakub;Lipková,Nikola;Forgáčová,Michaela;Bokorová,Silvia;Mišenko,Pavol |
| EPI_ISL_17308687, EPI_ISL_17308688, EPI_ISL_17308689 | Laboratório de Virologia - Instituto Nacional de Saúde | Centers for Disease Control and Prevention - United States of America, Laboratório de Virologia - Instituto Nacional de Saúde | Almiro Tivane, Everardo Vega, Amanda Smith, Lijuan Wang, Neuza Nguenha, Loira Machalele, Sadia Alí, Délcio Muteto, Mirela Pale, Aunésia Marrurele, Félix Gundane, Judite Salência |
| EPI_ISL_17308690 | Laboratório de Virologia - Instituto Nacional de Saúde | Laboratório de Vírus Respiratórios e do Sarampo - Instituto Oswaldo Cruz, Laboratório de Virologia - Instituto Nacional de Saúde | Almiro Tivane, Paola Resende, Neuza Nguenha, Loira Machalele, Sadia Alí, Délcio Muteto, Aunésia Marrurele, Mirela Pale, Félix Gundane, Judite Salência, Marilda Siqueira |
| EPI_ISL_17308691, EPI_ISL_17308692 | Laboratório de Virologia - Instituto Nacional de Saúde | Centers for Disease Control and Prevention - United States of America, Laboratório de Virologia - Instituto Nacional de Saúde | Almiro Tivane, Everardo Vega, Amanda Smith, Lijuan Wang, Neuza Nguenha, Loira Machalele, Sadia Alí, Délcio Muteto, Mirela Pale, Aunésia Marrurele, Félix Gundane, Judite Salência |
| EPI_ISL_17308693, EPI_ISL_17308694 | Laboratório de Virologia - Instituto Nacional de Saúde | Laboratório de Vírus Respiratórios e do Sarampo - Instituto Oswaldo Cruz, Laboratório de Virologia - Instituto Nacional de Saúde | Almiro Tivane, Paola Resende, Neuza Nguenha, Loira Machalele, Sadia Alí, Délcio Muteto, Aunésia Marrurele, Mirela Pale, Félix Gundane, Judite Salência, Marilda Siqueira |
| EPI_ISL_17308695 | Laboratório de Virologia - Instituto Nacional de Saúde | Centers for Disease Control and Prevention - United States of America, Laboratório de Virologia - Instituto Nacional de Saúde | Almiro Tivane, Everardo Vega, Amanda Smith, Lijuan Wang, Neuza Nguenha, Loira Machalele, Sadia Alí, Délcio Muteto, Mirela Pale, Aunésia Marrurele, Félix Gundane, Judite Salência |
| EPI_ISL_17308696, EPI_ISL_17308697 | Laboratório de Virologia - Instituto Nacional de Saúde | Laboratório de Vírus Respiratórios e do Sarampo - Instituto Oswaldo Cruz, Laboratório de Virologia - Instituto Nacional de Saúde | Almiro Tivane, Paola Resende, Neuza Nguenha, Loira Machalele, Sadia Alí, Délcio Muteto, Aunésia Marrurele, Mirela Pale, Félix Gundane, Judite Salência, Marilda Siqueira |
| EPI_ISL_17308698, EPI_ISL_17308699, EPI_ISL_17308700, EPI_ISL_17308701 | Laboratório de Virologia - Instituto Nacional de Saúde | Centers for Disease Control and Prevention - United States of America, Laboratório de Virologia - Instituto Nacional de Saúde | Almiro Tivane, Everardo Vega, Amanda Smith, Lijuan Wang, Neuza Nguenha, Loira Machalele, Sadia Alí, Délcio Muteto, Mirela Pale, Aunésia Marrurele, Félix Gundane, Judite Salência |
| EPI_ISL_17308702, EPI_ISL_17308703, EPI_ISL_17308704, EPI_ISL_17308705, EPI_ISL_17308706 | Laboratório de Virologia - Instituto Nacional de Saúde | Laboratório de Vírus Respiratórios e do Sarampo - Instituto Oswaldo Cruz, Laboratório de Virologia - Instituto Nacional de Saúde | Almiro Tivane, Paola Resende, Neuza Nguenha, Loira Machalele, Sadia Alí, Délcio Muteto, Aunésia Marrurele, Mirela Pale, Félix Gundane, Judite Salência, Marilda Siqueira |
| EPI_ISL_17308707 | Laboratório de Virologia - Instituto Nacional de Saúde | Centers for Disease Control and Prevention - United States of America, Laboratório de Virologia - Instituto Nacional de Saúde | Almiro Tivane, Everardo Vega, Amanda Smith, Lijuan Wang, Neuza Nguenha, Loira Machalele, Sadia Alí, Délcio Muteto, Mirela Pale, Aunésia Marrurele, Félix Gundane, Judite Salência |
| EPI_ISL_17308708 | Laboratório de Virologia - Instituto Nacional de | Laboratório de Vírus Respiratórios e do Sarampo - | Almiro Tivane, Paola Resende, Neuza Nguenha, Loira Machalele, Sadia Alí, Délcio Muteto, Aunésia Marrurele, Mirela Pale, Félix Gundane, Judite Salência, Marilda Siqueira |

| | | | |
|---|---|---|---|
| | Saúde | Instituto Oswaldo Cruz, Laboratório de Virologia - Instituto Nacional de Saúde | |
| EPI_ISL_17308709 | Laboratório de Virologia - Instituto Nacional de Saúde | Centers for Disease Control and Prevention - United States of America, Laboratório de Virologia - Instituto Nacional de Saúde | Almiro Tivane, Everardo Vega, Amanda Smith, Lijuan Wang, Neuza Nguenha, Loira Machalele, Sadia Alí, Délcio Muteto, Mirela Pale, Aunésia Marrurele, Félix Gundane, Judite Salência |
| EPI_ISL_17308710 | Laboratório de Virologia - Instituto Nacional de Saúde | Laboratório de Vírus Respiratórios e do Sarampo - Instituto Oswaldo Cruz, Laboratório de Virologia - Instituto Nacional de Saúde | Almiro Tivane, Paola Resende, Neuza Nguenha, Loira Machalele, Sadia Alí, Délcio Muteto, Aunésia Marrurele, Mirela Pale, Félix Gundane, Judite Salência, Marilda Siqueira |
| EPI_ISL_17308711, EPI_ISL_17308712 | Laboratório de Virologia - Instituto Nacional de Saúde | Centers for Disease Control and Prevention - United States of America, Laboratório de Virologia - Instituto Nacional de Saúde | Almiro Tivane, Everardo Vega, Amanda Smith, Lijuan Wang, Neuza Nguenha, Loira Machalele, Sadia Alí, Délcio Muteto, Mirela Pale, Aunésia Marrurele, Félix Gundane, Judite Salência |
| EPI_ISL_17308713 | Laboratório de Virologia - Instituto Nacional de Saúde | Laboratório de Vírus Respiratórios e do Sarampo - Instituto Oswaldo Cruz, Laboratório de Virologia - Instituto Nacional de Saúde | Almiro Tivane, Paola Resende, Neuza Nguenha, Loira Machalele, Sadia Alí, Délcio Muteto, Aunésia Marrurele, Mirela Pale, Félix Gundane, Judite Salência, Marilda Siqueira |
| EPI_ISL_17308714, EPI_ISL_17308715, EPI_ISL_17308716 | Laboratório de Virologia - Instituto Nacional de Saúde | Centers for Disease Control and Prevention - United States of America, Laboratório de Virologia - Instituto Nacional de Saúde | Almiro Tivane, Everardo Vega, Amanda Smith, Lijuan Wang, Neuza Nguenha, Loira Machalele, Sadia Alí, Délcio Muteto, Mirela Pale, Aunésia Marrurele, Félix Gundane, Judite Salência |
| EPI_ISL_17308717, EPI_ISL_17308718, EPI_ISL_17308719, EPI_ISL_17308720, EPI_ISL_17308721, EPI_ISL_17308722, EPI_ISL_17308723 | Laboratório de Virologia - Instituto Nacional de Saúde | Laboratório de Vírus Respiratórios e do Sarampo - Instituto Oswaldo Cruz, Laboratório de Virologia - Instituto Nacional de Saúde | Almiro Tivane, Paola Resende, Neuza Nguenha, Loira Machalele, Sadia Alí, Délcio Muteto, Aunésia Marrurele, Mirela Pale, Félix Gundane, Judite Salência, Marilda Siqueira |
| EPI_ISL_17308724, EPI_ISL_17308725 | Laboratório de Virologia - Instituto Nacional de Saúde | Centers for Disease Control and Prevention - United States of America, Laboratório de Virologia - Instituto Nacional de Saúde | Almiro Tivane, Everardo Vega, Amanda Smith, Lijuan Wang, Neuza Nguenha, Loira Machalele, Sadia Alí, Délcio Muteto, Mirela Pale, Aunésia Marrurele, Félix Gundane, Judite Salência |
| EPI_ISL_17308726, EPI_ISL_17308727 | Laboratório de Virologia - Instituto Nacional de Saúde | Laboratório de Vírus Respiratórios e do Sarampo - Instituto Oswaldo Cruz, Laboratório de Virologia - Instituto Nacional de Saúde | Almiro Tivane, Paola Resende, Neuza Nguenha, Loira Machalele, Sadia Alí, Délcio Muteto, Aunésia Marrurele, Mirela Pale, Félix Gundane, Judite Salência, Marilda Siqueira |
| EPI_ISL_17308728 | Laboratório de Virologia - Instituto Nacional de Saúde | Centers for Disease Control and Prevention - United States of America, Laboratório de Virologia - Instituto Nacional de Saúde | Almiro Tivane, Everardo Vega, Amanda Smith, Lijuan Wang, Neuza Nguenha, Loira Machalele, Sadia Alí, Délcio Muteto, Mirela Pale, Aunésia Marrurele, Félix Gundane, Judite Salência |
| EPI_ISL_17368037, EPI_ISL_17368038, EPI_ISL_17368039, EPI_ISL_17368040 | Edendale | National Institute for Communicable Diseases of the National Health Laboratory Service | Everatt J, Kekana D, Mahlangu B, Stock N, Ntozini B, Ntuli N, Mnguni A, Nzimande A, Ismail A, Bhiman JN, Wolter N |
| EPI_ISL_17368041, EPI_ISL_17368042 | Klerksdorp | National Institute for Communicable Diseases of the National Health Laboratory Service | Everatt J, Kekana D, Mahlangu B, Stock N, Ntozini B, Ntuli N, Mnguni A, Nzimande A, Ismail A, Bhiman JN, Wolter N |
| EPI_ISL_17368043, EPI_ISL_17368044 | Mapulaneng | National Institute for Communicable Diseases of the National Health Laboratory Service | Everatt J, Kekana D, Mahlangu B, Stock N, Ntozini B, Ntuli N, Mnguni A, Nzimande A, Ismail A, Bhiman JN, Wolter N |
| EPI_ISL_17417578, EPI_ISL_17417579, EPI_ISL_17417580, EPI_ISL_17417581, EPI_ISL_17417582, EPI_ISL_17417583, EPI_ISL_17417584, EPI_ISL_17417585, EPI_ISL_17417586, EPI_ISL_17417589, EPI_ISL_17417591, EPI_ISL_17417592, EPI_ISL_17417594, EPI_ISL_17417595, EPI_ISL_17417596, EPI_ISL_17417597, EPI_ISL_17417598, EPI_ISL_17417599, EPI_ISL_17417600, EPI_ISL_17417601, EPI_ISL_17417602, EPI_ISL_17417603, EPI_ISL_17417604 | | | |
| see above | Centers for Disease Control and Prevention (CDC) | Centers for Disease Control and Prevention (CDC) | Wang,L., Lumandas,M.U., Arguelles,V.L., Morin,J., Aggarwal,M., Tatusov,R. and Vega,E. |
| EPI_ISL_17481634, EPI_ISL_17481635, EPI_ISL_17481643, EPI_ISL_17481644, EPI_ISL_17481646, EPI_ISL_17481661 | National Public Health Institute of Slovakia | Laboratory of Genomics and Bioinformatics, Comenius University Science Park | Szemes,Tomas; Kalinakova,Anna; Kotvasova,Barbora; Sevcikova,Lucia; Vrablova,Terezia; Sedlackova,Tatiana; Rusnakova,Diana; Bohmer,Miroslav; Budis,Jaroslav; Styk,Jakub; Lipkova,Nikola; Forgacova,Michaela; Bokorova,Silvia; Misenko,Pavol |
| EPI_ISL_17508474 | University of Washington | University of Washington | Greninger,A.L., Makhsous,N., Kuypers,J., Shean,R.C. and Jerome,K.R. |
| EPI_ISL_17559319 | Red Cross | National Institute for Communicable Diseases of the National Health Laboratory Service | Everatt J, Kekana D, Mahlangu B, Stock N, Ntozini B, Ntuli N, Mnguni A, Nzimande A, Ismail A, Bhiman JN, Wolter N |
| EPI_ISL_17559321 | East ridge Clinic | National Institute for Communicable Diseases of the National Health Laboratory Service | Everatt J, Kekana D, Mahlangu B, Stock N, Ntozini B, Ntuli N, Mnguni A, Nzimande A, Ismail A, Bhiman JN, Wolter N |
| EPI_ISL_17559323 | Edendale | National Institute for Communicable Diseases of the National Health Laboratory Service | Everatt J, Kekana D, Mahlangu B, Stock N, Ntozini B, Ntuli N, Mnguni A, Nzimande A, Ismail A, Bhiman JN, Wolter N |
| EPI_ISL_17559324, EPI_ISL_17559327 | Rahima Moosa | National Institute for Communicable Diseases of the National Health Laboratory Service | Everatt J, Kekana D, Mahlangu B, Stock N, Ntozini B, Ntuli N, Mnguni A, Nzimande A, Ismail A, Bhiman JN, Wolter N |
| EPI_ISL_17559328, EPI_ISL_17559329 | Red Cross | National Institute for Communicable Diseases of the National Health Laboratory Service | Everatt J, Kekana D, Mahlangu B, Stock N, Ntozini B, Ntuli N, Mnguni A, Nzimande A, Ismail A, Bhiman JN, Wolter N |
| EPI_ISL_17559330, EPI_ISL_17559333, EPI_ISL_17559334 | Rahima Moosa | National Institute for Communicable Diseases of the National Health Laboratory Service | Everatt J, Kekana D, Mahlangu B, Stock N, Ntozini B, Ntuli N, Mnguni A, Nzimande A, Ismail A, Bhiman JN, Wolter N |
| EPI_ISL_17559337 | East ridge Clinic | National Institute for Communicable Diseases of the National Health Laboratory Service | Everatt J, Kekana D, Mahlangu B, Stock N, Ntozini B, Ntuli N, Mnguni A, Nzimande A, Ismail A, Bhiman JN, Wolter N |
| EPI_ISL_17559339, EPI_ISL_17559341, EPI_ISL_17559342, EPI_ISL_17559343 | Mitchells Plain | National Institute for Communicable Diseases of the National Health Laboratory Service | Everatt J, Kekana D, Mahlangu B, Stock N, Ntozini B, Ntuli N, Mnguni A, Nzimande A, Ismail A, Bhiman JN, Wolter N |
| EPI_ISL_17559344, EPI_ISL_17559345, EPI_ISL_17559346, EPI_ISL_17559347 | Red Cross | National Institute for Communicable Diseases of the National Health Laboratory Service | Everatt J, Kekana D, Mahlangu B, Stock N, Ntozini B, Ntuli N, Mnguni A, Nzimande A, Ismail A, Bhiman JN, Wolter N |
| EPI_ISL_17559349, EPI_ISL_17559352, EPI_ISL_17559354, EPI_ISL_17559355, EPI_ISL_17559363, EPI_ISL_17559364 | Rahima Moosa | National Institute for Communicable Diseases of the National Health Laboratory Service | Everatt J, Kekana D, Mahlangu B, Stock N, Ntozini B, Ntuli N, Mnguni A, Nzimande A, Ismail A, Bhiman JN, Wolter N |
| EPI_ISL_17559366, EPI_ISL_17559367, EPI_ISL_17559369 | Red Cross | National Institute for Communicable Diseases of the National Health Laboratory Service | Everatt J, Kekana D, Mahlangu B, Stock N, Ntozini B, Ntuli N, Mnguni A, Nzimande A, Ismail A, Bhiman JN, Wolter N |
| EPI_ISL_17559371 | Rahima Moosa | National Institute for Communicable Diseases of the National Health Laboratory Service | Everatt J, Kekana D, Mahlangu B, Stock N, Ntozini B, Ntuli N, Mnguni A, Nzimande A, Ismail A, Bhiman JN, Wolter N |
| EPI_ISL_17559372 | Mitchells Plain | National Institute for Communicable Diseases of the National Health Laboratory Service | Everatt J, Kekana D, Mahlangu B, Stock N, Ntozini B, Ntuli N, Mnguni A, Nzimande A, Ismail A, Bhiman JN, Wolter N |
| EPI_ISL_17559373 | East ridge Clinic | National Institute for Communicable Diseases of the National Health Laboratory Service | Everatt J, Kekana D, Mahlangu B, Stock N, Ntozini B, Ntuli N, Mnguni A, Nzimande A, Ismail A, Bhiman JN, Wolter N |
| EPI_ISL_17559374, EPI_ISL_17559375 | Mitchells Plain | National Institute for Communicable Diseases of the National Health Laboratory Service | Everatt J, Kekana D, Mahlangu B, Stock N, Ntozini B, Ntuli N, Mnguni A, Nzimande A, Ismail A, Bhiman JN, Wolter N |
| EPI_ISL_17559378 | Red Cross | National Institute for Communicable Diseases of the National Health Laboratory Service | Everatt J, Kekana D, Mahlangu B, Stock N, Ntozini B, Ntuli N, Mnguni A, Nzimande A, Ismail A, Bhiman JN, Wolter N |
| EPI_ISL_17559379 | Rahima Moosa | National Institute for Communicable Diseases of the National Health Laboratory Service | Everatt J, Kekana D, Mahlangu B, Stock N, Ntozini B, Ntuli N, Mnguni A, Nzimande A, Ismail A, Bhiman JN, Wolter N |
| EPI_ISL_17559380 | Edendale Gateway Clinic | National Institute for Communicable Diseases of the National Health Laboratory Service | Everatt J, Kekana D, Mahlangu B, Stock N, Ntozini B, Ntuli N, Mnguni A, Nzimande A, Ismail A, Bhiman JN, Wolter N |
| EPI_ISL_17559385 | Red Cross | National Institute for Communicable Diseases of the National Health Laboratory Service | Everatt J, Kekana D, Mahlangu B, Stock N, Ntozini B, Ntuli N, Mnguni A, Nzimande A, Ismail A, Bhiman JN, Wolter N |
| EPI_ISL_17559387, EPI_ISL_17559388, EPI_ISL_17559390, EPI_ISL_17559391, EPI_ISL_17559393, EPI_ISL_17559395 | Rahima Moosa | National Institute for Communicable Diseases of the National Health Laboratory Service | Everatt J, Kekana D, Mahlangu B, Stock N, Ntozini B, Ntuli N, Mnguni A, Nzimande A, Ismail A, Bhiman JN, Wolter N |
| EPI_ISL_17559399 | East ridge Clinic | National Institute for Communicable Diseases of the National Health Laboratory Service | Everatt J, Kekana D, Mahlangu B, Stock N, Ntozini B, Ntuli N, Mnguni A, Nzimande A, Ismail A, Bhiman JN, Wolter N |
| EPI_ISL_17559403 | Tintswalo | National Institute for Communicable Diseases of the National Health Laboratory Service | Everatt J, Kekana D, Mahlangu B, Stock N, Ntozini B, Ntuli N, Mnguni A, Nzimande A, Ismail A, Bhiman JN, Wolter N |
| EPI_ISL_17559466, EPI_ISL_17559467, EPI_ISL_17559468 | Laboratory of Respiratory Viruses and Measles, Oswaldo Cruz Institute, FIOCRUZ | Laboratorio de Alta Complexidade do Instituto Fernandes Figueira - LACIFF | Paola Resende, Fernando Motta, Elisa Cavalcante Pereira, Bruna Mendonça da Silva, Jéssica Graça Macedo de Carvalho, Larissa Macedo Pinto, Victor Guimaraes, Letícia Lima, Letícia Scalioni, Marilda Siqueira on behalf of the Fiocruz COVID-19 Genomic Surveillance Network |
| EPI_ISL_17583025, EPI_ISL_17583026, EPI_ISL_17583027, EPI_ISL_17583028, EPI_ISL_17583029, EPI_ISL_17583030, EPI_ISL_17583663, EPI_ISL_17583664, EPI_ISL_17583665, EPI_ISL_17583666, EPI_ISL_17583667, EPI_ISL_17583668, EPI_ISL_17583669, EPI_ISL_17583670 | | | |
| see above | West of Scotland Specialist Virology Centre | West of Scotland Specialist Virology Centre | Lynne Ferguson, Imogen Johnston-Menzies, Rory Gunson |
| EPI_ISL_17673340, EPI_ISL_17673341, EPI_ISL_17673342, EPI_ISL_17673343, EPI_ISL_17673344, EPI_ISL_17673345, EPI_ISL_17673346, EPI_ISL_17673347, EPI_ISL_17673348, EPI_ISL_17673349, EPI_ISL_17673350, EPI_ISL_17673351, EPI_ISL_17673352, EPI_ISL_17673353 | | | |
| see above | University of Washington, Virology | University of Washington, Virology | Sereewit,J., Hajian,P., Xie,H. and Greninger,A.L. |

| | | | |
|---|---|---|---|
| EPI_ISL_17773648, EPI_ISL_17773649, EPI_ISL_17773650, EPI_ISL_17773652, EPI_ISL_17773653, EPI_ISL_17773654, EPI_ISL_17773655, EPI_ISL_17773656, EPI_ISL_17773657, EPI_ISL_17773658, EPI_ISL_17773660, EPI_ISL_17773661 | | | |
| see above | Laboratorio Central de Saude Publica do Distrito Federal (LACEN/DF) | Laboratory of Respiratory Viruses and Measles, Oswaldo Cruz Institute, FIOCRUZ | Paola Resende, Fernando Motta, Elisa Cavalcante Pereira, Leticia Ferreira Lima, Bruna Mendonça da Silva, Jéssica Graça Macedo de Carvalho, Larissa Macedo Pinto, Victor Guimaraes, Leticia Scalioni, Rodrigo Ribeiro Rodrigues, Marilda Siqueira on behalf of the Fiocruz COVID-19 Genomic Surveillance Network |
| EPI_ISL_17773665 | Universidade Federal de São Paulo, Departamento de Medicina, Disciplina de Doenças Infecciosas e Parasitárias. Laboratório de Virologia | Laboratory of Respiratory Viruses and Measles, Oswaldo Cruz Institute, FIOCRUZ | Paola Resende, Fernando Motta, Elisa Cavalcante Pereira, Leticia Ferreira Lima, Bruna Mendonça da Silva, Jéssica Graça Macedo de Carvalho, Larissa Macedo Pinto, Victor Guimaraes, Leticia Scalioni, Rodrigo Ribeiro Rodrigues, Nancy Bellei, Marilda Siqueira on behalf of the Fiocruz COVID-19 Genomic Surveillance Network |
| EPI_ISL_17778166, EPI_ISL_17778298, EPI_ISL_17778391 | Laboratorio Central de Saude Publica do Distrito Federal (LACEN/DF) | Instituto Oswaldo Cruz FIOCRUZ - Laboratory of Respiratory Viruses and Measles (LVRS) | Paola Resende, Fernando Motta, Elisa Cavalcante Pereira, Leticia Ferreira Lima, Bruna Mendonça da Silva, Jéssica Graça Macedo de Carvalho, Larissa Macedo Pinto, Victor Guimaraes, Leticia Scalioni, Rodrigo Ribeiro Rodrigues, Marilda Siqueira on behalf of the Fiocruz COVID-19 Genomic Surveillance Network |
| EPI_ISL_17782655, EPI_ISL_17782657 | Kitasato University, Infection Control and Immunology | Kitasato University, Infection Control and Immunology | Ito,T., Sawada,A., Saito,A., Ishikura,K., Kawashima,H., Nakayama,T. and Katayama,K. |
| EPI_ISL_17782659 | University of Washington, Virology | University of Washington, Virology | Sereewit,J., Hajian,P., Xie,H. and Greninger,A.L. |
| EPI_ISL_17808729, EPI_ISL_17808730 | Central public health laboratory of the state of Amapa | Evandro Chagas Institute, Laboratory of Respiratory Viruses, National InfluenzaCenter | Mirleide Santos; Fernando Tavares; Edivaldo Junior; Luana Barbagelata; Amanda Mendes; Wanderley Dias; Delana Melo; Agata Monique; Edvaldo Penha; |
| EPI_ISL_17808737, EPI_ISL_17808739, EPI_ISL_17808740, EPI_ISL_17808741, EPI_ISL_17808742, EPI_ISL_17808743, EPI_ISL_17808744, EPI_ISL_17808745, EPI_ISL_17808746, EPI_ISL_17808747, EPI_ISL_17808748, EPI_ISL_17808749, EPI_ISL_17808750, EPI_ISL_17808751, EPI_ISL_17808752, EPI_ISL_17808753, EPI_ISL_17808754, EPI_ISL_17808755, EPI_ISL_17808756, EPI_ISL_17808757, EPI_ISL_17808758, EPI_ISL_17808759, EPI_ISL_17808760, EPI_ISL_17808761, EPI_ISL_17808762, EPI_ISL_17808763, EPI_ISL_17808764, EPI_ISL_17808765, EPI_ISL_17808766, EPI_ISL_17808767, EPI_ISL_17808768, EPI_ISL_17808769, EPI_ISL_17808770, EPI_ISL_17808771, EPI_ISL_17808773, EPI_ISL_17808774, EPI_ISL_17808775, EPI_ISL_17808776, EPI_ISL_17808777, EPI_ISL_17808778, EPI_ISL_17808779, EPI_ISL_17808780, EPI_ISL_17808781, EPI_ISL_17808782, EPI_ISL_17808783, EPI_ISL_17808784, EPI_ISL_17808785, EPI_ISL_17808786, EPI_ISL_17808787, EPI_ISL_17808788, EPI_ISL_17808789, EPI_ISL_17808790, EPI_ISL_17808792, EPI_ISL_17808793, EPI_ISL_17808794, EPI_ISL_17808795, EPI_ISL_17808797, EPI_ISL_17808798, EPI_ISL_17808799, EPI_ISL_17808800, EPI_ISL_17808802, EPI_ISL_17808803, EPI_ISL_17808804, EPI_ISL_17808805, EPI_ISL_17808806, EPI_ISL_17808807, EPI_ISL_17808808, EPI_ISL_17808809, EPI_ISL_17808810, EPI_ISL_17808811, EPI_ISL_17808812, EPI_ISL_17808814, EPI_ISL_17808815, EPI_ISL_17808816, EPI_ISL_17808817, EPI_ISL_17808819, EPI_ISL_17808820, EPI_ISL_17808822, EPI_ISL_17808824, EPI_ISL_17808826, EPI_ISL_17808827, EPI_ISL_17808828, EPI_ISL_17808831, EPI_ISL_17808832, EPI_ISL_17808834, EPI_ISL_17808835, EPI_ISL_17808836, EPI_ISL_17808837 | | | |
| see above | Valley Wise Health | Arizona State University | Holland, LaRinda A.; Holland, Steven C.; Smith, Matthew F.; Leonard, Victoria R.; Murugan, Vel; Nordstrom, Lora; Mulrow, Mary; Salgado, Raquel; White, Michael; Lim, Efrem S. |
| EPI_ISL_17950137, EPI_ISL_17950138, EPI_ISL_17950139, EPI_ISL_17950140, EPI_ISL_17950141, EPI_ISL_17950142, EPI_ISL_17950143, EPI_ISL_17950144, EPI_ISL_17950145, EPI_ISL_17950146, EPI_ISL_17950147, EPI_ISL_17950148, EPI_ISL_17950149, EPI_ISL_17950151, EPI_ISL_17950152, EPI_ISL_17950153, EPI_ISL_17950154, EPI_ISL_17950155, EPI_ISL_17950156, EPI_ISL_17950157, EPI_ISL_17950158, EPI_ISL_17950159, EPI_ISL_17950163, EPI_ISL_17950164, EPI_ISL_17950165, EPI_ISL_17950166, EPI_ISL_17950167, EPI_ISL_17950171, EPI_ISL_17950172, EPI_ISL_17950173, EPI_ISL_17950174, EPI_ISL_17950175, EPI_ISL_17950176, EPI_ISL_17950177, EPI_ISL_17950178, EPI_ISL_17950179, EPI_ISL_17950180, EPI_ISL_17950183, EPI_ISL_17950186, EPI_ISL_17950188, EPI_ISL_17950189, EPI_ISL_17950191, EPI_ISL_17950192, EPI_ISL_17950193, EPI_ISL_17950194, EPI_ISL_17950196, EPI_ISL_17950197, EPI_ISL_17950199, EPI_ISL_17950202, EPI_ISL_17950203, EPI_ISL_17950204, EPI_ISL_17950205, EPI_ISL_17950206, EPI_ISL_17950208, EPI_ISL_17950209, EPI_ISL_17950211, EPI_ISL_17950212, EPI_ISL_17950213, EPI_ISL_17950214, EPI_ISL_17950216, EPI_ISL_17950217, EPI_ISL_17950219, EPI_ISL_17950220, EPI_ISL_17950221, EPI_ISL_17950222, EPI_ISL_17950224, EPI_ISL_17950225, EPI_ISL_17950226, EPI_ISL_17950228, EPI_ISL_17950229, EPI_ISL_17950230, EPI_ISL_17950232, EPI_ISL_17950233, EPI_ISL_17950234, EPI_ISL_17950235, EPI_ISL_17950237, EPI_ISL_17950238, EPI_ISL_17950239, EPI_ISL_17950240, EPI_ISL_17950242, EPI_ISL_17950243, EPI_ISL_17950245, EPI_ISL_17950246, EPI_ISL_17950247, EPI_ISL_17950249, EPI_ISL_17950250, EPI_ISL_17950251, EPI_ISL_17950252 | | | |
| see above | Arizona State University | Arizona State University | Holland,S.C., Holland,L.A., Smith,M.F., Leonard,V.R., Murugan,V., Nordstrom,L., Mulrow,M., Salgado,R., White,M. and Lim,E.S. |
| EPI_ISL_17950996, EPI_ISL_17950997, EPI_ISL_17950998, EPI_ISL_17950999, EPI_ISL_17951000, EPI_ISL_17951001, EPI_ISL_17951002, EPI_ISL_17951003, EPI_ISL_17951004, EPI_ISL_17951005, EPI_ISL_17951006, EPI_ISL_17951007, EPI_ISL_17951008, EPI_ISL_17951009, EPI_ISL_17951010, EPI_ISL_17951011, EPI_ISL_17951012, EPI_ISL_17951013, EPI_ISL_17951014, EPI_ISL_17951015, EPI_ISL_17951016, EPI_ISL_17951017, EPI_ISL_17951018, EPI_ISL_17951019, EPI_ISL_17951020, EPI_ISL_17951021, EPI_ISL_17951022, EPI_ISL_17951023, EPI_ISL_17951024, EPI_ISL_17951025, EPI_ISL_17951026, EPI_ISL_17951027, EPI_ISL_17951028, EPI_ISL_17951029, EPI_ISL_17951030, EPI_ISL_17951031, EPI_ISL_17951032, EPI_ISL_17951033, EPI_ISL_17951034, EPI_ISL_17951035, EPI_ISL_17951036, EPI_ISL_17951037, EPI_ISL_17951038, EPI_ISL_17951039, EPI_ISL_17951040, EPI_ISL_17951041, EPI_ISL_17951042, EPI_ISL_17951043, EPI_ISL_17951044, EPI_ISL_17951045, EPI_ISL_17951046 | | | |
| see above | Molecular Biology Laboratory, Pedro de Elizalde Hospital | Virology Laboratory, Ricardo Gutiérrez Children's Hospital | Acuña, Dolores; Goya, Stephanie; Nabaes Jodar, Mercedes S.; Montoto, L; Wenk, G; Sevilla, ME; Miño, L; Valeri, C; Bokser, V; Mistchenko, Alicia S.; Viegas, Mariana |
| EPI_ISL_17964306, EPI_ISL_17964307, EPI_ISL_17964309, EPI_ISL_17964310, EPI_ISL_17964311, EPI_ISL_17964312, EPI_ISL_17964313, EPI_ISL_17964314, EPI_ISL_17964315, EPI_ISL_17964316, EPI_ISL_17964317, EPI_ISL_17964318, EPI_ISL_17964319, EPI_ISL_17964320, EPI_ISL_17964321, EPI_ISL_17964322, EPI_ISL_17964323, EPI_ISL_17964324, EPI_ISL_17964325, EPI_ISL_17964326, EPI_ISL_17964327, EPI_ISL_17964328, EPI_ISL_17964329, EPI_ISL_17964330, EPI_ISL_17964331, EPI_ISL_17964332, EPI_ISL_17964333, EPI_ISL_17964334, EPI_ISL_17964335, EPI_ISL_17964336, EPI_ISL_17964337, EPI_ISL_17964338, EPI_ISL_17964339, EPI_ISL_17964340, EPI_ISL_17964341, EPI_ISL_17964342, EPI_ISL_17964343, EPI_ISL_17964344, EPI_ISL_17964345, EPI_ISL_17964346, EPI_ISL_17964347, EPI_ISL_17964348, EPI_ISL_17964349, EPI_ISL_17964350, EPI_ISL_17964352, EPI_ISL_17964355, EPI_ISL_17964356, EPI_ISL_17964359, EPI_ISL_17964360, EPI_ISL_17964363, EPI_ISL_17964365, EPI_ISL_17964366, EPI_ISL_17964367, EPI_ISL_17964368, EPI_ISL_17964369, EPI_ISL_17964370, EPI_ISL_17964371, EPI_ISL_17964372, EPI_ISL_17964373, EPI_ISL_17964374, EPI_ISL_17964375 | | | |
| see above | University of Zambia Medical School | Department of Medical Microbiology, University Medical Centre Utrecht | Annefleur C. Langedijk, Bram Vrancken, Robert Jan Lebbink, Rachel C. Pieciak, Philippe Lemey, Louis J. Bont, Christopher J Gill |
| EPI_ISL_17992452, EPI_ISL_17992454, EPI_ISL_17992456, EPI_ISL_17992458 | The South African Red Cross Society - Western Cape Provincial Office | National Institute for Communicable Diseases of the National Health Laboratory Service | Everatt J, Kekana D, Mahlangu B, Stock N, Ntozini B, Ntuli N, Mnguni A, Nzimande A, Ismail A, Bhiman JN, Wolter N |
| EPI_ISL_17992460, EPI_ISL_17992462, EPI_ISL_17992464 | Rahima Moosa Mother and Child Hospital | National Institute for Communicable Diseases of the National Health Laboratory Service | Everatt J, Kekana D, Mahlangu B, Stock N, Ntozini B, Ntuli N, Mnguni A, Nzimande A, Ismail A, Bhiman JN, Wolter N |
| EPI_ISL_17992470, EPI_ISL_17992473 | Mitchells Plain Hospital | National Institute for Communicable Diseases of the National Health Laboratory Service | Everatt J, Kekana D, Mahlangu B, Stock N, Ntozini B, Ntuli N, Mnguni A, Nzimande A, Ismail A, Bhiman JN, Wolter N |
| EPI_ISL_17992475, EPI_ISL_17992477, EPI_ISL_17992479, EPI_ISL_17992481, EPI_ISL_17992483, EPI_ISL_17992485 | The South African Red Cross Society - Western Cape Provincial Office | National Institute for Communicable Diseases of the National Health Laboratory Service | Everatt J, Kekana D, Mahlangu B, Stock N, Ntozini B, Ntuli N, Mnguni A, Nzimande A, Ismail A, Bhiman JN, Wolter N |
| EPI_ISL_17992487 | Mitchells Plain Hospital | National Institute for Communicable Diseases of the National Health Laboratory Service | Everatt J, Kekana D, Mahlangu B, Stock N, Ntozini B, Ntuli N, Mnguni A, Nzimande A, Ismail A, Bhiman JN, Wolter N |
| EPI_ISL_17992489, EPI_ISL_17992492, EPI_ISL_17992494 | Rahima Moosa Mother and Child Hospital | National Institute for Communicable Diseases of the National Health Laboratory Service | Everatt J, Kekana D, Mahlangu B, Stock N, Ntozini B, Ntuli N, Mnguni A, Nzimande A, Ismail A, Bhiman JN, Wolter N |
| EPI_ISL_17992496 | Matikwana Hospital | National Institute for Communicable Diseases of the National Health Laboratory Service | Everatt J, Kekana D, Mahlangu B, Stock N, Ntozini B, Ntuli N, Mnguni A, Nzimande A, Ismail A, Bhiman JN, Wolter N |
| EPI_ISL_17992498 | Rahima Moosa Mother and Child Hospital | National Institute for Communicable Diseases of the National Health Laboratory Service | Everatt J, Kekana D, Mahlangu B, Stock N, Ntozini B, Ntuli N, Mnguni A, Nzimande A, Ismail A, Bhiman JN, Wolter N |
| EPI_ISL_17995589, EPI_ISL_17995590, EPI_ISL_17995591, EPI_ISL_17995592, EPI_ISL_17995593, EPI_ISL_17995594, EPI_ISL_17995595, EPI_ISL_17995596, EPI_ISL_17995597, EPI_ISL_17995598, EPI_ISL_17995599, EPI_ISL_17995600, EPI_ISL_17995601, EPI_ISL_17995602, EPI_ISL_17995603, EPI_ISL_17995604, EPI_ISL_17995605, EPI_ISL_17995606, EPI_ISL_17995607, EPI_ISL_17995608, EPI_ISL_17995609, EPI_ISL_17995610, EPI_ISL_17995611, EPI_ISL_17995612, EPI_ISL_17995613, EPI_ISL_17995614, EPI_ISL_17995615, EPI_ISL_17995616, EPI_ISL_17995617, EPI_ISL_17995618, EPI_ISL_17995619, EPI_ISL_17995620, EPI_ISL_17995621, EPI_ISL_17995622, EPI_ISL_17995623, EPI_ISL_17995624, EPI_ISL_17995625, EPI_ISL_17995626, EPI_ISL_17995627, EPI_ISL_17995628, EPI_ISL_17995629, EPI_ISL_17995630, EPI_ISL_17995631, EPI_ISL_17995632, EPI_ISL_17995633, EPI_ISL_17995634, EPI_ISL_17995635, EPI_ISL_17995636, EPI_ISL_17995637, EPI_ISL_17995638, EPI_ISL_17995639, EPI_ISL_17995640, EPI_ISL_17995641, EPI_ISL_17995642, EPI_ISL_17995643, EPI_ISL_17995644, EPI_ISL_17995645, EPI_ISL_17995646, EPI_ISL_17995647, EPI_ISL_17995648, EPI_ISL_17995649, EPI_ISL_17995650, EPI_ISL_17995651, EPI_ISL_17995652, EPI_ISL_17995653, EPI_ISL_17995654, EPI_ISL_17995655, EPI_ISL_17995656, EPI_ISL_17995657, EPI_ISL_17995658, EPI_ISL_17995659, EPI_ISL_17995660, EPI_ISL_17995661, EPI_ISL_17995662, EPI_ISL_17995663, EPI_ISL_17995664, EPI_ISL_17995665, EPI_ISL_17995666, EPI_ISL_17995667, EPI_ISL_17995668, EPI_ISL_17995669, EPI_ISL_17995670, EPI_ISL_17995671, EPI_ISL_17995672, EPI_ISL_17995673, EPI_ISL_17995674, EPI_ISL_17995675, EPI_ISL_17995676, EPI_ISL_17995677, EPI_ISL_17995678, EPI_ISL_17995679, EPI_ISL_17995680, EPI_ISL_17995681, EPI_ISL_17995682, EPI_ISL_17995683, EPI_ISL_17995684, EPI_ISL_17995685, EPI_ISL_17995686, EPI_ISL_17995687, EPI_ISL_17995688, EPI_ISL_17995689, EPI_ISL_17995690, EPI_ISL_17995691, EPI_ISL_17995692, EPI_ISL_17995693, EPI_ISL_17995694, EPI_ISL_17995695, EPI_ISL_17995696, EPI_ISL_17995697, EPI_ISL_17995698, EPI_ISL_17995699, EPI_ISL_17995700, EPI_ISL_17995701, EPI_ISL_17995702, EPI_ISL_17995703, EPI_ISL_17995704, EPI_ISL_17995705, EPI_ISL_17995706, EPI_ISL_17995707, EPI_ISL_17995708, EPI_ISL_17995709, EPI_ISL_17995710, EPI_ISL_17995711, EPI_ISL_17995712, EPI_ISL_17995713, EPI_ISL_17995714, EPI_ISL_17995715, EPI_ISL_17995716, EPI_ISL_17995717, EPI_ISL_17995718, EPI_ISL_17995719, EPI_ISL_17995720, EPI_ISL_17995721, EPI_ISL_17995722, EPI_ISL_17995723, EPI_ISL_17995724, EPI_ISL_17995725, EPI_ISL_17995726, EPI_ISL_17995727, EPI_ISL_17995728, EPI_ISL_17995729, EPI_ISL_17995730, EPI_ISL_17995731, EPI_ISL_17995732, EPI_ISL_17995733, EPI_ISL_17995734, EPI_ISL_17995735, EPI_ISL_17995736, EPI_ISL_17995737, EPI_ISL_17995738, EPI_ISL_17995739, EPI_ISL_17995740, EPI_ISL_17995741, EPI_ISL_17995742, EPI_ISL_17995743, EPI_ISL_17995744, EPI_ISL_17995745, EPI_ISL_17995746, EPI_ISL_17995747, EPI_ISL_17995748, EPI_ISL_17995749, EPI_ISL_17995750, EPI_ISL_17995751, EPI_ISL_17995752, EPI_ISL_17995753, EPI_ISL_17995754, EPI_ISL_17995755, EPI_ISL_17995756, EPI_ISL_17995757, EPI_ISL_17995758, EPI_ISL_17995759, EPI_ISL_17995760, EPI_ISL_17995761, EPI_ISL_17995762, EPI_ISL_17995763, EPI_ISL_17995764, EPI_ISL_17995765, EPI_ISL_17995766, EPI_ISL_17995767, EPI_ISL_17995768, EPI_ISL_17995769, EPI_ISL_17995770, EPI_ISL_17995771, EPI_ISL_17995772, EPI_ISL_17995773, EPI_ISL_17995774, EPI_ISL_17995775, EPI_ISL_17995776, EPI_ISL_17995777, EPI_ISL_17995778 | | | |
| see above | Robert Koch-Institute Nationales Referenzzentrum für Influenza | Robert Koch-Institute Nationales Referenzzentrum für Influenza | Sophie Kondgen, Janine Reiche |
| EPI_ISL_18042517 | Research Institute for Tropical Medicine, Department of Health Compound, Virology Section | WHO Collaborating Centre for Reference and Research on Influenza | Jonjee Morin, Catherine Dacasin, Vina Lea Arguelles, Xiaomin Dong, Steven Edwards, Yi-Mo Deng, Clyde Dapat, Ian Barr |
| EPI_ISL_18054314, EPI_ISL_18054332, EPI_ISL_18054349, EPI_ISL_18054351, EPI_ISL_18054367 | Kenya Medical Research Institute - KEMRI Wellcome Trust Research Programme | Kenya Medical Research Institute - KEMRI Wellcome Trust Research Programme | Lambisia,A.W., Lewa,C., Mutunga,M., Okanda,D., Githinji,G., Nokes,J.D. and Agoti,C.N. |
| EPI_ISL_18089334 | Department of Virology, National Institute of Health, Islamabad, Pakistan | Department of Virology, National Institute of Health, Islamabad, Pakistan | Massab Umair, Syed Adnan Haider, Qasim Ali, Muhammad Ammar, Zunera Jamal, and Muhammad Salman |
| EPI_ISL_18090692 | Westmead Institute for Medical Research & Sydney Infectious Disease Institute | Westmead Institute for Medical Research & Sydney Infectious Disease Institute | Pangesti,K.N.A., Ansari,H.R., Bayoumi,A., Kesson,A.M., Hill-Cawthorne,G.A. and Abd El Ghany,M. |
| EPI_ISL_18094390, EPI_ISL_18094391, EPI_ISL_18094392 | Department of Virology, National Institute of Health, Islamabad, Pakistan | Department of Virology, National Institute of Health, Islamabad, Pakistan | Massab Umair, Syed Adnan Haider, Muhammad Ammar, Zunera Jamal, and Muhammad Salman |
| EPI_ISL_18143445, EPI_ISL_18143447, EPI_ISL_18143451, EPI_ISL_18143454 | The Brotman Baty Institute for Precision Medicine | The Brotman Baty Institute for Precision Medicine | Frazar,C.D., Lee,J., Ryke,E., Gamboa,L., McDermot,E., Stone,J., Kolar,T., Han,P.D., Sibley,T.R., Truong,M., Reinhart,D., Wolf,C.R., Boeckh,M., Englund,J.A., Lutz,B.R., Waghmare,A., Viboud,C., Starita,L.M., Shendure,J., Bedford,T. and Chu,H.Y. |
| EPI_ISL_18143456 | The Brotman Baty Institute for Precision Medicine | The Brotman Baty Institute for Precision Medicine | Frazar,C.D., Lee,J., Ryke,E., Gamboa,L., McDermot,E., Stone,J., Kolar,T., Han,P.D., Sibley,T.R., Truong,M., Reinhart,D., Wolf,C.R., Duchin,J., Boeckh,M., Englund,J.A., Lutz,B.R., Waghmare,A., Viboud,C., Starita,L.M., Chu,H.Y., Bedford,T. and Shendure,J. |
| EPI_ISL_18143460 | The Brotman Baty Institute for Precision Medicine | The Brotman Baty Institute for Precision Medicine | Frazar,C.D., Lee,J., Ryke,E., Gamboa,L., McDermot,E., Stone,J., Kolar,T., Han,P.D., Sibley,T.R., Truong,M., Reinhart,D., Wolf,C.R., Boeckh,M., Englund,J.A., Lutz,B.R., Waghmare,A., Viboud,C., Starita,L.M., Shendure,J., Bedford,T. and Chu,H.Y. |
| EPI_ISL_18143463 | The Brotman Baty Institute for Precision Medicine | The Brotman Baty Institute for Precision Medicine | Frazar,C.D., Lee,J., Ryke,E., Gamboa,L., McDermot,E., Stone,J., Kolar,T., Han,P.D., Sibley,T.R., Truong,M., Reinhart,D., Wolf,C.R., Duchin,J., Boeckh,M., Englund,J.A., Lutz,B.R., Waghmare,A., Viboud,C., Starita,L.M., Chu,H.Y., Bedford,T. and Shendure,J. |
| EPI_ISL_18143464, EPI_ISL_18143465, EPI_ISL_18143467, EPI_ISL_18143469, EPI_ISL_18143471, EPI_ISL_18143472, EPI_ISL_18143476, EPI_ISL_18143478, EPI_ISL_18143479, EPI_ISL_18143480, EPI_ISL_18143483, EPI_ISL_18143484, EPI_ISL_18143486, EPI_ISL_18143489, EPI_ISL_18143490, EPI_ISL_18143494, EPI_ISL_18143495, EPI_ISL_18143496, EPI_ISL_18143497, EPI_ISL_18143499, EPI_ISL_18143500, EPI_ISL_18143502, EPI_ISL_18143503, EPI_ISL_18143506, EPI_ISL_18143507, EPI_ISL_18143508, EPI_ISL_18143509, EPI_ISL_18143510, EPI_ISL_18143511, EPI_ISL_18143512, EPI_ISL_18143513, EPI_ISL_18143514, EPI_ISL_18143517, EPI_ISL_18143518, EPI_ISL_18143520 | | | |
| see above | The Brotman Baty Institute for Precision Medicine | The Brotman Baty Institute for Precision Medicine | Frazar,C.D., Lee,J., Ryke,E., Gamboa,L., McDermot,E., Stone,J., Kolar,T., Han,P.D., Sibley,T.R., Truong,M., Reinhart,D., Wolf,C.R., Boeckh,M., Englund,J.A., Lutz,B.R., Waghmare,A., Viboud,C., Starita,L.M., Shendure,J., Bedford,T. and Chu,H.Y. |
| EPI_ISL_18143522 | The Brotman Baty Institute for Precision Medicine | The Brotman Baty Institute for Precision Medicine | Frazar,C.D., Lee,J., Ryke,E., Gamboa,L., McDermot,E., Stone,J., Kolar,T., Han,P.D., Sibley,T.R., Truong,M., Reinhart,D., Wolf,C.R., Duchin,J., Boeckh,M., Englund,J.A., Lutz,B.R., Waghmare,A., Viboud,C., Starita,L.M., Chu,H.Y., Bedford,T. and Shendure,J. |
| EPI_ISL_18143527, EPI_ISL_18143528, EPI_ISL_18143529, EPI_ISL_18143530, EPI_ISL_18143531, EPI_ISL_18143533, EPI_ISL_18143536, EPI_ISL_18143537, EPI_ISL_18143539, EPI_ISL_18143540, EPI_ISL_18143542, EPI_ISL_18143544, EPI_ISL_18143547, EPI_ISL_18143548, EPI_ISL_18143553, EPI_ISL_18143554, EPI_ISL_18143555, EPI_ISL_18143556, EPI_ISL_18143557, EPI_ISL_18143560, EPI_ISL_18143562, EPI_ISL_18143564, EPI_ISL_18143566, EPI_ISL_18143571, EPI_ISL_18143572, EPI_ISL_18143575, EPI_ISL_18143576, EPI_ISL_18143578, EPI_ISL_18143583, EPI_ISL_18143585, EPI_ISL_18143589, EPI_ISL_18143591, EPI_ISL_18143592, EPI_ISL_18143599, EPI_ISL_18143600, EPI_ISL_18143601, EPI_ISL_18143604, EPI_ISL_18143605, EPI_ISL_18143606, EPI_ISL_18143607, EPI_ISL_18143608, EPI_ISL_18143609, EPI_ISL_18143610, EPI_ISL_18143611, EPI_ISL_18143613, EPI_ISL_18143614, EPI_ISL_18143615, EPI_ISL_18143618, EPI_ISL_18143620, EPI_ISL_18143621, EPI_ISL_18143622, EPI_ISL_18143623, EPI_ISL_18143624, EPI_ISL_18143626, EPI_ISL_18143629, EPI_ISL_18143636, EPI_ISL_18143638, EPI_ISL_18143639, EPI_ISL_18143640, EPI_ISL_18143641, EPI_ISL_18143643, EPI_ISL_18143644, EPI_ISL_18143645, EPI_ISL_18143647, EPI_ISL_18143648, EPI_ISL_18143649, EPI_ISL_18143651, EPI_ISL_18143652, EPI_ISL_18143654, EPI_ISL_18143655, EPI_ISL_18143656, EPI_ISL_18143657, EPI_ISL_18143658, EPI_ISL_18143660, EPI_ISL_18143662, EPI_ISL_18143663, EPI_ISL_18143664, EPI_ISL_18143668, EPI_ISL_18143672, EPI_ISL_18143675, EPI_ISL_18143677, EPI_ISL_18143679, EPI_ISL_18143680, EPI_ISL_18143685, EPI_ISL_18143686, EPI_ISL_18143687, EPI_ISL_18143688, EPI_ISL_18143692, EPI_ISL_18143694, EPI_ISL_18143695, EPI_ISL_18143699, EPI_ISL_18143700, EPI_ISL_18143701, EPI_ISL_18143703, EPI_ISL_18143704, EPI_ISL_18143707, EPI_ISL_18143709, EPI_ISL_18143711, EPI_ISL_18143714, EPI_ISL_18143715 | | | |

| | | | |
|---|---|---|---|
| see above | The Brotman Baty Institute for Precision Medicine | The Brotman Baty Institute for Precision Medicine | Frazar,C.D., Lee,J., Ryke,E., Gamboa,L., McDermot,E., Stone,J., Kolar,T., Han,P.D., Sibley,T.R., Truong,M., Reinhart,D., Wolf,C.R., Boeckh,M., Englund,J.A., Lutz,B.R., Waghmare,A., Viboud,C., Starita,L.M., Shendure,J., Bedford,T. and Chu,H.Y. |
| EPI_ISL_18204747, EPI_ISL_18204754, EPI_ISL_18204756, EPI_ISL_18204758, EPI_ISL_18204760, EPI_ISL_18204771, EPI_ISL_18204773, EPI_ISL_18204775, EPI_ISL_18204777, EPI_ISL_18204780, EPI_ISL_18204782, EPI_ISL_18204784, EPI_ISL_18204786, EPI_ISL_18204838, EPI_ISL_18204856, EPI_ISL_18204861, EPI_ISL_18204867 | | | |
| see above | Microbiology Department. Complexo Hospitalario Universitario de Vigo | Microbiology Department. Complexo Hospitalario Universitario de Vigo | Daviña C, Martinez L, Perez-Castro S |
| EPI_ISL_18228265, EPI_ISL_18228290, EPI_ISL_18228291, EPI_ISL_18228292, EPI_ISL_18228293, EPI_ISL_18228294, EPI_ISL_18228295, EPI_ISL_18240645, EPI_ISL_18240646, EPI_ISL_18240647, EPI_ISL_18240648, EPI_ISL_18240649, EPI_ISL_18240650, EPI_ISL_18240651 | | | |
| see above | Institut Pasteur de Dakar | Institut Pasteur de Dakar | Jallow, Mamadou Malado; Diagne, Moussa Moise; Sankhe, Safietou; Mendy, Marie Pedapa; Ndiaye, Ndiende Koba; Sy, Sara; Kiori, Davy; Goudiaby, Deborah; Dia, Ndongo |
| EPI_ISL_18262238, EPI_ISL_18262240 | Instituto de Medicina Tropical Alexander von Humboldt, Universidad Peruana Cayetano Heredia | Laboratorio de Genómica Microbiana, Universidad Peruana Cayetano Heredia | Ericka Meza, Diego Cuicapuza, Anne Martínez-Ventura, Brenda Ayzanoa, Janet Huancachoque, César Ugarte, Carlos Zamudio, Pablo Tsukayama |
| EPI_ISL_18265062, EPI_ISL_18265063, EPI_ISL_18265064, EPI_ISL_18265065, EPI_ISL_18265066, EPI_ISL_18265067, EPI_ISL_18272375, EPI_ISL_18272376, EPI_ISL_18272377, EPI_ISL_18272378, EPI_ISL_18272379, EPI_ISL_18272380, EPI_ISL_18272381, EPI_ISL_18272382, EPI_ISL_18272386, EPI_ISL_18272387, EPI_ISL_18272388, EPI_ISL_18272389, EPI_ISL_18272390, EPI_ISL_18272391, EPI_ISL_18272392, EPI_ISL_18272393, EPI_ISL_18272394, EPI_ISL_18272398, EPI_ISL_18272399, EPI_ISL_18272400, EPI_ISL_18272401, EPI_ISL_18272402, EPI_ISL_18272403, EPI_ISL_18272404, EPI_ISL_18272405, EPI_ISL_18277074, EPI_ISL_18277075, EPI_ISL_18277076, EPI_ISL_18277077, EPI_ISL_18277078, EPI_ISL_18277079, EPI_ISL_18277080, EPI_ISL_18277081, EPI_ISL_18277082, EPI_ISL_18277083, EPI_ISL_18277084, EPI_ISL_18277085, EPI_ISL_18277086, EPI_ISL_18277087, EPI_ISL_18277132, EPI_ISL_18277133, EPI_ISL_18277134, EPI_ISL_18277135 | | | |
| see above | West of Scotland Specialist Virology Centre | West of Scotland Specialist Virology Centre | Lynne Ferguson, Imogen Johnston-Menzies, Rory Gunson |
| EPI_ISL_18279085 | National Institute of Public Health | National Institute of Public Health | Chhorvann CHHEA, Darapheak CHAU, Sitha PRUM, Visal CHHE, Sokha DUL, Sengly MANH, Phally VY, Borann SAR, Chanthap LON, Jessica E. Manning, Sophana Chea, Savuth CHIN |
| EPI_ISL_18321008, EPI_ISL_18321009, EPI_ISL_18321010, EPI_ISL_18321011, EPI_ISL_18321012, EPI_ISL_18321013, EPI_ISL_18321014, EPI_ISL_18321015, EPI_ISL_18321016, EPI_ISL_18321017, EPI_ISL_18321018, EPI_ISL_18321019, EPI_ISL_18321020, EPI_ISL_18321034, EPI_ISL_18321035, EPI_ISL_18321037, EPI_ISL_18321039, EPI_ISL_18321040, EPI_ISL_18321042, EPI_ISL_18321044, EPI_ISL_18321045, EPI_ISL_18321046, EPI_ISL_18321048, EPI_ISL_18321049, EPI_ISL_18321051, EPI_ISL_18321054, EPI_ISL_18321055, EPI_ISL_18321078, EPI_ISL_18321080, EPI_ISL_18321081, EPI_ISL_18321082, EPI_ISL_18321083, EPI_ISL_18321084, EPI_ISL_18321085, EPI_ISL_18321088, EPI_ISL_18321089, EPI_ISL_18321091, EPI_ISL_18321092, EPI_ISL_18321094, EPI_ISL_18321095, EPI_ISL_18321096, EPI_ISL_18321097, EPI_ISL_18321098, EPI_ISL_18321099, EPI_ISL_18321100, EPI_ISL_18321101 | | | |
| see above | National Virus Reference Laboratory | National Virus Reference Laboratory | Michael Carr, Charlene Bennett, Jonathan Dean, Daniel Hare, Cillian F De Gascun |
| EPI_ISL_18321558 | Servicio de Microbiología Complejo Hospitalario Universitario Nuestra Señora de Candelaria | Institute of Technology and Renewable Energy (ITER) Genomics Division | Julia,Alcoba-Florez;Rafaela,González-Montelongo;Diego,García-Martínez de Artola;Adrián,Muñoz-Barrera;Helena,Gil-Campesino;Oscar,Diez-Gil;Jose Miguel,Lorenzo-Salazar;Carlos,Flores |
| EPI_ISL_18329452, EPI_ISL_18329453, EPI_ISL_18329454, EPI_ISL_18329455, EPI_ISL_18329456, EPI_ISL_18329457, EPI_ISL_18329458, EPI_ISL_18329459, EPI_ISL_18329460, EPI_ISL_18329462 | Microbiology and Virology Depertment, Fondazione IRCCS Policlinico San Matteo, Pavia | Microbiology and Virology Depertment, Fondazione IRCCS Policlinico San Matteo, Pavia | Gugliemo Ferrari, Federica Giardina, Antonino Pitrolo, Fausto Baldanti |
| EPI_ISL_18334078, EPI_ISL_18334079 | Kitasato University, Infection Control and Immunology, Yukie Katayama Omura Satoshi Memorial Institute | Kitasato University, Infection Control and Immunology, Yukie Katayama Omura Satoshi Memorial Institute | Takai-Todaka,R., Ito,T., Haga,K., Ishiyama,R., Igarashi,T., Katayama,Y., Nakayama,T. and Katayama,K. |
| EPI_ISL_18334163, EPI_ISL_18334166, EPI_ISL_18334167, EPI_ISL_18334169, EPI_ISL_18334171, EPI_ISL_18334173, EPI_ISL_18334175, EPI_ISL_18334176, EPI_ISL_18334177, EPI_ISL_18334178, EPI_ISL_18334179, EPI_ISL_18334180, EPI_ISL_18334185, EPI_ISL_18334187, EPI_ISL_18334190, EPI_ISL_18334191, EPI_ISL_18334194, EPI_ISL_18334195, EPI_ISL_18334196, EPI_ISL_18334199, EPI_ISL_18334201, EPI_ISL_18334204, EPI_ISL_18334205, EPI_ISL_18334206, EPI_ISL_18334214, EPI_ISL_18334215, EPI_ISL_18334216, EPI_ISL_18334220, EPI_ISL_18334221, EPI_ISL_18334222, EPI_ISL_18334225, EPI_ISL_18334230, EPI_ISL_18334231, EPI_ISL_18334232, EPI_ISL_18334236 | | | |
| see above | Baylor College of Medicine, Division of Pediatric Tropical Medicine | Baylor College of Medicine, Division of Pediatric Tropical Medicine | Avadhanula,V., Agustinho,D.P., Menon,V.K., Chemaly,R.F., Shah,D.P., Qin,X., Surathu,A., Doddapaneni,H., Muzny,D.M., Metcalf,G.A., Gregeen,S.J., Gibbs,R.A., Petrosino,J.J., Sedlazeck,F.J. and Piedra,P.A. |
| EPI_ISL_1834083, EPI_ISL_1834084, EPI_ISL_1834085, EPI_ISL_1834086, EPI_ISL_1834087, EPI_ISL_1834088, EPI_ISL_1834089, EPI_ISL_1834090, EPI_ISL_1834091, EPI_ISL_1834092, EPI_ISL_1834093, EPI_ISL_1834094, EPI_ISL_1834095, EPI_ISL_1834096, EPI_ISL_1834097, EPI_ISL_1834098, EPI_ISL_1834099, EPI_ISL_1834100, EPI_ISL_1834101, EPI_ISL_1834102, EPI_ISL_1834103, EPI_ISL_1834104, EPI_ISL_1834105, EPI_ISL_1834106, EPI_ISL_1834107, EPI_ISL_1834108, EPI_ISL_1834109, EPI_ISL_1834110, EPI_ISL_1834112, EPI_ISL_1834113, EPI_ISL_1834114, EPI_ISL_1834115, EPI_ISL_1834116, EPI_ISL_1834117, EPI_ISL_1834118, EPI_ISL_1834119, EPI_ISL_1834120, EPI_ISL_1834121, EPI_ISL_1834123, EPI_ISL_1834124, EPI_ISL_1834125, EPI_ISL_1834126, EPI_ISL_1834127, EPI_ISL_1834128, EPI_ISL_1834129, EPI_ISL_1834130, EPI_ISL_1834131, EPI_ISL_1834132, EPI_ISL_1834133, EPI_ISL_1834134, EPI_ISL_1834135, EPI_ISL_1834136, EPI_ISL_1834137, EPI_ISL_1834138, EPI_ISL_1834139, EPI_ISL_1834140, EPI_ISL_1834141, EPI_ISL_1834142, EPI_ISL_1834143, EPI_ISL_1834144, EPI_ISL_1834145, EPI_ISL_1834146, EPI_ISL_1834147, EPI_ISL_1834148, EPI_ISL_1834149 | | | |
| see above | Royal Children's Hospital | WHO Influenza Centre for Reference and Research on Influenza | Angela Todd, Yi-Mo Deng, Annette Alafaci, Naomi Komadina |
| EPI_ISL_1834150, EPI_ISL_1834151, EPI_ISL_1834152, EPI_ISL_1834153, EPI_ISL_1834154, EPI_ISL_1834155, EPI_ISL_1834158, EPI_ISL_1834159, EPI_ISL_1834160, EPI_ISL_1834161, EPI_ISL_1834162, EPI_ISL_1834163, EPI_ISL_1834164, EPI_ISL_1834165, EPI_ISL_1834166, EPI_ISL_1834167, EPI_ISL_1834168, EPI_ISL_1834169, EPI_ISL_1834170, EPI_ISL_1834171, EPI_ISL_1834172, EPI_ISL_1834173 | | | |
| see above | Respiratory Virus Unit, National Infection Service, Public Health England | National Infection Service, Public Health England | Zambon M, Talts T, Ellis J, Miah S, Platt S |
| EPI_ISL_18374665 | Mitchells Plain Hospital | National Institute for Communicable Diseases of the National Health Laboratory Service | Everatt J, Kekana D, Mahlangu B, Stock N, Ntozini B, Ntuli N, Mnguni A, Nzimande A, Ismail A, Bhiman JN, Wolter N |
| EPI_ISL_18374668 | Rahima Moosa Mother and Child Hospital | National Institute for Communicable Diseases of the National Health Laboratory Service | Everatt J, Kekana D, Mahlangu B, Stock N, Ntozini B, Ntuli N, Mnguni A, Nzimande A, Ismail A, Bhiman JN, Wolter N |
| EPI_ISL_18387876, EPI_ISL_18387878, EPI_ISL_18387879, EPI_ISL_18387880, EPI_ISL_18387881, EPI_ISL_18387883, EPI_ISL_18387885, EPI_ISL_18387886, EPI_ISL_18387888, EPI_ISL_18387890, EPI_ISL_18387891, EPI_ISL_18387892, EPI_ISL_18387893, EPI_ISL_18387894, EPI_ISL_18387895, EPI_ISL_18387896, EPI_ISL_18387897, EPI_ISL_18387899, EPI_ISL_18387901, EPI_ISL_18387903, EPI_ISL_18387904, EPI_ISL_18387906, EPI_ISL_18387908, EPI_ISL_18387909, EPI_ISL_18387911, EPI_ISL_18387912, EPI_ISL_18387913, EPI_ISL_18387914, EPI_ISL_18387915, EPI_ISL_18387916, EPI_ISL_18387919, EPI_ISL_18387921, EPI_ISL_18387922, EPI_ISL_18387923, EPI_ISL_18387925, EPI_ISL_18387926, EPI_ISL_18387927, EPI_ISL_18387928, EPI_ISL_18387929, EPI_ISL_18387930, EPI_ISL_18387931, EPI_ISL_18387933, EPI_ISL_18387934, EPI_ISL_18387937, EPI_ISL_18387938, EPI_ISL_18387939, EPI_ISL_18387941, EPI_ISL_18387943, EPI_ISL_18387944, EPI_ISL_18387946, EPI_ISL_18387948, EPI_ISL_18387949, EPI_ISL_18387950, EPI_ISL_18387953, EPI_ISL_18387955, EPI_ISL_18387956, EPI_ISL_18387958, EPI_ISL_18387959, EPI_ISL_18387960, EPI_ISL_18387962, EPI_ISL_18387963 | | | |
| see above | The Brotman Baty Institute for Precision Medicine | The Brotman Baty Institute for Precision Medicine | Frazar,C.D., Lee,J., Ryke,E., Gamboa,L., McDermot,E., Kolar,T., Han,P.D., Sibley,T.R., Reinhart,D., Grindstaff,S., Truong,M., Babu,T.M., Feldstein,L.R., Saydah,S., Briggs-Hagen,M., Casto,A., Ehmen,B., Englund,J.A., Fortmann,S.P., Kuntz,J.L., Lockwood,T., Midgley,C.M., Mularski,R.A., Ogilvie,T., Reich,S., Schmidt,M.A., Smith,N., Starita,L., Stone,J., Vandermeer,M., Weil,A.A., Wolf,C.R., Chu,H.Y. and Naleway,A.L. |
| EPI_ISL_18446761, EPI_ISL_18446762, EPI_ISL_18446765, EPI_ISL_18446766, EPI_ISL_18446767, EPI_ISL_18446768, EPI_ISL_18446769, EPI_ISL_18446771, EPI_ISL_18446772, EPI_ISL_18446773, EPI_ISL_18446774, EPI_ISL_18446775, EPI_ISL_18446776, EPI_ISL_18446778, EPI_ISL_18446779, EPI_ISL_18446780, EPI_ISL_18446781, EPI_ISL_18446782, EPI_ISL_18446783, EPI_ISL_18446784, EPI_ISL_18446786, EPI_ISL_18446787, EPI_ISL_18446788, EPI_ISL_18446790, EPI_ISL_18446791, EPI_ISL_18446793, EPI_ISL_18446794, EPI_ISL_18446795, EPI_ISL_18446796, EPI_ISL_18446797, EPI_ISL_18446798, EPI_ISL_18446799, EPI_ISL_18446800, EPI_ISL_18446802, EPI_ISL_18446803, EPI_ISL_18446805, EPI_ISL_18446806, EPI_ISL_18446807, EPI_ISL_18446808, EPI_ISL_18446809 | | | |
| see above | Vittore Buzzi Children Hospital, Pediatric Department | Laboratory of Infectious Diseases, Department of Biomedical and Clinical Sciences, University of Milan | Alessia Lai, Annalisa Bergna, Valentina Fabiano, Carla della Ventura, Giulia Fumagalli, Alessandra Mari, Martina Loiodice, Gian Vincenzo Zuccotti, Gianguglielmo Zehender |
| EPI_ISL_18452361, EPI_ISL_18452363, EPI_ISL_18452364, EPI_ISL_18452365, EPI_ISL_18452366, EPI_ISL_18452367, EPI_ISL_18452368, EPI_ISL_18452369, EPI_ISL_18452370, EPI_ISL_18452371 | The Brotman Baty Institute for Precision Medicine | The Brotman Baty Institute for Precision Medicine | Frazar,C.D., Lee,J., Ryke,E., Gamboa,L., McDermot,E., Kolar,T., Han,P.D., Sibley,T.R., Reinhart,D., Grindstaff,S., Truong,M., Babu,T.M., Feldstein,L.R., Saydah,S., Briggs-Hagen,M., Casto,A., Ehmen,B., Englund,J.A., Fortmann,S.P., Kuntz,J.L., Lockwood,T., Midgley,C.M., Mularski,R.A., Ogilvie,T., Reich,S., Schmidt,M.A., Smith,N., Starita,L., Stone,J., Vandermeer,M., Weil,A.A., Wolf,C.R., Chu,H.Y. and Naleway,A.L. |
| EPI_ISL_18482717, EPI_ISL_18482718, EPI_ISL_18482719, EPI_ISL_18482720, EPI_ISL_18482723, EPI_ISL_18482724, EPI_ISL_18482728, EPI_ISL_18482729, EPI_ISL_18482730, EPI_ISL_18482731, EPI_ISL_18482734, EPI_ISL_18482735, EPI_ISL_18482736, EPI_ISL_18482740, EPI_ISL_18482741, EPI_ISL_18482744, EPI_ISL_18482746, EPI_ISL_18482748, EPI_ISL_18482749, EPI_ISL_18482751, EPI_ISL_18482752, EPI_ISL_18482753, EPI_ISL_18482756, EPI_ISL_18482757, EPI_ISL_18482759, EPI_ISL_18482760, EPI_ISL_18482761, EPI_ISL_18482762, EPI_ISL_18482765, EPI_ISL_18482766, EPI_ISL_18482767, EPI_ISL_18482769, EPI_ISL_18482771, EPI_ISL_18482772, EPI_ISL_18482773, EPI_ISL_18482775, EPI_ISL_18482776, EPI_ISL_18482777, EPI_ISL_18482778, EPI_ISL_18482779, EPI_ISL_18482780, EPI_ISL_18482781, EPI_ISL_18482782, EPI_ISL_18482784, EPI_ISL_18482785, EPI_ISL_18482786, EPI_ISL_18482787, EPI_ISL_18482788, EPI_ISL_18482791, EPI_ISL_18482792, EPI_ISL_18482793, EPI_ISL_18482795, EPI_ISL_18482797, EPI_ISL_18482799, EPI_ISL_18482800, EPI_ISL_18482801, EPI_ISL_18482802, EPI_ISL_18482804, EPI_ISL_18482805, EPI_ISL_18482807, EPI_ISL_18482808, EPI_ISL_18482809, EPI_ISL_18482811, EPI_ISL_18482813, EPI_ISL_18482814, EPI_ISL_18482815, EPI_ISL_18482816, EPI_ISL_18482817, EPI_ISL_18482819, EPI_ISL_18482820 | | | |
| see above | Beijing Children's Hospital, Capital Medical University, Laboratory of Infection and Virology | Beijing Children's Hospital, Capital Medical University, Laboratory of Infection and Virology | Li,F., Zhu,Y. and Chen,X. |
| EPI_ISL_18507281, EPI_ISL_18507283, EPI_ISL_18507286, EPI_ISL_18507287, EPI_ISL_18507288, EPI_ISL_18507291, EPI_ISL_18507292, EPI_ISL_18507293, EPI_ISL_18507294, EPI_ISL_18507295, EPI_ISL_18507296, EPI_ISL_18507297 | | | |
| see above | Victorian Infectious Diseases Reference Laboratory, Translational Diagnostics | Victorian Infectious Diseases Reference Laboratory, Translational Diagnostics | Steinig JOACHIM,E. |
| EPI_ISL_18509832, EPI_ISL_18509835, EPI_ISL_18509838, EPI_ISL_18509840, EPI_ISL_18509842, EPI_ISL_18509844, EPI_ISL_18509846, EPI_ISL_18509849, EPI_ISL_18509882, EPI_ISL_18509884, EPI_ISL_18509895, EPI_ISL_18509897, EPI_ISL_18509899, EPI_ISL_18509901, EPI_ISL_18509903, EPI_ISL_18510219 | | | |
| see above | Institut Pasteur de Dakar | Institut Pasteur de Dakar | Jallow,Mamadou Malado; Diagne,Moussa Moise; Sankhe,Safietou; Diop,Seynabou Mbaye Ba Souna; Mendy,Marie Pedapa; Sy,Sara; Ndiaye,Ndiende Koba; Kiory, Davy Evrard; Goudiaby,Deborah; Dia,Ndongo |
| EPI_ISL_18537458 | RELAB - BIOGROUP - Plateau technique St Denis | National Reference Center for Viruses of Respiratory Infections, Institut Pasteur, Paris | Léa Avon, Marion Barbet, Marine Bernard, Emma Bezot, Angela Brisebarre, Océane Dehan, Flora Donati, Vanessa Guimaraes, Banujaa Jeyarajah, Florian Préjean, Yannis Rahou, Sylvie van der Werf, Frédéric Lemoine, Samar Berreira Ibraim, Jérôme Bourret, Kévin Da Silva, Maud Vanpeene, Vincent Enouf, Marie-Anne Rameix-Welti, Yanis CHAIB |
| EPI_ISL_18537459 | Sentinelles Ile-de-France | National Reference Center for Viruses of Respiratory Infections, Institut Pasteur, Paris | Léa Avon, Marion Barbet, Marine Bernard, Emma Bezot, Angela Brisebarre, Océane Dehan, Flora Donati, Vanessa Guimaraes, Banujaa Jeyarajah, Florian Préjean, Yannis Rahou, Sylvie van der Werf, Frédéric Lemoine, Samar Berreira Ibraim, Jérôme Bourret, Kévin Da Silva, Maud Vanpeene, Vincent Enouf, Marie-Anne Rameix-Welti, Patricia LUBELSKI |
| EPI_ISL_18537460 | Sentinelles Province | National Reference Center for Viruses of Respiratory Infections, Institut Pasteur, Paris | Léa Avon, Marion Barbet, Marine Bernard, Emma Bezot, Angela Brisebarre, Océane Dehan, Flora Donati, Vanessa Guimaraes, Banujaa Jeyarajah, Florian Préjean, Yannis Rahou, Sylvie van der Werf, Frédéric Lemoine, Samar Berreira Ibraim, Jérôme Bourret, Kévin Da Silva, Maud Vanpeene, Vincent Enouf, Marie-Anne Rameix-Welti, Eric VAN MELKEBEKE |
| EPI_ISL_18537461 | Sentinelles Province | National Reference Center for Viruses of Respiratory Infections, Institut Pasteur, Paris | Léa Avon, Marion Barbet, Marine Bernard, Emma Bezot, Angela Brisebarre, Océane Dehan, Flora Donati, Vanessa Guimaraes, Banujaa Jeyarajah, Florian Préjean, Yannis Rahou, Sylvie van der Werf, Frédéric Lemoine, Samar Berreira Ibraim, Jérôme Bourret, Kévin Da Silva, Maud Vanpeene, Vincent Enouf, Marie-Anne Rameix-Welti, Gwenaelle MAHE |
| EPI_ISL_18537462 | Sentinelles Ile-de-France | National Reference Center for Viruses of Respiratory Infections, Institut Pasteur, Paris | Léa Avon, Marion Barbet, Marine Bernard, Emma Bezot, Angela Brisebarre, Océane Dehan, Flora Donati, Vanessa Guimaraes, Banujaa Jeyarajah, Florian Préjean, Yannis Rahou, Sylvie van der Werf, Frédéric Lemoine, Samar Berreira Ibraim, Jérôme Bourret, Kévin Da Silva, Maud Vanpeene, Vincent Enouf, Marie-Anne Rameix-Welti, Frédéric URBAIN |
| EPI_ISL_18537463, EPI_ISL_18537464 | RELAB - BIOGROUP - Plateau technique St Denis | National Reference Center for Viruses of Respiratory Infections, Institut Pasteur, Paris | Léa Avon, Marion Barbet, Marine Bernard, Emma Bezot, Angela Brisebarre, Océane Dehan, Flora Donati, Vanessa Guimaraes, Banujaa Jeyarajah, Florian Préjean, Yannis Rahou, Sylvie van der Werf, Frédéric Lemoine, Samar Berreira Ibraim, Jérôme Bourret, Kévin Da Silva, Maud Vanpeene, Vincent Enouf, Marie-Anne Rameix-Welti, Yanis CHAIB |
| EPI_ISL_18537465 | Sentinelles Ile-de-France | National Reference Center for Viruses of Respiratory Infections, Institut Pasteur, Paris | Léa Avon, Marion Barbet, Marine Bernard, Emma Bezot, Angela Brisebarre, Océane Dehan, Flora Donati, Vanessa Guimaraes, Banujaa Jeyarajah, Florian Préjean, Yannis Rahou, Sylvie van der Werf, Frédéric Lemoine, Samar Berreira Ibraim, Jérôme Bourret, Kévin Da Silva, Maud Vanpeene, Vincent Enouf, Marie-Anne Rameix-Welti, Marie-Anne COSSON |
| EPI_ISL_18537468 | RELAB - BIOGROUP - Plateau technique St Denis | National Reference Center for Viruses of Respiratory Infections, Institut Pasteur, Paris | Léa Avon, Marion Barbet, Marine Bernard, Emma Bezot, Angela Brisebarre, Océane Dehan, Flora Donati, Vanessa Guimaraes, Banujaa Jeyarajah, Florian Préjean, Yannis Rahou, Sylvie van der Werf, Frédéric Lemoine, Samar Berreira Ibraim, Jérôme Bourret, Kévin Da Silva, Maud Vanpeene, Vincent Enouf, Marie-Anne Rameix-Welti, Yanis CHAIB |
| EPI_ISL_18537469 | Hôpital Ambroise Paré - Service de Microbiologie et hygiène | National Reference Center for Viruses of Respiratory Infections, Institut Pasteur, Paris | Léa Avon, Marion Barbet, Marine Bernard, Emma Bezot, Angela Brisebarre, Océane Dehan, Flora Donati, Vanessa Guimaraes, Banujaa Jeyarajah, Florian Préjean, Yannis Rahou, Sylvie van der Werf, Frédéric Lemoine, Samar Berreira Ibraim, Jérôme Bourret, Kévin Da Silva, Maud Vanpeene, Vincent Enouf, Marie-Anne Rameix-Welti, Elyanne GAULT |
| EPI_ISL_18537472, EPI_ISL_18537474, EPI_ISL_18537479, EPI_ISL_18537480 | RELAB - BIOGROUP - Plateau technique St Denis | National Reference Center for Viruses of Respiratory Infections, Institut Pasteur, Paris | Léa Avon, Marion Barbet, Marine Bernard, Emma Bezot, Angela Brisebarre, Océane Dehan, Flora Donati, Vanessa Guimaraes, Banujaa Jeyarajah, Florian Préjean, Yannis Rahou, Sylvie van der Werf, Frédéric Lemoine, Samar Berreira Ibraim, Jérôme Bourret, Kévin Da Silva, Maud Vanpeene, Vincent Enouf, Marie-Anne Rameix-Welti, Yanis CHAIB |
| EPI_ISL_18537482 | Sentinelles Ile-de-France | National Reference Center for Viruses of | Léa Avon, Marion Barbet, Marine Bernard, Emma Bezot, Angela Brisebarre, Océane Dehan, Flora Donati, Vanessa Guimaraes, Banujaa Jeyarajah, Florian Préjean, Yannis Rahou, Sylvie van der Werf, Frédéric Lemoine, Samar Berreira |

| | | | |
|---|---|---|---|
| | | Respiratory Infections, Institut Pasteur, Paris | Ibraim, Jérôme Bourret, Kévin Da Silva, Maud Vanpeene, Vincent Enouf, Marie-Anne Rameix-Welti, MARIE ANNE DAUMONT |
| EPI_ISL_18537483, EPI_ISL_18537484, EPI_ISL_18537487, EPI_ISL_18537489 | RELAB - BIOGROUP - Plateau technique St Denis | National Reference Center for Viruses of Respiratory Infections, Institut Pasteur, Paris | Léa Avon, Marion Barbet, Marine Bernard, Emma Bezot, Angela Brisebarre, Océane Dehan, Flora Donati, Vanessa Guimaraes, Banujaa Jeyarajah, Florian Préjean, Yannis Rahou, Sylvie van der Werf, Frédéric Lemoine, Samar Berreira Ibraim, Jérôme Bourret, Kévin Da Silva, Maud Vanpeene, Vincent Enouf, Marie-Anne Rameix-Welti, Yanis CHAIB |
| EPI_ISL_18537494 | Hôpital Robert Debre, Service Microbiologie | National Reference Center for Viruses of Respiratory Infections, Institut Pasteur, Paris | Léa Avon, Marion Barbet, Marine Bernard, Emma Bezot, Angela Brisebarre, Océane Dehan, Flora Donati, Vanessa Guimaraes, Banujaa Jeyarajah, Florian Préjean, Yannis Rahou, Sylvie van der Werf, Frédéric Lemoine, Samar Berreira Ibraim, Jérôme Bourret, Kévin Da Silva, Maud Vanpeene, Vincent Enouf, Marie-Anne Rameix-Welti, Stéphane BONACORSI |
| EPI_ISL_18537495, EPI_ISL_18537496, EPI_ISL_18537497 | RELAB - BIOGROUP - Plateau technique St Denis | National Reference Center for Viruses of Respiratory Infections, Institut Pasteur, Paris | Léa Avon, Marion Barbet, Marine Bernard, Emma Bezot, Angela Brisebarre, Océane Dehan, Flora Donati, Vanessa Guimaraes, Banujaa Jeyarajah, Florian Préjean, Yannis Rahou, Sylvie van der Werf, Frédéric Lemoine, Samar Berreira Ibraim, Jérôme Bourret, Kévin Da Silva, Maud Vanpeene, Vincent Enouf, Marie-Anne Rameix-Welti, Yanis CHAIB |
| EPI_ISL_18537503 | RELAB - Laboratoire Biomer | National Reference Center for Viruses of Respiratory Infections, Institut Pasteur, Paris | Léa Avon, Marion Barbet, Marine Bernard, Emma Bezot, Angela Brisebarre, Océane Dehan, Flora Donati, Vanessa Guimaraes, Banujaa Jeyarajah, Florian Préjean, Yannis Rahou, Sylvie van der Werf, Frédéric Lemoine, Samar Berreira Ibraim, Jérôme Bourret, Kévin Da Silva, Maud Vanpeene, Vincent Enouf, Marie-Anne Rameix-Welti, Alexandra JACQUES |
| EPI_ISL_18537504 | RELAB - BIOGROUP - Plateau technique St Denis | National Reference Center for Viruses of Respiratory Infections, Institut Pasteur, Paris | Léa Avon, Marion Barbet, Marine Bernard, Emma Bezot, Angela Brisebarre, Océane Dehan, Flora Donati, Vanessa Guimaraes, Banujaa Jeyarajah, Florian Préjean, Yannis Rahou, Sylvie van der Werf, Frédéric Lemoine, Samar Berreira Ibraim, Jérôme Bourret, Kévin Da Silva, Maud Vanpeene, Vincent Enouf, Marie-Anne Rameix-Welti, Yanis CHAIB |
| EPI_ISL_18537505 | RELAB - Laboratoire Biomer | National Reference Center for Viruses of Respiratory Infections, Institut Pasteur, Paris | Léa Avon, Marion Barbet, Marine Bernard, Emma Bezot, Angela Brisebarre, Océane Dehan, Flora Donati, Vanessa Guimaraes, Banujaa Jeyarajah, Florian Préjean, Yannis Rahou, Sylvie van der Werf, Frédéric Lemoine, Samar Berreira Ibraim, Jérôme Bourret, Kévin Da Silva, Maud Vanpeene, Vincent Enouf, Marie-Anne Rameix-Welti, Alexandra JACQUES |
| EPI_ISL_18537506, EPI_ISL_18537507, EPI_ISL_18537508, EPI_ISL_18537510, EPI_ISL_18537511, EPI_ISL_18537512 | RELAB - BIOGROUP - Plateau technique St Denis | National Reference Center for Viruses of Respiratory Infections, Institut Pasteur, Paris | Léa Avon, Marion Barbet, Marine Bernard, Emma Bezot, Angela Brisebarre, Océane Dehan, Flora Donati, Vanessa Guimaraes, Banujaa Jeyarajah, Florian Préjean, Yannis Rahou, Sylvie van der Werf, Frédéric Lemoine, Samar Berreira Ibraim, Jérôme Bourret, Kévin Da Silva, Maud Vanpeene, Vincent Enouf, Marie-Anne Rameix-Welti, Yanis CHAIB |
| EPI_ISL_18568000, EPI_ISL_18568001 | RELAB - Laboratoire CBM 25 | National Reference Center for Viruses of Respiratory Infections, Institut Pasteur, Paris | Léa Avon, Marion Barbet, Marine Bernard, Emma Bezot, Angela Brisebarre, Océane Dehan, Flora Donati, Vanessa Guimaraes, Banujaa Jeyarajah, Florian Préjean, Yannis Rahou, Sylvie van der Werf, Frédéric Lemoine, Samar Berreira Ibraim, Jérôme Bourret, Kévin Da Silva, Maud Vanpeene, Vincent Enouf, Marie-Anne Rameix-Welti, Sophie GIRARD |
| EPI_ISL_18568002 | Sentinelles Province | National Reference Center for Viruses of Respiratory Infections, Institut Pasteur, Paris | Léa Avon, Marion Barbet, Marine Bernard, Emma Bezot, Angela Brisebarre, Océane Dehan, Flora Donati, Vanessa Guimaraes, Banujaa Jeyarajah, Florian Préjean, Yannis Rahou, Sylvie van der Werf, Frédéric Lemoine, Samar Berreira Ibraim, Jérôme Bourret, Kévin Da Silva, Maud Vanpeene, Vincent Enouf, Marie-Anne Rameix-Welti, Alexandre OZANNE |
| EPI_ISL_18568003, EPI_ISL_18568004, EPI_ISL_18568005, EPI_ISL_18568006, EPI_ISL_18568007, EPI_ISL_18568008, EPI_ISL_18568009, EPI_ISL_18568010, EPI_ISL_18568011, EPI_ISL_18568012, EPI_ISL_18568013, EPI_ISL_18568014, EPI_ISL_18568015, EPI_ISL_18568016 | | | |
| see above | RELAB - Biogroup - Laborizon Bretagne | National Reference Center for Viruses of Respiratory Infections, Institut Pasteur, Paris | Léa Avon, Marion Barbet, Marine Bernard, Emma Bezot, Angela Brisebarre, Océane Dehan, Flora Donati, Vanessa Guimaraes, Banujaa Jeyarajah, Florian Préjean, Yannis Rahou, Sylvie van der Werf, Frédéric Lemoine, Samar Berreira Ibraim, Jérôme Bourret, Kévin Da Silva, Maud Vanpeene, Vincent Enouf, Marie-Anne Rameix-Welti, Jean-François COMES |
| EPI_ISL_18568017 | Sentinelles Province | National Reference Center for Viruses of Respiratory Infections, Institut Pasteur, Paris | Léa Avon, Marion Barbet, Marine Bernard, Emma Bezot, Angela Brisebarre, Océane Dehan, Flora Donati, Vanessa Guimaraes, Banujaa Jeyarajah, Florian Préjean, Yannis Rahou, Sylvie van der Werf, Frédéric Lemoine, Samar Berreira Ibraim, Jérôme Bourret, Kévin Da Silva, Maud Vanpeene, Vincent Enouf, Marie-Anne Rameix-Welti, Eric VAN MELKEBEKE |
| EPI_ISL_18568018 | Sentinelles Province | National Reference Center for Viruses of Respiratory Infections, Institut Pasteur, Paris | Léa Avon, Marion Barbet, Marine Bernard, Emma Bezot, Angela Brisebarre, Océane Dehan, Flora Donati, Vanessa Guimaraes, Banujaa Jeyarajah, Florian Préjean, Yannis Rahou, Sylvie van der Werf, Frédéric Lemoine, Samar Berreira Ibraim, Jérôme Bourret, Kévin Da Silva, Maud Vanpeene, Vincent Enouf, Marie-Anne Rameix-Welti, Yannick FREYMANN |
| EPI_ISL_18568019, EPI_ISL_18568020, EPI_ISL_18568021, EPI_ISL_18568022 | RELAB - Medical Analysis Laboratory Cab-Lenys | National Reference Center for Viruses of Respiratory Infections, Institut Pasteur, Paris | Léa Avon, Marion Barbet, Marine Bernard, Emma Bezot, Angela Brisebarre, Océane Dehan, Flora Donati, Vanessa Guimaraes, Banujaa Jeyarajah, Florian Préjean, Yannis Rahou, Sylvie van der Werf, Frédéric Lemoine, Samar Berreira Ibraim, Jérôme Bourret, Kévin Da Silva, Maud Vanpeene, Vincent Enouf, Marie-Anne Rameix-Welti, Nadège GOURGUILLON |
| EPI_ISL_18568023, EPI_ISL_18568024, EPI_ISL_18568025, EPI_ISL_18568026, EPI_ISL_18568027 | RELAB - Selas Biogroup Lorraine | National Reference Center for Viruses of Respiratory Infections, Institut Pasteur, Paris | Léa Avon, Marion Barbet, Marine Bernard, Emma Bezot, Angela Brisebarre, Océane Dehan, Flora Donati, Vanessa Guimaraes, Banujaa Jeyarajah, Florian Préjean, Yannis Rahou, Sylvie van der Werf, Frédéric Lemoine, Samar Berreira Ibraim, Jérôme Bourret, Kévin Da Silva, Maud Vanpeene, Vincent Enouf, Marie-Anne Rameix-Welti, Alexandra JACQUES |
| EPI_ISL_18568028, EPI_ISL_18568029, EPI_ISL_18568030 | Centre Hospitalier (CH) d'Argenteuil (Victor Dupouy) | National Reference Center for Viruses of Respiratory Infections, Institut Pasteur, Paris | Léa Avon, Marion Barbet, Marine Bernard, Emma Bezot, Angela Brisebarre, Océane Dehan, Flora Donati, Vanessa Guimaraes, Banujaa Jeyarajah, Florian Préjean, Yannis Rahou, Sylvie van der Werf, Frédéric Lemoine, Samar Berreira Ibraim, Jérôme Bourret, Kévin Da Silva, Maud Vanpeene, Vincent Enouf, Marie-Anne Rameix-Welti, L COURDAVAULT |
| EPI_ISL_18568031, EPI_ISL_18568032 | Hôpital Ambroise Paré - Service de Microbiologie et hygiène | National Reference Center for Viruses of Respiratory Infections, Institut Pasteur, Paris | Léa Avon, Marion Barbet, Marine Bernard, Emma Bezot, Angela Brisebarre, Océane Dehan, Flora Donati, Vanessa Guimaraes, Banujaa Jeyarajah, Florian Préjean, Yannis Rahou, Sylvie van der Werf, Frédéric Lemoine, Samar Berreira Ibraim, Jérôme Bourret, Kévin Da Silva, Maud Vanpeene, Vincent Enouf, Marie-Anne Rameix-Welti, Elyanne GAULT |
| EPI_ISL_18568033 | Sentinelles Ile-de-France | National Reference Center for Viruses of Respiratory Infections, Institut Pasteur, Paris | Léa Avon, Marion Barbet, Marine Bernard, Emma Bezot, Angela Brisebarre, Océane Dehan, Flora Donati, Vanessa Guimaraes, Banujaa Jeyarajah, Florian Préjean, Yannis Rahou, Sylvie van der Werf, Frédéric Lemoine, Samar Berreira Ibraim, Jérôme Bourret, Kévin Da Silva, Maud Vanpeene, Vincent Enouf, Marie-Anne Rameix-Welti, Hajer KANOUN-DRIRA |
| EPI_ISL_18568034 | Hôpital Necker - Enfants - Malades,Laboratoire De Virologie | National Reference Center for Viruses of Respiratory Infections, Institut Pasteur, Paris | Léa Avon, Marion Barbet, Marine Bernard, Emma Bezot, Angela Brisebarre, Océane Dehan, Flora Donati, Vanessa Guimaraes, Banujaa Jeyarajah, Florian Préjean, Yannis Rahou, Sylvie van der Werf, Frédéric Lemoine, Samar Berreira Ibraim, Jérôme Bourret, Kévin Da Silva, Maud Vanpeene, Vincent Enouf, Marie-Anne Rameix-Welti, Marianne LERUEZ-VILLE |
| EPI_ISL_18568035, EPI_ISL_18568036, EPI_ISL_18568037, EPI_ISL_18568038, EPI_ISL_18568039 | RELAB - BIOGROUP - Plateau technique St Denis | National Reference Center for Viruses of Respiratory Infections, Institut Pasteur, Paris | Léa Avon, Marion Barbet, Marine Bernard, Emma Bezot, Angela Brisebarre, Océane Dehan, Flora Donati, Vanessa Guimaraes, Banujaa Jeyarajah, Florian Préjean, Yannis Rahou, Sylvie van der Werf, Frédéric Lemoine, Samar Berreira Ibraim, Jérôme Bourret, Kévin Da Silva, Maud Vanpeene, Vincent Enouf, Marie-Anne Rameix-Welti, Yanis CHAIB |
| EPI_ISL_18568040, EPI_ISL_18568041 | Sentinelles Ile-de-France | National Reference Center for Viruses of Respiratory Infections, Institut Pasteur, Paris | Léa Avon, Marion Barbet, Marine Bernard, Emma Bezot, Angela Brisebarre, Océane Dehan, Flora Donati, Vanessa Guimaraes, Banujaa Jeyarajah, Florian Préjean, Yannis Rahou, Sylvie van der Werf, Frédéric Lemoine, Samar Berreira Ibraim, Jérôme Bourret, Kévin Da Silva, Maud Vanpeene, Vincent Enouf, Marie-Anne Rameix-Welti, Marie-Anne COSSON |
| EPI_ISL_18568042, EPI_ISL_18568043, EPI_ISL_18568044 | Institut Fédératif De Biologie- Laboratoire De Virologie | National Reference Center for Viruses of Respiratory Infections, Institut Pasteur, Paris | Léa Avon, Marion Barbet, Marine Bernard, Emma Bezot, Angela Brisebarre, Océane Dehan, Flora Donati, Vanessa Guimaraes, Banujaa Jeyarajah, Florian Préjean, Yannis Rahou, Sylvie van der Werf, Frédéric Lemoine, Samar Berreira Ibraim, Jérôme Bourret, Kévin Da Silva, Maud Vanpeene, Vincent Enouf, Marie-Anne Rameix-Welti, Jean-Michel MANSUY |
| EPI_ISL_18568045 | SOS Médecins Nantes | National Reference Center for Viruses of Respiratory Infections, Institut Pasteur, Paris | Léa Avon, Marion Barbet, Marine Bernard, Emma Bezot, Angela Brisebarre, Océane Dehan, Flora Donati, Vanessa Guimaraes, Banujaa Jeyarajah, Florian Préjean, Yannis Rahou, Sylvie van der Werf, Frédéric Lemoine, Samar Berreira Ibraim, Jérôme Bourret, Kévin Da Silva, Maud Vanpeene, Vincent Enouf, Marie-Anne Rameix-Welti, Boris DUMONT |
| EPI_ISL_18568342, EPI_ISL_18569153, EPI_ISL_18569154, EPI_ISL_18569155, EPI_ISL_18569158, EPI_ISL_18569159, EPI_ISL_18569160 | Respiratory Virus Unit / Reference Microbiology Services / UK Health Security Agency | Reference Microbiology Services / UK Health Security Agency | Zambon M. Talts T. Kele B. Miah S |
| EPI_ISL_18578933 | University of Washington, Department of Laboratory Medicine | University of Washington, Department of Laboratory Medicine | Greninger,A.L., Makhsous,N., Kuypers,J., Shean,R.C. and Jerome,K.R. |
| EPI_ISL_18591750, EPI_ISL_18591753, EPI_ISL_18591757, EPI_ISL_18591758, EPI_ISL_18591760, EPI_ISL_18591761, EPI_ISL_18591762, EPI_ISL_18591765, EPI_ISL_18591767, EPI_ISL_18591771, EPI_ISL_18591772, EPI_ISL_18591774, EPI_ISL_18591775, EPI_ISL_18591776, EPI_ISL_18591777, EPI_ISL_18591780, EPI_ISL_18591781, EPI_ISL_18591784 | | | |
| see above | Respiratory Virus Unit / Reference Microbiology Services / UK Health Security Agency | Reference Microbiology Services / UK Health Security Agency | Zambon M. Talts T. Kele B. Miah S |
| EPI_ISL_18592436, EPI_ISL_18592439, EPI_ISL_18592440, EPI_ISL_18592441, EPI_ISL_18592442 | HOSPITAL UNIVERSITARIO CENTRAL DE ASTURIAS | Laboratorio de Virología HUCA | Pérez-Martínez Z, Boga JA, Rojo S, González-Alba JM, Ochoa-Varela C, Rodríguez-Pérez M, Melón S, Alvarez-Argüelles ME |
| EPI_ISL_2156812, EPI_ISL_2156813, EPI_ISL_2156814 | Royal Children's Hospital | WHO Influenza Centre for Reference and Research on Influenza | Angela Todd, Yi-Mo Deng, Annette Alafaci, Naomi Komadina |
| EPI_ISL_2543761 | National Center for Communicable Diseases | WHO Collaborating Centre for Reference and Research on Influenza | Xiaomin Dong, Darmaa Bardach, Yi-Mo Deng, Ammar Aziz, Naomi Komadina |
| EPI_ISL_2543762, EPI_ISL_2543764, EPI_ISL_2543765, EPI_ISL_2543766, EPI_ISL_2543767, EPI_ISL_2543768, EPI_ISL_2543769, EPI_ISL_2543770, EPI_ISL_2543771, EPI_ISL_2543772, EPI_ISL_2543773, EPI_ISL_2543774, EPI_ISL_2543775, EPI_ISL_2543776, EPI_ISL_2543777, EPI_ISL_2543778, EPI_ISL_2543779, EPI_ISL_2543780, EPI_ISL_2543781 | | | |
| see above | Monash Medical Centre | WHO Collaborating Centre for Reference and Research on Influenza | Xiaomin Dong, Michelle Francis, Tony Korman, Yi-Mo Deng, Ammar Aziz, Naomi Komadina |
| EPI_ISL_2543782, EPI_ISL_2543783, EPI_ISL_2543784, EPI_ISL_2543785, EPI_ISL_2543786, EPI_ISL_2543787, EPI_ISL_2543788, EPI_ISL_2543789, EPI_ISL_2543790, EPI_ISL_2543791, EPI_ISL_2543792, EPI_ISL_2543793, EPI_ISL_2543794, EPI_ISL_2543795, EPI_ISL_2543796, EPI_ISL_2543797, EPI_ISL_2543798, EPI_ISL_2543799, EPI_ISL_2543800, EPI_ISL_2543801, EPI_ISL_2543802, EPI_ISL_2543803, EPI_ISL_2543804, EPI_ISL_2543805, EPI_ISL_2543806, EPI_ISL_2543807, EPI_ISL_2543808, EPI_ISL_2543809, EPI_ISL_2543810 | | | |
| see above | Royal Children's Hospital | WHO Collaborating Centre for Reference and Research on Influenza | Xiaomin Dong, Annette Alafaci,Yi-Mo Deng, Ammar Aziz, Naomi Komadina |
| EPI_ISL_2543811, EPI_ISL_2543812, EPI_ISL_2543813, EPI_ISL_2543814, EPI_ISL_2543815, EPI_ISL_2543816, EPI_ISL_2543817, EPI_ISL_2543818, EPI_ISL_2543819, EPI_ISL_2543820, EPI_ISL_2543821, EPI_ISL_2543822, EPI_ISL_2543823, EPI_ISL_2543824, EPI_ISL_2543825, EPI_ISL_2543826 | | | |
| see above | Institut Pasteur de Cote d'Ivoire | WHO Collaborating Centre for Reference and Research on Influenza | Xiaomin Dong, Herve Kadjo, Yi-Mo Deng, Ammar Aziz, Naomi Komadina |
| EPI_ISL_2543827, EPI_ISL_2543828, EPI_ISL_2543829, EPI_ISL_2543830, EPI_ISL_2543831, EPI_ISL_2543832, EPI_ISL_2543833, EPI_ISL_2543834, EPI_ISL_2543835, EPI_ISL_2543836, EPI_ISL_2543837, EPI_ISL_2543838, EPI_ISL_2543839, EPI_ISL_2543840, EPI_ISL_2543841, EPI_ISL_2543842, EPI_ISL_2543843, EPI_ISL_2543844 | | | |
| see above | National Center for Communicable Diseases | WHO Collaborating Centre for Reference and Research on Influenza | Xiaomin Dong, Darmaa Bardach, Yi-Mo Deng, Ammar Aziz, Naomi Komadina |
| EPI_ISL_2543845, EPI_ISL_2543846, EPI_ISL_2543847 | NIC, National Institute of Health | WHO Collaborating Centre for Reference and Research on Influenza | Xiaomin Dong, Pilailuk Akkapaiboon Okada, Yi-Mo Deng, Ammar Aziz, Naomi Komadina |
| EPI_ISL_2543922 | KEMRI Wellcome Trust Research Programme | KEMRI Wellcome Trust Research Programme | Agoti,C.N., Otieno,J.R., Munywoki,P.K., Mwihuri,A.G., Cane,P.A., Nokes,D.J., Kellam,P. and Cotten,M.L. |
| EPI_ISL_2543931 | Pathogen Diagnostic Center, Institut Pasteur of Shanghai, Chinese Academy of Sciences | Pathogen Diagnostic Center, Institut Pasteur of Shanghai, Chinese Academy of Sciences | Fu,X., He,Z., Lan,K., Zhang,C., Dong,W., Cheng,Y. and Hu,Y. |
| EPI_ISL_2543932 | Microbiology Department, The University of Hong Kong, Queen Mary Hospital, University Pathology Building | Microbiology Department, The University of Hong Kong, Queen Mary Hospital, University Pathology Building | Zhang,K., He,J., Cheng,Z., Zhou,J., Bose,M., Henrickson,K.J. and Zheng,B. |
| EPI_ISL_2543933 | Influenza Group, National Institute of Virology | Influenza Group, National Institute of Virology | Choudhary,M.L., Wadhwa,B., Jadhav,S.M., Chadha,M.S. and Mourya,D.T. |
| EPI_ISL_2543934, EPI_ISL_2543935, EPI_ISL_2543936 | J. Craig Venter Institute | J. Craig Venter Institute | Shabman,R., Das,S.R., Puri,V., Fedorova,N., Amedeo,P., Williams,M., Shrivastava,S. and Halasa,N. |
| EPI_ISL_2543937, EPI_ISL_2543938, EPI_ISL_2543939, EPI_ISL_2543940 | Virology, Graduate School of Medicine, Tohoku University | Virology, Graduate School of Medicine, Tohoku University | Malasao,R., Furuse,Y., Okamoto,M., Dapat,C., Saito,M., Saito-Obata,M., Tamaki,R., Segubre-Mercado,E., Lupisan,S. and Oshitani,H. |
| EPI_ISL_2543941 | Division of Biosafety Evaluation and Control, Korea National Institute of Health, Korea Centers for Disease Control and Prevention | Division of Biosafety Evaluation and Control, Korea National Institute of Health, Korea Centers for Disease Control and Prevention | Yun,M.-R., Lee,W.-J., Kim,A.-R., Lee,H.S., Kim,K., Kim,S.S., Kim,Y.-J. and Kim,D.-W. |
| EPI_ISL_2543943 | Emerging Viral Infections, Oxford University Clinical | Emerging Viral Infections, Oxford University Clinical | Do,L.A.H., Wilm,A., van Doorn,H.R., Lam,H.M., Sukumaran,R., Tran,A.T., Nguyen,B.H., Tran,T.T.L., Tran,Q.H., Vo,Q.B., Tran Dac,N.A., Trinh,H.N., Nguyen,t.T.H., Le Binh,B.T., Le,K., Nguyen,M.T., Thai,Q.T., Vo,T.V., Ngo,N.Q.M., |

| | | | |
|---|---|---|---|
| | Research Unit | Research Unit | Dang,t.K.H., Cao,N.H., Tran,T.V., Ho,L.V., Farrar,J., de Jong,M.D., Chen,S., Nagaranjan,N., Bryant,J.E. and Hibberd,M.L. |
| EPI_ISL_2543949 | Medical Microbiology, University Medical Center Utrecht | Medical Microbiology, University Medical Center Utrecht | Tan,L., Lemey,P., Viveen,M. and Coenjaerts,F.E.J. |
| EPI_ISL_2543952, EPI_ISL_2543953, EPI_ISL_2543954 | J. Craig Venter Institute | J. Craig Venter Institute | Lorenzi,H., Town,C., Halpin,R., Bera,J., Ransier,A., Fedorova,N., Stockwell,T., Amedeo,P., Appalla,L., Bishop,B., Edworthy,P., Gupta,N., Hoover,J., Katzel,D., Li,K., Schobel,S., Shrivastava,S., Thovarai,V., Wang,S., Rebuffo-Scheer,C., Fan,J., He,J., Kehl,S.C., Lederboer,N., Jurgens,L.A., Bose,M.E., Beck,E.T., Kumar,S., Gerna,G., Wentworth,D.E. and Henrickson,K.J. |
| EPI_ISL_2543955 | Medical Microbiology, University Medical Center Utrecht | Medical Microbiology, University Medical Center Utrecht | Tan,L., Lemey,P., Viveen,M. and Coenjaerts,F. |
| EPI_ISL_2543956 | Medical Microbiology, University Medical Center Utrecht | Medical Microbiology, University Medical Center Utrecht | Tan,L., Lemey,P., Viveen,M. and Coenjaerts,F.E.J. |
| EPI_ISL_2543957 | Medical Microbiology, University Medical Center Utrecht | Medical Microbiology, University Medical Center Utrecht | Tan,L., Lemey,P., Viveen,M. and Coenjaerts,F. |
| EPI_ISL_2543958, EPI_ISL_2543959, EPI_ISL_2543960 | Medical Microbiology, University Medical Center Utrecht | Medical Microbiology, University Medical Center Utrecht | Tan,L., Lemey,P., Viveen,M. and Coenjaerts,F.E.J. |
| EPI_ISL_2543967 | J. Craig Venter Institute | J. Craig Venter Institute | Lorenzi,H., Town,C., Halpin,R., Bera,J., Ransier,A., Fedorova,N., Stockwell,T., Amedeo,P., Appalla,L., Bishop,B., Edworthy,P., Gupta,N., Hoover,J., Katzel,D., Li,K., Schobel,S., Shrivastava,S., Thovarai,V., Wang,S., Rebuffo-Scheer,C., Fan,J., He,J., Kehl,S.C., Lederboer,N., Jurgens,L.A., Bose,M.E., Beck,E.T., Kumar,S., Noyola,D.E., Wentworth,D.E. and Henrickson,K.J. |
| EPI_ISL_2543969 | Mirus Bio Corporation | Mirus Bio Corporation | Brazas,R.M. |
| EPI_ISL_2543970, EPI_ISL_2543971, EPI_ISL_2543972, EPI_ISL_2543973, EPI_ISL_2543974, EPI_ISL_2543975, EPI_ISL_2543976, EPI_ISL_2543977, EPI_ISL_2543978, EPI_ISL_2543979, EPI_ISL_2543980, EPI_ISL_2543981, EPI_ISL_2543982 | | | |
| see above | J. Craig Venter Institute | J. Craig Venter Institute | Das,S.R., Halpin,R.A., Shilts,M., Puri,V., Akopov,A., Fedorova,N., Stockwell,T., Amedeo,P., Bishop,B., Katzel,D., Schobel,S., Shrivastava,S. and Hartert,T. |
| EPI_ISL_2543983 | J. Craig Venter Institute | J. Craig Venter Institute | Shabman,R., Das,S.R., Shilts,M., Fedorova,N., Puri,V., Shrivastava,S., Amedeo,P., Hu,L., Durbin,A., Rocchi,I., Williams,T. and Hartert,T. |
| EPI_ISL_2543984, EPI_ISL_2543985, EPI_ISL_2543986, EPI_ISL_2543987, EPI_ISL_2543988, EPI_ISL_2543989, EPI_ISL_2543990 | Lab Medicine, UW | Lab Medicine, UW | Greninger,A.L., Makhsous,N., Kuypers,J.M., Shean,R.C. and Jerome,K.R. |
| EPI_ISL_2543991, EPI_ISL_2543992, EPI_ISL_2543993, EPI_ISL_2543995 | J. Craig Venter Institute | J. Craig Venter Institute | Das,S.R., Halpin,R.A., Puri,V., Akopov,A., Fedorova,N., Tsitrin,T., Stockwell,T., Amedeo,P., Bishop,B., Gupta,N., Hoover,J., Katzel,D., Schobel,S., Shrivastava,S., Wentworth,D.E. and Caserta,M. |
| EPI_ISL_2543996, EPI_ISL_2543997, EPI_ISL_2543998, EPI_ISL_2543999, EPI_ISL_2544000, EPI_ISL_2544001, EPI_ISL_2544002, EPI_ISL_2544003 | J. Craig Venter Institute | J. Craig Venter Institute | Das,S.R., Halpin,R.A., Puri,V., Akopov,A., Fedorova,N., Stockwell,T., Amedeo,P., Bishop,B., Katzel,D., Schobel,S., Shrivastava,S., Wentworth,D.E. and Caserta,M. |
| EPI_ISL_2544004, EPI_ISL_2544005, EPI_ISL_2544006, EPI_ISL_2544007, EPI_ISL_2544008, EPI_ISL_2544009, EPI_ISL_2544010, EPI_ISL_2544011, EPI_ISL_2544012, EPI_ISL_2544013 | J. Craig Venter Institute | J. Craig Venter Institute | Das,S.R., Halpin,R.A., Puri,V., Akopov,A., Fedorova,N., Stockwell,T., Amedeo,P., Bishop,B., Katzel,D., Schobel,S., Shrivastava,S., Hall,C.B., Tesini,B.L., Schnabel,K.C., Walsh,E.E. and Caserta,M. |
| EPI_ISL_2544014, EPI_ISL_2544015, EPI_ISL_2544016, EPI_ISL_2544017, EPI_ISL_2544018, EPI_ISL_2544019 | J. Craig Venter Institute | J. Craig Venter Institute | Shabman,R., Fedorova,N., Puri,V., Shrivastava,S., Amedeo,P., Isom,R., Hu,L., Pickett,B., Novotny,M., Durbin,A., Rocchi,I., Williams,T., Hall,C.B., Tesini,B.L., Schnabel,K.C., Walsh,E.E. and Caserta,M. |
| EPI_ISL_2544020 | Broad Institute of MIT & Harvard | Broad Institute of MIT & Harvard | Newman,R.M., Zody,M.C., DeVincenzo,J.P., Grad,Y., Lipsitch,M., Murphy,R., Fitzgerald,M., Young,S., Gargeya,S., Poon,T.W., Charlebois,P., Weiner,B., Yang,X., Piper,M.E., McCowan,C., Ireland,A., Levin,J., Malboeuf,C., Qu,J., Chapman,S.B., Murphy,C., Wortman,J., Nusbaum,C. and Birren,B. |
| EPI_ISL_2544021, EPI_ISL_2544022, EPI_ISL_2544023, EPI_ISL_2544024 | J. Craig Venter Institute | J. Craig Venter Institute | Das,S., Halpin,R.A., Bera,J., Fedorova,N., Tsitrin,T., Stockwell,T., Amedeo,P., Bishop,B., Gupta,N., Hoover,J., Katzel,D., Schobel,S., Shrivastava,S., Hartert,T., Moore,M., Chappell,J., Larkin,E., Wentworth,D.E. and Anderson,L.J. |
| EPI_ISL_2544025 | Infectious Diseases, St. Jude Children's Research Hospital | Infectious Diseases, St. Jude Children's Research Hospital | DeVincenzo,J., Jones,B.G., Kim,Y.-I. and Hurwitz,J.L. |
| EPI_ISL_2544026, EPI_ISL_2544027, EPI_ISL_2544028 | J. Craig Venter Institute | J. Craig Venter Institute | Lorenzi,H., Halpin,R., Nguyen,A., Fedorova,N., Overton,L., Stockwell,T., Amedeo,P., Bishop,B., Chen,H., Edworthy,P., Gupta,N., Katzel,D., Li,K., Schobel,S., Shrivastava,S., Thovarai,V., Wang,S., Shapiro,C.A., Williams,J., Peebles,S., Chi,M., Wentworth,D.E. and Moore,M.L. |
| EPI_ISL_2544029 | J. Craig Venter Institute | J. Craig Venter Institute | Lorenzi,H., Town,C., Halpin,R., Bera,J., Ransier,A., Fedorova,N., Stockwell,T., Amedeo,P., Appalla,L., Bishop,B., Edworthy,P., Gupta,N., Hoover,J., Katzel,D., Li,K., Schobel,S., Shrivastava,S., Thovarai,V., Wang,S., Rebuffo-Scheer,C., Fan,J., He,J., Kehl,S.C., Lederboer,N., Jurgens,L.A., Bose,M.E., Beck,E.T., Kumar,S., Wentworth,D.E. and Henrickson,K.J. |
| EPI_ISL_2544030, EPI_ISL_2544031 | Pediatrics - Infectious Diseases, Medical College of Wisconsin | Pediatrics - Infectious Diseases, Medical College of Wisconsin | Rebuffo-Scheer,C., Bose,M.E., He,J., Khaja,S., Ulatowski,M., Beck,E.T., Fan,J., Kumar,S., Nelson,M.I. and Henrickson,K.J. |
| EPI_ISL_2544032 | Molecular Virology and Microbiology, Baylor College of Medicine | Molecular Virology and Microbiology, Baylor College of Medicine | Tapia,L.I. and Piedra,P.A. |
| EPI_ISL_2544033 | J. Craig Venter Institute | J. Craig Venter Institute | Lorenzi,H., Town,C., Halpin,R., Bera,J., Ransier,A., Fedorova,N., Stockwell,T., Amedeo,P., Appalla,L., Bishop,B., Edworthy,P., Gupta,N., Hoover,J., Katzel,D., Li,K., Schobel,S., Shrivastava,S., Thovarai,V., Wang,S., Rebuffo-Scheer,C., Fan,J., He,J., Kehl,S.C., Lederboer,N., Jurgens,L.A., Bose,M.E., Beck,E.T., Kumar,S., Kok,T., Wentworth,D.E. and Henrickson,K.J. |
| EPI_ISL_2544034, EPI_ISL_2544035, EPI_ISL_2544036, EPI_ISL_2544037, EPI_ISL_2544038, EPI_ISL_2544039, EPI_ISL_2544040, EPI_ISL_2544041 | J. Craig Venter Institute | J. Craig Venter Institute | Shabman,R., Das,S.R., Shilts,M., Fedorova,N., Puri,V., Shrivastava,S., Amedeo,P., Williams,M., Barratt,K., Mitchell,J. and Jennings,L. |
| EPI_ISL_2544044 | J. Craig Venter Institute | J. Craig Venter Institute | Lorenzi,H., Town,C., Halpin,R., Bera,J., Ransier,A., Fedorova,N., Stockwell,T., Amedeo,P., Appalla,L., Bishop,B., Edworthy,P., Gupta,N., Hoover,J., Katzel,D., Schobel,S., Shrivastava,S., Thovarai,V., Wang,S., Rebuffo-Scheer,C., Fan,J., He,J., Kehl,S.C., Lederboer,N., Jurgens,L.A., Bose,M.E., Beck,E.T., Kumar,S., Videla,C., Wentworth,D.E. and Henrickson,K.J. |
| EPI_ISL_2544045, EPI_ISL_2544046 | J. Craig Venter Institute | J. Craig Venter Institute | Lorenzi,H., Town,C., Halpin,R., Bera,J., Ransier,A., Fedorova,N., Stockwell,T., Amedeo,P., Appalla,L., Bishop,B., Edworthy,P., Gupta,N., Hoover,J., Katzel,D., Li,K., Schobel,S., Shrivastava,S., Thovarai,V., Wang,S., Rebuffo-Scheer,C., Fan,J., He,J., Kehl,S.C., Lederboer,N., Jurgens,L.A., Bose,M.E., Beck,E.T., Kumar,S., Videla,C., Wentworth,D.E. and Henrickson,K.J. |
| EPI_ISL_2544051 | Virology Laboratory, Dr. Ricardo Gutierrez Children Hospital | Virology Laboratory, Dr. Ricardo Gutierrez Children Hospital | Goya,S., Valinotto,L.E., Tittarelli,E., Rojo,G.L., Greninger,A., Luso,S., Natale,M., Mistchenko,A.S. and Viegas,M. |
| EPI_ISL_2544052 | Virology Laboratory, Dr. Ricardo Gutierrez Children Hospital | Virology Laboratory, Dr. Ricardo Gutierrez Children Hospital | Goya,S., Valinotto,L.E., Tittarelli,E., Rojo,G.L., Greninger,A., Zaiat,J., Marti,M., Mistchenko,A.S. and Viegas,M. |
| EPI_ISL_2544055, EPI_ISL_2544056, EPI_ISL_2544057, EPI_ISL_2544058, EPI_ISL_2544059, EPI_ISL_2544060, EPI_ISL_2544061, EPI_ISL_2544062, EPI_ISL_2544063, EPI_ISL_2544064 | J. Craig Venter Institute | J. Craig Venter Institute | Wentworth,D.E., Halpin,R.A., Bera,J., Lin,X., Fedorova,N., Tsitrin,T., McLellan,M., Stockwell,T., Amedeo,P., Bishop,B., Gupta,N., Hoover,J., Katzel,D., Schobel,S., Shrivastava,S., Garcia,J., Laguna-Torres,V.A., Leguia,M., Benavides,J.G. and Halsey,E. |
| EPI_ISL_2574728 | Epidemiology and Demography Department, KEMRI-Wellcome Trust Research Programme | Epidemiology and Demography Department, KEMRI-Wellcome Trust Research Programme | Otieno,J.R., Kamau,E.M., Oketch,J.W., Ngoi,J.M., Agoti,C.N., Gichuki,A.M., Otieno,G.P., Ngama,M., Cane,P.A., Kellam,P., Cotten,M., Lemey,P. and Nokes,D.J. |
| EPI_ISL_2578661 | Pediatrics - Infectious Diseases, Medical College of Wisconsin | Pediatrics - Infectious Diseases, Medical College of Wisconsin | Rebuffo-Scheer,C., Bose,M.E., He,J., Khaja,S., Ulatowski,M., Beck,E.T., Fan,J., Kumar,S., Nelson,M.I. and Henrickson,K.J. |
| EPI_ISL_2578662 | Beijing Key Laboratory of Etiology of Viral Diseases in Children; Laboratory of Virology, Capital Institute of Pediatrics | Beijing Key Laboratory of Etiology of Viral Diseases in Children; Laboratory of Virology, Capital Institute of Pediatrics | Cui,G., Zhu,R., Deng,J., Zhao,L., Sun,Y., Wang,F. and Qian,Y. |
| EPI_ISL_2578663, EPI_ISL_2578664, EPI_ISL_2578665, EPI_ISL_2578666, EPI_ISL_2578667, EPI_ISL_2578668, EPI_ISL_2578669, EPI_ISL_2578670, EPI_ISL_2578671, EPI_ISL_2578672, EPI_ISL_2578673, EPI_ISL_2578674, EPI_ISL_2578675, EPI_ISL_2578676 | | | |
| see above | Epidemiology and Demography Department, KEMRI-Wellcome Trust Research Programme | Epidemiology and Demography Department, KEMRI-Wellcome Trust Research Programme | Otieno,J.R., Kamau,E.M., Oketch,J.W., Ngoi,J.M., Agoti,C.N., Gichuki,A.M., Otieno,G.P., Ngama,M., Cane,P.A., Kellam,P., Cotten,M., Lemey,P. and Nokes,D.J. |
| EPI_ISL_2578677, EPI_ISL_2578678 | Center for Infectious Diseases, School of Public Health, University of Texas Health Science Center | Center for Infectious Diseases, School of Public Health, University of Texas Health Science Center | Bahl,J., Hixson,J., Kim,D.-K., Qiu,X., Piedra,P.A., Piedra,F.-A., Avadhanula,V. and Machado,A.A. |
| EPI_ISL_2578679 | Marie Bashir Institute for Infectious Diseases and Biosecurity & Sydney Medical School, The University of Sydney, Westmead Institute for Medical Research | Marie Bashir Institute for Infectious Diseases and Biosecurity & Sydney Medical School, The University of Sydney, Westmead Institute for Medical Research | Eden,J.-S., Kok,J., Dwyer,D.E., Fernandez,M., Carter,I. and Holmes,E.C. |
| EPI_ISL_2578680, EPI_ISL_2578681, EPI_ISL_2578682, EPI_ISL_2578683, EPI_ISL_2578684, EPI_ISL_2578685, EPI_ISL_2578686, EPI_ISL_2578687, EPI_ISL_2578688, EPI_ISL_2578689, EPI_ISL_2578690 | | | |
| see above | Epidemiology and Demography Department, KEMRI-Wellcome Trust Research Programme | Epidemiology and Demography Department, KEMRI-Wellcome Trust Research Programme | Otieno,J.R., Kamau,E.M., Oketch,J.W., Ngoi,J.M., Agoti,C.N., Gichuki,A.M., Otieno,G.P., Ngama,M., Cane,P.A., Kellam,P., Cotten,M., Lemey,P. and Nokes,D.J. |
| EPI_ISL_2578691 | Department of Pediatrics, Center of Excellence in Clinical Virology, Chulalongkorn | Department of Pediatrics, Center of Excellence in Clinical Virology, Chulalongkorn | Thongpan,I. |
| EPI_ISL_2578692 | Center for Infectious Diseases, School of Public Health, University of Texas Health Science Center | Center for Infectious Diseases, School of Public Health, University of Texas Health Science Center | Bahl,J., Hixson,J., Kim,D.-K., Qiu,X., Piedra,P.A., Piedra,F.-A., Avadhanula,V. and Machado,A.A. |
| EPI_ISL_2578693, EPI_ISL_2578694, EPI_ISL_2578695, EPI_ISL_2578696, EPI_ISL_2578697, EPI_ISL_2578698, EPI_ISL_2578699, EPI_ISL_2578700, EPI_ISL_2578701, EPI_ISL_2578702, EPI_ISL_2578703, EPI_ISL_2578704, EPI_ISL_2578705, EPI_ISL_2578706, EPI_ISL_2578707, EPI_ISL_2578708, EPI_ISL_2578709, EPI_ISL_2578710, EPI_ISL_2578711 | | | |
| see above | Epidemiology and Demography Department, KEMRI-Wellcome Trust Research Programme | Epidemiology and Demography Department, KEMRI-Wellcome Trust Research Programme | Otieno,J.R., Kamau,E.M., Oketch,J.W., Ngoi,J.M., Agoti,C.N., Gichuki,A.M., Otieno,G.P., Ngama,M., Cane,P.A., Kellam,P., Cotten,M., Lemey,P. and Nokes,D.J. |
| EPI_ISL_2578712 | Pediatrics, University of New Mexico | Pediatrics, University of New Mexico | Kothari,A., Kennedy,J.L., Schwalm,K.C., Putt,C., Denson,J.L. and Dinwiddie,D.L. |

| | | | |
|---|---|---|---|
| EPI_ISL_2578713, EPI_ISL_2578714 | Center for Infectious Diseases, School of Public Health, University of Texas Health Science Center | Center for Infectious Diseases, School of Public Health, University of Texas Health Science Center | Bahl,J., Hixson,J., Kim,D.-K., Qiu,X., Piedra,P.A., Piedra,F.-A., Avadhanula,V. and Machado,A.A. |
| EPI_ISL_2578715, EPI_ISL_2578716, EPI_ISL_2578717 | Marie Bashir Institute for Infectious Diseases and Biosecurity & Sydney Medical School, The University of Sydney, Westmead Institute for Medical Research | Marie Bashir Institute for Infectious Diseases and Biosecurity & Sydney Medical School, The University of Sydney, Westmead Institute for Medical Research | Eden,J.-S., Kok,J., Dwyer,D.E., Fernandez,M., Carter,I. and Holmes,E.C. |
| EPI_ISL_2578718, EPI_ISL_2578719, EPI_ISL_2578720, EPI_ISL_2578721, EPI_ISL_2578722, EPI_ISL_2578723, EPI_ISL_2578724, EPI_ISL_2578725, EPI_ISL_2578726, EPI_ISL_2578727, EPI_ISL_2578728, EPI_ISL_2578729, EPI_ISL_2578730, EPI_ISL_2578731, EPI_ISL_2578732, EPI_ISL_2578733, EPI_ISL_2578734 | | | |
| see above | Epidemiology and Demography Department, KEMRI-Wellcome Trust Research Programme | Epidemiology and Demography Department, KEMRI-Wellcome Trust Research Programme | Otieno,J.R., Kamau,E.M., Oketch,J.W., Ngoi,J.M., Agoti,C.N., Gichuki,A.M., Otieno,G.P., Ngama,M., Cane,P.A., Kellam,P., Cotten,M., Lemey,P. and Nokes,D.J. |
| EPI_ISL_2578767, EPI_ISL_2578768 | Marie Bashir Institute for Infectious Diseases and Biosecurity & Sydney Medical School, The University of Sydney, Westmead Institute for Medical Research | Marie Bashir Institute for Infectious Diseases and Biosecurity & Sydney Medical School, The University of Sydney, Westmead Institute for Medical Research | Eden,J.-S., Kok,J., Dwyer,D.E., Fernandez,M., Carter,I. and Holmes,E.C. |
| EPI_ISL_2578769, EPI_ISL_2578770, EPI_ISL_2578771, EPI_ISL_2578772, EPI_ISL_2578773, EPI_ISL_2578774, EPI_ISL_2578775, EPI_ISL_2578776 | Epidemiology and Demography Department, KEMRI-Wellcome Trust Research Programme | Epidemiology and Demography Department, KEMRI-Wellcome Trust Research Programme | Otieno,J.R., Kamau,E.M., Oketch,J.W., Ngoi,J.M., Agoti,C.N., Gichuki,A.M., Otieno,G.P., Ngama,M., Cane,P.A., Kellam,P., Cotten,M., Lemey,P. and Nokes,D.J. |
| EPI_ISL_2578777, EPI_ISL_2578778 | J. Craig Venter Institute | J. Craig Venter Institute | Tan,G., Pickett,B., Fedorova,N., Amedeo,P., Hu,L., Christensen,J., Miller,J., Durbin,A., Williams,T., Arumemi,F., Cadiz,C., Alanis,R., Balmseda,A., Wiliams,T., Schiller,A., Patel,M., Kubale,J. and Gordon,A. |
| EPI_ISL_2578779, EPI_ISL_2578780 | Virology, Graduate School of Medicine, Tohoku University | Virology, Graduate School of Medicine, Tohoku University | Malasao,R., Furuse,Y., Okamoto,M., Dapat,C., Saito,M., Saito-Obata,M., Tamaki,R., Segubre-Mercado,E., Lupisan,S. and Oshitani,H. |
| EPI_ISL_2578787 | Department of Pediatrics, Center of Excellence in Clinical Virology, Chulalongkorn | Department of Pediatrics, Center of Excellence in Clinical Virology, Chulalongkorn | Thongpan,I. |
| EPI_ISL_2578788, EPI_ISL_2578789 | Marie Bashir Institute for Infectious Diseases and Biosecurity & Sydney Medical School, The University of Sydney, Westmead Institute for Medical Research | Marie Bashir Institute for Infectious Diseases and Biosecurity & Sydney Medical School, The University of Sydney, Westmead Institute for Medical Research | Eden,J.-S., Kok,J., Dwyer,D.E., Fernandez,M., Carter,I. and Holmes,E.C. |
| EPI_ISL_2578799, EPI_ISL_2578800, EPI_ISL_2578801 | Epidemiology and Demography Department, KEMRI-Wellcome Trust Research Programme | Epidemiology and Demography Department, KEMRI-Wellcome Trust Research Programme | Otieno,J.R., Kamau,E.M., Oketch,J.W., Ngoi,J.M., Agoti,C.N., Gichuki,A.M., Otieno,G.P., Ngama,M., Cane,P.A., Kellam,P., Cotten,M., Lemey,P. and Nokes,D.J. |
| EPI_ISL_2578802 | Virology, Graduate School of Medicine, Tohoku University | Virology, Graduate School of Medicine, Tohoku University | Malasao,R., Furuse,Y., Okamoto,M., Dapat,C., Saito,M., Saito-Obata,M., Tamaki,R., Segubre-Mercado,E., Lupisan,S. and Oshitani,H. |
| EPI_ISL_2578803 | Department of Pediatrics, Center of Excellence in Clinical Virology, Chulalongkorn | Department of Pediatrics, Center of Excellence in Clinical Virology, Chulalongkorn | Thongpan,I. |
| EPI_ISL_2578804, EPI_ISL_2578805, EPI_ISL_2578806, EPI_ISL_2578807, EPI_ISL_2578808 | Epidemiology and Demography Department, KEMRI-Wellcome Trust Research Programme | Epidemiology and Demography Department, KEMRI-Wellcome Trust Research Programme | Otieno,J.R., Kamau,E.M., Oketch,J.W., Ngoi,J.M., Agoti,C.N., Gichuki,A.M., Otieno,G.P., Ngama,M., Cane,P.A., Kellam,P., Cotten,M., Lemey,P. and Nokes,D.J. |
| EPI_ISL_2578809, EPI_ISL_2578810 | Virology, Graduate School of Medicine, Tohoku University | Virology, Graduate School of Medicine, Tohoku University | Malasao,R., Furuse,Y., Okamoto,M., Dapat,C., Saito,M., Saito-Obata,M., Tamaki,R., Segubre-Mercado,E., Lupisan,S. and Oshitani,H. |
| EPI_ISL_2578811, EPI_ISL_2578812, EPI_ISL_2578813 | Marie Bashir Institute for Infectious Diseases and Biosecurity & Sydney Medical School, The University of Sydney, Westmead Institute for Medical Research | Marie Bashir Institute for Infectious Diseases and Biosecurity & Sydney Medical School, The University of Sydney, Westmead Institute for Medical Research | Eden,J.-S., Kok,J., Dwyer,D.E., Fernandez,M., Carter,I. and Holmes,E.C. |
| EPI_ISL_2578814, EPI_ISL_2578815 | Epidemiology and Demography Department, KEMRI-Wellcome Trust Research Programme | Epidemiology and Demography Department, KEMRI-Wellcome Trust Research Programme | Otieno,J.R., Kamau,E.M., Oketch,J.W., Ngoi,J.M., Agoti,C.N., Gichuki,A.M., Otieno,G.P., Ngama,M., Cane,P.A., Kellam,P., Cotten,M., Lemey,P. and Nokes,D.J. |
| EPI_ISL_2578816 | J. Craig Venter Institute | J. Craig Venter Institute | Tan,G., Pickett,B., Fedorova,N., Amedeo,P., Hu,L., Christensen,J., Miller,J., Durbin,A., Williams,T., Arumemi,F., Cadiz,C., Alanis,R., Balmseda,A., Wiliams,T., Schiller,A., Patel,M., Kubale,J. and Gordon,A. |
| EPI_ISL_2578817 | Department of Pediatrics, Center of Excellence in Clinical Virology, Chulalongkorn | Department of Pediatrics, Center of Excellence in Clinical Virology, Chulalongkorn | Thongpan,I. |
| EPI_ISL_2578818, EPI_ISL_2578819, EPI_ISL_2578820, EPI_ISL_2578821 | Epidemiology and Demography Department, KEMRI-Wellcome Trust Research Programme | Epidemiology and Demography Department, KEMRI-Wellcome Trust Research Programme | Otieno,J.R., Kamau,E.M., Oketch,J.W., Ngoi,J.M., Agoti,C.N., Gichuki,A.M., Otieno,G.P., Ngama,M., Cane,P.A., Kellam,P., Cotten,M., Lemey,P. and Nokes,D.J. |
| EPI_ISL_2578825, EPI_ISL_2578826 | Marie Bashir Institute for Infectious Diseases and Biosecurity & Sydney Medical School, The University of Sydney, Westmead Institute for Medical Research | Marie Bashir Institute for Infectious Diseases and Biosecurity & Sydney Medical School, The University of Sydney, Westmead Institute for Medical Research | Eden,J.-S., Kok,J., Dwyer,D.E., Fernandez,M., Carter,I. and Holmes,E.C. |
| EPI_ISL_2579776 | Center for Infectious Diseases, School of Public Health, University of Texas Health Science Center | Center for Infectious Diseases, School of Public Health, University of Texas Health Science Center | Bahl,J., Hixson,J., Kim,D.-K., Qiu,X., Piedra,P.A., Piedra,F.-A., Avadhanula,V. and Machado,A.A. |
| EPI_ISL_2579857 | Influenza Group, National Institute of Virology | Influenza Group, National Institute of Virology | Choudhary,M.L., Wadhwa,B., Jadhav,S.M., Chadha,M.S. and Mourya,D.T. |
| EPI_ISL_2579858, EPI_ISL_2579859, EPI_ISL_2579860 | Center for Infectious Diseases, School of Public Health, University of Texas Health Science Center | Center for Infectious Diseases, School of Public Health, University of Texas Health Science Center | Bahl,J., Hixson,J., Kim,D.-K., Qiu,X., Piedra,P.A., Piedra,F.-A., Avadhanula,V. and Machado,A.A. |
| EPI_ISL_2579886, EPI_ISL_2579887, EPI_ISL_2579888, EPI_ISL_2579889, EPI_ISL_2579890, EPI_ISL_2579891, EPI_ISL_2579892, EPI_ISL_2579894, EPI_ISL_2579895, EPI_ISL_2579896, EPI_ISL_2579897 | | | |
| see above | Broad Institute of MIT & Harvard | Broad Institute of MIT & Harvard | Newman,R.M., Zody,M.C., DeVincenzo,J.P., Grad,Y., Lipsitch,M., Murphy,R., Fitzgerald,M., Young,S., Gargeya,S., Poon,T.W., Charlebois,P., Weiner,B., Yang,X., Piper,M.E., McCowan,C., Ireland,A., Levin,J., Malboeuf,C., Qu,J., Chapman,S.B., Murphy,C., Wortman,J., Nusbaum,C. and Birren,B. |
| EPI_ISL_2579900 | J. Craig Venter Institute | J. Craig Venter Institute | Shrivastava,S., Halpin,R.A., Puri,V., Fedorova,N.B., Stockwell,T., Amedeo,P., Katzel,D., Schobel,S., Pickett,B.E., Moore,M., Chappell,J., Larkin,E., Wentworth,D.E., Anderson,L.J. and Hartert,T. |
| EPI_ISL_2582147 | Experimental Pathology, Immunology, and Microbiology, American University of Beirut | Experimental Pathology, Immunology, and Microbiology, American University of Beirut | Hamdan,F., Ezzeddine,A., Elbahesh,H. and Zaraket,H. |
| EPI_ISL_2582157, EPI_ISL_2582160, EPI_ISL_2582161 | Epidemiology and Demography Department, KEMRI-Wellcome Trust Research Programme | Epidemiology and Demography Department, KEMRI-Wellcome Trust Research Programme | Otieno,J.R., Kamau,E.M., Oketch,J.W., Ngoi,J.M., Agoti,C.N., Gichuki,A.M., Otieno,G.P., Ngama,M., Cane,P.A., Kellam,P., Cotten,M., Lemey,P. and Nokes,D.J. |
| EPI_ISL_2582166 | Virology, Graduate School of Medicine, Tohoku University | Virology, Graduate School of Medicine, Tohoku University | Malasao,R., Furuse,Y., Okamoto,M., Dapat,C., Saito,M., Saito-Obata,M., Tamaki,R., Segubre-Mercado,E., Lupisan,S. and Oshitani,H. |
| EPI_ISL_2582167 | Lab Medicine, UW | Lab Medicine, UW | Greninger,A.L., Makhsous,N., Kuypers,J.M., Shean,R.C. and Jerome,K.R. |
| EPI_ISL_2582169 | Pediatrics - Infectious Diseases, Medical College of Wisconsin | Pediatrics - Infectious Diseases, Medical College of Wisconsin | Rebuffo-Scheer,C., Bose,M.E., He,J., Khaja,S., Ulatowski,M., Beck,E.T., Fan,J., Kumar,S., Nelson,M.I. and Henrickson,K.J. |
| EPI_ISL_2582173 | Virology, Graduate School of Medicine, Tohoku University | Virology, Graduate School of Medicine, Tohoku University | Malasao,R., Furuse,Y., Okamoto,M., Dapat,C., Saito,M., Saito-Obata,M., Tamaki,R., Segubre-Mercado,E., Lupisan,S. and Oshitani,H. |
| EPI_ISL_2582176 | Laboratory Medicine, UW Virology | Laboratory Medicine, UW Virology | Lin,M.J., Tait,A. and Greninger,A.L. |
| EPI_ISL_2582177 | Pediatrics, University of New Mexico | Pediatrics, University of New Mexico | Kothari,A., Kennedy,J.L., Schwalm,K.C., Putt,C., Denson,J.L. and Dinwiddie,D.L. |
| EPI_ISL_2582180 | Broad Institute of MIT & Harvard | Broad Institute of MIT & Harvard | Newman,R.M., Zody,M.C., DeVincenzo,J.P., Grad,Y., Lipsitch,M., Murphy,R., Fitzgerald,M., Young,S., Gargeya,S., Poon,T.W., Charlebois,P., Weiner,B., Yang,X., Piper,M.E., McCowan,C., Ireland,A., Levin,J., Malboeuf,C., Qu,J., Chapman,S.B., Murphy,C., Wortman,J., Nusbaum,C. and Birren,B. |
| EPI_ISL_2582181, EPI_ISL_2582183 | Virology, Graduate School of Medicine, Tohoku University | Virology, Graduate School of Medicine, Tohoku University | Malasao,R., Furuse,Y., Okamoto,M., Dapat,C., Saito,M., Saito-Obata,M., Tamaki,R., Segubre-Mercado,E., Lupisan,S. and Oshitani,H. |
| EPI_ISL_2582187, EPI_ISL_2582196 | J. Craig Venter Institute | J. Craig Venter Institute | Tan,G., Pickett,B., Fedorova,N., Amedeo,P., Isom,R., Hu,L., Christensen,J., Miller,J., Novotny,M., Durbin,A., Rocchi,I., Williams,T., Arumemi,F. and Das,S. |
| EPI_ISL_2582197 | Lab Medicine, UW | Lab Medicine, UW | Greninger,A.L., Makhsous,N., Kuypers,J.M., Shean,R.C. and Jerome,K.R. |
| EPI_ISL_2582201 | Epidemiology and Demography Department, KEMRI-Wellcome Trust Research Programme | Epidemiology and Demography Department, KEMRI-Wellcome Trust Research Programme | Otieno,J.R., Kamau,E.M., Oketch,J.W., Ngoi,J.M., Agoti,C.N., Gichuki,A.M., Otieno,G.P., Ngama,M., Cane,P.A., Kellam,P., Cotten,M., Lemey,P. and Nokes,D.J. |
| EPI_ISL_2582221 | J. Craig Venter Institute | J. Craig Venter Institute | Tan,G., Pickett,B., Fedorova,N., Amedeo,P., Hu,L., Christensen,J., Miller,J., Durbin,A., Williams,T., Arumemi,F., Cadiz,C., Alanis,R., Balmseda,A., Wiliams,T., Schiller,A., Patel,M., Kubale,J. and Gordon,A. |
| EPI_ISL_2582223, EPI_ISL_2582226, EPI_ISL_2582227 | Marie Bashir Institute for Infectious Diseases and Biosecurity & Sydney Medical School, The University of Sydney, Westmead Institute for Medical Research | Marie Bashir Institute for Infectious Diseases and Biosecurity & Sydney Medical School, The University of Sydney, Westmead Institute for Medical Research | Eden,J.-S., Kok,J., Dwyer,D.E., Fernandez,M., Carter,I. and Holmes,E.C. |
| EPI_ISL_2582230, EPI_ISL_2582231 | J. Craig Venter Institute | J. Craig Venter Institute | Tan,G., Pickett,B., Fedorova,N., Amedeo,P., Hu,L., Christensen,J., Miller,J., Durbin,A., Williams,T., Arumemi,F., Cadiz,C., Alanis,R., Balmseda,A., Wiliams,T., Schiller,A., Patel,M., Kubale,J. and Gordon,A. |
| EPI_ISL_2582233 | Laboratory Medicine, UW Virology | Laboratory Medicine, UW Virology | Lin,M.J., Tait,A. and Greninger,A.L. |
| EPI_ISL_2582235, EPI_ISL_2582238 | J. Craig Venter Institute | J. Craig Venter Institute | Tan,G., Pickett,B., Fedorova,N., Amedeo,P., Hu,L., Christensen,J., Miller,J., Durbin,A., Williams,T., Arumemi,F., Cadiz,C., Alanis,R., Balmseda,A., Wiliams,T., Schiller,A., Patel,M., Kubale,J. and Gordon,A. |
| EPI_ISL_2582239, EPI_ISL_2582241, EPI_ISL_2582244 | Marie Bashir Institute for Infectious Diseases and Biosecurity & Sydney Medical School, The | Marie Bashir Institute for Infectious Diseases and Biosecurity & Sydney Medical School, The | Eden,J.-S., Kok,J., Dwyer,D.E., Fernandez,M., Carter,I. and Holmes,E.C. |

| | | | |
|---|---|---|---|
| | University of Sydney, Westmead Institute for Medical Research | University of Sydney, Westmead Institute for Medical Research | |
| EPI_ISL_2582246 | Pediatrics, University of New Mexico | Pediatrics, University of New Mexico | Kothari,A., Kennedy,J.L., Schwalm,K.C., Putt,C., Denson,J.L. and Dinwiddie,D.L. |
| EPI_ISL_2582247 | Laboratory Medicine, UW Virology | Laboratory Medicine, UW Virology | Lin,M.J., Tait,A. and Greninger,A.L. |
| EPI_ISL_2582250, EPI_ISL_2582251, EPI_ISL_2582253 | Marie Bashir Institute for Infectious Diseases and Biosecurity & Sydney Medical School, The University of Sydney, Westmead Institute for Medical Research | Marie Bashir Institute for Infectious Diseases and Biosecurity & Sydney Medical School, The University of Sydney, Westmead Institute for Medical Research | Eden,J.-S., Kok,J., Dwyer,D.E., Fernandez,M., Carter,I. and Holmes,E.C. |
| EPI_ISL_2582256, EPI_ISL_2582257, EPI_ISL_2582260 | Epidemiology and Demography Department, KEMRI-Wellcome Trust Research Programme | Epidemiology and Demography Department, KEMRI-Wellcome Trust Research Programme | Otieno,J.R., Kamau,E.M., Oketch,J.W., Ngoi,J.M., Agoti,C.N., Gichuki,A.M., Otieno,G.P., Ngama,M., Cane,P.A., Kellam,P., Cotten,M., Lemey,P. and Nokes,D.J. |
| EPI_ISL_2582261 | Pediatrics, University of New Mexico | Pediatrics, University of New Mexico | Kothari,A., Kennedy,J.L., Schwalm,K.C., Putt,C., Denson,J.L. and Dinwiddie,D.L. |
| EPI_ISL_2582264, EPI_ISL_2582265 | Marie Bashir Institute for Infectious Diseases and Biosecurity & Sydney Medical School, The University of Sydney, Westmead Institute for Medical Research | Marie Bashir Institute for Infectious Diseases and Biosecurity & Sydney Medical School, The University of Sydney, Westmead Institute for Medical Research | Eden,J.-S., Kok,J., Dwyer,D.E., Fernandez,M., Carter,I. and Holmes,E.C. |
| EPI_ISL_2582267 | Epidemiology and Demography Department, KEMRI-Wellcome Trust Research Programme | Epidemiology and Demography Department, KEMRI-Wellcome Trust Research Programme | Otieno,J.R., Kamau,E.M., Oketch,J.W., Ngoi,J.M., Agoti,C.N., Gichuki,A.M., Otieno,G.P., Ngama,M., Cane,P.A., Kellam,P., Cotten,M., Lemey,P. and Nokes,D.J. |
| EPI_ISL_2582271 | J. Craig Venter Institute | J. Craig Venter Institute | Tan,G., Pickett,B., Fedorova,N., Amedeo,P., Hu,L., Christensen,J., Miller,J., Durbin,A., Williams,T., Arumemi,F., Cadiz,C., Alanis,R., Balmseda,A., Wiliams,T., Schiller,A., Patel,M., Kubale,J. and Gordon,A. |
| EPI_ISL_2582274, EPI_ISL_2582275, EPI_ISL_2582278 | Department of Experimental Modeling and Pathogenesis of Infectious Diseases, Federal Research Center of Fundamental and Translational Medicine | Department of Experimental Modeling and Pathogenesis of Infectious Diseases, Federal Research Center of Fundamental and Translational Medicine | Dubovitskiy,N.A., Sobolev,I.A., Kurskaya,O.G., Sharshov,K.A., Anoshina,A.V., Leonova,N.V., Murashkina,T.A., Solomatina,M.V., Derko,A.A., Saroyan,T.A., Kabilov,M.R., Alikina,T.Y. and Shestopalov,A.M. |
| EPI_ISL_2582279, EPI_ISL_2582281, EPI_ISL_2582284 | Epidemiology and Demography Department, KEMRI-Wellcome Trust Research Programme | Epidemiology and Demography Department, KEMRI-Wellcome Trust Research Programme | Otieno,J.R., Kamau,E.M., Oketch,J.W., Ngoi,J.M., Agoti,C.N., Gichuki,A.M., Otieno,G.P., Ngama,M., Cane,P.A., Kellam,P., Cotten,M., Lemey,P. and Nokes,D.J. |
| EPI_ISL_2582285 | Department of Pediatrics, Center of Excellence in Clinical Virology, Chulalongkorn | Department of Pediatrics, Center of Excellence in Clinical Virology, Chulalongkorn | Thongpan,I. |
| EPI_ISL_2582292 | Epidemiology and Demography Department, KEMRI-Wellcome Trust Research Programme | Epidemiology and Demography Department, KEMRI-Wellcome Trust Research Programme | Otieno,J.R., Kamau,E.M., Oketch,J.W., Ngoi,J.M., Agoti,C.N., Gichuki,A.M., Otieno,G.P., Ngama,M., Cane,P.A., Kellam,P., Cotten,M., Lemey,P. and Nokes,D.J. |
| EPI_ISL_2582296 | Department of Pediatrics, Center of Excellence in Clinical Virology, Chulalongkorn | Department of Pediatrics, Center of Excellence in Clinical Virology, Chulalongkorn | Thongpan,I. |
| EPI_ISL_2582298, EPI_ISL_2582299, EPI_ISL_2582302, EPI_ISL_2582303 | J. Craig Venter Institute | J. Craig Venter Institute | Shabman,R., Fedorova,N., Puri,V., Shrivastava,S., Amedeo,P., Isom,R., Hu,L., Pickett,B., Novotny,M., Durbin,A., Rocchi,I., Williams,T., Hall,C.B., Tesini,B.L., Schnabel,K.C., Walsh,E.E. and Caserta,M. |
| EPI_ISL_2582306, EPI_ISL_2582307, EPI_ISL_2582309, EPI_ISL_2582312, EPI_ISL_2582313, EPI_ISL_2582316, EPI_ISL_2582318, EPI_ISL_2582319, EPI_ISL_2582321, EPI_ISL_2582324, EPI_ISL_2582325, EPI_ISL_2582327, EPI_ISL_2582330, EPI_ISL_2582332, EPI_ISL_2582333, EPI_ISL_2582335, EPI_ISL_2582337, EPI_ISL_2582340, EPI_ISL_2582341, EPI_ISL_2582342, EPI_ISL_2582344, EPI_ISL_2582345, EPI_ISL_2582348, EPI_ISL_2582349, EPI_ISL_2582351, EPI_ISL_2582354, EPI_ISL_2582355, EPI_ISL_2582357, EPI_ISL_2582360, EPI_ISL_2582361, EPI_ISL_2582363, EPI_ISL_2582366, EPI_ISL_2582367, EPI_ISL_2582370, EPI_ISL_2582371, EPI_ISL_2582373, EPI_ISL_2582375, EPI_ISL_2582378, EPI_ISL_2582379, EPI_ISL_2582381, EPI_ISL_2582383, EPI_ISL_2582386, EPI_ISL_2582387, EPI_ISL_2582389, EPI_ISL_2582391, EPI_ISL_2582393, EPI_ISL_2582395, EPI_ISL_2582397, EPI_ISL_2582399, EPI_ISL_2582402, EPI_ISL_2582403, EPI_ISL_2582405, EPI_ISL_2582407, EPI_ISL_2582410, EPI_ISL_2582411, EPI_ISL_2582413, EPI_ISL_2582416, EPI_ISL_2582417, EPI_ISL_2582419, EPI_ISL_2582422, EPI_ISL_2582423, EPI_ISL_2582426, EPI_ISL_2582427, EPI_ISL_2582430, EPI_ISL_2582431, EPI_ISL_2582434, EPI_ISL_2582435, EPI_ISL_2582438, EPI_ISL_2582439, EPI_ISL_2582442, EPI_ISL_2582443, EPI_ISL_2582445, EPI_ISL_2582448, EPI_ISL_2582449, EPI_ISL_2582451, EPI_ISL_2582456, EPI_ISL_2582457, EPI_ISL_2582460, EPI_ISL_2582461, EPI_ISL_2582463, EPI_ISL_2582465, EPI_ISL_2582468, EPI_ISL_2582469, EPI_ISL_2582471, EPI_ISL_2582476, EPI_ISL_2582477, EPI_ISL_2582479, EPI_ISL_2582481, EPI_ISL_2582484, EPI_ISL_2582485, EPI_ISL_2582487, EPI_ISL_2582490, EPI_ISL_2582491, EPI_ISL_2582495, EPI_ISL_2582497, EPI_ISL_2582500, EPI_ISL_2582502, EPI_ISL_2582503, EPI_ISL_2582505, EPI_ISL_2582508, EPI_ISL_2582509 | | | |
| see above | Broad Institute of MIT & Harvard | Broad Institute of MIT & Harvard | Newman,R.M., Zody,M.C., DeVincenzo,J.P., Grad,Y., Lipsitch,M., Murphy,R., Fitzgerald,M., Young,S., Gargeya,S., Poon,T.W., Charlebois,P., Weiner,B., Yang,X., Piper,M.E., McCowan,C., Ireland,A., Levin,J., Malboeuf,C., Qu,J., Chapman,S.B., Murphy,C., Wortman,J., Nusbaum,C. and Birren,B. |
| EPI_ISL_2582510, EPI_ISL_2582512 | Marie Bashir Institute for Infectious Diseases and Biosecurity & Sydney Medical School, The University of Sydney, Westmead Institute for Medical Research | Marie Bashir Institute for Infectious Diseases and Biosecurity & Sydney Medical School, The University of Sydney, Westmead Institute for Medical Research | Eden,J.-S., Kok,J., Dwyer,D.E., Fernandez,M., Carter,I. and Holmes,E.C. |
| EPI_ISL_2582513, EPI_ISL_2582515, EPI_ISL_2582517, EPI_ISL_2582519 | Broad Institute of MIT & Harvard | Broad Institute of MIT & Harvard | Newman,R.M., Zody,M.C., DeVincenzo,J.P., Grad,Y., Lipsitch,M., Murphy,R., Fitzgerald,M., Young,S., Gargeya,S., Poon,T.W., Charlebois,P., Weiner,B., Yang,X., Piper,M.E., McCowan,C., Ireland,A., Levin,J., Malboeuf,C., Qu,J., Chapman,S.B., Murphy,C., Wortman,J., Nusbaum,C. and Birren,B. |
| EPI_ISL_2582521 | J. Craig Venter Institute | J. Craig Venter Institute | Shabman,R., Fedorova,N., Puri,V., Shrivastava,S., Amedeo,P., Isom,R., Hu,L., Pickett,B., Novotny,M., Durbin,A., Rocchi,I., Williams,T., Hall,C.B., Tesini,B.L., Schnabel,K.C., Walsh,E.E. and Caserta,M. |
| EPI_ISL_2582536 | Laboratory Medicine, UW Virology | Laboratory Medicine, UW Virology | Lin,M.J., Tait,A. and Greninger,A.L. |
| EPI_ISL_2582538, EPI_ISL_2582540, EPI_ISL_2582541 | Lab Medicine, UW | Lab Medicine, UW | Greninger,A.L., Makhsous,N., Kuypers,J.M., Shean,R.C. and Jerome,K.R. |
| EPI_ISL_2582543, EPI_ISL_2582545, EPI_ISL_2582547 | J. Craig Venter Institute | J. Craig Venter Institute | Tan,G., Pickett,B., Fedorova,N., Amedeo,P., Isom,R., Hu,L., Christensen,J., Miller,J., Novotny,M., Durbin,A., Rocchi,I., Williams,T., Arumemi,F. and Das,S. |
| EPI_ISL_2582551 | Lab Medicine, UW | Lab Medicine, UW | Greninger,A.L., Makhsous,N., Kuypers,J.M., Shean,R.C. and Jerome,K.R. |
| EPI_ISL_2582553 | J. Craig Venter Institute | J. Craig Venter Institute | Tan,G., Pickett,B., Fedorova,N., Amedeo,P., Isom,R., Hu,L., Christensen,J., Miller,J., Novotny,M., Durbin,A., Rocchi,I., Williams,T., Arumemi,F. and Das,S. |
| EPI_ISL_2582561 | Central Laboratory, Guangzhou Women and Children's Medical Center | Central Laboratory, Guangzhou Women and Children's Medical Center | Xie,J.H., Zhu,B., Zhong,J.Y., Chen,Y. and Zhang,Y.Y. |
| EPI_ISL_2582563 | The Second Department, Lanzhou Institute of Biological Products Co | The Second Department, Lanzhou Institute of Biological Products Co | Zhu,C., Fu,S., Yu,L. and Zhou,X. |
| EPI_ISL_2582565, EPI_ISL_2582567 | Epidemiology and Demography Department, KEMRI-Wellcome Trust Research Programme | Epidemiology and Demography Department, KEMRI-Wellcome Trust Research Programme | Otieno,J.R., Kamau,E.M., Oketch,J.W., Ngoi,J.M., Agoti,C.N., Gichuki,A.M., Otieno,G.P., Ngama,M., Cane,P.A., Kellam,P., Cotten,M., Lemey,P. and Nokes,D.J. |
| EPI_ISL_2582569, EPI_ISL_2582571 | J. Craig Venter Institute | J. Craig Venter Institute | Tan,G., Pickett,B., Fedorova,N., Amedeo,P., Isom,R., Hu,L., Christensen,J., Miller,J., Novotny,M., Durbin,A., Rocchi,I., Williams,T., Arumemi,F. and Das,S. |
| EPI_ISL_2582573 | Pediatrics - Infectious Diseases, Medical College of Wisconsin | Pediatrics - Infectious Diseases, Medical College of Wisconsin | Rebuffo-Scheer,C., Bose,M.E., He,J., Khaja,S., Ulatowski,M., Beck,E.T., Fan,J., Kumar,S., Nelson,M.I. and Henrickson,K.J. |
| EPI_ISL_2582577, EPI_ISL_2582579, EPI_ISL_2582581, EPI_ISL_2582583, EPI_ISL_2582585, EPI_ISL_2582587, EPI_ISL_2582588, EPI_ISL_2582590, EPI_ISL_2582592, EPI_ISL_2582594, EPI_ISL_2582596, EPI_ISL_2582598, EPI_ISL_2582600, EPI_ISL_2582601, EPI_ISL_2582603, EPI_ISL_2582605, EPI_ISL_2582607, EPI_ISL_2582609, EPI_ISL_2582610, EPI_ISL_2582612, EPI_ISL_2582614, EPI_ISL_2582616, EPI_ISL_2582618 | | | |
| see above | Broad Institute of MIT & Harvard | Broad Institute of MIT & Harvard | Newman,R.M., Zody,M.C., DeVincenzo,J.P., Grad,Y., Lipsitch,M., Murphy,R., Fitzgerald,M., Young,S., Gargeya,S., Poon,T.W., Charlebois,P., Weiner,B., Yang,X., Piper,M.E., McCowan,C., Ireland,A., Levin,J., Malboeuf,C., Qu,J., Chapman,S.B., Murphy,C., Wortman,J., Nusbaum,C. and Birren,B. |
| EPI_ISL_2582621 | Academy of Military Medical Sciences, Institute of Microbiology and Epidemiology | Academy of Military Medical Sciences, Institute of Microbiology and Epidemiology | Gu,H.J., Sun,S.J., Chen,R. and Yang,P.H. |
| EPI_ISL_2582623, EPI_ISL_2582625 | Broad Institute of MIT & Harvard | Broad Institute of MIT & Harvard | Newman,R.M., Zody,M.C., DeVincenzo,J.P., Grad,Y., Lipsitch,M., Murphy,R., Fitzgerald,M., Young,S., Gargeya,S., Poon,T.W., Charlebois,P., Weiner,B., Yang,X., Piper,M.E., McCowan,C., Ireland,A., Levin,J., Malboeuf,C., Qu,J., Chapman,S.B., Murphy,C., Wortman,J., Nusbaum,C. and Birren,B. |
| EPI_ISL_2582630 | Pediatrics - Infectious Diseases, Medical College of Wisconsin | Pediatrics - Infectious Diseases, Medical College of Wisconsin | Rebuffo-Scheer,C., Bose,M.E., He,J., Khaja,S., Ulatowski,M., Beck,E.T., Fan,J., Kumar,S., Nelson,M.I. and Henrickson,K.J. |
| EPI_ISL_2582631, EPI_ISL_2582633 | J. Craig Venter Institute | J. Craig Venter Institute | Shabman,R., Fedorova,N., Puri,V., Shrivastava,S., Amedeo,P., Isom,R., Hu,L., Pickett,B., Novotny,M., Durbin,A., Rocchi,I., Williams,T., Hall,C.B., Tesini,B.L., Schnabel,K.C., Walsh,E.E. and Caserta,M. |
| EPI_ISL_2582635 | Broad Institute of MIT & Harvard | Broad Institute of MIT & Harvard | Newman,R.M., Zody,M.C., DeVincenzo,J.P., Grad,Y., Lipsitch,M., Murphy,R., Fitzgerald,M., Young,S., Gargeya,S., Poon,T.W., Charlebois,P., Weiner,B., Yang,X., Piper,M.E., McCowan,C., Ireland,A., Levin,J., Malboeuf,C., Qu,J., Chapman,S.B., Murphy,C., Wortman,J., Nusbaum,C. and Birren,B. |
| EPI_ISL_2582641, EPI_ISL_2582644, EPI_ISL_2582646 | Marie Bashir Institute for Infectious Diseases and Biosecurity & Sydney Medical School, The University of Sydney, Westmead Institute for Medical Research | Marie Bashir Institute for Infectious Diseases and Biosecurity & Sydney Medical School, The University of Sydney, Westmead Institute for Medical Research | Eden,J.-S., Kok,J., Dwyer,D.E., Fernandez,M., Carter,I. and Holmes,E.C. |
| EPI_ISL_2582648, EPI_ISL_2582650, EPI_ISL_2582653 | Pediatrics, University of New Mexico | Pediatrics, University of New Mexico | Kothari,A., Kennedy,J.L., Schwalm,K.C., Putt,C., Denson,J.L. and Dinwiddie,D.L. |
| EPI_ISL_2582669, EPI_ISL_2582670, EPI_ISL_2582673 | Epidemiology and Demography Department, KEMRI-Wellcome Trust Research Programme | Epidemiology and Demography Department, KEMRI-Wellcome Trust Research Programme | Otieno,J.R., Kamau,E.M., Oketch,J.W., Ngoi,J.M., Agoti,C.N., Gichuki,A.M., Otieno,G.P., Ngama,M., Cane,P.A., Kellam,P., Cotten,M., Lemey,P. and Nokes,D.J. |
| EPI_ISL_2582675 | Suguru Takeuchi Nagoya University Graduate School of Medicine, Department of Pediatrics | Suguru Takeuchi Nagoya University Graduate School of Medicine, Department of Pediatrics | Takeuchi,S., Kawada,J. and Ito,Y. |
| EPI_ISL_2582678 | Epidemiology and Demography Department, KEMRI-Wellcome Trust Research Programme | Epidemiology and Demography Department, KEMRI-Wellcome Trust Research Programme | Otieno,J.R., Kamau,E.M., Oketch,J.W., Ngoi,J.M., Agoti,C.N., Gichuki,A.M., Otieno,G.P., Ngama,M., Cane,P.A., Kellam,P., Cotten,M., Lemey,P. and Nokes,D.J. |
| EPI_ISL_2582681 | Experimental Pathology, Immunology, and Microbiology, American University of Beirut | Experimental Pathology, Immunology, and Microbiology, American University of Beirut | Ezzeddine,A.M. |
| EPI_ISL_2582683, EPI_ISL_2582686, EPI_ISL_2582688, EPI_ISL_2582691, EPI_ISL_2582694, EPI_ISL_2582696, EPI_ISL_2582699, EPI_ISL_2582701 | Broad Institute of MIT & Harvard | Broad Institute of MIT & Harvard | Newman,R.M., Zody,M.C., DeVincenzo,J.P., Grad,Y., Lipsitch,M., Murphy,R., Fitzgerald,M., Young,S., Gargeya,S., Poon,T.W., Charlebois,P., Weiner,B., Yang,X., Piper,M.E., McCowan,C., Ireland,A., Levin,J., Malboeuf,C., Qu,J., Chapman,S.B., Murphy,C., Wortman,J., Nusbaum,C. and Birren,B. |
| EPI_ISL_2582703 | J. Craig Venter Institute | J. Craig Venter Institute | Shabman,R., Fedorova,N., Puri,V., Shrivastava,S., Amedeo,P., Isom,R., Hu,L., Pickett,B., Novotny,M., Durbin,A., Rocchi,I., Williams,T., Hall,C.B., Tesini,B.L., Schnabel,K.C., Walsh,E.E. and Caserta,M. |

| | | | |
|---|---|---|---|
| EPI_ISL_2582706, EPI_ISL_2582709, EPI_ISL_2582711, EPI_ISL_2582714, EPI_ISL_2582716, EPI_ISL_2582719, EPI_ISL_2582722, EPI_ISL_2582724, EPI_ISL_2582727, EPI_ISL_2582729, EPI_ISL_2582732, EPI_ISL_2582735, EPI_ISL_2582738, EPI_ISL_2582740, EPI_ISL_2582743, EPI_ISL_2582745, EPI_ISL_2582747, EPI_ISL_2582750, EPI_ISL_2582751, EPI_ISL_2582756, EPI_ISL_2582758, EPI_ISL_2582760, EPI_ISL_2582762, EPI_ISL_2582764 | | | |
| see above | Broad Institute of MIT & Harvard | Broad Institute of MIT & Harvard | Newman,R.M., Zody,M.C., DeVincenzo,J.P., Grad,Y., Lipsitch,M., Murphy,R., Fitzgerald,M., Young,S., Gargeya,S., Poon,T.W., Charlebois,P., Weiner,B., Yang,X., Piper,M.E., McCowan,C., Ireland,A., Levin,J., Malboeuf,C., Qu,J., Chapman,S.B., Murphy,C., Wortman,J., Nusbaum,C. and Birren,B. |
| EPI_ISL_2582766 | Infectious Disease Initiative, Broad Institute | Infectious Disease Initiative, Broad Institute | Newman,R.M., Zody,M.C., DeVincenzo,J.P., Grad,Y., Lipsitch,M., Murphy,R., Fitzgerald,M., Young,S., Gargeya,S., Poon,T.W., Charlebois,P., Weiner,B., Yang,X., Piper,M.E., McCowan,C., Ireland,A., Levin,J., Malboeuf,C., Qu,J., Chapman,S.B., Murphy,C., Wortman,J., Nusbaum,C. and Birren,B. |
| EPI_ISL_2582768 | Broad Institute of MIT & Harvard | Broad Institute of MIT & Harvard | Newman,R.M., Zody,M.C., DeVincenzo,J.P., Grad,Y., Lipsitch,M., Murphy,R., Fitzgerald,M., Young,S., Gargeya,S., Poon,T.W., Charlebois,P., Weiner,B., Yang,X., Piper,M.E., McCowan,C., Ireland,A., Levin,J., Malboeuf,C., Qu,J., Chapman,S.B., Murphy,C., Wortman,J., Nusbaum,C. and Birren,B. |
| EPI_ISL_2582771 | Marie Bashir Institute for Infectious Diseases and Biosecurity & Sydney Medical School, The University of Sydney, Westmead Institute for Medical Research | Marie Bashir Institute for Infectious Diseases and Biosecurity & Sydney Medical School, The University of Sydney, Westmead Institute for Medical Research | Eden,J.-S., Kok,J., Dwyer,D.E., Fernandez,M., Carter,I. and Holmes,E.C. |
| EPI_ISL_2582775, EPI_ISL_2582778 | J. Craig Venter Institute | J. Craig Venter Institute | Shabman,R., Fedorova,N., Puri,V., Shrivastava,S., Amedeo,P., Isom,R., Hu,L., Pickett,B., Novotny,M., Durbin,A., Rocchi,I., Williams,T., Hall,C.B., Tesini,B.L., Schnabel,K.C., Walsh,E.E. and Caserta,M. |
| EPI_ISL_2582780 | Broad Institute of MIT & Harvard | Broad Institute of MIT & Harvard | Newman,R.M., Zody,M.C., DeVincenzo,J.P., Grad,Y., Lipsitch,M., Murphy,R., Fitzgerald,M., Young,S., Gargeya,S., Poon,T.W., Charlebois,P., Weiner,B., Yang,X., Piper,M.E., McCowan,C., Ireland,A., Levin,J., Malboeuf,C., Qu,J., Chapman,S.B., Murphy,C., Wortman,J., Nusbaum,C. and Birren,B. |
| EPI_ISL_2582782 | J. Craig Venter Institute | J. Craig Venter Institute | Shabman,R., Fedorova,N., Puri,V., Shrivastava,S., Amedeo,P., Isom,R., Hu,L., Pickett,B., Novotny,M., Durbin,A., Rocchi,I., Williams,T., Hall,C.B., Tesini,B.L., Schnabel,K.C., Walsh,E.E. and Caserta,M. |
| EPI_ISL_2582784 | Broad Institute of MIT & Harvard | Broad Institute of MIT & Harvard | Newman,R.M., Zody,M.C., DeVincenzo,J.P., Grad,Y., Lipsitch,M., Murphy,R., Fitzgerald,M., Young,S., Gargeya,S., Poon,T.W., Charlebois,P., Weiner,B., Yang,X., Piper,M.E., McCowan,C., Ireland,A., Levin,J., Malboeuf,C., Qu,J., Chapman,S.B., Murphy,C., Wortman,J., Nusbaum,C. and Birren,B. |
| EPI_ISL_2582787 | Lab Medicine, UW | Lab Medicine, UW | Greninger,A.L., Makhsous,N., Kuypers,J.M., Shean,R.C. and Jerome,K.R. |
| EPI_ISL_2582789 | Epidemiology and Demography Department, KEMRI-Wellcome Trust Research Programme | Epidemiology and Demography Department, KEMRI-Wellcome Trust Research Programme | Otieno,J.R., Kamau,E.M., Oketch,J.W., Ngoi,J.M., Agoti,C.N., Gichuki,A.M., Otieno,G.P., Ngama,M., Cane,P.A., Kellam,P., Cotten,M., Lemey,P. and Nokes,D.J. |
| EPI_ISL_2582796 | Central Laboratory, Guangzhou Women and Children's Medical Center | Central Laboratory, Guangzhou Women and Children's Medical Center | Xie,J.H., Zhu,B., Zhong,J.Y., Chen,Y. and Zhang,Y.Y. |
| EPI_ISL_2582798 | Lab Medicine, UW | Lab Medicine, UW | Greninger,A.L., Makhsous,N., Kuypers,J.M., Shean,R.C. and Jerome,K.R. |
| EPI_ISL_2582801 | Department of Laboratory Medicine, Lin-Kou Chang-Gung Memorial Hospital | Department of Laboratory Medicine, Lin-Kou Chang-Gung Memorial Hospital | Tsao,K.-C., Gong,Y.-N., Yang,S.-L., Chen,G.-W., Chen,Y.-W., Huang,Y.-C. and Liu,Y.-C. |
| EPI_ISL_2582804 | Epidemiology and Demography Department, KEMRI-Wellcome Trust Research Programme | Epidemiology and Demography Department, KEMRI-Wellcome Trust Research Programme | Otieno,J.R., Kamau,E.M., Oketch,J.W., Ngoi,J.M., Agoti,C.N., Gichuki,A.M., Otieno,G.P., Ngama,M., Cane,P.A., Kellam,P., Cotten,M., Lemey,P. and Nokes,D.J. |
| EPI_ISL_2582805 | J. Craig Venter Institute | J. Craig Venter Institute | Tan,G., Pickett,B., Fedorova,N., Amedeo,P., Isom,R., Hu,L., Christensen,J., Miller,J., Novotny,M., Durbin,A., Rocchi,I., Williams,T., Arumemi,F. and Das,S. |
| EPI_ISL_2582808, EPI_ISL_2582810, EPI_ISL_2582813, EPI_ISL_2582819, EPI_ISL_2582820, EPI_ISL_2582823, EPI_ISL_2582826 | Broad Institute of MIT & Harvard | Broad Institute of MIT & Harvard | Newman,R.M., Zody,M.C., DeVincenzo,J.P., Grad,Y., Lipsitch,M., Murphy,R., Fitzgerald,M., Young,S., Gargeya,S., Poon,T.W., Charlebois,P., Weiner,B., Yang,X., Piper,M.E., McCowan,C., Ireland,A., Levin,J., Malboeuf,C., Qu,J., Chapman,S.B., Murphy,C., Wortman,J., Nusbaum,C. and Birren,B. |
| EPI_ISL_2582832, EPI_ISL_2582834 | Laboratory Medicine, UW Virology | Laboratory Medicine, UW Virology | Lin,M.J., Tait,A. and Greninger,A.L. |
| EPI_ISL_2582836, EPI_ISL_2582838 | Department of Experimental Modeling and Pathogenesis of Infectious Diseases, Federal Research Center of Fundamental and Translational Medicine | Department of Experimental Modeling and Pathogenesis of Infectious Diseases, Federal Research Center of Fundamental and Translational Medicine | Dubovitskiy,N.A., Sobolev,I.A., Kurskaya,O.G., Sharshov,K.A., Anoshina,A.V., Leonova,N.V., Murashkina,T.A., Solomatina,M.V., Derko,A.A., Saroyan,T.A., Kabilov,M.R., Alikina,T.Y. and Shestopalov,A.M. |
| EPI_ISL_2582841 | Marie Bashir Institute for Infectious Diseases and Biosecurity & Sydney Medical School, The University of Sydney, Westmead Institute for Medical Research | Marie Bashir Institute for Infectious Diseases and Biosecurity & Sydney Medical School, The University of Sydney, Westmead Institute for Medical Research | Eden,J.-S., Kok,J., Dwyer,D.E., Fernandez,M., Carter,I. and Holmes,E.C. |
| EPI_ISL_2582844 | Broad Institute of MIT & Harvard | Broad Institute of MIT & Harvard | Newman,R.M., Zody,M.C., DeVincenzo,J.P., Grad,Y., Lipsitch,M., Murphy,R., Fitzgerald,M., Young,S., Gargeya,S., Poon,T.W., Charlebois,P., Weiner,B., Yang,X., Piper,M.E., McCowan,C., Ireland,A., Levin,J., Malboeuf,C., Qu,J., Chapman,S.B., Murphy,C., Wortman,J., Nusbaum,C. and Birren,B. |
| EPI_ISL_2582847 | J. Craig Venter Institute | J. Craig Venter Institute | Shabman,R., Fedorova,N., Puri,V., Shrivastava,S., Amedeo,P., Isom,R., Hu,L., Pickett,B., Novotny,M., Durbin,A., Rocchi,I., Williams,T., Hall,C.B., Tesini,B.L., Schnabel,K.C., Walsh,E.E. and Caserta,M. |
| EPI_ISL_2582849, EPI_ISL_2582851 | Center for Infectious Diseases, School of Public Health, University of Texas Health Science Center | Center for Infectious Diseases, School of Public Health, University of Texas Health Science Center | Bahl,J., Hixson,J., Kim,D.-K., Qiu,X., Piedra,P.A., Piedra,F.-A., Avadhanula,V. and Machado,A.A. |
| EPI_ISL_2582854 | Broad Institute of MIT & Harvard | Broad Institute of MIT & Harvard | Newman,R.M., Zody,M.C., DeVincenzo,J.P., Grad,Y., Lipsitch,M., Murphy,R., Fitzgerald,M., Young,S., Gargeya,S., Poon,T.W., Charlebois,P., Weiner,B., Yang,X., Piper,M.E., McCowan,C., Ireland,A., Levin,J., Malboeuf,C., Qu,J., Chapman,S.B., Murphy,C., Wortman,J., Nusbaum,C. and Birren,B. |
| EPI_ISL_2582857, EPI_ISL_2582859, EPI_ISL_2582861 | Pediatrics - Infectious Diseases, Medical College of Wisconsin | Pediatrics - Infectious Diseases, Medical College of Wisconsin | Rebuffo-Scheer,C., Bose,M.E., He,J., Khaja,S., Ulatowski,M., Beck,E.T., Fan,J., Kumar,S., Nelson,M.I. and Henrickson,K.J. |
| EPI_ISL_2582864 | Gansu Center for Disease Control and Prevention, Pathogen Laboratory | Gansu Center for Disease Control and Prevention, Pathogen Laboratory | Qiao,R., Chen,J., Wu,H. and Yu,D. |
| EPI_ISL_2582866 | Epidemiology and Demography Department, KEMRI-Wellcome Trust Research Programme | Epidemiology and Demography Department, KEMRI-Wellcome Trust Research Programme | Otieno,J.R., Kamau,E.M., Oketch,J.W., Ngoi,J.M., Agoti,C.N., Gichuki,A.M., Otieno,G.P., Ngama,M., Cane,P.A., Kellam,P., Cotten,M., Lemey,P. and Nokes,D.J. |
| EPI_ISL_2582869 | Pediatrics - Infectious Diseases, Medical College of Wisconsin | Pediatrics - Infectious Diseases, Medical College of Wisconsin | Rebuffo-Scheer,C., Bose,M.E., He,J., Khaja,S., Ulatowski,M., Beck,E.T., Fan,J., Kumar,S., Nelson,M.I. and Henrickson,K.J. |
| EPI_ISL_2582873, EPI_ISL_2582875 | J. Craig Venter Institute | J. Craig Venter Institute | Shabman,R., Fedorova,N., Puri,V., Shrivastava,S., Amedeo,P., Isom,R., Hu,L., Pickett,B., Novotny,M., Durbin,A., Rocchi,I., Williams,T., Hall,C.B., Tesini,B.L., Schnabel,K.C., Walsh,E.E. and Caserta,M. |
| EPI_ISL_2582878 | Broad Institute of MIT & Harvard | Broad Institute of MIT & Harvard | Newman,R.M., Zody,M.C., DeVincenzo,J.P., Grad,Y., Lipsitch,M., Murphy,R., Fitzgerald,M., Young,S., Gargeya,S., Poon,T.W., Charlebois,P., Weiner,B., Yang,X., Piper,M.E., McCowan,C., Ireland,A., Levin,J., Malboeuf,C., Qu,J., Chapman,S.B., Murphy,C., Wortman,J., Nusbaum,C. and Birren,B. |
| EPI_ISL_2582880 | Medicine, University of Washington, 300 9th Ave, Harborview Research & Training Building | Medicine, University of Washington, 300 9th Ave, Harborview Research & Training Building | Chu,H., Scott,E. and Roychoudhury,P. |
| EPI_ISL_2582883 | Broad Institute of MIT & Harvard | Broad Institute of MIT & Harvard | Newman,R.M., Zody,M.C., DeVincenzo,J.P., Grad,Y., Lipsitch,M., Murphy,R., Fitzgerald,M., Young,S., Gargeya,S., Poon,T.W., Charlebois,P., Weiner,B., Yang,X., Piper,M.E., McCowan,C., Ireland,A., Levin,J., Malboeuf,C., Qu,J., Chapman,S.B., Murphy,C., Wortman,J., Nusbaum,C. and Birren,B. |
| EPI_ISL_2582886 | J. Craig Venter Institute | J. Craig Venter Institute | Shabman,R., Fedorova,N., Puri,V., Shrivastava,S., Amedeo,P., Isom,R., Hu,L., Pickett,B., Novotny,M., Durbin,A., Rocchi,I., Williams,T., Hall,C.B., Tesini,B.L., Schnabel,K.C., Walsh,E.E. and Caserta,M. |
| EPI_ISL_2582892, EPI_ISL_2582893 | Pediatrics - Infectious Diseases, Medical College of Wisconsin | Pediatrics - Infectious Diseases, Medical College of Wisconsin | Rebuffo-Scheer,C., Bose,M.E., He,J., Khaja,S., Ulatowski,M., Beck,E.T., Fan,J., Kumar,S., Nelson,M.I. and Henrickson,K.J. |
| EPI_ISL_2582901 | J. Craig Venter Institute | J. Craig Venter Institute | Shabman,R., Fedorova,N., Puri,V., Shrivastava,S., Amedeo,P., Isom,R., Hu,L., Pickett,B., Novotny,M., Durbin,A., Rocchi,I., Williams,T., Hall,C.B., Tesini,B.L., Schnabel,K.C., Walsh,E.E. and Caserta,M. |
| EPI_ISL_2582904 | Broad Institute of MIT & Harvard | Broad Institute of MIT & Harvard | Newman,R.M., Zody,M.C., DeVincenzo,J.P., Grad,Y., Lipsitch,M., Murphy,R., Fitzgerald,M., Young,S., Gargeya,S., Poon,T.W., Charlebois,P., Weiner,B., Yang,X., Piper,M.E., McCowan,C., Ireland,A., Levin,J., Malboeuf,C., Qu,J., Chapman,S.B., Murphy,C., Wortman,J., Nusbaum,C. and Birren,B. |
| EPI_ISL_2582906 | Department of Pediatrics, Center of Excellence in Clinical Virology, Chulalongkorn | Department of Pediatrics, Center of Excellence in Clinical Virology, Chulalongkorn | Thongpan,I. |
| EPI_ISL_2582908, EPI_ISL_2582910 | Epidemiology and Demography Department, KEMRI-Wellcome Trust Research Programme | Epidemiology and Demography Department, KEMRI-Wellcome Trust Research Programme | Otieno,J.R., Kamau,E.M., Oketch,J.W., Ngoi,J.M., Agoti,C.N., Gichuki,A.M., Otieno,G.P., Ngama,M., Cane,P.A., Kellam,P., Cotten,M., Lemey,P. and Nokes,D.J. |
| EPI_ISL_2582923, EPI_ISL_2582926, EPI_ISL_2582927, EPI_ISL_2582930, EPI_ISL_2582933 | J. Craig Venter Institute | J. Craig Venter Institute | Shabman,R., Fedorova,N., Puri,V., Shrivastava,S., Amedeo,P., Isom,R., Hu,L., Pickett,B., Novotny,M., Durbin,A., Rocchi,I., Williams,T., Hall,C.B., Tesini,B.L., Schnabel,K.C., Walsh,E.E. and Caserta,M. |
| EPI_ISL_2582935, EPI_ISL_2582938, EPI_ISL_2582941 | Broad Institute of MIT & Harvard | Broad Institute of MIT & Harvard | Newman,R.M., Zody,M.C., DeVincenzo,J.P., Grad,Y., Lipsitch,M., Murphy,R., Fitzgerald,M., Young,S., Gargeya,S., Poon,T.W., Charlebois,P., Weiner,B., Yang,X., Piper,M.E., McCowan,C., Ireland,A., Levin,J., Malboeuf,C., Qu,J., Chapman,S.B., Murphy,C., Wortman,J., Nusbaum,C. and Birren,B. |
| EPI_ISL_2582946, EPI_ISL_2582949 | J. Craig Venter Institute | J. Craig Venter Institute | Shabman,R., Fedorova,N., Puri,V., Shrivastava,S., Amedeo,P., Isom,R., Hu,L., Pickett,B., Novotny,M., Durbin,A., Rocchi,I., Williams,T., Hall,C.B., Tesini,B.L., Schnabel,K.C., Walsh,E.E. and Caserta,M. |
| EPI_ISL_2582951 | Broad Institute of MIT & Harvard | Broad Institute of MIT & Harvard | Newman,R.M., Zody,M.C., DeVincenzo,J.P., Grad,Y., Lipsitch,M., Murphy,R., Fitzgerald,M., Young,S., Gargeya,S., Poon,T.W., Charlebois,P., Weiner,B., Yang,X., Piper,M.E., McCowan,C., Ireland,A., Levin,J., Malboeuf,C., Qu,J., Chapman,S.B., Murphy,C., Wortman,J., Nusbaum,C. and Birren,B. |
| EPI_ISL_2582954 | J. Craig Venter Institute | J. Craig Venter Institute | Shabman,R., Fedorova,N., Puri,V., Shrivastava,S., Amedeo,P., Isom,R., Hu,L., Pickett,B., Novotny,M., Durbin,A., Rocchi,I., Williams,T., Hall,C.B., Tesini,B.L., Schnabel,K.C., Walsh,E.E. and Caserta,M. |
| EPI_ISL_2583023, EPI_ISL_2583024 | Epidemiology and Demography Department, KEMRI-Wellcome Trust Research Programme | Epidemiology and Demography Department, KEMRI-Wellcome Trust Research Programme | Otieno,J.R., Kamau,E.M., Oketch,J.W., Ngoi,J.M., Agoti,C.N., Gichuki,A.M., Otieno,G.P., Ngama,M., Cane,P.A., Kellam,P., Cotten,M., Lemey,P. and Nokes,D.J. |
| EPI_ISL_2583026 | Medicine, University of Washington, 300 9th Ave, Harborview Research & Training Building | Medicine, University of Washington, 300 9th Ave, Harborview Research & Training Building | Chu,H., Scott,E. and Roychoudhury,P. |
| EPI_ISL_2583029 | Broad Institute of MIT & Harvard | Broad Institute of MIT & Harvard | Newman,R.M., Zody,M.C., DeVincenzo,J.P., Grad,Y., Lipsitch,M., Murphy,R., Fitzgerald,M., Young,S., Gargeya,S., Poon,T.W., Charlebois,P., Weiner,B., Yang,X., Piper,M.E., McCowan,C., Ireland,A., Levin,J., Malboeuf,C., Qu,J., Chapman,S.B., Murphy,C., Wortman,J., Nusbaum,C. and Birren,B. |
| EPI_ISL_2583031 | Pediatrics - Infectious Diseases, Medical College of Wisconsin | Pediatrics - Infectious Diseases, Medical College of Wisconsin | Rebuffo-Scheer,C., Bose,M.E., He,J., Khaja,S., Ulatowski,M., Beck,E.T., Fan,J., Kumar,S., Nelson,M.I. and Henrickson,K.J. |

| | | | |
|---|---|---|---|
| EPI_ISL_2583034 | Broad Institute of MIT & Harvard | Broad Institute of MIT & Harvard | Newman,R.M., Zody,M.C., DeVincenzo,J.P., Grad,Y., Lipsitch,M., Murphy,R., Fitzgerald,M., Young,S., Gargeya,S., Poon,T.W., Charlebois,P., Weiner,B., Yang,X., Piper,M.E., McCowan,C., Ireland,A., Levin,J., Malboeuf,C., Qu,J., Chapman,S.B., Murphy,C., Wortman,J., Nusbaum,C. and Birren,B. |
| EPI_ISL_2583038 | Pediatrics, University of New Mexico | Pediatrics, University of New Mexico | Kothari,A., Kennedy,J.L., Schwalm,K.C., Putt,C., Denson,J.L. and Dinwiddie,D.L. |
| EPI_ISL_2583040 | Broad Institute of MIT & Harvard | Broad Institute of MIT & Harvard | Newman,R.M., Zody,M.C., DeVincenzo,J.P., Grad,Y., Lipsitch,M., Murphy,R., Fitzgerald,M., Young,S., Gargeya,S., Poon,T.W., Charlebois,P., Weiner,B., Yang,X., Piper,M.E., McCowan,C., Ireland,A., Levin,J., Malboeuf,C., Qu,J., Chapman,S.B., Murphy,C., Wortman,J., Nusbaum,C. and Birren,B. |
| EPI_ISL_2583042 | Pediatrics, University of New Mexico | Pediatrics, University of New Mexico | Kothari,A., Kennedy,J.L., Schwalm,K.C., Putt,C., Denson,J.L. and Dinwiddie,D.L. |
| EPI_ISL_2583045, EPI_ISL_2583046, EPI_ISL_2583048, EPI_ISL_2583050, EPI_ISL_2583054, EPI_ISL_2583055 | Broad Institute of MIT & Harvard | Broad Institute of MIT & Harvard | Newman,R.M., Zody,M.C., DeVincenzo,J.P., Grad,Y., Lipsitch,M., Murphy,R., Fitzgerald,M., Young,S., Gargeya,S., Poon,T.W., Charlebois,P., Weiner,B., Yang,X., Piper,M.E., McCowan,C., Ireland,A., Levin,J., Malboeuf,C., Qu,J., Chapman,S.B., Murphy,C., Wortman,J., Nusbaum,C. and Birren,B. |
| EPI_ISL_2583057 | Pediatrics - Infectious Diseases, Medical College of Wisconsin | Pediatrics - Infectious Diseases, Medical College of Wisconsin | Rebuffo-Scheer,C., Bose,M.E., He,J., Khaja,S., Ulatowski,M., Beck,E.T., Fan,J., Kumar,S., Nelson,M.I. and Henrickson,K.J. |
| EPI_ISL_2583061, EPI_ISL_2583062, EPI_ISL_2583063 | Pediatrics, University of New Mexico | Pediatrics, University of New Mexico | Kothari,A., Kennedy,J.L., Schwalm,K.C., Putt,C., Denson,J.L. and Dinwiddie,D.L. |
| EPI_ISL_2583065 | Pediatrics - Infectious Diseases, Medical College of Wisconsin | Pediatrics - Infectious Diseases, Medical College of Wisconsin | Rebuffo-Scheer,C., Bose,M.E., He,J., Khaja,S., Ulatowski,M., Beck,E.T., Fan,J., Kumar,S., Nelson,M.I. and Henrickson,K.J. |
| EPI_ISL_2583068, EPI_ISL_2583072, EPI_ISL_2583074 | Broad Institute of MIT & Harvard | Broad Institute of MIT & Harvard | Newman,R.M., Zody,M.C., DeVincenzo,J.P., Grad,Y., Lipsitch,M., Murphy,R., Fitzgerald,M., Young,S., Gargeya,S., Poon,T.W., Charlebois,P., Weiner,B., Yang,X., Piper,M.E., McCowan,C., Ireland,A., Levin,J., Malboeuf,C., Qu,J., Chapman,S.B., Murphy,C., Wortman,J., Nusbaum,C. and Birren,B. |
| EPI_ISL_2583076 | Pediatrics, University of New Mexico | Pediatrics, University of New Mexico | Kothari,A., Kennedy,J.L., Schwalm,K.C., Putt,C., Denson,J.L. and Dinwiddie,D.L. |
| EPI_ISL_2583078, EPI_ISL_2583080, EPI_ISL_2583082 | Broad Institute of MIT & Harvard | Broad Institute of MIT & Harvard | Newman,R.M., Zody,M.C., DeVincenzo,J.P., Grad,Y., Lipsitch,M., Murphy,R., Fitzgerald,M., Young,S., Gargeya,S., Poon,T.W., Charlebois,P., Weiner,B., Yang,X., Piper,M.E., McCowan,C., Ireland,A., Levin,J., Malboeuf,C., Qu,J., Chapman,S.B., Murphy,C., Wortman,J., Nusbaum,C. and Birren,B. |
| EPI_ISL_2583083 | Pediatrics - Infectious Diseases, Medical College of Wisconsin | Pediatrics - Infectious Diseases, Medical College of Wisconsin | Rebuffo-Scheer,C., Bose,M.E., He,J., Khaja,S., Ulatowski,M., Beck,E.T., Fan,J., Kumar,S., Nelson,M.I. and Henrickson,K.J. |
| EPI_ISL_2583085, EPI_ISL_2583087 | Broad Institute of MIT & Harvard | Broad Institute of MIT & Harvard | Newman,R.M., Zody,M.C., DeVincenzo,J.P., Grad,Y., Lipsitch,M., Murphy,R., Fitzgerald,M., Young,S., Gargeya,S., Poon,T.W., Charlebois,P., Weiner,B., Yang,X., Piper,M.E., McCowan,C., Ireland,A., Levin,J., Malboeuf,C., Qu,J., Chapman,S.B., Murphy,C., Wortman,J., Nusbaum,C. and Birren,B. |
| EPI_ISL_2583091 | Epidemiology and Demography Department, KEMRI-Wellcome Trust Research Programme | Epidemiology and Demography Department, KEMRI-Wellcome Trust Research Programme | Otieno,J.R., Kamau,E.M., Oketch,J.W., Ngoi,J.M., Agoti,C.N., Gichuki,A.M., Otieno,G.P., Ngama,M., Cane,P.A., Kellam,P., Cotten,M., Lemey,P. and Nokes,D.J. |
| EPI_ISL_2583092 | Broad Institute of MIT & Harvard | Broad Institute of MIT & Harvard | Newman,R.M., Zody,M.C., DeVincenzo,J.P., Grad,Y., Lipsitch,M., Murphy,R., Fitzgerald,M., Young,S., Gargeya,S., Poon,T.W., Charlebois,P., Weiner,B., Yang,X., Piper,M.E., McCowan,C., Ireland,A., Levin,J., Malboeuf,C., Qu,J., Chapman,S.B., Murphy,C., Wortman,J., Nusbaum,C. and Birren,B. |
| EPI_ISL_2585205 | J. Craig Venter Institute | J. Craig Venter Institute | Lorenzi,H., Town,C., Halpin,R., Bera,J., Ransier,A., Fedorova,N., Stockwell,T., Amedeo,P., Appalla,L., Bishop,B., Edworthy,P., Gupta,N., Hoover,J., Katzel,D., Li,K., Schobel,S., Shrivastava,S., Thovarai,V., Wang,S., Rebuffo-Scheer,C., Fan,J., He,J., Kehl,S.C., Lederboer,N., Jurgens,L.A., Bose,M.E., Beck,E.T., Kumar,S., Noyola,D.E., Wentworth,D.E. and Henrickson,K.J. |
| EPI_ISL_2585256, EPI_ISL_2585258 | J. Craig Venter Institute | J. Craig Venter Institute | Shabman,R., Das,S.R., Shilts,M., Fedorova,N., Puri,V., Shrivastava,S., Amedeo,P., Williams,M., Barratt,K., Mitchell,J. and Jennings,L. |
| EPI_ISL_2585259 | J. Craig Venter Institute | J. Craig Venter Institute | Das,S., Halpin,R.A., Bera,J., Puri,V., Fedorova,N., Tsitrin,T., Stockwell,T., Amedeo,P., Bishop,B., Katzel,D., Schobel,S., Shrivastava,S., Hartert,T., Moore,M., Chappell,J., Larkin,E., Wentworth,D.E. and Anderson,L.J. |
| EPI_ISL_2585260 | J. Craig Venter Institute | J. Craig Venter Institute | Das,S.R., Halpin,R.A., Shilts,M., Puri,V., Akopov,A., Fedorova,N., Stockwell,T., Amedeo,P., Bishop,B., Katzel,D., Schobel,S., Shrivastava,S. and Hartert,T. |
| EPI_ISL_2585261 | J. Craig Venter Institute | J. Craig Venter Institute | Shabman,R., Das,S.R., Shilts,M., Fedorova,N., Puri,V., Shrivastava,S., Amedeo,P., Williams,M., Barratt,K., Mitchell,J. and Jennings,L. |
| EPI_ISL_2585265 | J. Craig Venter Institute | J. Craig Venter Institute | Das,S.R., Halpin,R.A., Shilts,M., Puri,V., Akopov,A., Fedorova,N., Stockwell,T., Amedeo,P., Bishop,B., Katzel,D., Schobel,S., Shrivastava,S. and Hartert,T. |
| EPI_ISL_2585267, EPI_ISL_2585268, EPI_ISL_2585269, EPI_ISL_2585275, EPI_ISL_2585276, EPI_ISL_2585277, EPI_ISL_2585279 | J. Craig Venter Institute | J. Craig Venter Institute | Shabman,R., Das,S.R., Shilts,M., Fedorova,N., Puri,V., Shrivastava,S., Amedeo,P., Williams,M., Barratt,K., Mitchell,J. and Jennings,L. |
| EPI_ISL_2585280 | J. Craig Venter Institute | J. Craig Venter Institute | Das,S., Halpin,R.A., Bera,J., Fedorova,N., Tsitrin,T., Stockwell,T., Amedeo,P., Bishop,B., Gupta,N., Hoover,J., Katzel,D., Schobel,S., Shrivastava,S., Hartert,T., Moore,M., Chappell,J., Larkin,E., Wentworth,D.E. and Anderson,L.J. |
| EPI_ISL_2585281 | J. Craig Venter Institute | J. Craig Venter Institute | Das,S.R., Halpin,R.A., Shilts,M., Puri,V., Akopov,A., Fedorova,N., Stockwell,T., Amedeo,P., Bishop,B., Katzel,D., Schobel,S., Shrivastava,S. and Hartert,T. |
| EPI_ISL_2585284 | J. Craig Venter Institute | J. Craig Venter Institute | Shabman,R., Das,S.R., Puri,V., Fedorova,N., Amedeo,P., Williams,M., Shrivastava,S. and Halasa,N. |
| EPI_ISL_2585285 | J. Craig Venter Institute | J. Craig Venter Institute | Das,S.R., Halpin,R.A., Shilts,M., Puri,V., Akopov,A., Fedorova,N., Stockwell,T., Amedeo,P., Bishop,B., Katzel,D., Schobel,S., Shrivastava,S. and Hartert,T. |
| EPI_ISL_2585286 | J. Craig Venter Institute | J. Craig Venter Institute | Das,S., Halpin,R.A., Bera,J., Fedorova,N., Tsitrin,T., Stockwell,T., Amedeo,P., Bishop,B., Gupta,N., Hoover,J., Katzel,D., Schobel,S., Shrivastava,S., Hartert,T., Moore,M., Chappell,J., Larkin,E., Wentworth,D.E. and Anderson,L.J. |
| EPI_ISL_2585288, EPI_ISL_2585290 | J. Craig Venter Institute | J. Craig Venter Institute | Das,S.R., Halpin,R.A., Shilts,M., Puri,V., Akopov,A., Fedorova,N., Stockwell,T., Amedeo,P., Bishop,B., Katzel,D., Schobel,S., Shrivastava,S. and Hartert,T. |
| EPI_ISL_2585291 | J. Craig Venter Institute | J. Craig Venter Institute | Shabman,R., Das,S.R., Shilts,M., Fedorova,N., Puri,V., Shrivastava,S., Amedeo,P., Williams,M., Barratt,K., Mitchell,J. and Jennings,L. |
| EPI_ISL_2585292 | Laboratory Medicine, UW Virology | Laboratory Medicine, UW Virology | Lin,M.J., Tait,A. and Greninger,A.L. |
| EPI_ISL_2585302 | J. Craig Venter Institute | J. Craig Venter Institute | Das,S., Halpin,R.A., Bera,J., Puri,V., Fedorova,N., Tsitrin,T., Stockwell,T., Amedeo,P., Bishop,B., Katzel,D., Schobel,S., Shrivastava,S., Hartert,T., Moore,M., Chappell,J., Larkin,E., Wentworth,D.E. and Anderson,L.J. |
| EPI_ISL_2585307 | J. Craig Venter Institute | J. Craig Venter Institute | Shabman,R., Das,S.R., Shilts,M., Fedorova,N., Puri,V., Shrivastava,S., Amedeo,P., Williams,M., Barratt,K., Mitchell,J. and Jennings,L. |
| EPI_ISL_2585309 | J. Craig Venter Institute | J. Craig Venter Institute | Das,S.R., Halpin,R.A., Shilts,M., Puri,V., Akopov,A., Fedorova,N., Stockwell,T., Amedeo,P., Bishop,B., Katzel,D., Schobel,S., Shrivastava,S. and Hartert,T. |
| EPI_ISL_2585311, EPI_ISL_2585312, EPI_ISL_2585313, EPI_ISL_2585316, EPI_ISL_2585318 | J. Craig Venter Institute | J. Craig Venter Institute | Shabman,R., Das,S.R., Shilts,M., Fedorova,N., Puri,V., Shrivastava,S., Amedeo,P., Williams,M., Barratt,K., Mitchell,J. and Jennings,L. |
| EPI_ISL_2585319 | J. Craig Venter Institute | J. Craig Venter Institute | Das,S., Halpin,R.A., Bera,J., Fedorova,N., Tsitrin,T., Stockwell,T., Amedeo,P., Bishop,B., Gupta,N., Hoover,J., Katzel,D., Schobel,S., Shrivastava,S., Hartert,T., Moore,M., Chappell,J., Larkin,E., Wentworth,D.E. and Anderson,L.J. |
| EPI_ISL_2585320, EPI_ISL_2585323, EPI_ISL_2585325, EPI_ISL_2585327, EPI_ISL_2585329, EPI_ISL_2585332, EPI_ISL_2587159 | J. Craig Venter Institute | J. Craig Venter Institute | Das,S.R., Halpin,R.A., Shilts,M., Puri,V., Akopov,A., Fedorova,N., Stockwell,T., Amedeo,P., Bishop,B., Katzel,D., Schobel,S., Shrivastava,S. and Hartert,T. |
| EPI_ISL_2587163 | J. Craig Venter Institute | J. Craig Venter Institute | Shabman,R., Das,S.R., Puri,V., Fedorova,N., Amedeo,P., Williams,M., Shrivastava,S. and Halasa,N. |
| EPI_ISL_2587342 | J. Craig Venter Institute | J. Craig Venter Institute | Shabman,R., Das,S.R., Shilts,M., Fedorova,N., Puri,V., Shrivastava,S., Amedeo,P., Williams,M., Barratt,K., Mitchell,J. and Jennings,L. |
| EPI_ISL_2587344 | J. Craig Venter Institute | J. Craig Venter Institute | Shabman,R., Das,S.R., Shilts,M., Fedorova,N., Puri,V., Shrivastava,S., Amedeo,P., Hu,L., Durbin,A., Rocchi,I., Williams,T. and Hartert,T. |
| EPI_ISL_2587345 | J. Craig Venter Institute | J. Craig Venter Institute | Das,S., Halpin,R.A., Bera,J., Fedorova,N., Tsitrin,T., Stockwell,T., Amedeo,P., Bishop,B., Gupta,N., Hoover,J., Katzel,D., Schobel,S., Shrivastava,S., Hartert,T., Moore,M., Chappell,J., Larkin,E., Wentworth,D.E. and Anderson,L.J. |
| EPI_ISL_2587349 | J. Craig Venter Institute | J. Craig Venter Institute | Shabman,R., Das,S.R., Shilts,M., Fedorova,N., Puri,V., Shrivastava,S., Amedeo,P., Hu,L., Durbin,A., Rocchi,I., Williams,T. and Hartert,T. |
| EPI_ISL_2587351, EPI_ISL_2587353 | J. Craig Venter Institute | J. Craig Venter Institute | Das,S., Halpin,R.A., Bera,J., Fedorova,N., Tsitrin,T., Stockwell,T., Amedeo,P., Bishop,B., Gupta,N., Hoover,J., Katzel,D., Schobel,S., Shrivastava,S., Hartert,T., Moore,M., Chappell,J., Larkin,E., Wentworth,D.E. and Anderson,L.J. |
| EPI_ISL_2587355, EPI_ISL_2587357 | J. Craig Venter Institute | J. Craig Venter Institute | Das,S.R., Halpin,R.A., Shilts,M., Puri,V., Akopov,A., Fedorova,N., Stockwell,T., Amedeo,P., Bishop,B., Katzel,D., Schobel,S., Shrivastava,S. and Hartert,T. |
| EPI_ISL_2587358 | J. Craig Venter Institute | J. Craig Venter Institute | Das,S., Halpin,R.A., Bera,J., Fedorova,N., Tsitrin,T., Stockwell,T., Amedeo,P., Bishop,B., Gupta,N., Hoover,J., Katzel,D., Schobel,S., Shrivastava,S., Hartert,T., Moore,M., Chappell,J., Larkin,E., Wentworth,D.E. and Anderson,L.J. |
| EPI_ISL_2587362, EPI_ISL_2587364 | J. Craig Venter Institute | J. Craig Venter Institute | Das,S.R., Halpin,R.A., Shilts,M., Puri,V., Akopov,A., Fedorova,N., Stockwell,T., Amedeo,P., Bishop,B., Katzel,D., Schobel,S., Shrivastava,S. and Hartert,T. |
| EPI_ISL_2587366 | J. Craig Venter Institute | J. Craig Venter Institute | Das,S., Halpin,R.A., Bera,J., Fedorova,N., Tsitrin,T., Stockwell,T., Amedeo,P., Bishop,B., Gupta,N., Hoover,J., Katzel,D., Schobel,S., Shrivastava,S., Hartert,T., Moore,M., Chappell,J., Larkin,E., Wentworth,D.E. and Anderson,L.J. |
| EPI_ISL_2587379, EPI_ISL_2587381, EPI_ISL_2587383, EPI_ISL_2587385, EPI_ISL_2587387 | J. Craig Venter Institute | J. Craig Venter Institute | Shabman,R., Das,S.R., Shilts,M., Fedorova,N., Puri,V., Shrivastava,S., Amedeo,P., Williams,M., Barratt,K., Mitchell,J. and Jennings,L. |
| EPI_ISL_2587389 | J. Craig Venter Institute | J. Craig Venter Institute | Shabman,R., Das,S.R., Puri,V., Fedorova,N., Amedeo,P., Williams,M., Shrivastava,S. and Halasa,N. |
| EPI_ISL_2587391, EPI_ISL_2587393, EPI_ISL_2587396, EPI_ISL_2587398, EPI_ISL_2587400, EPI_ISL_2587402, EPI_ISL_2587404 | J. Craig Venter Institute | J. Craig Venter Institute | Shabman,R., Das,S.R., Shilts,M., Fedorova,N., Puri,V., Shrivastava,S., Amedeo,P., Williams,M., Barratt,K., Mitchell,J. and Jennings,L. |
| EPI_ISL_2587406 | J. Craig Venter Institute | J. Craig Venter Institute | Das,S., Halpin,R.A., Bera,J., Puri,V., Fedorova,N., Tsitrin,T., Stockwell,T., Amedeo,P., Bishop,B., Katzel,D., Schobel,S., Shrivastava,S., Hartert,T., Moore,M., Chappell,J., Larkin,E., Wentworth,D.E. and Anderson,L.J. |
| EPI_ISL_2587408, EPI_ISL_2587410 | J. Craig Venter Institute | J. Craig Venter Institute | Shabman,R., Das,S.R., Shilts,M., Fedorova,N., Puri,V., Shrivastava,S., Amedeo,P., Williams,M., Barratt,K., Mitchell,J. and Jennings,L. |
| EPI_ISL_2587412 | J. Craig Venter Institute | J. Craig Venter Institute | Shabman,R., Das,S.R., Shilts,M., Fedorova,N., Puri,V., Shrivastava,S., Amedeo,P., Hu,L., Durbin,A., Rocchi,I., Williams,T. and Hartert,T. |
| EPI_ISL_2587414, EPI_ISL_2587415 | J. Craig Venter Institute | J. Craig Venter Institute | Das,S., Halpin,R.A., Bera,J., Fedorova,N., Tsitrin,T., Stockwell,T., Amedeo,P., Bishop,B., Gupta,N., Hoover,J., Katzel,D., Schobel,S., Shrivastava,S., Hartert,T., Moore,M., Chappell,J., Larkin,E., Wentworth,D.E. and Anderson,L.J. |
| EPI_ISL_2587417, EPI_ISL_2587421, EPI_ISL_2587423, EPI_ISL_2587425, EPI_ISL_2587429, EPI_ISL_2587431, EPI_ISL_2587433 | J. Craig Venter Institute | J. Craig Venter Institute | Das,S.R., Halpin,R.A., Shilts,M., Puri,V., Akopov,A., Fedorova,N., Stockwell,T., Amedeo,P., Bishop,B., Katzel,D., Schobel,S., Shrivastava,S. and Hartert,T. |
| EPI_ISL_2587434 | J. Craig Venter Institute | J. Craig Venter Institute | Das,S., Halpin,R.A., Bera,J., Puri,V., Fedorova,N., Tsitrin,T., Stockwell,T., Amedeo,P., Bishop,B., Katzel,D., Schobel,S., Shrivastava,S., Hartert,T., Moore,M., Chappell,J., Larkin,E., Wentworth,D.E. and Anderson,L.J. |
| EPI_ISL_2587436, EPI_ISL_2587438 | J. Craig Venter Institute | J. Craig Venter Institute | Das,S., Halpin,R.A., Bera,J., Fedorova,N., Tsitrin,T., Stockwell,T., Amedeo,P., Bishop,B., Gupta,N., Hoover,J., Katzel,D., Schobel,S., Shrivastava,S., Hartert,T., Moore,M., Chappell,J., Larkin,E., Wentworth,D.E. and Anderson,L.J. |
| EPI_ISL_2587440, EPI_ISL_2587442, EPI_ISL_2587444, EPI_ISL_2587446, EPI_ISL_2587447, EPI_ISL_2587451, EPI_ISL_2587452, EPI_ISL_2587454, EPI_ISL_2587456, EPI_ISL_2587458 | J. Craig Venter Institute | J. Craig Venter Institute | Das,S.R., Halpin,R.A., Shilts,M., Puri,V., Akopov,A., Fedorova,N., Stockwell,T., Amedeo,P., Bishop,B., Katzel,D., Schobel,S., Shrivastava,S. and Hartert,T. |
| EPI_ISL_2587469 | J. Craig Venter Institute | J. Craig Venter Institute | Das,S., Halpin,R.A., Bera,J., Fedorova,N., Tsitrin,T., Stockwell,T., Amedeo,P., Bishop,B., Gupta,N., Hoover,J., Katzel,D., Schobel,S., Shrivastava,S., Hartert,T., Moore,M., Chappell,J., Larkin,E., Wentworth,D.E. and Anderson,L.J. |
| EPI_ISL_2587471, EPI_ISL_2587473, EPI_ISL_2587494 | J. Craig Venter Institute | J. Craig Venter Institute | Das,S.R., Halpin,R.A., Shilts,M., Puri,V., Akopov,A., Fedorova,N., Stockwell,T., Amedeo,P., Bishop,B., Katzel,D., Schobel,S., Shrivastava,S. and Hartert,T. |

| | | | |
|---|---|---|---|
| EPI_ISL_2588290, EPI_ISL_2588292 | J. Craig Venter Institute | J. Craig Venter Institute | Das,S., Halpin,R.A., Bera,J., Fedorova,N., Tsitrin,T., Stockwell,T., Amedeo,P., Bishop,B., Gupta,N., Hoover,J., Katzel,D., Schobel,S., Shrivastava,S., Hartert,T., Moore,M., Chappell,J., Larkin,E., Wentworth,D.E. and Anderson,L.J. |
| EPI_ISL_2588301, EPI_ISL_2588302, EPI_ISL_2588303 | J. Craig Venter Institute | J. Craig Venter Institute | Das,S.R., Halpin,R.A., Shilts,M., Puri,V., Akopov,A., Fedorova,N., Stockwell,T., Amedeo,P., Bishop,B., Katzel,D., Schobel,S., Shrivastava,S. and Hartert,T. |
| EPI_ISL_2588304 | J. Craig Venter Institute | J. Craig Venter Institute | Das,S., Halpin,R.A., Bera,J., Fedorova,N., Tsitrin,T., Stockwell,T., Amedeo,P., Bishop,B., Gupta,N., Hoover,J., Katzel,D., Schobel,S., Shrivastava,S., Hartert,T., Moore,M., Chappell,J., Larkin,E., Wentworth,D.E. and Anderson,L.J. |
| EPI_ISL_2588305, EPI_ISL_2588306 | J. Craig Venter Institute | J. Craig Venter Institute | Das,S.R., Halpin,R.A., Shilts,M., Puri,V., Akopov,A., Fedorova,N., Stockwell,T., Amedeo,P., Bishop,B., Katzel,D., Schobel,S., Shrivastava,S. and Hartert,T. |
| EPI_ISL_2588347 | J. Craig Venter Institute | J. Craig Venter Institute | Das,S., Halpin,R.A., Bera,J., Fedorova,N., Tsitrin,T., Stockwell,T., Amedeo,P., Bishop,B., Gupta,N., Hoover,J., Katzel,D., Schobel,S., Shrivastava,S., Hartert,T., Moore,M., Chappell,J., Larkin,E., Wentworth,D.E. and Anderson,L.J. |
| EPI_ISL_2588348 | J. Craig Venter Institute | J. Craig Venter Institute | Das,S., Halpin,R.A., Bera,J., Puri,V., Fedorova,N., Tsitrin,T., Stockwell,T., Amedeo,P., Bishop,B., Katzel,D., Schobel,S., Shrivastava,S., Hartert,T., Moore,M., Chappell,J., Larkin,E., Wentworth,D.E. and Anderson,L.J. |
| EPI_ISL_2588349, EPI_ISL_2588350, EPI_ISL_2588351 | J. Craig Venter Institute | J. Craig Venter Institute | Das,S., Halpin,R.A., Bera,J., Fedorova,N., Tsitrin,T., Stockwell,T., Amedeo,P., Bishop,B., Gupta,N., Hoover,J., Katzel,D., Schobel,S., Shrivastava,S., Hartert,T., Moore,M., Chappell,J., Larkin,E., Wentworth,D.E. and Anderson,L.J. |
| EPI_ISL_2588352 | J. Craig Venter Institute | J. Craig Venter Institute | Das,S., Halpin,R.A., Bera,J., Puri,V., Fedorova,N., Tsitrin,T., Stockwell,T., Amedeo,P., Bishop,B., Katzel,D., Schobel,S., Shrivastava,S., Hartert,T., Moore,M., Chappell,J., Larkin,E., Wentworth,D.E. and Anderson,L.J. |
| EPI_ISL_2588353, EPI_ISL_2588354, EPI_ISL_2588355 | J. Craig Venter Institute | J. Craig Venter Institute | Das,S.R., Halpin,R.A., Shilts,M., Puri,V., Akopov,A., Fedorova,N., Stockwell,T., Amedeo,P., Bishop,B., Katzel,D., Schobel,S., Shrivastava,S. and Hartert,T. |
| EPI_ISL_2588356 | J. Craig Venter Institute | J. Craig Venter Institute | Shabman,R., Das,S.R., Shilts,M., Fedorova,N., Puri,V., Shrivastava,S., Amedeo,P., Williams,M., Barratt,K., Mitchell,J. and Jennings,L. |
| EPI_ISL_2588357, EPI_ISL_2588358, EPI_ISL_2588359, EPI_ISL_2588360, EPI_ISL_2588361, EPI_ISL_2588362 | J. Craig Venter Institute | J. Craig Venter Institute | Das,S., Halpin,R.A., Bera,J., Fedorova,N., Tsitrin,T., Stockwell,T., Amedeo,P., Bishop,B., Gupta,N., Hoover,J., Katzel,D., Schobel,S., Shrivastava,S., Hartert,T., Moore,M., Chappell,J., Larkin,E., Wentworth,D.E. and Anderson,L.J. |
| EPI_ISL_2588363, EPI_ISL_2588364, EPI_ISL_2588365, EPI_ISL_2588366, EPI_ISL_2588367, EPI_ISL_2588368, EPI_ISL_2588369, EPI_ISL_2588370, EPI_ISL_2588371, EPI_ISL_2588372, EPI_ISL_2588373, EPI_ISL_2588374, EPI_ISL_2588375 | | | |
| see above | J. Craig Venter Institute | J. Craig Venter Institute | Das,S.R., Halpin,R.A., Shilts,M., Puri,V., Akopov,A., Fedorova,N., Stockwell,T., Amedeo,P., Bishop,B., Katzel,D., Schobel,S., Shrivastava,S. and Hartert,T. |
| EPI_ISL_2588376 | J. Craig Venter Institute | J. Craig Venter Institute | Shabman,R., Das,S.R., Puri,V., Fedorova,N., Amedeo,P., Williams,M., Shrivastava,S. and Halasa,N. |
| EPI_ISL_2588377, EPI_ISL_2588378 | J. Craig Venter Institute | J. Craig Venter Institute | Das,S.R., Halpin,R.A., Shilts,M., Puri,V., Akopov,A., Fedorova,N., Stockwell,T., Amedeo,P., Bishop,B., Katzel,D., Schobel,S., Shrivastava,S. and Hartert,T. |
| EPI_ISL_2588379 | J. Craig Venter Institute | J. Craig Venter Institute | Wentworth,D.E., Halpin,R.A., Bera,J., Lin,X., Fedorova,N., Tsitrin,T., McLellan,M., Stockwell,T., Amedeo,P., Bishop,B., Gupta,N., Hoover,J., Katzel,D., Schobel,S., Shrivastava,S., Garcia,J., Laguna-Torres,V.A., Leguia,M., Benavides,J.G. and Halsey,E. |
| EPI_ISL_2588380 | J. Craig Venter Institute | J. Craig Venter Institute | Das,S.R., Halpin,R.A., Shilts,M., Puri,V., Akopov,A., Fedorova,N., Stockwell,T., Amedeo,P., Bishop,B., Katzel,D., Schobel,S., Shrivastava,S. and Hartert,T. |
| EPI_ISL_2588437 | J. Craig Venter Institute | J. Craig Venter Institute | Shabman,R., Das,S.R., Puri,V., Fedorova,N., Amedeo,P., Williams,M., Shrivastava,S. and Halasa,N. |
| EPI_ISL_2588597 | J. Craig Venter Institute | J. Craig Venter Institute | Das,S., Halpin,R.A., Bera,J., Fedorova,N., Tsitrin,T., Stockwell,T., Amedeo,P., Bishop,B., Gupta,N., Hoover,J., Katzel,D., Schobel,S., Shrivastava,S., Hartert,T., Moore,M., Chappell,J., Larkin,E., Wentworth,D.E. and Anderson,L.J. |
| EPI_ISL_2588598, EPI_ISL_2588599, EPI_ISL_2588600 | J. Craig Venter Institute | J. Craig Venter Institute | Das,S.R., Halpin,R.A., Shilts,M., Puri,V., Akopov,A., Fedorova,N., Stockwell,T., Amedeo,P., Bishop,B., Katzel,D., Schobel,S., Shrivastava,S. and Hartert,T. |
| EPI_ISL_2588794 | Virology Laboratory, Dr. Ricardo Gutierrez Children Hospital | Virology Laboratory, Dr. Ricardo Gutierrez Children Hospital | Goya,S., Rojo,G.L., Valinotto,L.E., Mistchenko,A.S. and Viegas,M. |
| EPI_ISL_2588795 | Virology Laboratory, Dr. Ricardo Gutierrez Children Hospital | Virology Laboratory, Dr. Ricardo Gutierrez Children Hospital | Goya,S., Valinotto,L.E., Tittarelli,E., Rojo,G.L., Greninger,A., Zaiat,J., Marti,M., Mistchenko,A.S. and Viegas,M. |
| EPI_ISL_2588796, EPI_ISL_2588818 | Virology Laboratory, Dr. Ricardo Gutierrez Children Hospital | Virology Laboratory, Dr. Ricardo Gutierrez Children Hospital | Goya,S., Valinotto,L.E., Tittarelli,E., Rojo,G.L., Greninger,A., Luso,S., Natale,M., Mistchenko,A.S. and Viegas,M. |
| EPI_ISL_2588827 | Virology Laboratory, Dr. Ricardo Gutierrez Children Hospital | Virology Laboratory, Dr. Ricardo Gutierrez Children Hospital | Goya,S., Valinotto,L.E., Tittarelli,E., Rojo,G.L., Greninger,A., Zaiat,J., Marti,M., Mistchenko,A.S. and Viegas,M. |
| EPI_ISL_2588828 | Virology Laboratory, Dr. Ricardo Gutierrez Children Hospital | Virology Laboratory, Dr. Ricardo Gutierrez Children Hospital | Goya,S., Rojo,G.L., Valinotto,L.E., Mistchenko,A.S. and Viegas,M. |
| EPI_ISL_2592492 | J. Craig Venter Institute | J. Craig Venter Institute | Shabman,R., Fedorova,N., Puri,V., Shrivastava,S., Amedeo,P., Isom,R., Hu,L., Pickett,B., Novotny,M., Durbin,A., Rocchi,I., Williams,T., Hall,C.B., Tesini,B.L., Schnabel,K.C., Walsh,E.E. and Caserta,M. |
| EPI_ISL_2592529 | Pediatrics - Infectious Diseases, Medical College of Wisconsin | Pediatrics - Infectious Diseases, Medical College of Wisconsin | Rebuffo-Scheer,C., Bose,M.E., He,J., Khaja,S., Ulatowski,M., Beck,E.T., Fan,J., Kumar,S., Nelson,M.I. and Henrickson,K.J. |
| EPI_ISL_2592567, EPI_ISL_2592568, EPI_ISL_2592569, EPI_ISL_2592570, EPI_ISL_2592571, EPI_ISL_2592572 | Broad Institute of MIT & Harvard | Broad Institute of MIT & Harvard | Newman,R.M., Zody,M.C., DeVincenzo,J.P., Grad,Y., Lipsitch,M., Murphy,R., Fitzgerald,M., Young,S., Gargeya,S., Poon,T.W., Charlebois,P., Weiner,B., Yang,X., Piper,M.E., McCowan,C., Ireland,A., Levin,J., Malboeuf,C., Qu,J., Chapman,S.B., Murphy,C., Wortman,J., Nusbaum,C. and Birren,B. |
| EPI_ISL_2592783 | Pediatrics - Infectious Diseases, Medical College of Wisconsin | Pediatrics - Infectious Diseases, Medical College of Wisconsin | Rebuffo-Scheer,C., Bose,M.E., He,J., Khaja,S., Ulatowski,M., Beck,E.T., Fan,J., Kumar,S., Nelson,M.I. and Henrickson,K.J. |
| EPI_ISL_2593173 | Virology, University of Washington | Virology, University of Washington | Greninger,A.L., Shean,R.C. and Makhsous,N. |
| EPI_ISL_2594878, EPI_ISL_2594879, EPI_ISL_2595159, EPI_ISL_2595160 | Center for Infectious Diseases, School of Public Health, University of Texas Health Science Center | Center for Infectious Diseases, School of Public Health, University of Texas Health Science Center | Bahl,J., Hixson,J., Kim,D.-K., Qiu,X., Piedra,P.A., Piedra,F.-A., Avadhanula,V. and Machado,A.A. |
| EPI_ISL_2595173 | Beijing Key Laboratory of Etiology of Viral Diseases in Children; Laboratory of Virology, Capital Institute of Pediatrics | Beijing Key Laboratory of Etiology of Viral Diseases in Children; Laboratory of Virology, Capital Institute of Pediatrics | Cui,G., Zhu,R., Deng,J., Zhao,L., Sun,Y., Wang,F. and Qian,Y. |
| EPI_ISL_2595181, EPI_ISL_2595182 | Center for Infectious Diseases, School of Public Health, University of Texas Health Science Center | Center for Infectious Diseases, School of Public Health, University of Texas Health Science Center | Bahl,J., Hixson,J., Kim,D.-K., Qiu,X., Piedra,P.A., Piedra,F.-A., Avadhanula,V. and Machado,A.A. |
| EPI_ISL_2595188, EPI_ISL_2595192, EPI_ISL_2595196, EPI_ISL_2595197, EPI_ISL_2595198, EPI_ISL_2595205 | Epidemiology and Demography Department, KEMRI-Wellcome Trust Research Programme | Epidemiology and Demography Department, KEMRI-Wellcome Trust Research Programme | Otieno,J.R., Kamau,E.M., Oketch,J.W., Ngoi,J.M., Agoti,C.N., Gichuki,A.M., Otieno,G.P., Ngama,M., Cane,P.A., Kellam,P., Cotten,M., Lemey,P. and Nokes,D.J. |
| EPI_ISL_2595236, EPI_ISL_2595277, EPI_ISL_2595278, EPI_ISL_2595279 | Broad Institute of MIT & Harvard | Broad Institute of MIT & Harvard | Newman,R.M., Zody,M.C., DeVincenzo,J.P., Grad,Y., Lipsitch,M., Murphy,R., Fitzgerald,M., Young,S., Gargeya,S., Poon,T.W., Charlebois,P., Weiner,B., Yang,X., Piper,M.E., McCowan,C., Ireland,A., Levin,J., Malboeuf,C., Qu,J., Chapman,S.B., Murphy,C., Wortman,J., Nusbaum,C. and Birren,B. |
| EPI_ISL_2595315 | Epidemiology and Demography Department, KEMRI-Wellcome Trust Research Programme | Epidemiology and Demography Department, KEMRI-Wellcome Trust Research Programme | Otieno,J.R., Kamau,E.M., Oketch,J.W., Ngoi,J.M., Agoti,C.N., Gichuki,A.M., Otieno,G.P., Ngama,M., Cane,P.A., Kellam,P., Cotten,M., Lemey,P. and Nokes,D.J. |
| EPI_ISL_2595321 | J. Craig Venter Institute | J. Craig Venter Institute | Shrivastava,S., Halpin,R.A., Puri,V., Fedorova,N.B., Stockwell,T., Amedeo,P., Katzel,D., Schobel,S., Pickett,B.E., Moore,M., Chappell,J., Larkin,E., Wentworth,D.E., Anderson,L.J. and Hartert,T. |
| EPI_ISL_2595357 | Epidemiology and Demography Department, KEMRI-Wellcome Trust Research Programme | Epidemiology and Demography Department, KEMRI-Wellcome Trust Research Programme | Otieno,J.R., Kamau,E.M., Oketch,J.W., Ngoi,J.M., Agoti,C.N., Gichuki,A.M., Otieno,G.P., Ngama,M., Cane,P.A., Kellam,P., Cotten,M., Lemey,P. and Nokes,D.J. |
| EPI_ISL_2595477 | Marie Bashir Institute for Infectious Diseases and Biosecurity & Sydney Medical School, The University of Sydney, Westmead Institute for Medical Research | Marie Bashir Institute for Infectious Diseases and Biosecurity & Sydney Medical School, The University of Sydney, Westmead Institute for Medical Research | Eden,J.-S., Kok,J., Dwyer,D.E., Fernandez,M., Carter,I. and Holmes,E.C. |
| EPI_ISL_2595540, EPI_ISL_2595541 | Epidemiology and Demography Department, KEMRI-Wellcome Trust Research Programme | Epidemiology and Demography Department, KEMRI-Wellcome Trust Research Programme | Otieno,J.R., Kamau,E.M., Oketch,J.W., Ngoi,J.M., Agoti,C.N., Gichuki,A.M., Otieno,G.P., Ngama,M., Cane,P.A., Kellam,P., Cotten,M., Lemey,P. and Nokes,D.J. |
| EPI_ISL_2595546 | Department of Pediatrics, Center of Excellence in Clinical Virology, Chulalongkorn | Department of Pediatrics, Center of Excellence in Clinical Virology, Chulalongkorn | Thongpan,I. |
| EPI_ISL_2595548, EPI_ISL_2595550 | Epidemiology and Demography Department, KEMRI-Wellcome Trust Research Programme | Epidemiology and Demography Department, KEMRI-Wellcome Trust Research Programme | Otieno,J.R., Kamau,E.M., Oketch,J.W., Ngoi,J.M., Agoti,C.N., Gichuki,A.M., Otieno,G.P., Ngama,M., Cane,P.A., Kellam,P., Cotten,M., Lemey,P. and Nokes,D.J. |
| EPI_ISL_2595591, EPI_ISL_2595592, EPI_ISL_2595604 | Kazuya Shirato National Institute of Infectious Diseases, Virology III | Kazuya Shirato National Institute of Infectious Diseases, Virology III | Shirato,K., Sato,K., Dapat,I., Nao,N., Omiya,S., Matsuyama,S., Takeda,M. and Nishimura,H. |
| EPI_ISL_2595605, EPI_ISL_2595607, EPI_ISL_2595608, EPI_ISL_2595609 | Marie Bashir Institute for Infectious Diseases and Biosecurity & Sydney Medical School, The University of Sydney, Westmead Institute for Medical Research | Marie Bashir Institute for Infectious Diseases and Biosecurity & Sydney Medical School, The University of Sydney, Westmead Institute for Medical Research | Eden,J.-S., Kok,J., Dwyer,D.E., Fernandez,M., Carter,I. and Holmes,E.C. |
| EPI_ISL_2595626, EPI_ISL_2595627, EPI_ISL_2595628, EPI_ISL_2595631, EPI_ISL_2595633, EPI_ISL_2595634, EPI_ISL_2595636, EPI_ISL_2595638, EPI_ISL_2595639, EPI_ISL_2595640, EPI_ISL_2595641, EPI_ISL_2595642, EPI_ISL_2595644 | | | |
| see above | Epidemiology and Demography Department, KEMRI-Wellcome Trust Research Programme | Epidemiology and Demography Department, KEMRI-Wellcome Trust Research Programme | Otieno,J.R., Kamau,E.M., Oketch,J.W., Ngoi,J.M., Agoti,C.N., Gichuki,A.M., Otieno,G.P., Ngama,M., Cane,P.A., Kellam,P., Cotten,M., Lemey,P. and Nokes,D.J. |
| EPI_ISL_2595651 | Lab Medicine, UW | Lab Medicine, UW | Greninger,A.L., Makhsous,N., Kuypers,J.M., Shean,R.C. and Jerome,K.R. |
| EPI_ISL_2595653, EPI_ISL_2595654, EPI_ISL_2595655, EPI_ISL_2595657, EPI_ISL_2595658, EPI_ISL_2595660, EPI_ISL_2595661, EPI_ISL_2595663, EPI_ISL_2595665, EPI_ISL_2595672, EPI_ISL_2595673 | | | |
| see above | Epidemiology and Demography Department, KEMRI-Wellcome Trust Research Programme | Epidemiology and Demography Department, KEMRI-Wellcome Trust Research Programme | Otieno,J.R., Kamau,E.M., Oketch,J.W., Ngoi,J.M., Agoti,C.N., Gichuki,A.M., Otieno,G.P., Ngama,M., Cane,P.A., Kellam,P., Cotten,M., Lemey,P. and Nokes,D.J. |
| EPI_ISL_2595681, EPI_ISL_2595684 | Marie Bashir Institute for Infectious Diseases and Biosecurity & Sydney Medical School, The University of Sydney, Westmead Institute for Medical Research | Marie Bashir Institute for Infectious Diseases and Biosecurity & Sydney Medical School, The University of Sydney, Westmead Institute for Medical Research | Eden,J.-S., Kok,J., Dwyer,D.E., Fernandez,M., Carter,I. and Holmes,E.C. |
| EPI_ISL_2595691 | J. Craig Venter Institute | J. Craig Venter Institute | Tan,G., Pickett,B., Fedorova,N., Amedeo,P., Hu,L., Christensen,J., Miller,J., Durbin,A., Williams,T., Arumemi,F., Cadiz,C., Alanis,R., Balmseda,A., Wiliams,T., Schiller,A., Patel,M., Kubale,J. and Gordon,A. |
| EPI_ISL_2595696 | Pediatrics, University of New Mexico | Pediatrics, University of New Mexico | Kothari,A., Kennedy,J.L., Schwalm,K.C., Putt,C., Denson,J.L. and Dinwiddie,D.L. |
| EPI_ISL_2595697 | Epidemiology and Demography Department, KEMRI-Wellcome Trust Research Programme | Epidemiology and Demography Department, KEMRI-Wellcome Trust Research Programme | Otieno,J.R., Kamau,E.M., Oketch,J.W., Ngoi,J.M., Agoti,C.N., Gichuki,A.M., Otieno,G.P., Ngama,M., Cane,P.A., Kellam,P., Cotten,M., Lemey,P. and Nokes,D.J. |

| | | | |
|---|---|---|---|
| EPI_ISL_2595699, EPI_ISL_2595700, EPI_ISL_2595701 | Pediatrics, University of New Mexico | Pediatrics, University of New Mexico | Kothari,A., Kennedy,J.L., Schwalm,K.C., Putt,C., Denson,J.L. and Dinwiddie,D.L. |
| EPI_ISL_2595702 | Epidemiology and Demography Department, KEMRI-Wellcome Trust Research Programme | Epidemiology and Demography Department, KEMRI-Wellcome Trust Research Programme | Otieno,J.R., Kamau,E.M., Oketch,J.W., Ngoi,J.M., Agoti,C.N., Gichuki,A.M., Otieno,G.P., Ngama,M., Cane,P.A., Kellam,P., Cotten,M., Lemey,P. and Nokes,D.J. |
| EPI_ISL_2595703, EPI_ISL_2595704, EPI_ISL_2595705 | Pediatrics, University of New Mexico | Pediatrics, University of New Mexico | Kothari,A., Kennedy,J.L., Schwalm,K.C., Putt,C., Denson,J.L. and Dinwiddie,D.L. |
| EPI_ISL_2595706, EPI_ISL_2595707, EPI_ISL_2595708, EPI_ISL_2595709, EPI_ISL_2595710, EPI_ISL_2595711, EPI_ISL_2595712 | Marie Bashir Institute for Infectious Diseases and Biosecurity & Sydney Medical School, The University of Sydney, Westmead Institute for Medical Research | Marie Bashir Institute for Infectious Diseases and Biosecurity & Sydney Medical School, The University of Sydney, Westmead Institute for Medical Research | Eden,J.-S., Kok,J., Dwyer,D.E., Fernandez,M., Carter,I. and Holmes,E.C. |
| EPI_ISL_2595713 | Department of Pediatrics, Center of Excellence in Clinical Virology, Chulalongkorn | Department of Pediatrics, Center of Excellence in Clinical Virology, Chulalongkorn | Thongpan,I. |
| EPI_ISL_2595714, EPI_ISL_2595715, EPI_ISL_2595716 | Marie Bashir Institute for Infectious Diseases and Biosecurity & Sydney Medical School, The University of Sydney, Westmead Institute for Medical Research | Marie Bashir Institute for Infectious Diseases and Biosecurity & Sydney Medical School, The University of Sydney, Westmead Institute for Medical Research | Eden,J.-S., Kok,J., Dwyer,D.E., Fernandez,M., Carter,I. and Holmes,E.C. |
| EPI_ISL_2595717 | J. Craig Venter Institute | J. Craig Venter Institute | Tan,G., Pickett,B., Fedorova,N., Amedeo,P., Hu,L., Christensen,J., Miller,J., Durbin,A., Williams,T., Arumemi,F., Cadiz,C., Alanis,R., Balmseda,A., Wiliams,T., Schiller,A., Patel,M., Kubale,J. and Gordon,A. |
| EPI_ISL_2595718, EPI_ISL_2595719 | Marie Bashir Institute for Infectious Diseases and Biosecurity & Sydney Medical School, The University of Sydney, Westmead Institute for Medical Research | Marie Bashir Institute for Infectious Diseases and Biosecurity & Sydney Medical School, The University of Sydney, Westmead Institute for Medical Research | Eden,J.-S., Kok,J., Dwyer,D.E., Fernandez,M., Carter,I. and Holmes,E.C. |
| EPI_ISL_2595722 | Department of Pediatrics, Center of Excellence in Clinical Virology, Chulalongkorn | Department of Pediatrics, Center of Excellence in Clinical Virology, Chulalongkorn | Thongpan,I. |
| EPI_ISL_2595723 | Gansu Center for Disease Control and Prevention, Pathogen Laboratory | Gansu Center for Disease Control and Prevention, Pathogen Laboratory | Qiao,R., Chen,J., Wu,H. and Yu,D. |
| EPI_ISL_2595724 | Laboratory Medicine, UW Virology | Laboratory Medicine, UW Virology | Lin,M.J., Tait,A. and Greninger,A.L. |
| EPI_ISL_2811729 | Royal Children's Hospital | WHO Collaborating Centre for Reference and Research on Influenza | Jean Moselen, Annette Alafaci, Yi-Mo Deng, Ammar Aziz, Naomi Komadina |
| EPI_ISL_2835616 | PathWest Laboratory Medicine WA Microbial Surveillance Unit | PathWest Laboratory Medicine WA Microbial Surveillance Unit | Chisha Sikazwe, Avram Levy, David Smith, Chris Blyth, Alice Michie, Cara Minney-Smith, David Speers |
| EPI_ISL_2839170, EPI_ISL_2839171, EPI_ISL_2839172, EPI_ISL_2839173, EPI_ISL_2839176, EPI_ISL_2839177 | PathWest Laboratory Medicine WA Microbial Surveillance Unit | PathWest Laboratory Medicine WA Microbial Surveillance Unit | "Chisha Sikazwe, Avram Levy, David Smith, Chris Blyth, Alice Michie, Cara Minney-Smith, David Speers" |
| EPI_ISL_2839186, EPI_ISL_2839187, EPI_ISL_2839188, EPI_ISL_2839189 | Centre for Infectious Diseases and Microbiology Laboratory Services | Centre for Infectious Diseases and Microbiology Laboratory Services | "John-Sebastian Eden, Jen Kok, Dominic Dwyer, Edward Holmes, Philip Britton, Alison Kesson, Elena Cutmore, Rachel Tulloch, Bethany Horsburgh" |
| EPI_ISL_2839190 | Departments of Clinical Microbiology and Infectious Diseases | Centre for Infectious Diseases and Microbiology Laboratory Services | "John-Sebastian Eden, Jen Kok, Dominic Dwyer, Edward Holmes, Philip Britton, Alison Kesson, Elena Cutmore, Rachel Tulloch, Bethany Horsburgh" |
| EPI_ISL_2839196 | Centre for Infectious Diseases and Microbiology Laboratory Services | Centre for Infectious Diseases and Microbiology Laboratory Services | "John-Sebastian Eden, Jen Kok, Dominic Dwyer, Edward Holmes, Philip Britton, Alison Kesson, Elena Cutmore, Rachel Tulloch, Bethany Horsburgh" |
| EPI_ISL_2839197 | Departments of Clinical Microbiology and Infectious Diseases | Centre for Infectious Diseases and Microbiology Laboratory Services | "John-Sebastian Eden, Jen Kok, Dominic Dwyer, Edward Holmes, Philip Britton, Alison Kesson, Elena Cutmore, Rachel Tulloch, Bethany Horsburgh" |
| EPI_ISL_2839198 | Centre for Infectious Diseases and Microbiology Laboratory Services | Centre for Infectious Diseases and Microbiology Laboratory Services | "John-Sebastian Eden, Jen Kok, Dominic Dwyer, Edward Holmes, Philip Britton, Alison Kesson, Elena Cutmore, Rachel Tulloch, Bethany Horsburgh" |
| EPI_ISL_2839199 | PathWest Laboratory Medicine WA Microbial Surveillance Unit | PathWest Laboratory Medicine WA Microbial Surveillance Unit | "Chisha Sikazwe, Avram Levy, David Smith, Chris Blyth, Alice Michie, Cara Minney-Smith, David Speers" |
| EPI_ISL_2839201, EPI_ISL_2839202, EPI_ISL_2839203, EPI_ISL_2839204, EPI_ISL_2839205, EPI_ISL_2839206, EPI_ISL_2839207, EPI_ISL_2839208, EPI_ISL_2839209, EPI_ISL_2839210, EPI_ISL_2839211 | | | |
| see above | Centre for Infectious Diseases and Microbiology Laboratory Services | Centre for Infectious Diseases and Microbiology Laboratory Services | "John-Sebastian Eden, Jen Kok, Dominic Dwyer, Edward Holmes, Philip Britton, Alison Kesson, Elena Cutmore, Rachel Tulloch, Bethany Horsburgh" |
| EPI_ISL_2839213, EPI_ISL_2839214, EPI_ISL_2839215, EPI_ISL_2839216, EPI_ISL_2839217, EPI_ISL_2839218 | Departments of Clinical Microbiology and Infectious Diseases | Centre for Infectious Diseases and Microbiology Laboratory Services | "John-Sebastian Eden, Jen Kok, Dominic Dwyer, Edward Holmes, Philip Britton, Alison Kesson, Elena Cutmore, Rachel Tulloch, Bethany Horsburgh" |
| EPI_ISL_2839219, EPI_ISL_2839220, EPI_ISL_2839221, EPI_ISL_2839222, EPI_ISL_2839223, EPI_ISL_2839224, EPI_ISL_2839225 | Centre for Infectious Diseases and Microbiology Laboratory Services | Centre for Infectious Diseases and Microbiology Laboratory Services | "John-Sebastian Eden, Jen Kok, Dominic Dwyer, Edward Holmes, Philip Britton, Alison Kesson, Elena Cutmore, Rachel Tulloch, Bethany Horsburgh" |
| EPI_ISL_2839227, EPI_ISL_2839228, EPI_ISL_2839229, EPI_ISL_2839230, EPI_ISL_2839231, EPI_ISL_2839232, EPI_ISL_2839233, EPI_ISL_2839234 | Departments of Clinical Microbiology and Infectious Diseases | Centre for Infectious Diseases and Microbiology Laboratory Services | "John-Sebastian Eden, Jen Kok, Dominic Dwyer, Edward Holmes, Philip Britton, Alison Kesson, Elena Cutmore, Rachel Tulloch, Bethany Horsburgh" |
| EPI_ISL_2839235, EPI_ISL_2839236, EPI_ISL_2839237, EPI_ISL_2839239, EPI_ISL_2839242, EPI_ISL_2839244, EPI_ISL_2839246, EPI_ISL_2839248, EPI_ISL_2839250, EPI_ISL_2839252, EPI_ISL_2839254, EPI_ISL_2839256, EPI_ISL_2839258, EPI_ISL_2839260, EPI_ISL_2839262, EPI_ISL_2839264, EPI_ISL_2839266, EPI_ISL_2839269, EPI_ISL_2839271, EPI_ISL_2839273, EPI_ISL_2839275, EPI_ISL_2839277, EPI_ISL_2839279, EPI_ISL_2839281, EPI_ISL_2839283, EPI_ISL_2839285, EPI_ISL_2839288, EPI_ISL_2839290, EPI_ISL_2839292 | | | |
| see above | Centre for Infectious Diseases and Microbiology Laboratory Services | Centre for Infectious Diseases and Microbiology Laboratory Services | "John-Sebastian Eden, Jen Kok, Dominic Dwyer, Edward Holmes, Philip Britton, Alison Kesson, Elena Cutmore, Rachel Tulloch, Bethany Horsburgh" |
| EPI_ISL_2839294, EPI_ISL_2839296, EPI_ISL_2839298, EPI_ISL_2839300, EPI_ISL_2839302, EPI_ISL_2839304, EPI_ISL_2839306, EPI_ISL_2839308, EPI_ISL_2839310, EPI_ISL_2839312 | Departments of Clinical Microbiology and Infectious Diseases | Centre for Infectious Diseases and Microbiology Laboratory Services | "John-Sebastian Eden, Jen Kok, Dominic Dwyer, Edward Holmes, Philip Britton, Alison Kesson, Elena Cutmore, Rachel Tulloch, Bethany Horsburgh" |
| EPI_ISL_2839313, EPI_ISL_2839316, EPI_ISL_2839318, EPI_ISL_2839320, EPI_ISL_2839322, EPI_ISL_2839324, EPI_ISL_2839326, EPI_ISL_2839328, EPI_ISL_2839330, EPI_ISL_2839332, EPI_ISL_2839334, EPI_ISL_2839337, EPI_ISL_2839339, EPI_ISL_2839341, EPI_ISL_2839343, EPI_ISL_2839345, EPI_ISL_2839347, EPI_ISL_2839349 | | | |
| see above | Centre for Infectious Diseases and Microbiology Laboratory Services | Centre for Infectious Diseases and Microbiology Laboratory Services | "John-Sebastian Eden, Jen Kok, Dominic Dwyer, Edward Holmes, Philip Britton, Alison Kesson, Elena Cutmore, Rachel Tulloch, Bethany Horsburgh" |
| EPI_ISL_2839351, EPI_ISL_2839354, EPI_ISL_2839356 | Departments of Clinical Microbiology and Infectious Diseases | Centre for Infectious Diseases and Microbiology Laboratory Services | "John-Sebastian Eden, Jen Kok, Dominic Dwyer, Edward Holmes, Philip Britton, Alison Kesson, Elena Cutmore, Rachel Tulloch, Bethany Horsburgh" |
| EPI_ISL_2839358, EPI_ISL_2839360, EPI_ISL_2839362 | Centre for Infectious Diseases and Microbiology Laboratory Services | Centre for Infectious Diseases and Microbiology Laboratory Services | "John-Sebastian Eden, Jen Kok, Dominic Dwyer, Edward Holmes, Philip Britton, Alison Kesson, Elena Cutmore, Rachel Tulloch, Bethany Horsburgh" |
| EPI_ISL_2839364, EPI_ISL_2839372, EPI_ISL_2839397, EPI_ISL_2839398, EPI_ISL_2839399, EPI_ISL_2839400, EPI_ISL_2839401, EPI_ISL_2839402, EPI_ISL_2839403, EPI_ISL_2839404, EPI_ISL_2839405, EPI_ISL_2839406, EPI_ISL_2839407, EPI_ISL_2839408, EPI_ISL_2839409, EPI_ISL_2839410, EPI_ISL_2839411, EPI_ISL_2839412, EPI_ISL_2839413, EPI_ISL_2839414, EPI_ISL_2839415, EPI_ISL_2839418, EPI_ISL_2839438, EPI_ISL_2839439, EPI_ISL_2839440, EPI_ISL_2839441, EPI_ISL_2839443, EPI_ISL_2839444, EPI_ISL_2839445, EPI_ISL_2839446, EPI_ISL_2839447, EPI_ISL_2839448 | | | |
| see above | PathWest Laboratory Medicine WA Microbial Surveillance Unit | PathWest Laboratory Medicine WA Microbial Surveillance Unit | "Chisha Sikazwe, Avram Levy, David Smith, Chris Blyth, Alice Michie, Cara Minney-Smith, David Speers" |
| EPI_ISL_2989613 | Monash Medical Centre | WHO Collaborating Centre for Reference and Research on Influenza | Xiaomin Dong, Michelle Francis, Tony Korman, Yi-Mo Deng, Ammar Aziz, Naomi Komadina |
| EPI_ISL_2991498 | PathWest Laboratory Medicine WA Microbial Surveillance Unit | PathWest Laboratory Medicine WA Microbial Surveillance Unit | Chisha Sikazwe, Avram Levy, David Smith, Chris Blyth, Alice Michie, Cara Minney-Smith, David Speers |
| EPI_ISL_412458 | Centro de Salud Sócrates Flores Vivas | University of Edinburgh | Gordon, A., Alanis,R., Balmseda,A., Schiller,A., Patel,M., Kubale,J., Tan,G., Pickett,B., Fedorova,N., Amedeo,P., Hu,L., Christensen,J., Miller,J., Durbin,A., Williams,T., Arumemi,F., Cadiz,C., Williams, T. |
| EPI_ISL_412643 | Virology Laboratory Ricardo Gutiérrez Children's Hospital | Virology Laboratory Ricardo Gutiérrez Children's Hospital | Goya, Stephanie.; Nabaes Jodar, Mercedes S.; Valinotto, Laura E.; Rojo, Gabriel L.; Zaiat, Jonathan; Marti, Marcelo A.; Mistchenko, Alicia S.; Viegas, M. |
| EPI_ISL_412857, EPI_ISL_412858 | Virology Laboratory, Ricardo Gutiérrez Children's Hospital | Virology Laboratory, Ricardo Gutiérrez Children's Hospital / Vanderbilt University Medical Center | Goya, Stephanie; Lucion, Maria Florencia; Juarez, Maria del Valle; Shilts, Meghan; Gentile, Angela; Mistchenko, Alicia S.; Das, Suman & Viegas, Mariana |
| EPI_ISL_412859 | WHO National Influenza Centre Russian Federation | WHO National Influenza Centre Russian Federation | Komissarova K.,Fadeev A., Komissarov A., Krivitskaya V. |
| EPI_ISL_412865 | Respiratory Virus Unit, Microbiology Services Colindale, Public Health England | Microbiology Services Colindale, Public Health England | Zambon M |
| EPI_ISL_412866 | Respiratory Virus Unit, Microbiology Services Colindale, Public Health England | Microbiology Services Colindale, Public Health England | Zambon M. |
| EPI_ISL_413222, EPI_ISL_413353 | Institut Pasteur de Madagascar | Institut Pasteur de Madagascar | Jean-Michel HERAUD |
| EPI_ISL_4602779 | VIC, Victorian Infectious Diseases Reference Laboratory | WHO Collaborating Centre for Reference and Research on Influenza | Xiaomin Dong, Annette Alafaci,Yi-Mo Deng, Ammar Aziz, Naomi Komadina |
| EPI_ISL_5522630 | Institut Pasteur de Côte d'Ivoire | WHO Collaborating Centre for Reference and | Xiaomin Dong, Herve Kadjo, Yi-Mo Deng, Ammar Aziz, Naomi Komadina |

| | | | |
|---|---|---|---|
| | | Research on Influenza | |
| EPI_ISL_6174135, EPI_ISL_6174136 | Cote d'ivoire, Institut Pasteur de Cote d'ivoire | WHO Collaborating Centre for Reference and Research on Influenza | Xiaomin Dong, Herve Kadjo, Yi-Mo Deng, Ammar Aziz, Naomi Komadina |
| EPI_ISL_6208719, EPI_ISL_6208724, EPI_ISL_6208725, EPI_ISL_6208726, EPI_ISL_6208727 | Egypt, Central Public Health Laboratory (CPHL) | WHO Collaborating Centre for Reference and Research on Influenza | Xiaomin Dong,Yi-Mo Deng, Amel Naguib, Naomi Komadina |
| EPI_ISL_6268619, EPI_ISL_6268624, EPI_ISL_6314483 | Department of Respiratory Medicine, Wuhan Children's Hospital | CAS Key Laboratory of Special Pathogens and Biosafety, Chinese Academy of Sciences | Decheng Wang, Yi Yan, Ying Li, Xiaoxia Lu, Di Liu |
| EPI_ISL_6494783, EPI_ISL_6494784, EPI_ISL_6494785, EPI_ISL_6494787, EPI_ISL_6494788, EPI_ISL_6494792, EPI_ISL_6494793, EPI_ISL_6494794, EPI_ISL_6494796, EPI_ISL_6494800, EPI_ISL_6494805, EPI_ISL_6494806, EPI_ISL_6494807, EPI_ISL_6494808, EPI_ISL_6494809, EPI_ISL_6494811, EPI_ISL_6494815, EPI_ISL_6494816, EPI_ISL_6494818, EPI_ISL_6494819, EPI_ISL_6494820, EPI_ISL_6494821, EPI_ISL_6494822, EPI_ISL_6494823, EPI_ISL_6494824, EPI_ISL_6494909, EPI_ISL_6494911, EPI_ISL_6494935, EPI_ISL_6494953, EPI_ISL_6494954, EPI_ISL_6494955, EPI_ISL_6494957, EPI_ISL_6494958, EPI_ISL_6494960, EPI_ISL_6494962, EPI_ISL_6494963, EPI_ISL_6494965, EPI_ISL_6494966, EPI_ISL_6494967, EPI_ISL_6494970, EPI_ISL_6494971, EPI_ISL_6494973, EPI_ISL_6494974, EPI_ISL_6494975, EPI_ISL_6494976, EPI_ISL_6494977, EPI_ISL_6494979, EPI_ISL_6494980, EPI_ISL_6494981, EPI_ISL_6494983, EPI_ISL_6494984, EPI_ISL_6494985, EPI_ISL_6494988, EPI_ISL_6494991, EPI_ISL_6494992, EPI_ISL_6494993, EPI_ISL_6494995, EPI_ISL_6494996, EPI_ISL_6494997, EPI_ISL_6494998, EPI_ISL_6495001, EPI_ISL_6495003, EPI_ISL_6495004, EPI_ISL_6495007, EPI_ISL_6495008, EPI_ISL_6495009, EPI_ISL_6495010, EPI_ISL_732337, EPI_ISL_732338, EPI_ISL_732340, EPI_ISL_732341, EPI_ISL_732342, EPI_ISL_732344, EPI_ISL_732345, EPI_ISL_732346, EPI_ISL_732347, EPI_ISL_732348, EPI_ISL_732359, EPI_ISL_732360, EPI_ISL_732361, EPI_ISL_732368, EPI_ISL_732369, EPI_ISL_732372 | | | |
| see above | Respiratory Virus Unit, National Infection Service, Public Health England | National Infection Service, Public Health England | Zambon M, Talts T, Ellis J, Miah S, Platt S |
| EPI_ISL_9003918 | National Institute for Communicable Diseases of the National Health Laboratory Service | National Institute for Communicable Diseases of the National Health Laboratory Service | Amoako DG, Everatt J, Mohale T, Ntuli N, Mahlangu B, Mnguni A, Ismail A, Bhiman JN, Wolter N |
