## Supplementary material for "ARTIC RSV amplicon sequencing reveals global RSV genotype dynamics": GISAID acknowledgement table: gisaid_rsv_acknowledgement_table_2023_12_08_10_B.pdf

We gratefully acknowledge the following Authors from the Originating laboratories responsible for obtaining the specimens, as well as the Submitting laboratories where the genome data were generated and shared via GISAID, on which this research is based.

All Submitters of data may be contacted directly via www.gisaid.org

Authors are sorted alphabetically.

| Accession ID | Originating Laboratory | Submitting Laboratory | Authors |
|---|---|---|---|
| EPI_ISL_1074025, EPI_ISL_1074026, EPI_ISL_1074027, EPI_ISL_1074030, EPI_ISL_1074032, EPI_ISL_1074033, EPI_ISL_1074069, EPI_ISL_1074070, EPI_ISL_1074071, EPI_ISL_1074072, EPI_ISL_1074073, EPI_ISL_1074074, EPI_ISL_1074075, EPI_ISL_1074076, EPI_ISL_1074077, EPI_ISL_1074078, EPI_ISL_1074079, EPI_ISL_1074080, EPI_ISL_1074081, EPI_ISL_1074082, EPI_ISL_1074083, EPI_ISL_1074084, EPI_ISL_1074085, EPI_ISL_1074086, EPI_ISL_1074087, EPI_ISL_1074091, EPI_ISL_1074092, EPI_ISL_1074093, EPI_ISL_1074095, EPI_ISL_1074096, EPI_ISL_1074097, EPI_ISL_1074101, EPI_ISL_1074102, EPI_ISL_1074107, EPI_ISL_1074109, EPI_ISL_1074110, EPI_ISL_1074116, EPI_ISL_1074117, EPI_ISL_1074119, EPI_ISL_1074121, EPI_ISL_1074122, EPI_ISL_1074123, EPI_ISL_1074128, EPI_ISL_1074129, EPI_ISL_1074130, EPI_ISL_1074131, EPI_ISL_1074132, EPI_ISL_1074133, EPI_ISL_1074134, EPI_ISL_1074142, EPI_ISL_1074143, EPI_ISL_1074144, EPI_ISL_1074147, EPI_ISL_1074148, EPI_ISL_1074155, EPI_ISL_1074162, EPI_ISL_1074163, EPI_ISL_1074174, EPI_ISL_1074175, EPI_ISL_1074176, EPI_ISL_1074177, EPI_ISL_1074178, EPI_ISL_1074179, EPI_ISL_1074180, EPI_ISL_1074181, EPI_ISL_1074182, EPI_ISL_1074185, EPI_ISL_1074186, EPI_ISL_1074189, EPI_ISL_1074192, EPI_ISL_1074193, EPI_ISL_1074196, EPI_ISL_1074197, EPI_ISL_1074198, EPI_ISL_1074199, EPI_ISL_1074200, EPI_ISL_1074201, EPI_ISL_1074202, EPI_ISL_1074203, EPI_ISL_1074205, EPI_ISL_1074206, EPI_ISL_1074207, EPI_ISL_1074215, EPI_ISL_1074216, EPI_ISL_1074217, EPI_ISL_1074218, EPI_ISL_1074219, EPI_ISL_1074220, EPI_ISL_1074221, EPI_ISL_1074222, EPI_ISL_1074223, EPI_ISL_1074224, EPI_ISL_1074225, EPI_ISL_1074226, EPI_ISL_1074229, EPI_ISL_1074230, EPI_ISL_1074231, EPI_ISL_1074232, EPI_ISL_1074235, EPI_ISL_1074236, EPI_ISL_1074237, EPI_ISL_1074238, EPI_ISL_1074239, EPI_ISL_1074241, EPI_ISL_1074242, EPI_ISL_1074247, EPI_ISL_1074251, EPI_ISL_1074252, EPI_ISL_1074254, EPI_ISL_1074258, EPI_ISL_1074261, EPI_ISL_1074263, EPI_ISL_1074264, EPI_ISL_1074265, EPI_ISL_1074266, EPI_ISL_1074268, EPI_ISL_1074269, EPI_ISL_1074283, EPI_ISL_1074284, EPI_ISL_1074285, EPI_ISL_1074286, EPI_ISL_1074287, EPI_ISL_1074288, EPI_ISL_1074289, EPI_ISL_1074290, EPI_ISL_1074291, EPI_ISL_1074292, EPI_ISL_1074293, EPI_ISL_1074294, EPI_ISL_1074295, EPI_ISL_1074296, EPI_ISL_1074297, EPI_ISL_1074298, EPI_ISL_1074299, EPI_ISL_1074300 | | | |
| see above | Virology Laboratory, Ricardo Gutiérrez Children's Hospital | Vanderbilt University Medical Center / Virology Laboratory, Ricardo Gutiérrez Children's Hospital | Goya, Stephanie; Lucion, Maria Florencia; Juarez, Maria del Valle; Shilts, Meghan; Gentile, Angela; Mistchenko, Alicia S.; Das, Suman# & Viegas, Mariana# (#contributed equally to this study) |
| EPI_ISL_10914986, EPI_ISL_10914987 | VIC, Victorian Infectious Diseases Reference Laboratory | WHO Collaborating Centre for Reference and Research on Influenza | Xiaomin Dong, Annette Alafaci,Yi-Mo Deng, Naomi Komadina, Ammar Aziz |
| EPI_ISL_10954009 | Respiratory Virus Unit, Specialised Microbiology and Laboratories Directorate, UK Health Security Agency | Respiratory Virus Unit, Specialised Microbiology and Laboratories Directorate, UK Health Security Agency | Zambon M, Talts T, Ellis J, Miah S, Platt S |
| EPI_ISL_11050303 | Phra Nakhon Si Ayutthaya Hospital | National Institute of Health, Department of Medical Sciences, Ministry of Public Health, Thailand | Siripaporn Phuygun; Pakorn Piromtong; Thanutsapa Thanadachakul; Natchaya Khiadsang; Sunthareeya Waicharoen; Malinee Chittaganpitch; Pilailuk Akkapaiboon Okada |
| EPI_ISL_11055769 | Rahima Moosa Mother and child Hospital | National Institute for Communicable Diseases of the National Health Laboratory Service | Amoako DG, Everatt J, Kekana D, Scheepers C, Mohale T, Ntuli N, Mahlangu B, Mnguni A, Ismail A, Bhiman JN, Wolter N |
| EPI_ISL_11055770 | Edendale Gateway clinic | National Institute for Communicable Diseases of the National Health Laboratory Service | Amoako DG, Everatt J, Kekana D, Scheepers C, Mohale T, Ntuli N, Mahlangu B, Mnguni A, Ismail A, Bhiman JN, Wolter N |
| EPI_ISL_11055771 | Rahima Moosa Mother and child Hospital | National Institute for Communicable Diseases of the National Health Laboratory Service | Amoako DG, Everatt J, Kekana D, Scheepers C, Mohale T, Ntuli N, Mahlangu B, Mnguni A, Ismail A, Bhiman JN, Wolter N |
| EPI_ISL_11055772 | Red cross childrens hospital | National Institute for Communicable Diseases of the National Health Laboratory Service | Amoako DG, Everatt J, Kekana D, Scheepers C, Mohale T, Ntuli N, Mahlangu B, Mnguni A, Ismail A, Bhiman JN, Wolter N |
| EPI_ISL_11055778, EPI_ISL_11055783 | Rahima Moosa Mother and child Hospital | National Institute for Communicable Diseases of the National Health Laboratory Service | Amoako DG, Everatt J, Kekana D, Scheepers C, Mohale T, Ntuli N, Mahlangu B, Mnguni A, Ismail A, Bhiman JN, Wolter N |
| EPI_ISL_11055787 | Agincourt clinic | National Institute for Communicable Diseases of the National Health Laboratory Service | Amoako DG, Everatt J, Kekana D, Scheepers C, Mohale T, Ntuli N, Mahlangu B, Mnguni A, Ismail A, Bhiman JN, Wolter N |
| EPI_ISL_11055792, EPI_ISL_11055793, EPI_ISL_11055794 | Red cross childrens hospital | National Institute for Communicable Diseases of the National Health Laboratory Service | Amoako DG, Everatt J, Kekana D, Scheepers C, Mohale T, Ntuli N, Mahlangu B, Mnguni A, Ismail A, Bhiman JN, Wolter N |
| EPI_ISL_11055795 | Klerksdorp | National Institute for Communicable Diseases of the National Health Laboratory Service | Amoako DG, Everatt J, Kekana D, Scheepers C, Mohale T, Ntuli N, Mahlangu B, Mnguni A, Ismail A, Bhiman JN, Wolter N |
| EPI_ISL_11055796 | Jouberton clinic | National Institute for Communicable Diseases of the National Health Laboratory Service | Amoako DG, Everatt J, Kekana D, Scheepers C, Mohale T, Ntuli N, Mahlangu B, Mnguni A, Ismail A, Bhiman JN, Wolter N |
| EPI_ISL_11055797, EPI_ISL_11055798, EPI_ISL_11055801, EPI_ISL_11055802 | Red cross childrens hospital | National Institute for Communicable Diseases of the National Health Laboratory Service | Amoako DG, Everatt J, Kekana D, Scheepers C, Mohale T, Ntuli N, Mahlangu B, Mnguni A, Ismail A, Bhiman JN, Wolter N |
| EPI_ISL_11055803 | Edendale hospital | National Institute for Communicable Diseases of the National Health Laboratory Service | Amoako DG, Everatt J, Kekana D, Scheepers C, Mohale T, Ntuli N, Mahlangu B, Mnguni A, Ismail A, Bhiman JN, Wolter N |
| EPI_ISL_11055804 | Mitchell's plain distric hospital | National Institute for Communicable Diseases of the National Health Laboratory Service | Amoako DG, Everatt J, Kekana D, Scheepers C, Mohale T, Ntuli N, Mahlangu B, Mnguni A, Ismail A, Bhiman JN, Wolter N |
| EPI_ISL_11055805 | Klerksdorp | National Institute for Communicable Diseases of the National Health Laboratory Service | Amoako DG, Everatt J, Kekana D, Scheepers C, Mohale T, Ntuli N, Mahlangu B, Mnguni A, Ismail A, Bhiman JN, Wolter N |
| EPI_ISL_11428307, EPI_ISL_11428308, EPI_ISL_11428310, EPI_ISL_11428311, EPI_ISL_11428312, EPI_ISL_11428313, EPI_ISL_11428314 | Respiratory Virus Unit, Reference Services, UK Health Security Agency | Reference Services, UK Health Security Agency | Zambon M, Talts T, Ellis J, Miah S, Platt S |
| EPI_ISL_11817018, EPI_ISL_11817022, EPI_ISL_11817029, EPI_ISL_11817031, EPI_ISL_11817033, EPI_ISL_11817034, EPI_ISL_11817035, EPI_ISL_11817037, EPI_ISL_11817040, EPI_ISL_11817044, EPI_ISL_11817047, EPI_ISL_11817048, EPI_ISL_11817050, EPI_ISL_11817052, EPI_ISL_11817054, EPI_ISL_11817055, EPI_ISL_11817057, EPI_ISL_11817058, EPI_ISL_11817059, EPI_ISL_11817060, EPI_ISL_11817063, EPI_ISL_11817064, EPI_ISL_11817067, EPI_ISL_11817068, EPI_ISL_11817073, EPI_ISL_11817074, EPI_ISL_11817077, EPI_ISL_11817079, EPI_ISL_11817081, EPI_ISL_11817084, EPI_ISL_11817085 | | | |
| see above | QLD, Royal Brisbane and Woman's Hospital | WHO Collaborating Centre for Reference and Research on Influenza | Xiaomin Dong, Yi-Mo Deng, Ammar Aziz, Naomi Komadina |
| EPI_ISL_11817087, EPI_ISL_11817089, EPI_ISL_11817093, EPI_ISL_11817098 | VIC, Royal Children's Hospital | WHO Collaborating Centre for Reference and Research on Influenza | Xiaomin Dong, Annette Alafaci,Yi-Mo Deng, Ammar Aziz, Naomi Komadina |
| EPI_ISL_11817110, EPI_ISL_11817111, EPI_ISL_11817112, EPI_ISL_11817113, EPI_ISL_11817114, EPI_ISL_11817115, EPI_ISL_11817116, EPI_ISL_11817117, EPI_ISL_11817118, EPI_ISL_11817119, EPI_ISL_11817120, EPI_ISL_11817121, EPI_ISL_11817122, EPI_ISL_11817123, EPI_ISL_11817124, EPI_ISL_11817125, EPI_ISL_11817126, EPI_ISL_11817127, EPI_ISL_11817128, EPI_ISL_11817129, EPI_ISL_11817130, EPI_ISL_11817131, EPI_ISL_11817132, EPI_ISL_11817133, EPI_ISL_11817134, EPI_ISL_11817135, EPI_ISL_11817136, EPI_ISL_11817137, EPI_ISL_11817138, EPI_ISL_11817139 | | | |
| see above | VIC, Monash Medical Centre | WHO Collaborating Centre for Reference and Research on Influenza | Xiaomin Dong, Michelle Francis, Tony Korman, Yi-Mo Deng, Ammar Aziz, Naomi Komadina |
| EPI_ISL_12529654 | Klerksdorp | National Institute for Communicable Diseases of the National Health Laboratory Service | Amoako DG, Everatt J, Kekana D, Scheepers C, Mohale T, Ntuli N, Mahlangu B, Mnguni A, Ismail A, Bhiman JN, Wolter N |
| EPI_ISL_12529655 | Rahima Moosa Mother and child Hospital | National Institute for Communicable Diseases of the National Health Laboratory Service | Amoako DG, Everatt J, Kekana D, Scheepers C, Mohale T, Ntuli N, Mahlangu B, Mnguni A, Ismail A, Bhiman JN, Wolter N |
| EPI_ISL_12529656 | Edendale hospital | National Institute for Communicable Diseases of the National Health Laboratory Service | Amoako DG, Everatt J, Kekana D, Scheepers C, Mohale T, Ntuli N, Mahlangu B, Mnguni A, Ismail A, Bhiman JN, Wolter N |
| EPI_ISL_12529657 | Klerksdorp | National Institute for Communicable Diseases of the National Health Laboratory Service | Amoako DG, Everatt J, Kekana D, Scheepers C, Mohale T, Ntuli N, Mahlangu B, Mnguni A, Ismail A, Bhiman JN, Wolter N |
| EPI_ISL_12529659 | Red cross childrens hospital | National Institute for Communicable Diseases of the National Health Laboratory Service | Amoako DG, Everatt J, Kekana D, Scheepers C, Mohale T, Ntuli N, Mahlangu B, Mnguni A, Ismail A, Bhiman JN, Wolter N |
| EPI_ISL_12529660 | Edendale hospital | National Institute for Communicable Diseases of the National Health Laboratory Service | Amoako DG, Everatt J, Kekana D, Scheepers C, Mohale T, Ntuli N, Mahlangu B, Mnguni A, Ismail A, Bhiman JN, Wolter N |
| EPI_ISL_12544916, EPI_ISL_12544917 | National Influenza Center, National Institute of Hygiene and Epidemiology (NIHE) | National Institute of Hygiene and Epidemiology (NIHE) National Influenza Centre, Virology Department | Ung Thi Hong Trang, Hoang Vu Mai Phuong, Nguyen Huy Hoang, Nguyen Le Khanh Hang, Le Thi Thanh, Nguyen Vu Son, Vuong Duc Cuong, Pham Thi Hien, Tran Thu Huong, Nguyen Co Thach, Nguyen Phuong Anh, Le Quynh Mai |
| EPI_ISL_12970400, EPI_ISL_12970402, EPI_ISL_12970404, EPI_ISL_12970406, EPI_ISL_12970410, EPI_ISL_12970424 | Department of Virology, Research Institute for Tropical Medicine | RVDT/RVD/DVD/CDC | Lijuan Wang, Mayan U. Lumandas, Vina Lea Arguelles, Jonjee Morin, Roman Tatusov and Everardo Vega |
| EPI_ISL_13231399, EPI_ISL_13231406, EPI_ISL_13231411, EPI_ISL_13231414, EPI_ISL_13231416, EPI_ISL_13231420, EPI_ISL_13231421, EPI_ISL_13231423, EPI_ISL_13231424, EPI_ISL_13231426, EPI_ISL_13231427, EPI_ISL_13231430, EPI_ISL_13231431, EPI_ISL_13231432, EPI_ISL_13231433, EPI_ISL_13231437, EPI_ISL_13231438, EPI_ISL_13231440 | | | |
| see above | Israel Central Virology laboratory, Ministry of Health | Israel Central Virology laboratory, Ministry of Health | Neta S. Zuckerman, Efrat Bucris, Miranda Geva, Hagar Morad, Or Zilbertzan, Ilana S. Fratty, Ital Nemet, Nofar Atari, Limor Kliker, Ella Mendelson, Michal Mandelboim |
| EPI_ISL_14084067, EPI_ISL_14084068, EPI_ISL_14084070, EPI_ISL_14084072, EPI_ISL_14084073, EPI_ISL_14084074, EPI_ISL_14084075, EPI_ISL_14084076, EPI_ISL_14084077, EPI_ISL_14084078, EPI_ISL_14084079, EPI_ISL_14084080, EPI_ISL_14084081, EPI_ISL_14084082, EPI_ISL_14084083, EPI_ISL_14084084, EPI_ISL_14084085, EPI_ISL_14084086, EPI_ISL_14084087 | | | |
| see above | Complexo Hospitalario Universitario de Vigo. Microbiology Department | Complexo Hospitalario Universitario de Vigo. Microbiology Department | Daviña, Carlos; Pizcueta, Juan; Perez, Sonia |
| EPI_ISL_14769847 | Tintswalo | National Institute for Communicable Diseases of the National Health Laboratory Service | Amoako DG, Everatt J, Kekana D, Mahlangu B, Stock N, Ntuli N, Mnguni A, Motsatsi G, Nzimande A, Ismail A, Bhiman JN, Wolter N |
| EPI_ISL_14769849, EPI_ISL_14769851, EPI_ISL_14769854, EPI_ISL_14769855, EPI_ISL_14769856 | Red Cross | National Institute for Communicable Diseases of the National Health Laboratory Service | Amoako DG, Everatt J, Kekana D, Mahlangu B, Stock N, Ntuli N, Mnguni A, Motsatsi G, Nzimande A, Ismail A, Bhiman JN, Wolter N |

| | | | |
|---|---|---|---|
| EPI_ISL_14769858 | Mitchells Plain | National Institute for Communicable Diseases of the National Health Laboratory Service | Amoako DG, Everatt J, Kekana D, Mahlangu B, Stock N, Ntuli N, Mnguni A, Motsatsi G, Nzimande A, Ismail A, Bhiman JN, Wolter N |
| EPI_ISL_14769860 | Rahima Moosa | National Institute for Communicable Diseases of the National Health Laboratory Service | Amoako DG, Everatt J, Kekana D, Mahlangu B, Stock N, Ntuli N, Mnguni A, Motsatsi G, Nzimande A, Ismail A, Bhiman JN, Wolter N |
| EPI_ISL_15004431, EPI_ISL_15004432, EPI_ISL_15004434, EPI_ISL_15004436, EPI_ISL_15004437 | National Institute of Hygiene, virology departement, National Influenza Center | Centers for Disease Control and Prevention, Coronavirus and Other Respiratory Viruses Division | Megha Aggarwal, Lijuan Wang, Ji In Park, Amanda Smith and Everardo Vega |
| EPI_ISL_15055329, EPI_ISL_15055330, EPI_ISL_15055331, EPI_ISL_15055332, EPI_ISL_15055333, EPI_ISL_15055334, EPI_ISL_15055335, EPI_ISL_15055336, EPI_ISL_15055337, EPI_ISL_15055338, EPI_ISL_15055339, EPI_ISL_15055340, EPI_ISL_15055341, EPI_ISL_15055342, EPI_ISL_15055343, EPI_ISL_15055344, EPI_ISL_15055345, EPI_ISL_15055346, EPI_ISL_15055347, EPI_ISL_15055348, EPI_ISL_15055349, EPI_ISL_15055350, EPI_ISL_15055351, EPI_ISL_15055352, EPI_ISL_15055353, EPI_ISL_15055354, EPI_ISL_15055355, EPI_ISL_15055356, EPI_ISL_15055357, EPI_ISL_15055358, EPI_ISL_15067694, EPI_ISL_15067695, EPI_ISL_15067696, EPI_ISL_15067697, EPI_ISL_15067698, EPI_ISL_15067699, EPI_ISL_15067700, EPI_ISL_15067701, EPI_ISL_15067702, EPI_ISL_15067703, EPI_ISL_15067704, EPI_ISL_15067705, EPI_ISL_15067706, EPI_ISL_15067707, EPI_ISL_15067708, EPI_ISL_15067709, EPI_ISL_15067710, EPI_ISL_15067711, EPI_ISL_15067712, EPI_ISL_15067713, EPI_ISL_15067714, EPI_ISL_15067715, EPI_ISL_15067716, EPI_ISL_15067717, EPI_ISL_15067718, EPI_ISL_15067719, EPI_ISL_15067720, EPI_ISL_15067721, EPI_ISL_15067722, EPI_ISL_15067723 | | | |
| see above | Virology Laboratory, Ricardo Gutiérrez Children's Hospital | Virology Laboratory, Ricardo Gutiérrez Children's Hospital | Acuña, Dolores; Goya, Stephanie; Nabaes Jodar, Mercedes S.; Lucion, M. Florencia; Juárez, María del Valle; Gentile, Ángela; Mistchenko, Alicia S.; Viegas, Mariana |
| EPI_ISL_15120684, EPI_ISL_15120685, EPI_ISL_15120686, EPI_ISL_15120687, EPI_ISL_15120691, EPI_ISL_15120693, EPI_ISL_15120695, EPI_ISL_15120696, EPI_ISL_15120697, EPI_ISL_15120698, EPI_ISL_15120699, EPI_ISL_15120700, EPI_ISL_15120701, EPI_ISL_15120702, EPI_ISL_15120704, EPI_ISL_15120705, EPI_ISL_15120707, EPI_ISL_15120708, EPI_ISL_15120709, EPI_ISL_15120710, EPI_ISL_15120711, EPI_ISL_15120712, EPI_ISL_15120714, EPI_ISL_15120720, EPI_ISL_15120722, EPI_ISL_15120724, EPI_ISL_15120725, EPI_ISL_15120732, EPI_ISL_15120733, EPI_ISL_15120734, EPI_ISL_15120735, EPI_ISL_15120736, EPI_ISL_15120737, EPI_ISL_15120740, EPI_ISL_15120741, EPI_ISL_15120742, EPI_ISL_15120744, EPI_ISL_15120746, EPI_ISL_15120747, EPI_ISL_15120748, EPI_ISL_15120749, EPI_ISL_15120750, EPI_ISL_15120751, EPI_ISL_15120752, EPI_ISL_15120753, EPI_ISL_15120754, EPI_ISL_15120755, EPI_ISL_15120756, EPI_ISL_15120757, EPI_ISL_15120758, EPI_ISL_15120759, EPI_ISL_15120760, EPI_ISL_15120761, EPI_ISL_15120762, EPI_ISL_15120763, EPI_ISL_15120764, EPI_ISL_15120765, EPI_ISL_15120766, EPI_ISL_15120767, EPI_ISL_15120768, EPI_ISL_15120769, EPI_ISL_15120779, EPI_ISL_15120784, EPI_ISL_15120787, EPI_ISL_15120788, EPI_ISL_15120789, EPI_ISL_15120790, EPI_ISL_15120791, EPI_ISL_15120792, EPI_ISL_15120793, EPI_ISL_15120794, EPI_ISL_15120795, EPI_ISL_15120796, EPI_ISL_15120797, EPI_ISL_15120798, EPI_ISL_15120799 | | | |
| see above | National Institute of Hygiene, virology departement, National Influenza Center | Centers for Disease Control and Prevention, Coronavirus and Other Respiratory Viruses Division | Megha Aggarwal, Lijuan Wang, Ji In Park, Amanda Smith and Everardo Vega |
| EPI_ISL_15421405 | Department of Infectious Diseases in Humans, Viral Diseases Laboratory, Sciensano | Department of Infectious Diseases in Humans, Viral Diseases Laboratory, Sciensano | François Dufrasne, Sarah Denayer, Steven Van Gucht, Cyril Barbezange |
| EPI_ISL_15601574 | Laboratory of Virology - Institute of Public Health, Republic of Macedonia | Laboratory of Virology - Institute of Public Health, Republic of Macedonia | Teodora Buzharova, Golubinka Boshevska, Elizabeta Jancheska |
| EPI_ISL_15728619 | Red Cross | National Institute for Communicable Diseases of the National Health Laboratory Service | Everatt J, Kekana D, Mahlangu B, Stock N, Ntozini B, Ntuli N, Mnguni A, Nzimande A, Ismail A, Bhiman JN, Wolter N |
| EPI_ISL_15728620, EPI_ISL_15728621 | Rahima Moosa | National Institute for Communicable Diseases of the National Health Laboratory Service | Everatt J, Kekana D, Mahlangu B, Stock N, Ntozini B, Ntuli N, Mnguni A, Nzimande A, Ismail A, Bhiman JN, Wolter N |
| EPI_ISL_15728622, EPI_ISL_15728624 | Red Cross | National Institute for Communicable Diseases of the National Health Laboratory Service | Everatt J, Kekana D, Mahlangu B, Stock N, Ntozini B, Ntuli N, Mnguni A, Nzimande A, Ismail A, Bhiman JN, Wolter N |
| EPI_ISL_15728625 | Matikwane | National Institute for Communicable Diseases of the National Health Laboratory Service | Everatt J, Kekana D, Mahlangu B, Stock N, Ntozini B, Ntuli N, Mnguni A, Nzimande A, Ismail A, Bhiman JN, Wolter N |
| EPI_ISL_15728627, EPI_ISL_15728628, EPI_ISL_15728629 | Red Cross | National Institute for Communicable Diseases of the National Health Laboratory Service | Everatt J, Kekana D, Mahlangu B, Stock N, Ntozini B, Ntuli N, Mnguni A, Nzimande A, Ismail A, Bhiman JN, Wolter N |
| EPI_ISL_15728631 | Livingstone Hospital | National Institute for Communicable Diseases of the National Health Laboratory Service | Everatt J, Kekana D, Mahlangu B, Stock N, Ntozini B, Ntuli N, Mnguni A, Nzimande A, Ismail A, Bhiman JN, Wolter N |
| EPI_ISL_15728632, EPI_ISL_15728633 | Red Cross | National Institute for Communicable Diseases of the National Health Laboratory Service | Everatt J, Kekana D, Mahlangu B, Stock N, Ntozini B, Ntuli N, Mnguni A, Nzimande A, Ismail A, Bhiman JN, Wolter N |
| EPI_ISL_15728634 | Mitchells Plain | National Institute for Communicable Diseases of the National Health Laboratory Service | Everatt J, Kekana D, Mahlangu B, Stock N, Ntozini B, Ntuli N, Mnguni A, Nzimande A, Ismail A, Bhiman JN, Wolter N |
| EPI_ISL_15750192, EPI_ISL_15750193, EPI_ISL_15750194, EPI_ISL_15750195 | Inserm UMR1137, Universite de Paris | Inserm UMR1137, Universite de Paris | Coppee,R., Chenane,H.R., Bridier-Nahmias,A., Tcherakian,C., Catherinot,E., Collin,G., Lebourgeois,S., Visseaux,B., Descamps,D., Vasse,M., Farfour,E., Coppee,R., Chenane,H.R., Bridier-Nahmias,A., Collin,G., Lebourgeois,S., Visseaux,B., Descamps,D. |
| EPI_ISL_15750833, EPI_ISL_15750834, EPI_ISL_15750835, EPI_ISL_15750836, EPI_ISL_15750837, EPI_ISL_15750838, EPI_ISL_15750839 | State Key Laboratory of Pathogen and Biosecurity, Beijing Institute of Microbiology and Epidemiology | State Key Laboratory of Pathogen and Biosecurity, Beijing Institute of Microbiology and Epidemiology | Jia,N., Cao,W.-C., Hu,Y.-L., Ye,R.-Z., Que,T.-C., Xia,L.-Y., Cui,X.-M., Zhang,Y.-W., Jiang,J.-F., Wang,Q.-H., Wang,Q., Jia,N., Cao,W.-C., Hu,Y.-L., Ye,R.-Z., Que,T.-C., Xia,L.-Y., Cui,X.-M., Zhang,Y.-W., Jiang,J.-F., Wang,Q.-H., Wang,Q. |
| EPI_ISL_15751422 | Pathology, Stanford University School of Medicine | Pathology, Stanford University School of Medicine | Huang,C., Doan,T., Sahoo,M.K., Pinsky,B.A. |
| EPI_ISL_15751943, EPI_ISL_15751944, EPI_ISL_15751945, EPI_ISL_15751946, EPI_ISL_15751947, EPI_ISL_15751948, EPI_ISL_15751949, EPI_ISL_15751950, EPI_ISL_15751951, EPI_ISL_15751952, EPI_ISL_15751953, EPI_ISL_15751954, EPI_ISL_15751955, EPI_ISL_15751956, EPI_ISL_15751957, EPI_ISL_15751958, EPI_ISL_15751959, EPI_ISL_15751960, EPI_ISL_15751961, EPI_ISL_15751962, EPI_ISL_15751963, EPI_ISL_15751964, EPI_ISL_15751965, EPI_ISL_15751966, EPI_ISL_15751967, EPI_ISL_15751968, EPI_ISL_15751969, EPI_ISL_15751970, EPI_ISL_15751971, EPI_ISL_15751972, EPI_ISL_15751973, EPI_ISL_15751974, EPI_ISL_15751975, EPI_ISL_15751976, EPI_ISL_15751977, EPI_ISL_15751978, EPI_ISL_15751979, EPI_ISL_15751980, EPI_ISL_15751981, EPI_ISL_15751982, EPI_ISL_15751983, EPI_ISL_15751984, EPI_ISL_15751985, EPI_ISL_15751986, EPI_ISL_15751987, EPI_ISL_15751988, EPI_ISL_15751989, EPI_ISL_15751990, EPI_ISL_15751991, EPI_ISL_15751992, EPI_ISL_15751993, EPI_ISL_15751994, EPI_ISL_15751995, EPI_ISL_15751996, EPI_ISL_15751997, EPI_ISL_15751998, EPI_ISL_15751999, EPI_ISL_15752000, EPI_ISL_15752001, EPI_ISL_15752002, EPI_ISL_15752003, EPI_ISL_15752004 | | | |
| see above | Respiratory Viruses Branch, Division of Viral Diseases | Respiratory Viruses Branch, Division of Viral Diseases | Wang,L., Ng,T.F.F., Castro,C.J., Marine,R.L., Magana,L.C., Esona,M., Peret,T.C.T., Thornburg,N.J., Wang,L., Ng,T.F.F., Castro,C.J., Marine,R.L., Magana,L.C., Esona,M., Peret,T.C.T., Thornburg,N.J. |
| EPI_ISL_15752977, EPI_ISL_15752978, EPI_ISL_15752979, EPI_ISL_15752980, EPI_ISL_15752981, EPI_ISL_15752982, EPI_ISL_15752984, EPI_ISL_15752985, EPI_ISL_15752986, EPI_ISL_15752987, EPI_ISL_15752988, EPI_ISL_15752990, EPI_ISL_15752992, EPI_ISL_15752993, EPI_ISL_15752994, EPI_ISL_15752995, EPI_ISL_15752996, EPI_ISL_15752997, EPI_ISL_15752998, EPI_ISL_15752999, EPI_ISL_15753000, EPI_ISL_15753001, EPI_ISL_15753002, EPI_ISL_15753003, EPI_ISL_15753005, EPI_ISL_15753006, EPI_ISL_15753007, EPI_ISL_15753008, EPI_ISL_15753009, EPI_ISL_15753010, EPI_ISL_15753011, EPI_ISL_15753012, EPI_ISL_15753013, EPI_ISL_15753014, EPI_ISL_15753015, EPI_ISL_15753016, EPI_ISL_15753017, EPI_ISL_15753018, EPI_ISL_15753019, EPI_ISL_15753020, EPI_ISL_15753021, EPI_ISL_15753022, EPI_ISL_15753023, EPI_ISL_15753024, EPI_ISL_15753025, EPI_ISL_15753026, EPI_ISL_15753027, EPI_ISL_15753028, EPI_ISL_15753029, EPI_ISL_15753030, EPI_ISL_15753032, EPI_ISL_15753035, EPI_ISL_15753036, EPI_ISL_15753037, EPI_ISL_15753038, EPI_ISL_15753039, EPI_ISL_15753040, EPI_ISL_15753041, EPI_ISL_15753043, EPI_ISL_15753044, EPI_ISL_15753045, EPI_ISL_15753046, EPI_ISL_15753048, EPI_ISL_15753049, EPI_ISL_15753050, EPI_ISL_15753051, EPI_ISL_15753052, EPI_ISL_15753053, EPI_ISL_15753054, EPI_ISL_15753055, EPI_ISL_15753056, EPI_ISL_15753058, EPI_ISL_15753059, EPI_ISL_15753060, EPI_ISL_15753061, EPI_ISL_15753062, EPI_ISL_15753064, EPI_ISL_15753066, EPI_ISL_15753067, EPI_ISL_15753068, EPI_ISL_15753069, EPI_ISL_15753071, EPI_ISL_15753072, EPI_ISL_15753073, EPI_ISL_15753074, EPI_ISL_15753075, EPI_ISL_15753076, EPI_ISL_15753077, EPI_ISL_15753078, EPI_ISL_15753079, EPI_ISL_15753080, EPI_ISL_15753082, EPI_ISL_15753083, EPI_ISL_15753084, EPI_ISL_15753085, EPI_ISL_15753086, EPI_ISL_15753087, EPI_ISL_15753088, EPI_ISL_15753089, EPI_ISL_15753090, EPI_ISL_15753091, EPI_ISL_15753092, EPI_ISL_15753093, EPI_ISL_15753094, EPI_ISL_15753096, EPI_ISL_15753097, EPI_ISL_15753098, EPI_ISL_15753099, EPI_ISL_15753100, EPI_ISL_15753101, EPI_ISL_15753103, EPI_ISL_15753104, EPI_ISL_15753105, EPI_ISL_15753106, EPI_ISL_15753107, EPI_ISL_15753109, EPI_ISL_15753110, EPI_ISL_15753111, EPI_ISL_15753112, EPI_ISL_15753114, EPI_ISL_15753115, EPI_ISL_15753116, EPI_ISL_15753117, EPI_ISL_15753118, EPI_ISL_15753119, EPI_ISL_15753121, EPI_ISL_15753122, EPI_ISL_15753124, EPI_ISL_15753125, EPI_ISL_15753126, EPI_ISL_15753127, EPI_ISL_15753128, EPI_ISL_15753129, EPI_ISL_15753132, EPI_ISL_15753133, EPI_ISL_15753134, EPI_ISL_15753135, EPI_ISL_15753137, EPI_ISL_15753138, EPI_ISL_15753139, EPI_ISL_15753140, EPI_ISL_15753141, EPI_ISL_15753142, EPI_ISL_15753144, EPI_ISL_15753145, EPI_ISL_15753146, EPI_ISL_15753147, EPI_ISL_15753148, EPI_ISL_15753150, EPI_ISL_15753151, EPI_ISL_15753152, EPI_ISL_15753153, EPI_ISL_15753154, EPI_ISL_15753155, EPI_ISL_15753156, EPI_ISL_15753158, EPI_ISL_15753159, EPI_ISL_15753161, EPI_ISL_15753162, EPI_ISL_15753163, EPI_ISL_15753164, EPI_ISL_15753165, EPI_ISL_15753166, EPI_ISL_15753167, EPI_ISL_15753168, EPI_ISL_15753170, EPI_ISL_15753171, EPI_ISL_15753172, EPI_ISL_15753174, EPI_ISL_15753175, EPI_ISL_15753177, EPI_ISL_15753178, EPI_ISL_15753179, EPI_ISL_15753180, EPI_ISL_15753181, EPI_ISL_15753182, EPI_ISL_15753184, EPI_ISL_15753185, EPI_ISL_15753186, EPI_ISL_15753187, EPI_ISL_15753188, EPI_ISL_15753189, EPI_ISL_15753190, EPI_ISL_15753192, EPI_ISL_15753193, EPI_ISL_15753194, EPI_ISL_15753195, EPI_ISL_15753196, EPI_ISL_15753197, EPI_ISL_15753198, EPI_ISL_15753199, EPI_ISL_15753201, EPI_ISL_15753202, EPI_ISL_15753203, EPI_ISL_15753204, EPI_ISL_15753205, EPI_ISL_15753206, EPI_ISL_15753207, EPI_ISL_15753208, EPI_ISL_15753209, EPI_ISL_15753210, EPI_ISL_15753211, EPI_ISL_15753212, EPI_ISL_15753213, EPI_ISL_15753214, EPI_ISL_15753215, EPI_ISL_15753217, EPI_ISL_15753219, EPI_ISL_15753220, EPI_ISL_15753221, EPI_ISL_15753223, EPI_ISL_15753224, EPI_ISL_15753225, EPI_ISL_15753226, EPI_ISL_15753228, EPI_ISL_15753229, EPI_ISL_15753230, EPI_ISL_15753231, EPI_ISL_15753232, EPI_ISL_15753233, EPI_ISL_15753234, EPI_ISL_15753236, EPI_ISL_15753237, EPI_ISL_15753238, EPI_ISL_15753239, EPI_ISL_15753240, EPI_ISL_15753241, EPI_ISL_15753242, EPI_ISL_15753243, EPI_ISL_15753244, EPI_ISL_15753245, EPI_ISL_15753246, EPI_ISL_15753247, EPI_ISL_15753248, EPI_ISL_15753249, EPI_ISL_15753250, EPI_ISL_15753251 | | | |
| see above | Department of Paediatrics, University of Oxford | Department of Paediatrics, University of Oxford | Lin,G.-L., Drysdale,S.B., Snape,M.D., O'Connor,D., Brown,A., MacIntyre-Cockett,G., Mellado-Gomez,E., de Cesare,M., Bonsall,D., Ansari,M.A., Oner,D., Aerssens,J., Butler,C., Bont,L., Openshaw,P., Martinon-Torres,F., Nair,H., Bowden,R., Golubchik,T., Pollard,A.J., Lin,G.-L. |
| EPI_ISL_15753573 | Eurasian Institute of zoonotic infections, The Federal Research Center of Fundamental and Translational Medicine | Eurasian Institute of zoonotic infections, The Federal Research Center of Fundamental and Translational Medicine | Dubovitskiy,N.A., Sobolev,I.A., Kurskaya,O.G., Simkina,O.A., Komissarova,T.V., Murashkina,T.A., Solomatina,M.V., Derko,A.A., Saroyan,T.A., Kabilov,M.R., Tupikin,A.E., Sharshov,K.A., Shestopalov,A.M. |
| EPI_ISL_15753578, EPI_ISL_15753579, EPI_ISL_15753580 | Eurasian Institute of zoonotic infections, The Federal Research Center of Fundamental and Translational Medicine | Eurasian Institute of zoonotic infections, The Federal Research Center of Fundamental and Translational Medicine | Dubovitskiy,N.A., Sobolev,I.A., Kurskaya,O.G., Anoshina,A.V., Leonova,N.V., Murashkina,T.A., Solomatina,M.V., Derko,A.A., Saroyan,T.A., Kabilov,M.R., Tupikin,A.E., Sharshov,K.A., Shestopalov,A.M. |
| EPI_ISL_15771617, EPI_ISL_15771618, EPI_ISL_15771621, EPI_ISL_15771622, EPI_ISL_15771625, EPI_ISL_15771626, EPI_ISL_15771628, EPI_ISL_15771630 | Respiratory Viruses Branch, Division of Viral Diseases, Centers for Disease Control and Prevention | Respiratory Viruses Branch, Division of Viral Diseases, Centers for Disease Control and Prevention | Wang,L., Ng,T.F.F., Castro,C.J., Marine,R.L., Magana,L.C., Esona,M., Peret,T.C.T. and Thornburg,N.J. |
| EPI_ISL_15772196 | Laboratory Medicine, UW Virology | Laboratory Medicine, UW Virology | Sereewit,J., Xie,H. and Greninger,A. |
| EPI_ISL_15773988, EPI_ISL_15773989, EPI_ISL_15774005, EPI_ISL_15774019, EPI_ISL_15774028, EPI_ISL_15774034, EPI_ISL_15774036, EPI_ISL_15774037, EPI_ISL_15774040, EPI_ISL_15774041, EPI_ISL_15774044, EPI_ISL_15774048, EPI_ISL_15774051, EPI_ISL_15774052, EPI_ISL_15774055, EPI_ISL_15774057, EPI_ISL_15774061, EPI_ISL_15774069 | | | |
| see above | QLD, Royal Brisbane and Woman's Hospital | WHO Collaborating Centre for Reference and Research on Influenza | Xiaomin Dong, Yi-Mo Deng, Ammar Aziz, Naomi Komadina |
| EPI_ISL_15820265 | Respiratory Viruses Branch, Division of Viral Diseases | Respiratory Viruses Branch, Division of Viral Diseases | Wang,L., Ng,T.F.F., Castro,C.J., Marine,R.L., Magana,L.C., Esona,M., Peret,T.C.T., Thornburg,N.J., Wang,L., Ng,T.F.F., Castro,C.J., Marine,R.L., Magana,L.C., Esona,M., Peret,T.C.T., Thornburg,N.J. |
| EPI_ISL_15895059 | Laboratorio Central de Saude Publica do Estado do Rio Grande do Sul (LACEN/RS) | Laboratory of Respiratory Viruses and Measles, Oswaldo Cruz Institute, FIOCRUZ | Paola Resende, Fernando Motta, Elisa Cavalcante Pereira, Bruna Mendonça da Silva, Jéssica Graça Macedo de Carvalho, Larissa Macedo Pinto, Victor Guimaraes, Ighor Leonardo Arantes, Leticia Scalioni, Anderson Brandao Leite, Marilda Siqueira on behalf of the Fiocruz COVID-19 Genomic Surveillance Network |
| EPI_ISL_15895095 | Laboratorio Central de Saude Publica do Estado do Espirito Santo (LACEN/ES) | Laboratory of Respiratory Viruses and Measles, Oswaldo Cruz Institute, FIOCRUZ | Paola Resende, Fernando Motta, Elisa Cavalcante Pereira, Ighor Arantes, Bruna Mendonça da Silva, Jéssica Graça Macedo de Carvalho, Larissa Macedo Pinto, Victor Guimaraes, Leticia Scalioni, Rodrigo Ribeiro Rodrigues, Marilda Siqueira on behalf of the Fiocruz COVID-19 Genomic Surveillance Network |
| EPI_ISL_15895128 | Universidade Federal de São Paulo, Departamento de Medicina, Disciplina de Doenças Infecciosas e Parasitárias. Laboratório de Virologia | Laboratory of Respiratory Viruses and Measles, Oswaldo Cruz Institute, FIOCRUZ | Paola Resende, Fernando Motta, Elisa Cavalcante Pereira, Ighor Arantes, Bruna Mendonça da Silva, Jéssica Graça Macedo de Carvalho, Larissa Macedo Pinto, Victor Guimaraes, Leticia Scalioni, Ana Helena Perosa, Gabriela Rodrigues Barbosa, Nancy Bellei e Marilda Siqueira on behalf of the Fiocruz COVID-19 Genomic Surveillance Network |
| EPI_ISL_15896140 | VIC, RCH Molecular Microbiology Dept. (Bio21) | WHO Collaborating Centre for Reference and Research on Influenza | Xiaomin Dong, Steven Edwards, Yi-Mo Deng, Ammar Aziz, Ian Barr |
| EPI_ISL_15896142, EPI_ISL_15896147, EPI_ISL_15896153, EPI_ISL_15896154 | QLD, Queensland Childrens Hospital | WHO Collaborating Centre for Reference and Research on Influenza | Xiaomin Dong, Steven Edwards, Yi-Mo Deng, Ammar Aziz, Ian Barr |
| EPI_ISL_15896157 | VIC, RCH Molecular Microbiology Dept. (Bio21) | WHO Collaborating Centre for Reference and Research on Influenza | Xiaomin Dong, Steven Edwards, Yi-Mo Deng, Ammar Aziz, Ian Barr |
| EPI_ISL_15896163 | QLD, Queensland Childrens Hospital | WHO Collaborating Centre for Reference and Research on Influenza | Xiaomin Dong, Steven Edwards, Yi-Mo Deng, Ammar Aziz, Ian Barr |

| | | | |
|---|---|---|---|
| EPI_ISL_15896165 | VIC, RCH Molecular Microbiology Dept. (Bio21) | WHO Collaborating Centre for Reference and Research on Influenza | Xiaomin Dong, Steven Edwards, Yi-Mo Deng, Ammar Aziz, Ian Barr |
| EPI_ISL_15896173, EPI_ISL_15896174 | QLD, Queensland Childrens Hospital | WHO Collaborating Centre for Reference and Research on Influenza | Xiaomin Dong, Steven Edwards, Yi-Mo Deng, Ammar Aziz, Ian Barr |
| EPI_ISL_15896180, EPI_ISL_15896184, EPI_ISL_15896185 | VIC, RCH Molecular Microbiology Dept. (Bio21) | WHO Collaborating Centre for Reference and Research on Influenza | Xiaomin Dong, Steven Edwards, Yi-Mo Deng, Ammar Aziz, Ian Barr |
| EPI_ISL_15896190, EPI_ISL_15896191, EPI_ISL_15896192 | NT, Royal Darwin Hospital | WHO Collaborating Centre for Reference and Research on Influenza | Xiaomin Dong, Steven Edwards, Yi-Mo Deng, Ammar Aziz, Ian Barr |
| EPI_ISL_15896196, EPI_ISL_15896197 | VIC, RCH Molecular Microbiology Dept. (Bio21) | WHO Collaborating Centre for Reference and Research on Influenza | Xiaomin Dong, Steven Edwards, Yi-Mo Deng, Ammar Aziz, Ian Barr |
| EPI_ISL_15896200, EPI_ISL_15896201 | NT, Royal Darwin Hospital | WHO Collaborating Centre for Reference and Research on Influenza | Xiaomin Dong, Steven Edwards, Yi-Mo Deng, Ammar Aziz, Ian Barr |
| EPI_ISL_15896204, EPI_ISL_15896205, EPI_ISL_15896206, EPI_ISL_15896207, EPI_ISL_15896208, EPI_ISL_15896209 | QLD, Queensland Childrens Hospital | WHO Collaborating Centre for Reference and Research on Influenza | Xiaomin Dong, Steven Edwards, Yi-Mo Deng, Ammar Aziz, Ian Barr |
| EPI_ISL_15896210, EPI_ISL_15896212, EPI_ISL_15896214, EPI_ISL_15896215, EPI_ISL_15896216, EPI_ISL_15896217, EPI_ISL_15896219, EPI_ISL_15896220, EPI_ISL_15896221 | VIC, RCH Molecular Microbiology Dept. (Bio21) | WHO Collaborating Centre for Reference and Research on Influenza | Xiaomin Dong, Steven Edwards, Yi-Mo Deng, Ammar Aziz, Ian Barr |
| EPI_ISL_15896223, EPI_ISL_15896224, EPI_ISL_15896225, EPI_ISL_15896226, EPI_ISL_15896227, EPI_ISL_15896228, EPI_ISL_15896229 | NT, Royal Darwin Hospital | WHO Collaborating Centre for Reference and Research on Influenza | Xiaomin Dong, Steven Edwards, Yi-Mo Deng, Ammar Aziz, Ian Barr |
| EPI_ISL_15896230, EPI_ISL_15896231, EPI_ISL_15896232, EPI_ISL_15896233, EPI_ISL_15896235, EPI_ISL_15896236, EPI_ISL_15896237, EPI_ISL_15896238, EPI_ISL_15896239, EPI_ISL_15896240, EPI_ISL_15896241, EPI_ISL_15896242 | | | |
| see above | QLD, Queensland Childrens Hospital | WHO Collaborating Centre for Reference and Research on Influenza | Xiaomin Dong, Steven Edwards, Yi-Mo Deng, Ammar Aziz, Ian Barr |
| EPI_ISL_15896243, EPI_ISL_15896244, EPI_ISL_15896245, EPI_ISL_15896246, EPI_ISL_15896247, EPI_ISL_15896248 | VIC, RCH Molecular Microbiology Dept. (Bio21) | WHO Collaborating Centre for Reference and Research on Influenza | Xiaomin Dong, Steven Edwards, Yi-Mo Deng, Ammar Aziz, Ian Barr |
| EPI_ISL_15896249, EPI_ISL_15896250, EPI_ISL_15896251, EPI_ISL_15896252, EPI_ISL_15896253, EPI_ISL_15896254, EPI_ISL_15896255, EPI_ISL_15896256, EPI_ISL_15896258, EPI_ISL_15896259, EPI_ISL_15896260, EPI_ISL_15896261, EPI_ISL_15896262 | | | |
| see above | NT, Royal Darwin Hospital | WHO Collaborating Centre for Reference and Research on Influenza | Xiaomin Dong, Steven Edwards, Yi-Mo Deng, Ammar Aziz, Ian Barr |
| EPI_ISL_15896263, EPI_ISL_15896264, EPI_ISL_15896265, EPI_ISL_15896266, EPI_ISL_15896267, EPI_ISL_15896268, EPI_ISL_15896269, EPI_ISL_15896270, EPI_ISL_15896271, EPI_ISL_15896272, EPI_ISL_15896273, EPI_ISL_15896274 | | | |
| see above | QLD, Queensland Childrens Hospital | WHO Collaborating Centre for Reference and Research on Influenza | Xiaomin Dong, Steven Edwards, Yi-Mo Deng, Ammar Aziz, Ian Barr |
| EPI_ISL_15896275, EPI_ISL_15896276, EPI_ISL_15896277, EPI_ISL_15896278 | VIC, RCH Molecular Microbiology Dept. (Bio21) | WHO Collaborating Centre for Reference and Research on Influenza | Xiaomin Dong, Steven Edwards, Yi-Mo Deng, Ammar Aziz, Ian Barr |
| EPI_ISL_15896279, EPI_ISL_15896280, EPI_ISL_15896281, EPI_ISL_15896282, EPI_ISL_15896283, EPI_ISL_15896284, EPI_ISL_15896285, EPI_ISL_15896286, EPI_ISL_15896287, EPI_ISL_15896288 | NT, Royal Darwin Hospital | WHO Collaborating Centre for Reference and Research on Influenza | Xiaomin Dong, Steven Edwards, Yi-Mo Deng, Ammar Aziz, Ian Barr |
| EPI_ISL_15896289, EPI_ISL_15896290, EPI_ISL_15896291 | QLD, Queensland Childrens Hospital | WHO Collaborating Centre for Reference and Research on Influenza | Xiaomin Dong, Steven Edwards, Yi-Mo Deng, Ammar Aziz, Ian Barr |
| EPI_ISL_15896292, EPI_ISL_15896293, EPI_ISL_15896295, EPI_ISL_15896296 | VIC, RCH Molecular Microbiology Dept. (Bio21) | WHO Collaborating Centre for Reference and Research on Influenza | Xiaomin Dong, Steven Edwards, Yi-Mo Deng, Ammar Aziz, Ian Barr |
| EPI_ISL_15896297, EPI_ISL_15896298 | NT, Royal Darwin Hospital | WHO Collaborating Centre for Reference and Research on Influenza | Xiaomin Dong, Steven Edwards, Yi-Mo Deng, Ammar Aziz, Ian Barr |
| EPI_ISL_15896299, EPI_ISL_15896300 | VIC, RCH Molecular Microbiology Dept. (Bio21) | WHO Collaborating Centre for Reference and Research on Influenza | Xiaomin Dong, Steven Edwards, Yi-Mo Deng, Ammar Aziz, Ian Barr |
| EPI_ISL_16006110, EPI_ISL_16006111, EPI_ISL_16006112, EPI_ISL_16006114, EPI_ISL_16006115, EPI_ISL_16006116, EPI_ISL_16006117, EPI_ISL_16006118 | Laboratory of Virology - Institute of Public Health Skopje, Macedonia | Laboratory of Virology - Institute of Public Health Skopje, Macedonia | Teodora Buzharova, Golubinka Boshevska, Elizabeta Jancheska |
| EPI_ISL_16132314, EPI_ISL_16132315, EPI_ISL_16132317, EPI_ISL_16132318, EPI_ISL_16132319, EPI_ISL_16132320, EPI_ISL_16132321, EPI_ISL_16132322, EPI_ISL_16132323, EPI_ISL_16132324, EPI_ISL_16132325, EPI_ISL_16132326, EPI_ISL_16132327 | | | |
| see above | University of WashingtonLaboratory Medicine and Pathology, UW Medicine | University of WashingtonLaboratory Medicine and Pathology, UW Medicine | Goya,S., Sereewit,J., Pfalmer,D., Nguyen,T., Sobolik,E.B. and Greninger,A.L. |
| EPI_ISL_16132380, EPI_ISL_16132381, EPI_ISL_16132383 | Infectious Disease Program, Broad Institute of Harvard and MIT | Infectious Disease Program, Broad Institute of Harvard and MIT | Adams,G., Uddin,R., Messer,K., Dobbins,S., Kotzen,B., Siddle,K.J., Petros,B., Paull,J., Brock-Fisher,T., Levine,Z., Krasilnikova,L., Negrete-Arenas,F., Tomkins-Tinch,C., Chaluvadi,S., Marmol,C., DeRuff,K., Loreth,C., Birren,B.W., Park,D.J., MacInnis,B.L., Sabeti,P.C., Rosenberg,E., Turbett,S., Lemieux,J.E., Adams,G., Uddin,R., Messer,K., Dobbins,S., Kotzen,B., Siddle,K.J., Petros,B., Paull,J., Brock-Fisher,T., Levine,Z., Krasilnikova,L., Negrete-Arenas,F., Tomkins-Tinch,C., Chaluvadi,S., Marmol,C., DeRuff,K., Loreth,C., Birren,B.W., Park,D.J., MacInnis,B.L., Sabeti,P.C., Rosenberg,E., Turbett,S., Lemieux,J.E. |
| EPI_ISL_16132413, EPI_ISL_16132414, EPI_ISL_16132415, EPI_ISL_16132416, EPI_ISL_16132417, EPI_ISL_16132418, EPI_ISL_16132419, EPI_ISL_16132420 | University of WashingtonLaboratory Medicine and Pathology, UW Medicine | University of WashingtonLaboratory Medicine and Pathology, UW Medicine | Goya,S., Sereewit,J., Pfalmer,D., Nguyen,T., Sobolik,E.B. and Greninger,A.L. |
| EPI_ISL_16132421 | University of Washington Laboratory Medicine and Pathology, UW Medicine | University of Washington Laboratory Medicine and Pathology, UW Medicine | Goya,S., Sereewit,J., Pfalmer,D., Nguyen,T., Sobolik,E.B. and Greninger,A.L. |
| EPI_ISL_16251637, EPI_ISL_16251648, EPI_ISL_16251649, EPI_ISL_16251655, EPI_ISL_16251669, EPI_ISL_16251670, EPI_ISL_16251674 | Microbiology Department. Complexo Hospitalario Universitario de Vigo | Microbiology Department. Complexo Hospitalario Universitario de Vigo | Daviña C, Martinez L, Perez-Castro S |
| EPI_ISL_16289246, EPI_ISL_16289248, EPI_ISL_16289249, EPI_ISL_16289253, EPI_ISL_16289271, EPI_ISL_16289279, EPI_ISL_16289294 | National Influenza Center, National Institute of Hygiene and Epidemiology (NIHE) | National Influenza Center, National Institute of Hygiene and Epidemiology (NIHE) | Ung Thi Hong Trang, Hoang Vu Mai Phuong, Nguyen Huy Hoang, Nguyen Le Khanh Hang, Le Thi Quynh Mai |
| EPI_ISL_1647470, EPI_ISL_1647471, EPI_ISL_1647472, EPI_ISL_1647473, EPI_ISL_1647474, EPI_ISL_1647475, EPI_ISL_1647476, EPI_ISL_1647477, EPI_ISL_1647478, EPI_ISL_1647479, EPI_ISL_1647480, EPI_ISL_1647481, EPI_ISL_1647482, EPI_ISL_1647483, EPI_ISL_1647484, EPI_ISL_1647485, EPI_ISL_1647486, EPI_ISL_1647487, EPI_ISL_1647488, EPI_ISL_1647489 | | | |
| see above | Instituto Nacional de Saúde | WHO Influenza Centre for Reference and Research on Influenza | Angela Todd, Yi-Mo Deng, Almiro Rogerio Tivane, Naomi Komadina |
| EPI_ISL_1647490, EPI_ISL_1647491, EPI_ISL_1647492, EPI_ISL_1647493, EPI_ISL_1647494, EPI_ISL_1647495, EPI_ISL_1647496, EPI_ISL_1647497, EPI_ISL_1647498, EPI_ISL_1647499, EPI_ISL_1647500, EPI_ISL_1647501, EPI_ISL_1647502, EPI_ISL_1647503, EPI_ISL_1647504, EPI_ISL_1647505, EPI_ISL_1647506, EPI_ISL_1647507, EPI_ISL_1647508, EPI_ISL_1647509, EPI_ISL_1647510, EPI_ISL_1647511, EPI_ISL_1647512, EPI_ISL_1647513, EPI_ISL_1647514, EPI_ISL_1647515, EPI_ISL_1647516, EPI_ISL_1647517, EPI_ISL_1647518, EPI_ISL_1647519, EPI_ISL_1647520, EPI_ISL_1647521, EPI_ISL_1647522, EPI_ISL_1647523, EPI_ISL_1647524, EPI_ISL_1647525, EPI_ISL_1647526, EPI_ISL_1647527, EPI_ISL_1647528, EPI_ISL_1647529, EPI_ISL_1647530, EPI_ISL_1647531, EPI_ISL_1647532, EPI_ISL_1647533, EPI_ISL_1647534, EPI_ISL_1647535, EPI_ISL_1647536, EPI_ISL_1647537, EPI_ISL_1647538, EPI_ISL_1647539, EPI_ISL_1647540, EPI_ISL_1647541, EPI_ISL_1647542, EPI_ISL_1647543, EPI_ISL_1647544, EPI_ISL_1647545, EPI_ISL_1647546, EPI_ISL_1647547, EPI_ISL_1647548, EPI_ISL_1647549, EPI_ISL_1647550, EPI_ISL_1647551, EPI_ISL_1647552, EPI_ISL_1647553, EPI_ISL_1647554, EPI_ISL_1647555, EPI_ISL_1647556, EPI_ISL_1647557, EPI_ISL_1647558, EPI_ISL_1647559, EPI_ISL_1647560, EPI_ISL_1647561, EPI_ISL_1647562, EPI_ISL_1647563, EPI_ISL_1647564, EPI_ISL_1647565, EPI_ISL_1647566, EPI_ISL_1647567, EPI_ISL_1647568, EPI_ISL_1647569, EPI_ISL_1647570, EPI_ISL_1647571, EPI_ISL_1647572, EPI_ISL_1647573, EPI_ISL_1647574, EPI_ISL_1647575, EPI_ISL_1647576, EPI_ISL_1647577, EPI_ISL_1647578, EPI_ISL_1647579, EPI_ISL_1647580, EPI_ISL_1647581, EPI_ISL_1647582, EPI_ISL_1647583, EPI_ISL_1647584, EPI_ISL_1647585, EPI_ISL_1647586, EPI_ISL_1647587, EPI_ISL_1647588, EPI_ISL_1647589, EPI_ISL_1647590, EPI_ISL_1647591, EPI_ISL_1647592, EPI_ISL_1647593, EPI_ISL_1647594, EPI_ISL_1647595, EPI_ISL_1647596, EPI_ISL_1647597, EPI_ISL_1647598, EPI_ISL_1647599, EPI_ISL_1647600 | | | |
| see above | Respiratory Virus Unit, National Infection Service, Public Health England | National Infection Service, Public Health England | Zambon M, Talts T, Ellis J, Miah S, Platt S |
| EPI_ISL_16533855, EPI_ISL_16533857, EPI_ISL_16533858, EPI_ISL_16533861, EPI_ISL_16533862, EPI_ISL_16533863, EPI_ISL_16533864, EPI_ISL_16533866 | Center for Virology, Medical University of Vienna | Center for Virology, Medical University of Vienna | Jeremy V. Camp, Monika Redlberger-Fritz |
| EPI_ISL_1653993, EPI_ISL_1653994, EPI_ISL_1653995, EPI_ISL_1653996, EPI_ISL_1653997, EPI_ISL_1653998, EPI_ISL_1653999 | Royal Children's Hospital | WHO Influenza Centre for Reference and Research on Influenza | Angela Todd, Yi-Mo Deng, Annette Alafaci, Naomi Komadina |
| EPI_ISL_16609708, EPI_ISL_16609709, EPI_ISL_16609711, EPI_ISL_16609712, EPI_ISL_16609713, EPI_ISL_16609714, EPI_ISL_16609715, EPI_ISL_16609716, EPI_ISL_16609717, EPI_ISL_16609718, EPI_ISL_16609719, EPI_ISL_16609720, EPI_ISL_16609721, EPI_ISL_16609722, EPI_ISL_16609723, EPI_ISL_16609724, EPI_ISL_16609725, EPI_ISL_16609726, EPI_ISL_16609727, EPI_ISL_16609728, EPI_ISL_16609729, EPI_ISL_16609730, EPI_ISL_16609732, EPI_ISL_16609735, EPI_ISL_16609736, EPI_ISL_16609737, EPI_ISL_16609738, EPI_ISL_16609739, EPI_ISL_16609740, EPI_ISL_16609743, EPI_ISL_16609744, EPI_ISL_16609745, EPI_ISL_16609751, EPI_ISL_16609753, EPI_ISL_16609756, EPI_ISL_16609757, EPI_ISL_16609761, EPI_ISL_16609762, EPI_ISL_16609763, EPI_ISL_16609765, EPI_ISL_16609767, EPI_ISL_16609768, EPI_ISL_16609769, EPI_ISL_16672799, EPI_ISL_16672800, EPI_ISL_16672801, EPI_ISL_16672802, EPI_ISL_16672803, EPI_ISL_16672804, EPI_ISL_16672805, EPI_ISL_16672806, EPI_ISL_16672807, EPI_ISL_16672808, EPI_ISL_16672809, EPI_ISL_16672810, EPI_ISL_16672811, EPI_ISL_16672812, EPI_ISL_16672814, EPI_ISL_16672815, EPI_ISL_16672816, EPI_ISL_16672817, EPI_ISL_16672818, EPI_ISL_16672819, EPI_ISL_16672821, EPI_ISL_16672822, EPI_ISL_16672823, EPI_ISL_16672824, EPI_ISL_16672825, EPI_ISL_16672830, EPI_ISL_16672832, EPI_ISL_16672834, EPI_ISL_16672835, EPI_ISL_16672854, EPI_ISL_16672855, EPI_ISL_16672857, EPI_ISL_16672858, EPI_ISL_16672859, EPI_ISL_16672860, EPI_ISL_16672861, EPI_ISL_16672862, EPI_ISL_16672863, EPI_ISL_16672864, EPI_ISL_16672865, EPI_ISL_16672866, EPI_ISL_16672867, EPI_ISL_16672868, EPI_ISL_16672870, EPI_ISL_16672871, EPI_ISL_16672872, EPI_ISL_16672873, EPI_ISL_16672874, EPI_ISL_16672875, EPI_ISL_16672877, EPI_ISL_16672878, EPI_ISL_16672879, EPI_ISL_16672880, EPI_ISL_16672881, EPI_ISL_16672883, EPI_ISL_16672884, EPI_ISL_16672885, EPI_ISL_16672886, EPI_ISL_16672888, EPI_ISL_16672890, EPI_ISL_16672891, EPI_ISL_16672917, EPI_ISL_16672918, EPI_ISL_16672921, EPI_ISL_16672922, EPI_ISL_16672923, EPI_ISL_16672924, EPI_ISL_16672925, EPI_ISL_16672932, EPI_ISL_16672933, EPI_ISL_16672935, EPI_ISL_16672937, EPI_ISL_16672939, EPI_ISL_16672941, EPI_ISL_16672945 | | | |
| see above | Respiratory Virus Unit / Reference Microbiology Services / UK Health Security Agency | Reference Microbiology Services / UK Health Security Agency | Zambon M. Talts T. Mosscrop L. Miah S |
| EPI_ISL_16681200, EPI_ISL_16681221, EPI_ISL_16681260, EPI_ISL_16681272, EPI_ISL_16681286, EPI_ISL_16681299, EPI_ISL_16681312, EPI_ISL_16681327, EPI_ISL_16681338 | Broad Institute of Harvard and Massachusetts Institute of Technology | Broad Institute of Harvard and Massachusetts Institute of Technology | Adams,G., Uddin,R., Messer,K., Dobbins,S., Kotzen,B., Siddle,K.J., Petros,B., Paull,J., Brock-Fisher,T., Levine,Z., Krasilnikova,L., Negrete-Arenas,F., Tomkins-Tinch,C., Chaluvadi,S., Marmol,C., DeRuff,K., Loreth,C., Birren,B.W., Park,D.J., MacInnis,B.L., Sabeti,P.C., Rosenberg,E., Turbett,S. and Lemieux,J.E. |

| | | | |
|---|---|---|---|
| EPI_ISL_16708974, EPI_ISL_16708978, EPI_ISL_16708983, EPI_ISL_16708994, EPI_ISL_16708999, EPI_ISL_16709000, EPI_ISL_16709001, EPI_ISL_16709006, EPI_ISL_16709007, EPI_ISL_16709008, EPI_ISL_16709010 | | | |
| see above | University Hospitals of Leicester NHS Trust Pathology Services / Leicester Royal Infirmary | Reference Microbiology Services / UK Health Security Agency | Williams T. Zambon M. Talts T. Mosscrop L. Holmes C. Miah S |
| EPI_ISL_16714277 | Reference Microbiology Services / UK Health Security Agency | Reference Microbiology Services / UK Health Security Agency | Zambon M. Talts T. Mosscrop L. Miah S |
| EPI_ISL_16714278, EPI_ISL_16714279, EPI_ISL_16714283, EPI_ISL_16714287, EPI_ISL_16714289, EPI_ISL_16714291, EPI_ISL_16714294, EPI_ISL_16714298, EPI_ISL_16714305, EPI_ISL_16714306, EPI_ISL_16714307, EPI_ISL_16714313, EPI_ISL_16714315, EPI_ISL_16714319, EPI_ISL_16714323, EPI_ISL_16714327, EPI_ISL_16714328, EPI_ISL_16714330, EPI_ISL_16714332, EPI_ISL_16714333, EPI_ISL_16714335, EPI_ISL_16714336, EPI_ISL_16714338, EPI_ISL_16714339, EPI_ISL_16714340, EPI_ISL_16714342, EPI_ISL_16714345, EPI_ISL_16714348, EPI_ISL_16714349, EPI_ISL_16714351, EPI_ISL_16714352, EPI_ISL_16714353, EPI_ISL_16714356, EPI_ISL_16714357, EPI_ISL_16714358, EPI_ISL_16714359, EPI_ISL_16714364, EPI_ISL_16714369, EPI_ISL_16714370, EPI_ISL_16714371, EPI_ISL_16714375, EPI_ISL_16714377, EPI_ISL_16714378, EPI_ISL_16714380, EPI_ISL_16714384, EPI_ISL_16714385, EPI_ISL_16714386, EPI_ISL_16714387, EPI_ISL_16714389, EPI_ISL_16714394, EPI_ISL_16714408, EPI_ISL_16714413, EPI_ISL_16714418, EPI_ISL_16714421, EPI_ISL_16714422, EPI_ISL_16714423, EPI_ISL_16714424, EPI_ISL_16714426, EPI_ISL_16714427, EPI_ISL_16714453, EPI_ISL_16714454, EPI_ISL_16714460, EPI_ISL_16714463, EPI_ISL_16714466, EPI_ISL_16714469, EPI_ISL_16714470, EPI_ISL_16714472 | | | |
| see above | University Hospital Southampton / Southampton Specialist Virology Centre | Reference Microbiology Services / UK Health Security Agency | Williams T. Zambon M. Talts T. Mosscrop L. Pelosi E. Miah S |
| EPI_ISL_16714572, EPI_ISL_16714573, EPI_ISL_16714574, EPI_ISL_16714576, EPI_ISL_16714578, EPI_ISL_16714580, EPI_ISL_16714581, EPI_ISL_16714583, EPI_ISL_16714584, EPI_ISL_16714585, EPI_ISL_16714586, EPI_ISL_16714587, EPI_ISL_16714589, EPI_ISL_16714590, EPI_ISL_16714592, EPI_ISL_16714594, EPI_ISL_16714595, EPI_ISL_16714596, EPI_ISL_16714597, EPI_ISL_16714598, EPI_ISL_16714599, EPI_ISL_16714600, EPI_ISL_16714602, EPI_ISL_16714603, EPI_ISL_16714604, EPI_ISL_16714605, EPI_ISL_16714607, EPI_ISL_16714608, EPI_ISL_16714609, EPI_ISL_16714610, EPI_ISL_16714612, EPI_ISL_16714613, EPI_ISL_16714614, EPI_ISL_16714615, EPI_ISL_16714616, EPI_ISL_16714618, EPI_ISL_16714620, EPI_ISL_16714621, EPI_ISL_16714622, EPI_ISL_16714623, EPI_ISL_16714624, EPI_ISL_16714625, EPI_ISL_16714626, EPI_ISL_16714627, EPI_ISL_16714628, EPI_ISL_16714629, EPI_ISL_16714630, EPI_ISL_16714631, EPI_ISL_16714632, EPI_ISL_16714633, EPI_ISL_16714634, EPI_ISL_16714635, EPI_ISL_16714636, EPI_ISL_16714637, EPI_ISL_16714638, EPI_ISL_16714639, EPI_ISL_16714640, EPI_ISL_16714641, EPI_ISL_16714642, EPI_ISL_16714643, EPI_ISL_16714644, EPI_ISL_16714645, EPI_ISL_16714646, EPI_ISL_16714647, EPI_ISL_16714648, EPI_ISL_16714649, EPI_ISL_16714650, EPI_ISL_16714651, EPI_ISL_16714652, EPI_ISL_16714653, EPI_ISL_16714654, EPI_ISL_16714655, EPI_ISL_16714656, EPI_ISL_16714657, EPI_ISL_16714658, EPI_ISL_16714659, EPI_ISL_16714660, EPI_ISL_16714661, EPI_ISL_16714662, EPI_ISL_16714663, EPI_ISL_16714664, EPI_ISL_16714665, EPI_ISL_16714666, EPI_ISL_16714667, EPI_ISL_16714668, EPI_ISL_16714669, EPI_ISL_16714670, EPI_ISL_16714671, EPI_ISL_16714672, EPI_ISL_16714673, EPI_ISL_16714674, EPI_ISL_16714675, EPI_ISL_16714676, EPI_ISL_16714677, EPI_ISL_16714678, EPI_ISL_16714679, EPI_ISL_16714680, EPI_ISL_16714681, EPI_ISL_16714682, EPI_ISL_16714683, EPI_ISL_16714684, EPI_ISL_16714685, EPI_ISL_16714686, EPI_ISL_16714687, EPI_ISL_16714705, EPI_ISL_16714706, EPI_ISL_16714707, EPI_ISL_16714708, EPI_ISL_16714709, EPI_ISL_16714710, EPI_ISL_16714711, EPI_ISL_16714712, EPI_ISL_16714713, EPI_ISL_16714714, EPI_ISL_16714715, EPI_ISL_16714716, EPI_ISL_16714717, EPI_ISL_16714718, EPI_ISL_16714719, EPI_ISL_16714720, EPI_ISL_16714721, EPI_ISL_16714722, EPI_ISL_16714723, EPI_ISL_16714724, EPI_ISL_16714725, EPI_ISL_16714726, EPI_ISL_16714727, EPI_ISL_16714728, EPI_ISL_16714729, EPI_ISL_16714730, EPI_ISL_16714731, EPI_ISL_16714732, EPI_ISL_16714733, EPI_ISL_16714734, EPI_ISL_16714735, EPI_ISL_16714736, EPI_ISL_16714737, EPI_ISL_16714738, EPI_ISL_16714739, EPI_ISL_16714741, EPI_ISL_16714742 | | | |
| see above | Respiratory Virus Unit / Reference Microbiology Services / UK Health Security Agency | Reference Microbiology Services / UK Health Security Agency | Zambon M. Talts T. Mosscrop L. Miah S |
| EPI_ISL_16714744 | NT, Royal Darwin Hospital | WHO Collaborating Centre for Reference and Research on Influenza | Xiaomin Dong, Steven Edwards, Yi-Mo Deng, Ammar Aziz, Ian Barr |
| EPI_ISL_16714746, EPI_ISL_16714748, EPI_ISL_16714750, EPI_ISL_16714751, EPI_ISL_16714752, EPI_ISL_16714753, EPI_ISL_16714754, EPI_ISL_16714755, EPI_ISL_16714756 | VIC, RCH Molecular Microbiology Dept. (Bio21) | WHO Collaborating Centre for Reference and Research on Influenza | Xiaomin Dong, Steven Edwards, Yi-Mo Deng, Ammar Aziz, Ian Barr |
| EPI_ISL_16714757, EPI_ISL_16714758, EPI_ISL_16714759, EPI_ISL_16714761, EPI_ISL_16714762, EPI_ISL_16714763, EPI_ISL_16714764, EPI_ISL_16714765, EPI_ISL_16714766, EPI_ISL_16714767, EPI_ISL_16714768, EPI_ISL_16714769, EPI_ISL_16714771, EPI_ISL_16714772, EPI_ISL_16714773, EPI_ISL_16714775, EPI_ISL_16714776, EPI_ISL_16714777, EPI_ISL_16714778, EPI_ISL_16714779 | | | |
| see above | NT, Royal Darwin Hospital | WHO Collaborating Centre for Reference and Research on Influenza | Xiaomin Dong, Steven Edwards, Yi-Mo Deng, Ammar Aziz, Ian Barr |
| EPI_ISL_16714782, EPI_ISL_16714783, EPI_ISL_16714784, EPI_ISL_16714785, EPI_ISL_16714787, EPI_ISL_16714788, EPI_ISL_16714792, EPI_ISL_16714793, EPI_ISL_16714794, EPI_ISL_16714796, EPI_ISL_16714798, EPI_ISL_16714801 | | | |
| see above | VIC, Royal Children's Hospital Victoria Australia | WHO Collaborating Centre for Reference and Research on Influenza | Xiaomin Dong, Steven Edwards, Yi-Mo Deng, Ammar Aziz, Ian Barr |
| EPI_ISL_16714802, EPI_ISL_16714804, EPI_ISL_16714805, EPI_ISL_16714806, EPI_ISL_16714807, EPI_ISL_16714808, EPI_ISL_16714811, EPI_ISL_16714812, EPI_ISL_16714813 | NT, Royal Darwin Hospital | WHO Collaborating Centre for Reference and Research on Influenza | Xiaomin Dong, Steven Edwards, Yi-Mo Deng, Ammar Aziz, Ian Barr |
| EPI_ISL_16737023, EPI_ISL_16737024, EPI_ISL_16737025, EPI_ISL_16737026, EPI_ISL_16737027, EPI_ISL_16737028 | Respiratory Virus Unit / Reference Microbiology Services / UK Health Security Agency | Reference Microbiology Services / UK Health Security Agency | Zambon M. Talts T. Mosscrop L. Miah S |
| EPI_ISL_16737031, EPI_ISL_16737032, EPI_ISL_16737034, EPI_ISL_16737035, EPI_ISL_16737036, EPI_ISL_16737037, EPI_ISL_16737039 | University Hospital Southampton / Southampton Specialist Virology Centre | Reference Microbiology Services / UK Health Security Agency | Williams T. Zambon M. Talts T. Mosscrop L. Pelosi E. Miah S |
| EPI_ISL_16746474, EPI_ISL_16746476, EPI_ISL_16746477 | Respiratory Virus Unit / Reference Microbiology Services / UK Health Security Agency | Reference Microbiology Services / UK Health Security Agency | Zambon M. Talts T. Mosscrop L. Miah S |
| EPI_ISL_16881187, EPI_ISL_16881188, EPI_ISL_16881189, EPI_ISL_16881190, EPI_ISL_16881191, EPI_ISL_16881192, EPI_ISL_16881193, EPI_ISL_16881194, EPI_ISL_16881195, EPI_ISL_16881196, EPI_ISL_16881197, EPI_ISL_16881198, EPI_ISL_16881199, EPI_ISL_16881200, EPI_ISL_16881201, EPI_ISL_16881202, EPI_ISL_16881203, EPI_ISL_16881204, EPI_ISL_16881205, EPI_ISL_16881206, EPI_ISL_16881207, EPI_ISL_16881208 | | | |
| see above | West of Scotland Specialist Virology Centre | West of Scotland Specialist Virology Centre | Lynne Ferguson, Imogen Johnston-Menzies, Rory Gunson |
| EPI_ISL_16905446, EPI_ISL_16905449, EPI_ISL_16905450 | National Institute of Hygiene, virology departement, National Influenza Center | Centers for Disease Control and Prevention, Coronavirus and Other Respiratory Viruses Division | Megha Aggarwal, Lijuan Wang, Ji In Park, Amanda Smith and Everardo Vega |
| EPI_ISL_16929303, EPI_ISL_16929305, EPI_ISL_16929307 | West of Scotland Specialist Virology Centre | West of Scotland Specialist Virology Centre | Lynne Ferguson, Imogen Johnston-Menzies, Rory Gunson |
| EPI_ISL_16959469 | Ranfurly Medical Centre, Auckland | Institute of Environmental Science and Research | Klarysse Berquist, Lauren Jelley, Una Ren, Meaghan O'Neill |
| EPI_ISL_16959470 | Rolleston Central Health, Christchurch | Institute of Environmental Science and Research | Klarysse Berquist, Lauren Jelley, Una Ren, Meaghan O'Neill |
| EPI_ISL_16959471 | Upper Hutt Health Centre | Institute of Environmental Science and Research | Klarysse Berquist, Lauren Jelley, Una Ren, Meaghan O'Neill |
| EPI_ISL_16959472 | Westland Medical Centre | Institute of Environmental Science and Research | Klarysse Berquist, Lauren Jelley, Una Ren, Meaghan O'Neill |
| EPI_ISL_16959475 | LabPlus | Institute of Environmental Science and Research | Klarysse Berquist, Lauren Jelley, Una Ren, Meaghan O'Neill |
| EPI_ISL_16959476, EPI_ISL_16959477, EPI_ISL_16959478, EPI_ISL_16959479, EPI_ISL_16959480, EPI_ISL_16959481, EPI_ISL_16959482, EPI_ISL_16959484, EPI_ISL_16959485, EPI_ISL_16959486, EPI_ISL_16959487, EPI_ISL_16959488, EPI_ISL_16959489, EPI_ISL_16959491, EPI_ISL_16959492, EPI_ISL_16959493, EPI_ISL_16959494, EPI_ISL_16959496, EPI_ISL_16959497, EPI_ISL_16959498, EPI_ISL_16959499, EPI_ISL_16959501, EPI_ISL_16959502, EPI_ISL_16959503, EPI_ISL_16959504, EPI_ISL_16959505, EPI_ISL_16959506, EPI_ISL_16959508, EPI_ISL_16959510, EPI_ISL_16959511, EPI_ISL_16959512, EPI_ISL_16959513, EPI_ISL_16959515, EPI_ISL_16959516, EPI_ISL_16959517, EPI_ISL_16959518, EPI_ISL_16959519, EPI_ISL_16959520, EPI_ISL_16959521, EPI_ISL_16959523, EPI_ISL_16959524, EPI_ISL_16959525, EPI_ISL_16959526, EPI_ISL_16959527, EPI_ISL_16959529 | | | |
| see above | ESR - Wallaceville | Institute of Environmental Science and Research | Klarysse Berquist, Lauren Jelley, Una Ren, Meaghan O'Neill |
| EPI_ISL_16959544 | Middlemore Hospital | Institute of Environmental Science and Research | Klarysse Berquist, Lauren Jelley, Una Ren, Meaghan O'Neill |
| EPI_ISL_16959552, EPI_ISL_16959553, EPI_ISL_16959554, EPI_ISL_16959555 | Hawkes Bay District Health Board | Institute of Environmental Science and Research | Klarysse Berquist, Lauren Jelley, Una Ren, Meaghan O'Neill |
| EPI_ISL_16959556, EPI_ISL_16959558, EPI_ISL_16959559, EPI_ISL_16959560 | Wellington SCL | Institute of Environmental Science and Research | Klarysse Berquist, Lauren Jelley, Una Ren, Meaghan O'Neill |
| EPI_ISL_16959586, EPI_ISL_16959589, EPI_ISL_16959590, EPI_ISL_16959610, EPI_ISL_16959611, EPI_ISL_16959624, EPI_ISL_16959631, EPI_ISL_16959650, EPI_ISL_16959654, EPI_ISL_16959657, EPI_ISL_16959658, EPI_ISL_16959675, EPI_ISL_16959678, EPI_ISL_16959689, EPI_ISL_16959696, EPI_ISL_16959701, EPI_ISL_16959704, EPI_ISL_16959705, EPI_ISL_16959707, EPI_ISL_16959716, EPI_ISL_16959717, EPI_ISL_16959723, EPI_ISL_16959726, EPI_ISL_16959729 | | | |
| see above | Middlemore Hospital | Institute of Environmental Science and Research | Klarysse Berquist, Lauren Jelley, Una Ren, Meaghan O'Neill |
| EPI_ISL_16959739, EPI_ISL_16959740 | Southern Community Lab - Dunedin | Institute of Environmental Science and Research | Klarysse Berquist, Lauren Jelley, Una Ren, Meaghan O'Neill |
| EPI_ISL_16959752, EPI_ISL_16959753, EPI_ISL_16959755, EPI_ISL_16959760, EPI_ISL_16959761, EPI_ISL_16959763 | Middlemore Hospital | Institute of Environmental Science and Research | Klarysse Berquist, Lauren Jelley, Una Ren, Meaghan O'Neill |
| EPI_ISL_16959784, EPI_ISL_16959785, EPI_ISL_16959786, EPI_ISL_16959787, EPI_ISL_16959789, EPI_ISL_16959790, EPI_ISL_16959791, EPI_ISL_16959793, EPI_ISL_16959794, EPI_ISL_16959795, EPI_ISL_16959796, EPI_ISL_16959797, EPI_ISL_16959798, EPI_ISL_16959799, EPI_ISL_16959800, EPI_ISL_16959801, EPI_ISL_16959802, EPI_ISL_16959804, EPI_ISL_16959805, EPI_ISL_16959807, EPI_ISL_16959808, EPI_ISL_16959809, EPI_ISL_16959811, EPI_ISL_16959812, EPI_ISL_16959813, EPI_ISL_16959815, EPI_ISL_16959816, EPI_ISL_16959817 | | | |
| see above | Wellington SCL | Institute of Environmental Science and Research | Klarysse Berquist, Lauren Jelley, Una Ren, Meaghan O'Neill |
| EPI_ISL_16959823, EPI_ISL_16959824, EPI_ISL_16959825, EPI_ISL_16959827, EPI_ISL_16959828, EPI_ISL_16959837, EPI_ISL_16959838, EPI_ISL_16959839, EPI_ISL_16959840 | Middlemore Hospital | Institute of Environmental Science and Research | Klarysse Berquist, Lauren Jelley, Una Ren, Meaghan O'Neill |
| EPI_ISL_16959841, EPI_ISL_16959842, EPI_ISL_16959843, EPI_ISL_16959844, EPI_ISL_16959845, EPI_ISL_16959848, EPI_ISL_16959850 | Canterbury Health Laboratory | Institute of Environmental Science and Research | Klarysse Berquist, Lauren Jelley, Una Ren, Meaghan O'Neill |
| EPI_ISL_16959853, EPI_ISL_16959854, EPI_ISL_16959855 | Labplus | Institute of Environmental Science and Research | Klarysse Berquist, Lauren Jelley, Una Ren, Meaghan O'Neill |
| EPI_ISL_16959857, EPI_ISL_16959859, EPI_ISL_16959864, EPI_ISL_16959865, EPI_ISL_16959868 | Southern Community Lab - Dunedin | Institute of Environmental Science and Research | Klarysse Berquist, Lauren Jelley, Una Ren, Meaghan O'Neill |
| EPI_ISL_16959874, EPI_ISL_16959875, EPI_ISL_16959876, EPI_ISL_16959877, EPI_ISL_16959878 | Middlemore Hospital | Institute of Environmental Science and Research | Klarysse Berquist, Lauren Jelley, Una Ren, Meaghan O'Neill |
| EPI_ISL_16959880, EPI_ISL_16959884, EPI_ISL_16959888, EPI_ISL_16959890 | Labplus | Institute of Environmental Science and Research | Klarysse Berquist, Lauren Jelley, Una Ren, Meaghan O'Neill |
| EPI_ISL_16959891, EPI_ISL_16959892, EPI_ISL_16959893, EPI_ISL_16959894, EPI_ISL_16959896, EPI_ISL_16959897, EPI_ISL_16959898, EPI_ISL_16959899, EPI_ISL_16959900, EPI_ISL_16959901, EPI_ISL_16959902, EPI_ISL_16959903, EPI_ISL_16959904, EPI_ISL_16959906, EPI_ISL_16959908, EPI_ISL_16959909, EPI_ISL_16959910, EPI_ISL_16959911, EPI_ISL_16959912, EPI_ISL_16959913, EPI_ISL_16959914, EPI_ISL_16959916, EPI_ISL_16959917 | | | |
| see above | Wellington SCL | Institute of Environmental Science and Research | Klarysse Berquist, Lauren Jelley, Una Ren, Meaghan O'Neill |
| EPI_ISL_16959921 | PathLab Bay of Plenty | Institute of Environmental Science and Research | Klarysse Berquist, Lauren Jelley, Una Ren, Meaghan O'Neill |
| EPI_ISL_16959929, EPI_ISL_16959930, EPI_ISL_16959931, | Canterbury Health Laboratory | Institute of Environmental Science and Research | Klarysse Berquist, Lauren Jelley, Una Ren, Meaghan O'Neill |

| | | | |
|---|---|---|---|
| EPI_ISL_16959932 | | | |
| EPI_ISL_16959933, EPI_ISL_16959934, EPI_ISL_16959935 | Hawkes Bay District Health Board | Institute of Environmental Science and Research | Klarysse Berquist, Lauren Jelley, Una Ren, Meaghan O'Neill |
| EPI_ISL_16959936, EPI_ISL_16959937, EPI_ISL_16959938, EPI_ISL_16959939, EPI_ISL_16959940, EPI_ISL_16959941 | Middlemore Hospital | Institute of Environmental Science and Research | Klarysse Berquist, Lauren Jelley, Una Ren, Meaghan O'Neill |
| EPI_ISL_16959947, EPI_ISL_16959950, EPI_ISL_16959956, EPI_ISL_16959958 | LabPlus | Institute of Environmental Science and Research | Klarysse Berquist, Lauren Jelley, Una Ren, Meaghan O'Neill |
| EPI_ISL_16959984 | MedLab South Nelson | Institute of Environmental Science and Research | Klarysse Berquist, Lauren Jelley, Una Ren, Meaghan O'Neill |
| EPI_ISL_16959986, EPI_ISL_16959987, EPI_ISL_16959988, EPI_ISL_16959989, EPI_ISL_16959990, EPI_ISL_16959991, EPI_ISL_16959992 | Canterbury Health Laboratory | Institute of Environmental Science and Research | Klarysse Berquist, Lauren Jelley, Una Ren, Meaghan O'Neill |
| EPI_ISL_16959993, EPI_ISL_16959994, EPI_ISL_16959996, EPI_ISL_16959998 | Southern Community Lab - Dunedin | Institute of Environmental Science and Research | Klarysse Berquist, Lauren Jelley, Una Ren, Meaghan O'Neill |
| EPI_ISL_16960000, EPI_ISL_16960001, EPI_ISL_16960006, EPI_ISL_16960009 | Waikato Hospital | Institute of Environmental Science and Research | Klarysse Berquist, Lauren Jelley, Una Ren, Meaghan O'Neill |
| EPI_ISL_16960014, EPI_ISL_16960015, EPI_ISL_16960016, EPI_ISL_16960017 | Middlemore Hospital | Institute of Environmental Science and Research | Klarysse Berquist, Lauren Jelley, Una Ren, Meaghan O'Neill |
| EPI_ISL_16960018, EPI_ISL_16960019, EPI_ISL_16960020 | Hawkes Bay District Health Board | Institute of Environmental Science and Research | Klarysse Berquist, Lauren Jelley, Una Ren, Meaghan O'Neill |
| EPI_ISL_16960022, EPI_ISL_16960027, EPI_ISL_16960028, EPI_ISL_16960031, EPI_ISL_16960033, EPI_ISL_16960035, EPI_ISL_16960040, EPI_ISL_16960041, EPI_ISL_16960043, EPI_ISL_16960044, EPI_ISL_16960053, EPI_ISL_16960060, EPI_ISL_16960063, EPI_ISL_16960067, EPI_ISL_16960069, EPI_ISL_16960075, EPI_ISL_16960078, EPI_ISL_16960079, EPI_ISL_16960080, EPI_ISL_16960083, EPI_ISL_16960087, EPI_ISL_16960090, EPI_ISL_16960091 | | | |
| see above | LabPlus | Institute of Environmental Science and Research | Klarysse Berquist, Lauren Jelley, Una Ren, Meaghan O'Neill |
| EPI_ISL_16960092, EPI_ISL_16960093, EPI_ISL_16960094, EPI_ISL_16960095, EPI_ISL_16960096, EPI_ISL_16960097, EPI_ISL_16960098 | Middlemore Hospital | Institute of Environmental Science and Research | Klarysse Berquist, Lauren Jelley, Una Ren, Meaghan O'Neill |
| EPI_ISL_16960102 | Wellington SCL | Institute of Environmental Science and Research | Klarysse Berquist, Lauren Jelley, Una Ren, Meaghan O'Neill |
| EPI_ISL_16960105, EPI_ISL_16960106, EPI_ISL_16960107, EPI_ISL_16960108, EPI_ISL_16960109, EPI_ISL_16960110 | Canterbury Health Laboratory | Institute of Environmental Science and Research | Klarysse Berquist, Lauren Jelley, Una Ren, Meaghan O'Neill |
| EPI_ISL_16960116 | Middlemore Hospital | Institute of Environmental Science and Research | Klarysse Berquist, Lauren Jelley, Una Ren, Meaghan O'Neill |
| EPI_ISL_16960119, EPI_ISL_16960120, EPI_ISL_16960121, EPI_ISL_16960122, EPI_ISL_16960123, EPI_ISL_16960124, EPI_ISL_16960126, EPI_ISL_16960127, EPI_ISL_16960128 | Southern Community Lab - Dunedin | Institute of Environmental Science and Research | Klarysse Berquist, Lauren Jelley, Una Ren, Meaghan O'Neill |
| EPI_ISL_16960133, EPI_ISL_16960134, EPI_ISL_16960135, EPI_ISL_16960137, EPI_ISL_16960138, EPI_ISL_16960141, EPI_ISL_16960142, EPI_ISL_16960143, EPI_ISL_16960144, EPI_ISL_16960145, EPI_ISL_16960146 | | | |
| see above | LabPlus | Institute of Environmental Science and Research | Klarysse Berquist, Lauren Jelley, Una Ren, Meaghan O'Neill |
| EPI_ISL_16960147, EPI_ISL_16960148 | Canterbury Health Laboratory | Institute of Environmental Science and Research | Klarysse Berquist, Lauren Jelley, Una Ren, Meaghan O'Neill |
| EPI_ISL_16960150 | Wellington SCL | Institute of Environmental Science and Research | Klarysse Berquist, Lauren Jelley, Una Ren, Meaghan O'Neill |
| EPI_ISL_17066778 | VIC, Royal Children's Hospital Victoria Australia | WHO Collaborating Centre for Reference and Research on Influenza | Xiaomin Dong, Steven Edwards, Yi-Mo Deng, Ammar Aziz, Ian Barr |
| EPI_ISL_17066780, EPI_ISL_17066781, EPI_ISL_17066783, EPI_ISL_17066785 | NT, Royal Darwin Hospital | WHO Collaborating Centre for Reference and Research on Influenza | Xiaomin Dong, Steven Edwards, Yi-Mo Deng, Ammar Aziz, Ian Barr |
| EPI_ISL_17066786, EPI_ISL_17066787, EPI_ISL_17066789, EPI_ISL_17066790, EPI_ISL_17066791, EPI_ISL_17066792, EPI_ISL_17066794, EPI_ISL_17066796, EPI_ISL_17066798, EPI_ISL_17066799, EPI_ISL_17066801, EPI_ISL_17066803, EPI_ISL_17066805, EPI_ISL_17066806, EPI_ISL_17066808, EPI_ISL_17066809, EPI_ISL_17066810, EPI_ISL_17066811, EPI_ISL_17066812, EPI_ISL_17066814, EPI_ISL_17066815, EPI_ISL_17066816 | | | |
| see above | QLD, Pathology Queensland - Royal Brisbane and Women's Hospital | WHO Collaborating Centre for Reference and Research on Influenza | Xiaomin Dong, Steven Edwards, Yi-Mo Deng, Ammar Aziz, Ian Barr |
| EPI_ISL_17066818, EPI_ISL_17066822, EPI_ISL_17066823, EPI_ISL_17066825, EPI_ISL_17066826, EPI_ISL_17066829, EPI_ISL_17066830, EPI_ISL_17066832 | VIC, Royal Children's Hospital Victoria Australia | WHO Collaborating Centre for Reference and Research on Influenza | Xiaomin Dong, Steven Edwards, Yi-Mo Deng, Ammar Aziz, Ian Barr |
| EPI_ISL_17089180, EPI_ISL_17089181, EPI_ISL_17089182, EPI_ISL_17089185, EPI_ISL_17089186, EPI_ISL_17089188, EPI_ISL_17089189, EPI_ISL_17089190, EPI_ISL_17089191 | University of Washington - Laboratory Medicine | University of Washington - Laboratory Medicine | Sereewit,J., Xie,H., Roychoudhury,P. and Greninger,A.L. |
| EPI_ISL_17104707 | National Institute of Public Health | National Institute of Public Health | Chhorvann CHHEA, Darapheak CHAU, Sitha PRUM, Visal CHHE, Sokha DUL, Sengly MANH, Phally VY, Borann SAR, Chanthap LON, Jessica E. Manning, Sophana Chea, Savuth CHIN |
| EPI_ISL_17127011, EPI_ISL_17127012, EPI_ISL_17127013, EPI_ISL_17127014, EPI_ISL_17127015 | Servicio de Microbiologia, Hospital Universitario Virgen del Rocio | Plataforma de Medicina Computacional, Fundación Progreso y Salud | José Antonio Lepe, Javier Pérez-Florido, María Lara Jiménez |
| EPI_ISL_17177049 | University Clinical Center Tuzla, Polyclinic for Laboratory Diagnostics, Institute of Microbiology, Department of Molecular Microbiology | Republic of Macedonia, Institute of Public Health Laboratory of Virology and Molecular Diagnostics | Nijaz Tihic, Jasmina Smajlovic, Emir Halilovic |
| EPI_ISL_17221667, EPI_ISL_17221668, EPI_ISL_17221669, EPI_ISL_17221670, EPI_ISL_17221672, EPI_ISL_17221673, EPI_ISL_17221675, EPI_ISL_17221677, EPI_ISL_17221678 | University of Washington - Virology | University of Washington - Virology | Sereewit,J., Xie,H., Roychoudhury,P. and Greninger,A.L. |
| EPI_ISL_17253611, EPI_ISL_17253612, EPI_ISL_17253613, EPI_ISL_17253614, EPI_ISL_17253615, EPI_ISL_17253620, EPI_ISL_17253623, EPI_ISL_17253624, EPI_ISL_17253626 | National Public Health Institute of Slovakia | Laboratory of Genomics and Bioinformatics, Comenius University Science Park | Szemes,Tomáš;Kaliňáková,Anna;Kotvasová,Barbora;Ševčíková,Lucia;Vrabľová,Terézia;Sedláčková,Tatiana;Rusňáková,Diana;Böhmer,Miroslav;Budiš,Jaroslav;Styk,Jakub;Lipková,Nikola;Forgáčová,Michaela;Bokorová,Silvia;Mišenko,Pavol |
| EPI_ISL_17258468, EPI_ISL_17258469, EPI_ISL_17258470, EPI_ISL_17258471, EPI_ISL_17258472, EPI_ISL_17258473, EPI_ISL_17258474, EPI_ISL_17258475, EPI_ISL_17258476, EPI_ISL_17258477, EPI_ISL_17258478, EPI_ISL_17258479, EPI_ISL_17258480, EPI_ISL_17258481, EPI_ISL_17258482, EPI_ISL_17258483, EPI_ISL_17258484, EPI_ISL_17258485, EPI_ISL_17258486, EPI_ISL_17258487, EPI_ISL_17258488, EPI_ISL_17258489, EPI_ISL_17258490, EPI_ISL_17258491, EPI_ISL_17258492, EPI_ISL_17258493, EPI_ISL_17258494, EPI_ISL_17258495, EPI_ISL_17258496, EPI_ISL_17258497, EPI_ISL_17258498, EPI_ISL_17258499, EPI_ISL_17258500, EPI_ISL_17258501, EPI_ISL_17258502, EPI_ISL_17258503, EPI_ISL_17258504, EPI_ISL_17258505, EPI_ISL_17258506, EPI_ISL_17258507, EPI_ISL_17258509, EPI_ISL_17258510, EPI_ISL_17258511, EPI_ISL_17258512, EPI_ISL_17258513, EPI_ISL_17258514, EPI_ISL_17258515, EPI_ISL_17258516, EPI_ISL_17258518, EPI_ISL_17258519, EPI_ISL_17258520, EPI_ISL_17258521, EPI_ISL_17258523, EPI_ISL_17258524, EPI_ISL_17258525, EPI_ISL_17258526 | | | |
| see above | Respiratory Virus Unit / Reference Microbiology Services / UK Health Security Agency | Reference Microbiology Services / UK Health Security Agency | Zambon M. Talts T. Mosscrop L. Miah S |
| EPI_ISL_17276532, EPI_ISL_17276533, EPI_ISL_17276534, EPI_ISL_17276535, EPI_ISL_17276536, EPI_ISL_17276537, EPI_ISL_17276539, EPI_ISL_17276540, EPI_ISL_17276543, EPI_ISL_17276544, EPI_ISL_17276545, EPI_ISL_17276547, EPI_ISL_17276549, EPI_ISL_17276550, EPI_ISL_17276551, EPI_ISL_17276552, EPI_ISL_17276554, EPI_ISL_17276557, EPI_ISL_17276560, EPI_ISL_17276561, EPI_ISL_17276562, EPI_ISL_17276563, EPI_ISL_17276565, EPI_ISL_17276569, EPI_ISL_17276571, EPI_ISL_17276572, EPI_ISL_17276574, EPI_ISL_17276575, EPI_ISL_17276576, EPI_ISL_17276577, EPI_ISL_17276578, EPI_ISL_17276579, EPI_ISL_17276580, EPI_ISL_17276581, EPI_ISL_17276582, EPI_ISL_17276583, EPI_ISL_17276584, EPI_ISL_17276585, EPI_ISL_17276586, EPI_ISL_17276587, EPI_ISL_17276589, EPI_ISL_17276590, EPI_ISL_17276591, EPI_ISL_17276592, EPI_ISL_17276593, EPI_ISL_17276594, EPI_ISL_17276595, EPI_ISL_17276596, EPI_ISL_17276597, EPI_ISL_17276598, EPI_ISL_17276599 | | | |
| see above | Respiratory Virus Unit , Reference Microbiology Services , UK Health Security Agency | Reference Microbiology Services , UK Health Security Agency | Zambon M. Talts T. Mosscrop L. Miah S |
| EPI_ISL_17276840, EPI_ISL_17276841, EPI_ISL_17276842, EPI_ISL_17276843, EPI_ISL_17276844, EPI_ISL_17276845, EPI_ISL_17276846, EPI_ISL_17276847, EPI_ISL_17276848, EPI_ISL_17276849, EPI_ISL_17276850, EPI_ISL_17276851, EPI_ISL_17276852, EPI_ISL_17276853, EPI_ISL_17276854, EPI_ISL_17276855, EPI_ISL_17276856, EPI_ISL_17276857, EPI_ISL_17276858, EPI_ISL_17276859, EPI_ISL_17276860, EPI_ISL_17276861, EPI_ISL_17276862, EPI_ISL_17276863, EPI_ISL_17276864, EPI_ISL_17276865, EPI_ISL_17276866, EPI_ISL_17276867, EPI_ISL_17276868, EPI_ISL_17276869, EPI_ISL_17276870, EPI_ISL_17276871, EPI_ISL_17276873, EPI_ISL_17276874, EPI_ISL_17276875, EPI_ISL_17276876, EPI_ISL_17276877, EPI_ISL_17276878, EPI_ISL_17276879, EPI_ISL_17276880, EPI_ISL_17276881, EPI_ISL_17276882, EPI_ISL_17276883, EPI_ISL_17276884, EPI_ISL_17276885, EPI_ISL_17276886, EPI_ISL_17276887, EPI_ISL_17276888, EPI_ISL_17276889, EPI_ISL_17276890, EPI_ISL_17276891, EPI_ISL_17276892, EPI_ISL_17276893, EPI_ISL_17276894, EPI_ISL_17276895, EPI_ISL_17276896, EPI_ISL_17276897, EPI_ISL_17276899, EPI_ISL_17276900, EPI_ISL_17276901, EPI_ISL_17276902, EPI_ISL_17276903, EPI_ISL_17276904, EPI_ISL_17276905, EPI_ISL_17276906, EPI_ISL_17276908, EPI_ISL_17276909, EPI_ISL_17276910, EPI_ISL_17276911, EPI_ISL_17276912, EPI_ISL_17276913, EPI_ISL_17276914, EPI_ISL_17276915, EPI_ISL_17287519, EPI_ISL_17287520, EPI_ISL_17287521, EPI_ISL_17287522, EPI_ISL_17287523, EPI_ISL_17287525, EPI_ISL_17287526 | | | |
| see above | Respiratory Virus Unit, Reference Microbiology Services, UK Health Security Agency | Reference Microbiology Services, UK Health Security Agency | Zambon M. Talts T. Mosscrop L. Miah S |
| EPI_ISL_17297218, EPI_ISL_17297219, EPI_ISL_17297220, EPI_ISL_17297221, EPI_ISL_17297223, EPI_ISL_17297224, EPI_ISL_17297226, EPI_ISL_17297229, EPI_ISL_17297230, EPI_ISL_17297232, EPI_ISL_17297235, EPI_ISL_17297236, EPI_ISL_17297237, EPI_ISL_17297238, EPI_ISL_17297239, EPI_ISL_17297240, EPI_ISL_17297241, EPI_ISL_17297243, EPI_ISL_17297244, EPI_ISL_17297245, EPI_ISL_17297246, EPI_ISL_17297248, EPI_ISL_17297249, EPI_ISL_17297250, EPI_ISL_17297251, EPI_ISL_17297252 | | | |
| see above | Respiratory Virus Unit / Reference Microbiology Services / UK Health Security Agency | Reference Microbiology Services / UK Health Security Agency | Zambon M. Talts T. Mosscrop L. Miah S |
| EPI_ISL_17368046, EPI_ISL_17368047, EPI_ISL_17368048, EPI_ISL_17368049 | Rahima Moosa | National Institute for Communicable Diseases of the National Health Laboratory Service | Everatt J, Kekana D, Mahlangu B, Stock N, Ntozini B, Ntuli N, Mnguni A, Nzimande A, Ismail A, Bhiman JN, Wolter N |
| EPI_ISL_17368050, EPI_ISL_17368051 | Red Cross | National Institute for Communicable Diseases of the National Health Laboratory Service | Everatt J, Kekana D, Mahlangu B, Stock N, Ntozini B, Ntuli N, Mnguni A, Nzimande A, Ismail A, Bhiman JN, Wolter N |
| EPI_ISL_17368052 | Matikwane | National Institute for Communicable Diseases of the National Health Laboratory Service | Everatt J, Kekana D, Mahlangu B, Stock N, Ntozini B, Ntuli N, Mnguni A, Nzimande A, Ismail A, Bhiman JN, Wolter N |
| EPI_ISL_17368054 | Red Cross | National Institute for Communicable Diseases of the National Health Laboratory Service | Everatt J, Kekana D, Mahlangu B, Stock N, Ntozini B, Ntuli N, Mnguni A, Nzimande A, Ismail A, Bhiman JN, Wolter N |
| EPI_ISL_17417577, EPI_ISL_17417587, EPI_ISL_17417588, EPI_ISL_17417590, EPI_ISL_17417593, EPI_ISL_17417605 | Centers for Disease Control and Prevention (CDC) | Centers for Disease Control and Prevention (CDC) | Wang,L., Lumandas,M.U., Arguelles,V.L., Morin,J., Aggarwal,M., Tatusov,R. and Vega,E. |

| | | | |
|---|---|---|---|
| EPI_ISL_17481632, EPI_ISL_17481636, EPI_ISL_17481649, EPI_ISL_17481652, EPI_ISL_17481653, EPI_ISL_17481654, EPI_ISL_17481658, EPI_ISL_17481660, EPI_ISL_17481662 | National Public Health Institute of Slovakia | Laboratory of Genomics and Bioinformatics, Comenius University Science Park | Szemes,Tomas; Kalinakova,Anna; Kotvasova,Barbora; Sevcikova,Lucia; Vrablova,Terezia; Sedlackova,Tatiana; Rusnakova,Diana; Bohmer,Miroslav; Budis,Jaroslav; Styk,Jakub; Lipkova,Nikola; Forgacova,Michaela; Bokorova,Silvia; Misenko,Pavol |
| EPI_ISL_17559357 | Edendale | National Institute for Communicable Diseases of the National Health Laboratory Service | Everatt J, Kekana D, Mahlangu B, Stock N, Ntozini B, Ntuli N, Mnguni A, Nzimande A, Ismail A, Bhiman JN, Wolter N |
| EPI_ISL_17559359, EPI_ISL_17559361 | Red Cross | National Institute for Communicable Diseases of the National Health Laboratory Service | Everatt J, Kekana D, Mahlangu B, Stock N, Ntozini B, Ntuli N, Mnguni A, Nzimande A, Ismail A, Bhiman JN, Wolter N |
| EPI_ISL_17559407 | Agincourt clinic | National Institute for Communicable Diseases of the National Health Laboratory Service | Everatt J, Kekana D, Mahlangu B, Stock N, Ntozini B, Ntuli N, Mnguni A, Nzimande A, Ismail A, Bhiman JN, Wolter N |
| EPI_ISL_17559419, EPI_ISL_17559450 | Servicio de Microbiologia, Hospital Universitario Virgen del Rocio, Sevilla, Spain | Plataforma de Medicina Computacional, Fundación Progreso y Salud | José Antonio Lepe, Javier Pérez-Florido, María Lara Jiménez |
| EPI_ISL_17559469 | Laboratory of Respiratory Viruses and Measles, Oswaldo Cruz Institute, FIOCRUZ | Laboratorio de Alta Complexidade do Instituto Fernandes Figueira - LACIFF | Paola Resende, Fernando Motta, Elisa Cavalcante Pereira, Bruna Mendonça da Silva, Jéssica Graça Macedo de Carvalho, Larissa Macedo Pinto, Victor Guimaraes, Letícia Lima, Letícia Scalioni, Marilda Siqueira on behalf of the Fiocruz COVID-19 Genomic Surveillance Network |
| EPI_ISL_17583146, EPI_ISL_17583147, EPI_ISL_17583148, EPI_ISL_17583149, EPI_ISL_17583150, EPI_ISL_17583151, EPI_ISL_17583152, EPI_ISL_17583153, EPI_ISL_17583312, EPI_ISL_17583313, EPI_ISL_17583314 | | | |
| see above | West of Scotland Specialist Virology Centre | West of Scotland Specialist Virology Centre | Lynne Ferguson, Imogen Johnston-Menzies, Rory Gunson |
| EPI_ISL_17597842 | Laboratório de Virologia - Instituto Nacional de Saúde | Centers for Disease Control and Prevention - United States of America, Laboratório de Virologia - Instituto Nacional de Saúde | Almiro Tivane, Everardo Vega, Amanda Smith, Lijuan Wang, Neuza Nguenha, Loira Machalele, Sadia Alí, Délcio Muteto, Mirela Pale, Aunésia Marrurele, Félix Gundane, Judite Salência |
| EPI_ISL_17597871, EPI_ISL_17597881, EPI_ISL_17597898 | Laboratório de Virologia - Instituto Nacional de Saúde | Laboratório de Vírus Respiratórios e do Sarampo - Instituto Oswaldo Cruz, Laboratório de Virologia - Instituto Nacional de Saúde | Almiro Tivane, Paola Resende, Neuza Nguenha, Loira Machalele, Sadia Alí, Délcio Muteto, Aunésia Marrurele, Mirela Pale, Félix Gundane, Judite Salência, Marilda Siqueira |
| EPI_ISL_1760383, EPI_ISL_1760384, EPI_ISL_1760385, EPI_ISL_1760386, EPI_ISL_1760387, EPI_ISL_1760388, EPI_ISL_1760389, EPI_ISL_1760390, EPI_ISL_1760391, EPI_ISL_1760392, EPI_ISL_1760393, EPI_ISL_1760394, EPI_ISL_1760395, EPI_ISL_1760396, EPI_ISL_1760397, EPI_ISL_1760398, EPI_ISL_1760399, EPI_ISL_1760400, EPI_ISL_1760401, EPI_ISL_1760402, EPI_ISL_1760403, EPI_ISL_1760404, EPI_ISL_1760405, EPI_ISL_1760406, EPI_ISL_1760407, EPI_ISL_1760408, EPI_ISL_1760409, EPI_ISL_1760410, EPI_ISL_1760411, EPI_ISL_1760412, EPI_ISL_1760413, EPI_ISL_1760414, EPI_ISL_1760415, EPI_ISL_1760416, EPI_ISL_1760417, EPI_ISL_1760418, EPI_ISL_1760419, EPI_ISL_1760420, EPI_ISL_1760421, EPI_ISL_1760422, EPI_ISL_1760423, EPI_ISL_1760424, EPI_ISL_1760425, EPI_ISL_1760426, EPI_ISL_1760427, EPI_ISL_1760428, EPI_ISL_1760429, EPI_ISL_1760430, EPI_ISL_1760431, EPI_ISL_1760432, EPI_ISL_1760433, EPI_ISL_1760434, EPI_ISL_1760435, EPI_ISL_1760436, EPI_ISL_1760437, EPI_ISL_1760438, EPI_ISL_1760439, EPI_ISL_1760440, EPI_ISL_1760441, EPI_ISL_1760442, EPI_ISL_1760443, EPI_ISL_1760444 | | | |
| see above | Royal Children's Hospital, 50 Flemington Rd., Park | WHO Influenza Centre for Reference and Research on Influenza | Angela Todd, Yi-Mo Deng, Annette Alafaci, Naomi Komadina |
| EPI_ISL_17673354, EPI_ISL_17673355, EPI_ISL_17673356, EPI_ISL_17673357, EPI_ISL_17673358, EPI_ISL_17673359, EPI_ISL_17673360, EPI_ISL_17673361, EPI_ISL_17673362, EPI_ISL_17673363, EPI_ISL_17673364 | | | |
| see above | University of Washington, Virology | University of Washington, Virology | Sereewit,J., Hajian,P., Xie,H. and Greninger,A.L. |
| EPI_ISL_17694327, EPI_ISL_17694328, EPI_ISL_17694329, EPI_ISL_17694331, EPI_ISL_17694333, EPI_ISL_17694334, EPI_ISL_17694335, EPI_ISL_17694339, EPI_ISL_17694340, EPI_ISL_17694341, EPI_ISL_17694344 | | | |
| see above | Respiratory Virus Unit / Reference Microbiology Services / UK Health Security Agency | Reference Microbiology Services / UK Health Security Agency | Zambon M. Talts T. Mosscrop L. Miah S |
| EPI_ISL_17773663, EPI_ISL_17773664 | Universidade Federal de São Paulo, Departamento de Medicina, Disciplina de Doenças Infecciosas e Parasitárias. Laboratório de Virologia | Laboratory of Respiratory Viruses and Measles, Oswaldo Cruz Institute, FIOCRUZ | Paola Resende, Fernando Motta, Elisa Cavalcante Pereira, Leticia Ferreira Lima, Bruna Mendonça da Silva, Jéssica Graça Macedo de Carvalho, Larissa Macedo Pinto, Victor Guimaraes, Leticia Scalioni, Rodrigo Ribeiro Rodrigues, Nancy Bellei, Marilda Siqueira on behalf of the Fiocruz COVID-19 Genomic Surveillance Network |
| EPI_ISL_17778084 | Laboratorio Central de Saude Publica do Distrito Federal (LACEN/DF) | Instituto Oswaldo Cruz FIOCRUZ - Laboratory of Respiratory Viruses and Measles (LVRS) | Paola Resende, Fernando Motta, Elisa Cavalcante Pereira, Leticia Ferreira Lima, Bruna Mendonça da Silva, Jéssica Graça Macedo de Carvalho, Larissa Macedo Pinto, Victor Guimaraes, Leticia Scalioni, Rodrigo Ribeiro Rodrigues, Marilda Siqueira on behalf of the Fiocruz COVID-19 Genomic Surveillance Network |
| EPI_ISL_17778104 | Universidade Federal de São Paulo, Departamento de Medicina, Disciplina de Doenças Infecciosas e Parasitárias. Laboratório de Virologia | Instituto Oswaldo Cruz FIOCRUZ - Laboratory of Respiratory Viruses and Measles (LVRS) | Paola Resende, Fernando Motta, Elisa Cavalcante Pereira, Leticia Ferreira Lima, Bruna Mendonça da Silva, Jéssica Graça Macedo de Carvalho, Larissa Macedo Pinto, Victor Guimaraes, Leticia Scalioni, Rodrigo Ribeiro Rodrigues, Marilda Siqueira on behalf of the Fiocruz COVID-19 Genomic Surveillance Network |
| EPI_ISL_17782656, EPI_ISL_17782658 | Kitasato University, Infection Control and Immunology | Kitasato University, Infection Control and Immunology | Ito,T., Sawada,A., Saito,A., Ishikura,K., Kawashima,H., Nakayama,T. and Katayama,K. |
| EPI_ISL_17808731, EPI_ISL_17808732, EPI_ISL_17808733, EPI_ISL_17808734 | Central public health laboratory of the state of Amapa | Evandro Chagas Institute, Laboratory of Respiratory Viruses, National InfluenzaCenter | Mirleide Santos; Fernando Tavares; Edivaldo Junior; Luana Barbagelata; Amanda Mendes; Wanderley Dias; Delana Melo; Agata Monique; Edvaldo Penha; |
| EPI_ISL_17808838, EPI_ISL_17808839, EPI_ISL_17808840, EPI_ISL_17808841, EPI_ISL_17808842, EPI_ISL_17808843, EPI_ISL_17808844, EPI_ISL_17808845, EPI_ISL_17808847, EPI_ISL_17808848, EPI_ISL_17808849, EPI_ISL_17808850, EPI_ISL_17808851, EPI_ISL_17808852, EPI_ISL_17808853, EPI_ISL_17808854, EPI_ISL_17808855, EPI_ISL_17808856, EPI_ISL_17808858, EPI_ISL_17808859, EPI_ISL_17808860, EPI_ISL_17808861, EPI_ISL_17808862 | | | |
| see above | Valley Wise Health | Arizona State University | Holland, LaRinda A.; Holland, Steven C.; Smith, Matthew F.; Leonard, Victoria R.; Murugan, Vel; Nordstrom, Lora; Mulrow, Mary; Salgado, Raquel; White, Michael; Lim, Efrem S. |
| EPI_ISL_17950136, EPI_ISL_17950160, EPI_ISL_17950161, EPI_ISL_17950162, EPI_ISL_17950168, EPI_ISL_17950169, EPI_ISL_17950170, EPI_ISL_17950181, EPI_ISL_17950182, EPI_ISL_17950184, EPI_ISL_17950185, EPI_ISL_17950187, EPI_ISL_17950190, EPI_ISL_17950195, EPI_ISL_17950198, EPI_ISL_17950200, EPI_ISL_17950201, EPI_ISL_17950207, EPI_ISL_17950210, EPI_ISL_17950215, EPI_ISL_17950227, EPI_ISL_17950231, EPI_ISL_17950236 | | | |
| see above | Arizona State University | Arizona State University | Holland,S.C., Holland,L.A., Smith,M.F., Leonard,V.R., Murugan,V., Nordstrom,L., Mulrow,M., Salgado,R., White,M. and Lim,E.S. |
| EPI_ISL_17951047, EPI_ISL_17951048, EPI_ISL_17951049, EPI_ISL_17951050 | Molecular Biology Laboratory, Pedro de Elizalde Hospital | Virology Laboratory, Ricardo Gutiérrez Children's Hospital | Acuña, Dolores; Goya, Stephanie; Nabaes Jodar, Mercedes S.; Montoto, L; Wenk, G; Sevilla, ME; Miño, L; Valeri, C; Bokser, V; Mistchenko, Alicia S.; Viegas, Mariana |
| EPI_ISL_17964351, EPI_ISL_17964353, EPI_ISL_17964354, EPI_ISL_17964357, EPI_ISL_17964358, EPI_ISL_17964361, EPI_ISL_17964362, EPI_ISL_17964364, EPI_ISL_17964376 | University of Zambia Medical School | Department of Medical Microbiology, University Medical Centre Utrecht | Annefleur C. Langedijk, Bram Vrancken, Robert Jan Lebbink, Rachel C. Pieciak, Philippe Lemey, Louis J. Bont, Christopher J Gill |
| EPI_ISL_17992500 | The South African Red Cross Society - Western Cape Provincial Office | National Institute for Communicable Diseases of the National Health Laboratory Service | Everatt J, Kekana D, Mahlangu B, Stock N, Ntozini B, Ntuli N, Mnguni A, Nzimande A, Ismail A, Bhiman JN, Wolter N |
| EPI_ISL_17992511 | Edendale Hospital | National Institute for Communicable Diseases of the National Health Laboratory Service | Everatt J, Kekana D, Mahlangu B, Stock N, Ntozini B, Ntuli N, Mnguni A, Nzimande A, Ismail A, Bhiman JN, Wolter N |
| EPI_ISL_17998372, EPI_ISL_17998373 | Dr. Sadick Saban's office | National Institute for Communicable Diseases of the National Health Laboratory Service | Everatt J, Kekana D, Mahlangu B, Stock N, Ntozini B, Ntuli N, Mnguni A, Nzimande A, Ismail A, Bhiman JN, Wolter N |
| EPI_ISL_17998374 | Dr Claire Castelyn's office | National Institute for Communicable Diseases of the National Health Laboratory Service | Everatt J, Kekana D, Mahlangu B, Stock N, Ntozini B, Ntuli N, Mnguni A, Nzimande A, Ismail A, Bhiman JN, Wolter N |
| EPI_ISL_17998375 | Dr. Sadick Saban's office | National Institute for Communicable Diseases of the National Health Laboratory Service | Everatt J, Kekana D, Mahlangu B, Stock N, Ntozini B, Ntuli N, Mnguni A, Nzimande A, Ismail A, Bhiman JN, Wolter N |
| EPI_ISL_17998376 | Dr Aysha Kola Adam's office | National Institute for Communicable Diseases of the National Health Laboratory Service | Everatt J, Kekana D, Mahlangu B, Stock N, Ntozini B, Ntuli N, Mnguni A, Nzimande A, Ismail A, Bhiman JN, Wolter N |
| EPI_ISL_17998378 | Dr. Sadick Saban's office | National Institute for Communicable Diseases of the National Health Laboratory Service | Everatt J, Kekana D, Mahlangu B, Stock N, Ntozini B, Ntuli N, Mnguni A, Nzimande A, Ismail A, Bhiman JN, Wolter N |
| EPI_ISL_17998379 | Dr. PDC Erasmus' office | National Institute for Communicable Diseases of the National Health Laboratory Service | Everatt J, Kekana D, Mahlangu B, Stock N, Ntozini B, Ntuli N, Mnguni A, Nzimande A, Ismail A, Bhiman JN, Wolter N |
| EPI_ISL_18005798, EPI_ISL_18005799, EPI_ISL_18005800, EPI_ISL_18005801, EPI_ISL_18005802, EPI_ISL_18005803, EPI_ISL_18005804, EPI_ISL_18005805, EPI_ISL_18005806, EPI_ISL_18005807, EPI_ISL_18005808, EPI_ISL_18005809, EPI_ISL_18005810, EPI_ISL_18005811, EPI_ISL_18005812, EPI_ISL_18005813, EPI_ISL_18005814, EPI_ISL_18005815, EPI_ISL_18005816, EPI_ISL_18005817, EPI_ISL_18005818, EPI_ISL_18005819, EPI_ISL_18005820, EPI_ISL_18005821, EPI_ISL_18005822, EPI_ISL_18005823, EPI_ISL_18005824, EPI_ISL_18005825, EPI_ISL_18005826, EPI_ISL_18005827, EPI_ISL_18005828, EPI_ISL_18005829, EPI_ISL_18005830, EPI_ISL_18005831, EPI_ISL_18005832, EPI_ISL_18005833, EPI_ISL_18005834, EPI_ISL_18005835, EPI_ISL_18005836, EPI_ISL_18005837, EPI_ISL_18005838, EPI_ISL_18005839, EPI_ISL_18005840, EPI_ISL_18005841, EPI_ISL_18005842, EPI_ISL_18005843, EPI_ISL_18005844, EPI_ISL_18005845, EPI_ISL_18005846, EPI_ISL_18005847, EPI_ISL_18005848, EPI_ISL_18005849, EPI_ISL_18005850, EPI_ISL_18005851, EPI_ISL_18005852, EPI_ISL_18005853, EPI_ISL_18005854, EPI_ISL_18005855, EPI_ISL_18005856, EPI_ISL_18005857, EPI_ISL_18005858, EPI_ISL_18005859, EPI_ISL_18005860, EPI_ISL_18005861, EPI_ISL_18005862, EPI_ISL_18005863, EPI_ISL_18005864, EPI_ISL_18005865, EPI_ISL_18005866, EPI_ISL_18005867, EPI_ISL_18005868, EPI_ISL_18005869, EPI_ISL_18005870, EPI_ISL_18005871, EPI_ISL_18005872, EPI_ISL_18005873, EPI_ISL_18005874, EPI_ISL_18005875, EPI_ISL_18005876, EPI_ISL_18005877, EPI_ISL_18005878, EPI_ISL_18005879, EPI_ISL_18005880, EPI_ISL_18005881, EPI_ISL_18005882, EPI_ISL_18005883, EPI_ISL_18005884, EPI_ISL_18005885, EPI_ISL_18005886, EPI_ISL_18005887, EPI_ISL_18005888, EPI_ISL_18005889, EPI_ISL_18005890, EPI_ISL_18005891, EPI_ISL_18005892, EPI_ISL_18005893, EPI_ISL_18005894, EPI_ISL_18005895, EPI_ISL_18005896, EPI_ISL_18005897, EPI_ISL_18005898, EPI_ISL_18005899, EPI_ISL_18005900, EPI_ISL_18005901, EPI_ISL_18005902, EPI_ISL_18005903, EPI_ISL_18005904, EPI_ISL_18005905, EPI_ISL_18005906, EPI_ISL_18005907, EPI_ISL_18005908, EPI_ISL_18005909, EPI_ISL_18005910, EPI_ISL_18005911, EPI_ISL_18005912, EPI_ISL_18005913, EPI_ISL_18005914, EPI_ISL_18005915, EPI_ISL_18005916, EPI_ISL_18005917, EPI_ISL_18005918, EPI_ISL_18005919, EPI_ISL_18005920, EPI_ISL_18005921, EPI_ISL_18005922, EPI_ISL_18005923, EPI_ISL_18005924, EPI_ISL_18005925, EPI_ISL_18005926, EPI_ISL_18005927, EPI_ISL_18005928, EPI_ISL_18005929, EPI_ISL_18005930, EPI_ISL_18005931, EPI_ISL_18005932, EPI_ISL_18005933, EPI_ISL_18005934, EPI_ISL_18005935, EPI_ISL_18005936, EPI_ISL_18005937, EPI_ISL_18005938, EPI_ISL_18005939, EPI_ISL_18005940, EPI_ISL_18005941, EPI_ISL_18005942, EPI_ISL_18005943, EPI_ISL_18005944, EPI_ISL_18005945, EPI_ISL_18005946, EPI_ISL_18005947, EPI_ISL_18005948, EPI_ISL_18005949, EPI_ISL_18005950, EPI_ISL_18005951, EPI_ISL_18005952 | | | |
| see above | Robert Koch-Institute Nationales Referenzzentrum für Influenza | Robert Koch-Institute Nationales Referenzzentrum für Influenza | Sophie Kondgen, Janine Reiche |
| EPI_ISL_18042405, EPI_ISL_18042406, EPI_ISL_18042407, EPI_ISL_18042408 | Respiratory Virus Unit / Reference Microbiology Services / UK Health Security Agency | Reference Microbiology Services / UK Health Security Agency | Zambon M. Talts T. Kele B. Miah S |
| EPI_ISL_18042523, EPI_ISL_18042524, EPI_ISL_18042526, EPI_ISL_18042527, EPI_ISL_18042528, EPI_ISL_18042531, EPI_ISL_18042532, EPI_ISL_18042533, EPI_ISL_18042535, EPI_ISL_18042536, EPI_ISL_18042537, EPI_ISL_18042539, EPI_ISL_18042540, EPI_ISL_18042541, EPI_ISL_18042542, EPI_ISL_18042545, EPI_ISL_18042548, EPI_ISL_18042550, EPI_ISL_18042551, EPI_ISL_18042553, EPI_ISL_18042555, EPI_ISL_18042556, EPI_ISL_18042557, EPI_ISL_18042558, EPI_ISL_18042560, EPI_ISL_18042561, EPI_ISL_18042562, EPI_ISL_18042563, EPI_ISL_18042564, EPI_ISL_18042567, EPI_ISL_18042568, EPI_ISL_18042569 | | | |
| see above | Research Institute for Tropical Medicine, Department of Health Compound, Virology Section | WHO Collaborating Centre for Reference and Research on Influenza | Jonjee Morin, Catherine Dacasin, Vina Lea Arguelles, Xiaomin Dong, Steven Edwards, Yi-Mo Deng, Clyde Dapat, Ian Barr |
| EPI_ISL_18090572, EPI_ISL_18090573, EPI_ISL_18090576, EPI_ISL_18090579, EPI_ISL_18090581, EPI_ISL_18090583, EPI_ISL_18090584, EPI_ISL_18090588, EPI_ISL_18090589, EPI_ISL_18090590, EPI_ISL_18090592, EPI_ISL_18090600, EPI_ISL_18090601, EPI_ISL_18090603, EPI_ISL_18090605, EPI_ISL_18090609, EPI_ISL_18090610, EPI_ISL_18090613, EPI_ISL_18090615, EPI_ISL_18090617, EPI_ISL_18090621, EPI_ISL_18090627, EPI_ISL_18090628, EPI_ISL_18090630, EPI_ISL_18090631, EPI_ISL_18090633, EPI_ISL_18090634, EPI_ISL_18090635, EPI_ISL_18090638, EPI_ISL_18090639, EPI_ISL_18090640, EPI_ISL_18090641, EPI_ISL_18090644, EPI_ISL_18090649, EPI_ISL_18090655, EPI_ISL_18090659, EPI_ISL_18090668, EPI_ISL_18090669, EPI_ISL_18090670, EPI_ISL_18090672, EPI_ISL_18090673, EPI_ISL_18090674, EPI_ISL_18090675, EPI_ISL_18090681, EPI_ISL_18090683, EPI_ISL_18090685, EPI_ISL_18090687, EPI_ISL_18090688, EPI_ISL_18090689, EPI_ISL_18090691, EPI_ISL_18090693 | | | |
| see above | Westmead Institute for Medical Research & Sydney Infectious Disease Institute | Westmead Institute for Medical Research & Sydney Infectious Disease Institute | Pangesti,K.N.A., Ansari,H.R., Bayoumi,A., Kesson,A.M., Hill-Cawthorne,G.A. and Abd El Ghany,M. |

| | | | |
|---|---|---|---|
| EPI_ISL_18094393 | Department of Virology, National Institute of Health, Islamabad, Pakistan | Department of Virology, National Institute of Health, Islamabad, Pakistan | Massab Umair, Syed Adnan Haider, Muhammad Ammar, Zunera Jamal, and Muhammad Salman |
| EPI_ISL_18143446, EPI_ISL_18143448 | The Brotman Baty Institute for Precision Medicine | The Brotman Baty Institute for Precision Medicine | Frazar,C.D., Lee,J., Gamboa,L., McDermot,E., Stone,J., Kolar,T., Han,P.D., Sibley,T.R., Truong,M., Reinhart,D., Wolf,C.R., Boeckh,M., Englund,J.A., Lutz,B.R., Waghmare,A., Viboud,C., Starita,L.M., Shendure,J., Bedford,T. and Chu,H.Y. |
| EPI_ISL_18143450 | The Brotman Baty Institute for Precision Medicine | The Brotman Baty Institute for Precision Medicine | Frazar,C.D., Lee,J., Ryke,E., Gamboa,L., McDermot,E., Stone,J., Kolar,T., Han,P.D., Sibley,T.R., Truong,M., Reinhart,D., Wolf,C.R., Duchin,J., Boeckh,M., Englund,J.A., Lutz,B.R., Waghmare,A., Viboud,C., Starita,L.M., Chu,H.Y., Bedford,T. and Shendure,J. |
| EPI_ISL_18143452 | The Brotman Baty Institute for Precision Medicine | The Brotman Baty Institute for Precision Medicine | Frazar,C.D., Lee,J., Gamboa,L., McDermot,E., Stone,J., Kolar,T., Han,P.D., Sibley,T.R., Truong,M., Reinhart,D., Wolf,C.R., Boeckh,M., Englund,J.A., Lutz,B.R., Waghmare,A., Viboud,C., Starita,L.M., Shendure,J., Bedford,T. and Chu,H.Y. |
| EPI_ISL_18143453 | The Brotman Baty Institute for Precision Medicine | The Brotman Baty Institute for Precision Medicine | Frazar,C.D., Lee,J., Ryke,E., Gamboa,L., McDermot,E., Stone,J., Kolar,T., Han,P.D., Sibley,T.R., Truong,M., Reinhart,D., Wolf,C.R., Duchin,J., Boeckh,M., Englund,J.A., Lutz,B.R., Waghmare,A., Viboud,C., Starita,L.M., Chu,H.Y., Bedford,T. and Shendure,J. |
| EPI_ISL_18143455, EPI_ISL_18143458, EPI_ISL_18143459, EPI_ISL_18143461 | The Brotman Baty Institute for Precision Medicine | The Brotman Baty Institute for Precision Medicine | Frazar,C.D., Lee,J., Gamboa,L., McDermot,E., Stone,J., Kolar,T., Han,P.D., Sibley,T.R., Truong,M., Reinhart,D., Wolf,C.R., Boeckh,M., Englund,J.A., Lutz,B.R., Waghmare,A., Viboud,C., Starita,L.M., Shendure,J., Bedford,T. and Chu,H.Y. |
| EPI_ISL_18143462, EPI_ISL_18143466, EPI_ISL_18143468, EPI_ISL_18143470, EPI_ISL_18143473 | The Brotman Baty Institute for Precision Medicine | The Brotman Baty Institute for Precision Medicine | Frazar,C.D., Lee,J., Ryke,E., Gamboa,L., McDermot,E., Stone,J., Kolar,T., Han,P.D., Sibley,T.R., Truong,M., Reinhart,D., Wolf,C.R., Duchin,J., Boeckh,M., Englund,J.A., Lutz,B.R., Waghmare,A., Viboud,C., Starita,L.M., Chu,H.Y., Bedford,T. and Shendure,J. |
| EPI_ISL_18143474 | The Brotman Baty Institute for Precision Medicine | The Brotman Baty Institute for Precision Medicine | Frazar,C.D., Lee,J., Gamboa,L., McDermot,E., Stone,J., Kolar,T., Han,P.D., Sibley,T.R., Truong,M., Reinhart,D., Wolf,C.R., Boeckh,M., Englund,J.A., Lutz,B.R., Waghmare,A., Viboud,C., Starita,L.M., Shendure,J., Bedford,T. and Chu,H.Y. |
| EPI_ISL_18143475, EPI_ISL_18143481, EPI_ISL_18143482 | The Brotman Baty Institute for Precision Medicine | The Brotman Baty Institute for Precision Medicine | Frazar,C.D., Lee,J., Ryke,E., Gamboa,L., McDermot,E., Stone,J., Kolar,T., Han,P.D., Sibley,T.R., Truong,M., Reinhart,D., Wolf,C.R., Duchin,J., Boeckh,M., Englund,J.A., Lutz,B.R., Waghmare,A., Viboud,C., Starita,L.M., Chu,H.Y., Bedford,T. and Shendure,J. |
| EPI_ISL_18143485 | The Brotman Baty Institute for Precision Medicine | The Brotman Baty Institute for Precision Medicine | Frazar,C.D., Lee,J., Gamboa,L., McDermot,E., Stone,J., Kolar,T., Han,P.D., Sibley,T.R., Truong,M., Reinhart,D., Wolf,C.R., Boeckh,M., Englund,J.A., Lutz,B.R., Waghmare,A., Viboud,C., Starita,L.M., Shendure,J., Bedford,T. and Chu,H.Y. |
| EPI_ISL_18143487, EPI_ISL_18143488 | The Brotman Baty Institute for Precision Medicine | The Brotman Baty Institute for Precision Medicine | Frazar,C.D., Lee,J., Ryke,E., Gamboa,L., McDermot,E., Stone,J., Kolar,T., Han,P.D., Sibley,T.R., Truong,M., Reinhart,D., Wolf,C.R., Duchin,J., Boeckh,M., Englund,J.A., Lutz,B.R., Waghmare,A., Viboud,C., Starita,L.M., Chu,H.Y., Bedford,T. and Shendure,J. |
| EPI_ISL_18143491 | The Brotman Baty Institute for Precision Medicine | The Brotman Baty Institute for Precision Medicine | Frazar,C.D., Lee,J., Gamboa,L., McDermot,E., Stone,J., Kolar,T., Han,P.D., Sibley,T.R., Truong,M., Reinhart,D., Wolf,C.R., Boeckh,M., Englund,J.A., Lutz,B.R., Waghmare,A., Viboud,C., Starita,L.M., Shendure,J., Bedford,T. and Chu,H.Y. |
| EPI_ISL_18143492, EPI_ISL_18143493 | The Brotman Baty Institute for Precision Medicine | The Brotman Baty Institute for Precision Medicine | Frazar,C.D., Lee,J., Ryke,E., Gamboa,L., McDermot,E., Stone,J., Kolar,T., Han,P.D., Sibley,T.R., Truong,M., Reinhart,D., Wolf,C.R., Duchin,J., Boeckh,M., Englund,J.A., Lutz,B.R., Waghmare,A., Viboud,C., Starita,L.M., Chu,H.Y., Bedford,T. and Shendure,J. |
| EPI_ISL_18143498 | The Brotman Baty Institute for Precision Medicine | The Brotman Baty Institute for Precision Medicine | Frazar,C.D., Lee,J., Gamboa,L., McDermot,E., Stone,J., Kolar,T., Han,P.D., Sibley,T.R., Truong,M., Reinhart,D., Wolf,C.R., Boeckh,M., Englund,J.A., Lutz,B.R., Waghmare,A., Viboud,C., Starita,L.M., Shendure,J., Bedford,T. and Chu,H.Y. |
| EPI_ISL_18143501 | The Brotman Baty Institute for Precision Medicine | The Brotman Baty Institute for Precision Medicine | Frazar,C.D., Lee,J., Ryke,E., Gamboa,L., McDermot,E., Stone,J., Kolar,T., Han,P.D., Sibley,T.R., Truong,M., Reinhart,D., Wolf,C.R., Duchin,J., Boeckh,M., Englund,J.A., Lutz,B.R., Waghmare,A., Viboud,C., Starita,L.M., Chu,H.Y., Bedford,T. and Shendure,J. |
| EPI_ISL_18143504 | The Brotman Baty Institute for Precision Medicine | The Brotman Baty Institute for Precision Medicine | Frazar,C.D., Lee,J., Gamboa,L., McDermot,E., Stone,J., Kolar,T., Han,P.D., Sibley,T.R., Truong,M., Reinhart,D., Wolf,C.R., Boeckh,M., Englund,J.A., Lutz,B.R., Waghmare,A., Viboud,C., Starita,L.M., Shendure,J., Bedford,T. and Chu,H.Y. |
| EPI_ISL_18143505 | The Brotman Baty Institute for Precision Medicine | The Brotman Baty Institute for Precision Medicine | Frazar,C.D., Lee,J., Ryke,E., Gamboa,L., McDermot,E., Stone,J., Kolar,T., Han,P.D., Sibley,T.R., Truong,M., Reinhart,D., Wolf,C.R., Duchin,J., Boeckh,M., Englund,J.A., Lutz,B.R., Waghmare,A., Viboud,C., Starita,L.M., Chu,H.Y., Bedford,T. and Shendure,J. |
| EPI_ISL_18143515, EPI_ISL_18143516 | The Brotman Baty Institute for Precision Medicine | The Brotman Baty Institute for Precision Medicine | Frazar,C.D., Lee,J., Gamboa,L., McDermot,E., Stone,J., Kolar,T., Han,P.D., Sibley,T.R., Truong,M., Reinhart,D., Wolf,C.R., Boeckh,M., Englund,J.A., Lutz,B.R., Waghmare,A., Viboud,C., Starita,L.M., Shendure,J., Bedford,T. and Chu,H.Y. |
| EPI_ISL_18143519, EPI_ISL_18143521, EPI_ISL_18143523, EPI_ISL_18143524, EPI_ISL_18143525 | The Brotman Baty Institute for Precision Medicine | The Brotman Baty Institute for Precision Medicine | Frazar,C.D., Lee,J., Ryke,E., Gamboa,L., McDermot,E., Stone,J., Kolar,T., Han,P.D., Sibley,T.R., Truong,M., Reinhart,D., Wolf,C.R., Duchin,J., Boeckh,M., Englund,J.A., Lutz,B.R., Waghmare,A., Viboud,C., Starita,L.M., Chu,H.Y., Bedford,T. and Shendure,J. |
| EPI_ISL_18143526 | The Brotman Baty Institute for Precision Medicine | The Brotman Baty Institute for Precision Medicine | Frazar,C.D., Lee,J., Gamboa,L., McDermot,E., Stone,J., Kolar,T., Han,P.D., Sibley,T.R., Truong,M., Reinhart,D., Wolf,C.R., Boeckh,M., Englund,J.A., Lutz,B.R., Waghmare,A., Viboud,C., Starita,L.M., Shendure,J., Bedford,T. and Chu,H.Y. |
| EPI_ISL_18143534 | The Brotman Baty Institute for Precision Medicine | The Brotman Baty Institute for Precision Medicine | Frazar,C.D., Lee,J., Ryke,E., Gamboa,L., McDermot,E., Stone,J., Kolar,T., Han,P.D., Sibley,T.R., Truong,M., Reinhart,D., Wolf,C.R., Duchin,J., Boeckh,M., Englund,J.A., Lutz,B.R., Waghmare,A., Viboud,C., Starita,L.M., Chu,H.Y., Bedford,T. and Shendure,J. |
| EPI_ISL_18143535 | The Brotman Baty Institute for Precision Medicine | The Brotman Baty Institute for Precision Medicine | Frazar,C.D., Lee,J., Gamboa,L., McDermot,E., Stone,J., Kolar,T., Han,P.D., Sibley,T.R., Truong,M., Reinhart,D., Wolf,C.R., Boeckh,M., Englund,J.A., Lutz,B.R., Waghmare,A., Viboud,C., Starita,L.M., Shendure,J., Bedford,T. and Chu,H.Y. |
| EPI_ISL_18143541, EPI_ISL_18143543, EPI_ISL_18143545 | The Brotman Baty Institute for Precision Medicine | The Brotman Baty Institute for Precision Medicine | Frazar,C.D., Lee,J., Ryke,E., Gamboa,L., McDermot,E., Stone,J., Kolar,T., Han,P.D., Sibley,T.R., Truong,M., Reinhart,D., Wolf,C.R., Duchin,J., Boeckh,M., Englund,J.A., Lutz,B.R., Waghmare,A., Viboud,C., Starita,L.M., Chu,H.Y., Bedford,T. and Shendure,J. |
| EPI_ISL_18143546, EPI_ISL_18143549, EPI_ISL_18143550, EPI_ISL_18143552 | The Brotman Baty Institute for Precision Medicine | The Brotman Baty Institute for Precision Medicine | Frazar,C.D., Lee,J., Gamboa,L., McDermot,E., Stone,J., Kolar,T., Han,P.D., Sibley,T.R., Truong,M., Reinhart,D., Wolf,C.R., Boeckh,M., Englund,J.A., Lutz,B.R., Waghmare,A., Viboud,C., Starita,L.M., Shendure,J., Bedford,T. and Chu,H.Y. |
| EPI_ISL_18143558, EPI_ISL_18143559 | The Brotman Baty Institute for Precision Medicine | The Brotman Baty Institute for Precision Medicine | Frazar,C.D., Lee,J., Ryke,E., Gamboa,L., McDermot,E., Stone,J., Kolar,T., Han,P.D., Sibley,T.R., Truong,M., Reinhart,D., Wolf,C.R., Duchin,J., Boeckh,M., Englund,J.A., Lutz,B.R., Waghmare,A., Viboud,C., Starita,L.M., Chu,H.Y., Bedford,T. and Shendure,J. |
| EPI_ISL_18143561 | The Brotman Baty Institute for Precision Medicine | The Brotman Baty Institute for Precision Medicine | Frazar,C.D., Lee,J., Gamboa,L., McDermot,E., Stone,J., Kolar,T., Han,P.D., Sibley,T.R., Truong,M., Reinhart,D., Wolf,C.R., Boeckh,M., Englund,J.A., Lutz,B.R., Waghmare,A., Viboud,C., Starita,L.M., Shendure,J., Bedford,T. and Chu,H.Y. |
| EPI_ISL_18143565 | The Brotman Baty Institute for Precision Medicine | The Brotman Baty Institute for Precision Medicine | Frazar,C.D., Lee,J., Ryke,E., Gamboa,L., McDermot,E., Stone,J., Kolar,T., Han,P.D., Sibley,T.R., Truong,M., Reinhart,D., Wolf,C.R., Duchin,J., Boeckh,M., Englund,J.A., Lutz,B.R., Waghmare,A., Viboud,C., Starita,L.M., Chu,H.Y., Bedford,T. and Shendure,J. |
| EPI_ISL_18143567 | The Brotman Baty Institute for Precision Medicine | The Brotman Baty Institute for Precision Medicine | Frazar,C.D., Lee,J., Gamboa,L., McDermot,E., Stone,J., Kolar,T., Han,P.D., Sibley,T.R., Truong,M., Reinhart,D., Wolf,C.R., Boeckh,M., Englund,J.A., Lutz,B.R., Waghmare,A., Viboud,C., Starita,L.M., Shendure,J., Bedford,T. and Chu,H.Y. |
| EPI_ISL_18143568 | The Brotman Baty Institute for Precision Medicine | The Brotman Baty Institute for Precision Medicine | Frazar,C.D., Lee,J., Ryke,E., Gamboa,L., McDermot,E., Stone,J., Kolar,T., Han,P.D., Sibley,T.R., Truong,M., Reinhart,D., Wolf,C.R., Duchin,J., Boeckh,M., Englund,J.A., Lutz,B.R., Waghmare,A., Viboud,C., Starita,L.M., Chu,H.Y., Bedford,T. and Shendure,J. |
| EPI_ISL_18143569, EPI_ISL_18143570 | The Brotman Baty Institute for Precision Medicine | The Brotman Baty Institute for Precision Medicine | Frazar,C.D., Lee,J., Gamboa,L., McDermot,E., Stone,J., Kolar,T., Han,P.D., Sibley,T.R., Truong,M., Reinhart,D., Wolf,C.R., Boeckh,M., Englund,J.A., Lutz,B.R., Waghmare,A., Viboud,C., Starita,L.M., Shendure,J., Bedford,T. and Chu,H.Y. |
| EPI_ISL_18143573 | The Brotman Baty Institute for Precision Medicine | The Brotman Baty Institute for Precision Medicine | Frazar,C.D., Lee,J., Ryke,E., Gamboa,L., McDermot,E., Stone,J., Kolar,T., Han,P.D., Sibley,T.R., Truong,M., Reinhart,D., Wolf,C.R., Duchin,J., Boeckh,M., Englund,J.A., Lutz,B.R., Waghmare,A., Viboud,C., Starita,L.M., Chu,H.Y., Bedford,T. and Shendure,J. |
| EPI_ISL_18143574, EPI_ISL_18143577 | The Brotman Baty Institute for Precision Medicine | The Brotman Baty Institute for Precision Medicine | Frazar,C.D., Lee,J., Gamboa,L., McDermot,E., Stone,J., Kolar,T., Han,P.D., Sibley,T.R., Truong,M., Reinhart,D., Wolf,C.R., Boeckh,M., Englund,J.A., Lutz,B.R., Waghmare,A., Viboud,C., Starita,L.M., Shendure,J., Bedford,T. and Chu,H.Y. |
| EPI_ISL_18143579 | The Brotman Baty Institute for Precision Medicine | The Brotman Baty Institute for Precision Medicine | Frazar,C.D., Lee,J., Ryke,E., Gamboa,L., McDermot,E., Stone,J., Kolar,T., Han,P.D., Sibley,T.R., Truong,M., Reinhart,D., Wolf,C.R., Duchin,J., Boeckh,M., Englund,J.A., Lutz,B.R., Waghmare,A., Viboud,C., Starita,L.M., Chu,H.Y., Bedford,T. and Shendure,J. |
| EPI_ISL_18143581, EPI_ISL_18143584 | The Brotman Baty Institute for Precision Medicine | The Brotman Baty Institute for Precision Medicine | Frazar,C.D., Lee,J., Gamboa,L., McDermot,E., Stone,J., Kolar,T., Han,P.D., Sibley,T.R., Truong,M., Reinhart,D., Wolf,C.R., Boeckh,M., Englund,J.A., Lutz,B.R., Waghmare,A., Viboud,C., Starita,L.M., Shendure,J., Bedford,T. and Chu,H.Y. |
| EPI_ISL_18143586 | The Brotman Baty Institute for Precision Medicine | The Brotman Baty Institute for Precision Medicine | Frazar,C.D., Lee,J., Ryke,E., Gamboa,L., McDermot,E., Stone,J., Kolar,T., Han,P.D., Sibley,T.R., Truong,M., Reinhart,D., Wolf,C.R., Duchin,J., Boeckh,M., Englund,J.A., Lutz,B.R., Waghmare,A., Viboud,C., Starita,L.M., Chu,H.Y., Bedford,T. and Shendure,J. |
| EPI_ISL_18143587 | The Brotman Baty Institute for Precision Medicine | The Brotman Baty Institute for Precision Medicine | Frazar,C.D., Lee,J., Gamboa,L., McDermot,E., Stone,J., Kolar,T., Han,P.D., Sibley,T.R., Truong,M., Reinhart,D., Wolf,C.R., Boeckh,M., Englund,J.A., Lutz,B.R., Waghmare,A., Viboud,C., Starita,L.M., Shendure,J., Bedford,T. and Chu,H.Y. |
| EPI_ISL_18143588 | The Brotman Baty Institute for Precision Medicine | The Brotman Baty Institute for Precision Medicine | Frazar,C.D., Lee,J., Ryke,E., Gamboa,L., McDermot,E., Stone,J., Kolar,T., Han,P.D., Sibley,T.R., Truong,M., Reinhart,D., Wolf,C.R., Duchin,J., Boeckh,M., Englund,J.A., Lutz,B.R., Waghmare,A., Viboud,C., Starita,L.M., Chu,H.Y., Bedford,T. and Shendure,J. |
| EPI_ISL_18143590 | The Brotman Baty Institute for Precision Medicine | The Brotman Baty Institute for Precision Medicine | Frazar,C.D., Lee,J., Gamboa,L., McDermot,E., Stone,J., Kolar,T., Han,P.D., Sibley,T.R., Truong,M., Reinhart,D., Wolf,C.R., Boeckh,M., Englund,J.A., Lutz,B.R., Waghmare,A., Viboud,C., Starita,L.M., Shendure,J., Bedford,T. and Chu,H.Y. |
| EPI_ISL_18143593, EPI_ISL_18143594, EPI_ISL_18143595 | The Brotman Baty Institute for Precision Medicine | The Brotman Baty Institute for Precision Medicine | Frazar,C.D., Lee,J., Ryke,E., Gamboa,L., McDermot,E., Stone,J., Kolar,T., Han,P.D., Sibley,T.R., Truong,M., Reinhart,D., Wolf,C.R., Duchin,J., Boeckh,M., Englund,J.A., Lutz,B.R., Waghmare,A., Viboud,C., Starita,L.M., Chu,H.Y., Bedford,T. and Shendure,J. |
| EPI_ISL_18143596, EPI_ISL_18143597, EPI_ISL_18143598, EPI_ISL_18143602 | The Brotman Baty Institute for Precision Medicine | The Brotman Baty Institute for Precision Medicine | Frazar,C.D., Lee,J., Gamboa,L., McDermot,E., Stone,J., Kolar,T., Han,P.D., Sibley,T.R., Truong,M., Reinhart,D., Wolf,C.R., Boeckh,M., Englund,J.A., Lutz,B.R., Waghmare,A., Viboud,C., Starita,L.M., Shendure,J., Bedford,T. and Chu,H.Y. |
| EPI_ISL_18143603, EPI_ISL_18143612, EPI_ISL_18143616 | The Brotman Baty Institute for Precision Medicine | The Brotman Baty Institute for Precision Medicine | Frazar,C.D., Lee,J., Ryke,E., Gamboa,L., McDermot,E., Stone,J., Kolar,T., Han,P.D., Sibley,T.R., Truong,M., Reinhart,D., Wolf,C.R., Duchin,J., Boeckh,M., Englund,J.A., Lutz,B.R., Waghmare,A., Viboud,C., Starita,L.M., Chu,H.Y., Bedford,T. and Shendure,J. |
| EPI_ISL_18143617 | The Brotman Baty Institute for Precision Medicine | The Brotman Baty Institute for Precision Medicine | Frazar,C.D., Lee,J., Gamboa,L., McDermot,E., Stone,J., Kolar,T., Han,P.D., Sibley,T.R., Truong,M., Reinhart,D., Wolf,C.R., Boeckh,M., Englund,J.A., Lutz,B.R., Waghmare,A., Viboud,C., Starita,L.M., Shendure,J., Bedford,T. and Chu,H.Y. |
| EPI_ISL_18143619 | The Brotman Baty Institute for Precision Medicine | The Brotman Baty Institute for Precision Medicine | Frazar,C.D., Lee,J., Ryke,E., Gamboa,L., McDermot,E., Stone,J., Kolar,T., Han,P.D., Sibley,T.R., Truong,M., Reinhart,D., Wolf,C.R., Duchin,J., Boeckh,M., Englund,J.A., Lutz,B.R., Waghmare,A., Viboud,C., Starita,L.M., Chu,H.Y., Bedford,T. and Shendure,J. |
| EPI_ISL_18143625 | The Brotman Baty Institute for Precision Medicine | The Brotman Baty Institute for Precision Medicine | Frazar,C.D., Lee,J., Gamboa,L., McDermot,E., Stone,J., Kolar,T., Han,P.D., Sibley,T.R., Truong,M., Reinhart,D., Wolf,C.R., Boeckh,M., Englund,J.A., Lutz,B.R., Waghmare,A., Viboud,C., Starita,L.M., Shendure,J., Bedford,T. and Chu,H.Y. |
| EPI_ISL_18143627, EPI_ISL_18143628 | The Brotman Baty Institute for Precision Medicine | The Brotman Baty Institute for Precision Medicine | Frazar,C.D., Lee,J., Ryke,E., Gamboa,L., McDermot,E., Stone,J., Kolar,T., Han,P.D., Sibley,T.R., Truong,M., Reinhart,D., Wolf,C.R., Duchin,J., Boeckh,M., Englund,J.A., Lutz,B.R., Waghmare,A., Viboud,C., Starita,L.M., Chu,H.Y., Bedford,T. and Shendure,J. |
| EPI_ISL_18143630 | The Brotman Baty Institute for Precision Medicine | The Brotman Baty Institute for Precision Medicine | Frazar,C.D., Lee,J., Gamboa,L., McDermot,E., Stone,J., Kolar,T., Han,P.D., Sibley,T.R., Truong,M., Reinhart,D., Wolf,C.R., Boeckh,M., Englund,J.A., Lutz,B.R., Waghmare,A., Viboud,C., Starita,L.M., Shendure,J., Bedford,T. and Chu,H.Y. |
| EPI_ISL_18143631, EPI_ISL_18143632, EPI_ISL_18143633, EPI_ISL_18143634, EPI_ISL_18143635, EPI_ISL_18143637, EPI_ISL_18143642 | The Brotman Baty Institute for Precision Medicine | The Brotman Baty Institute for Precision Medicine | Frazar,C.D., Lee,J., Ryke,E., Gamboa,L., McDermot,E., Stone,J., Kolar,T., Han,P.D., Sibley,T.R., Truong,M., Reinhart,D., Wolf,C.R., Duchin,J., Boeckh,M., Englund,J.A., Lutz,B.R., Waghmare,A., Viboud,C., Starita,L.M., Chu,H.Y., Bedford,T. and Shendure,J. |
| EPI_ISL_18143646 | The Brotman Baty Institute for Precision Medicine | The Brotman Baty Institute for Precision Medicine | Frazar,C.D., Lee,J., Gamboa,L., McDermot,E., Stone,J., Kolar,T., Han,P.D., Sibley,T.R., Truong,M., Reinhart,D., Wolf,C.R., Boeckh,M., Englund,J.A., Lutz,B.R., Waghmare,A., Viboud,C., Starita,L.M., Shendure,J., Bedford,T. and Chu,H.Y. |
| EPI_ISL_18143650, EPI_ISL_18143653, EPI_ISL_18143659, EPI_ISL_18143661 | The Brotman Baty Institute for Precision Medicine | The Brotman Baty Institute for Precision Medicine | Frazar,C.D., Lee,J., Ryke,E., Gamboa,L., McDermot,E., Stone,J., Kolar,T., Han,P.D., Sibley,T.R., Truong,M., Reinhart,D., Wolf,C.R., Duchin,J., Boeckh,M., Englund,J.A., Lutz,B.R., Waghmare,A., Viboud,C., Starita,L.M., Chu,H.Y., Bedford,T. and Shendure,J. |
| EPI_ISL_18143665, EPI_ISL_18143666, EPI_ISL_18143667 | The Brotman Baty Institute for Precision Medicine | The Brotman Baty Institute for Precision Medicine | Frazar,C.D., Lee,J., Gamboa,L., McDermot,E., Stone,J., Kolar,T., Han,P.D., Sibley,T.R., Truong,M., Reinhart,D., Wolf,C.R., Boeckh,M., Englund,J.A., Lutz,B.R., Waghmare,A., Viboud,C., Starita,L.M., Shendure,J., Bedford,T. and Chu,H.Y. |
| EPI_ISL_18143669, EPI_ISL_18143671, EPI_ISL_18143673, EPI_ISL_18143674 | The Brotman Baty Institute for Precision Medicine | The Brotman Baty Institute for Precision Medicine | Frazar,C.D., Lee,J., Ryke,E., Gamboa,L., McDermot,E., Stone,J., Kolar,T., Han,P.D., Sibley,T.R., Truong,M., Reinhart,D., Wolf,C.R., Duchin,J., Boeckh,M., Englund,J.A., Lutz,B.R., Waghmare,A., Viboud,C., Starita,L.M., Chu,H.Y., Bedford,T. and Shendure,J. |
| EPI_ISL_18143676 | The Brotman Baty Institute for Precision Medicine | The Brotman Baty Institute for Precision Medicine | Frazar,C.D., Lee,J., Gamboa,L., McDermot,E., Stone,J., Kolar,T., Han,P.D., Sibley,T.R., Truong,M., Reinhart,D., Wolf,C.R., Boeckh,M., Englund,J.A., Lutz,B.R., Waghmare,A., Viboud,C., Starita,L.M., Shendure,J., Bedford,T. and Chu,H.Y. |
| EPI_ISL_18143678, EPI_ISL_18143681, EPI_ISL_18143682, EPI_ISL_18143683 | The Brotman Baty Institute for Precision Medicine | The Brotman Baty Institute for Precision Medicine | Frazar,C.D., Lee,J., Ryke,E., Gamboa,L., McDermot,E., Stone,J., Kolar,T., Han,P.D., Sibley,T.R., Truong,M., Reinhart,D., Wolf,C.R., Duchin,J., Boeckh,M., Englund,J.A., Lutz,B.R., Waghmare,A., Viboud,C., Starita,L.M., Chu,H.Y., Bedford,T. and Shendure,J. |
| EPI_ISL_18143684, EPI_ISL_18143689, EPI_ISL_18143690 | The Brotman Baty Institute for Precision Medicine | The Brotman Baty Institute for Precision Medicine | Frazar,C.D., Lee,J., Gamboa,L., McDermot,E., Stone,J., Kolar,T., Han,P.D., Sibley,T.R., Truong,M., Reinhart,D., Wolf,C.R., Boeckh,M., Englund,J.A., Lutz,B.R., Waghmare,A., Viboud,C., Starita,L.M., Shendure,J., Bedford,T. and Chu,H.Y. |
| EPI_ISL_18143691 | The Brotman Baty Institute for Precision Medicine | The Brotman Baty Institute for Precision Medicine | Frazar,C.D., Lee,J., Ryke,E., Gamboa,L., McDermot,E., Stone,J., Kolar,T., Han,P.D., Sibley,T.R., Truong,M., Reinhart,D., Wolf,C.R., Duchin,J., Boeckh,M., Englund,J.A., Lutz,B.R., Waghmare,A., Viboud,C., Starita,L.M., Chu,H.Y., Bedford,T. |

| | | | |
|---|---|---|---|
| | | | and Shendure,J. |
| EPI_ISL_18143693, EPI_ISL_18143696, EPI_ISL_18143697 | The Brotman Baty Institute for Precision Medicine | The Brotman Baty Institute for Precision Medicine | Frazar,C.D., Lee,J., Gamboa,L., McDermot,E., Stone,J., Kolar,T., Han,P.D., Sibley,T.R., Truong,M., Reinhart,D., Wolf,C.R., Boeckh,M., Englund,J.A., Lutz,B.R., Waghmare,A., Viboud,C., Starita,L.M., Shendure,J., Bedford,T. and Chu,H.Y. |
| EPI_ISL_18143698, EPI_ISL_18143702, EPI_ISL_18143705, EPI_ISL_18143708, EPI_ISL_18143710, EPI_ISL_18143712 | The Brotman Baty Institute for Precision Medicine | The Brotman Baty Institute for Precision Medicine | Frazar,C.D., Lee,J., Ryke,E., Gamboa,L., McDermot,E., Stone,J., Kolar,T., Han,P.D., Sibley,T.R., Truong,M., Reinhart,D., Wolf,C.R., Duchin,J., Boeckh,M., Englund,J.A., Lutz,B.R., Waghmare,A., Viboud,C., Starita,L.M., Chu,H.Y., Bedford,T. and Shendure,J. |
| EPI_ISL_18143713, EPI_ISL_18143716 | The Brotman Baty Institute for Precision Medicine | The Brotman Baty Institute for Precision Medicine | Frazar,C.D., Lee,J., Gamboa,L., McDermot,E., Stone,J., Kolar,T., Han,P.D., Sibley,T.R., Truong,M., Reinhart,D., Wolf,C.R., Boeckh,M., Englund,J.A., Lutz,B.R., Waghmare,A., Viboud,C., Starita,L.M., Shendure,J., Bedford,T. and Chu,H.Y. |
| EPI_ISL_18204749, EPI_ISL_18204752, EPI_ISL_18204769, EPI_ISL_18204790, EPI_ISL_18204798, EPI_ISL_18204800, EPI_ISL_18204803, EPI_ISL_18204805, EPI_ISL_18204807, EPI_ISL_18204809, EPI_ISL_18204811, EPI_ISL_18204813, EPI_ISL_18204815, EPI_ISL_18204816, EPI_ISL_18204817, EPI_ISL_18204819, EPI_ISL_18204821, EPI_ISL_18204823, EPI_ISL_18204825, EPI_ISL_18204829, EPI_ISL_18204831, EPI_ISL_18204833, EPI_ISL_18204836, EPI_ISL_18204854, EPI_ISL_18204863, EPI_ISL_18204876, EPI_ISL_18204878, EPI_ISL_18204880, EPI_ISL_18204884 | | | |
| see above | Microbiology Department. Complexo Hospitalario Universitario de Vigo | Microbiology Department. Complexo Hospitalario Universitario de Vigo | Daviña C, Martinez L, Perez-Castro S |
| EPI_ISL_18228533, EPI_ISL_18228536, EPI_ISL_18228537, EPI_ISL_18228539, EPI_ISL_18228540, EPI_ISL_18228541, EPI_ISL_18228544, EPI_ISL_18228548, EPI_ISL_18228618, EPI_ISL_18228653, EPI_ISL_18228657, EPI_ISL_18228658, EPI_ISL_18228659, EPI_ISL_18228660, EPI_ISL_18228661, EPI_ISL_18228662, EPI_ISL_18228663, EPI_ISL_18228703, EPI_ISL_18228704, EPI_ISL_18228705, EPI_ISL_18228707, EPI_ISL_18228708, EPI_ISL_18228709, EPI_ISL_18228811, EPI_ISL_18229075, EPI_ISL_18229381, EPI_ISL_18229635, EPI_ISL_18229703 | | | |
| see above | Institut Pasteur de Dakar | Institut Pasteur de Dakar | Jallow, Mamadou Malado; Diagne, Moussa Moise; Sankhe, Safietou; Mendy, Marie Pedapa; Ndiaye, Ndiende Koba; Sy, Sara; Kiori, Davy; Goudiaby, Deborah; Dia, Ndongo |
| EPI_ISL_18231036, EPI_ISL_18231037, EPI_ISL_18231038, EPI_ISL_18231039, EPI_ISL_18231040, EPI_ISL_18231041, EPI_ISL_18231042, EPI_ISL_18231043, EPI_ISL_18231044, EPI_ISL_18231045, EPI_ISL_18231046, EPI_ISL_18231047, EPI_ISL_18231048, EPI_ISL_18231124, EPI_ISL_18231125, EPI_ISL_18231126, EPI_ISL_18231128, EPI_ISL_18231129, EPI_ISL_18231130, EPI_ISL_18231131 | | | |
| see above | Virology Laboratory, International Centre for Diarrhoeal Disease Research, Bangladesh (ICDDR,B) | Genome Centre, International Centre for Diarrhoeal Disease Research, Bangladesh (ICDDR,B) | Mohammad Jubair, Tasnim Jabin, Md. Mobarok Hossain, Shahriar Islam, Shovan Basak Moon, Mustafizur Rahman |
| EPI_ISL_18262241 | Instituto de Medicina Tropical Alexander von Humboldt, Universidad Peruana Cayetano Heredia | Laboratorio de Genómica Microbiana, Universidad Peruana Cayetano Heredia | Ericka Meza, Diego Cuicapuza, Anne Martínez-Ventura, Brenda Ayzanoa, Janet Huancachoque, César Ugarte, Carlos Zamudio, Pablo Tsukayama |
| EPI_ISL_18272383, EPI_ISL_18272384, EPI_ISL_18272385, EPI_ISL_18272395, EPI_ISL_18272396, EPI_ISL_18272397, EPI_ISL_18272406, EPI_ISL_18272407, EPI_ISL_18272408, EPI_ISL_18272409, EPI_ISL_18272410, EPI_ISL_18272411, EPI_ISL_18272412, EPI_ISL_18272413, EPI_ISL_18272414, EPI_ISL_18272415, EPI_ISL_18274506, EPI_ISL_18274507, EPI_ISL_18274508, EPI_ISL_18274509, EPI_ISL_18274510, EPI_ISL_18274511, EPI_ISL_18274512, EPI_ISL_18274513, EPI_ISL_18274514, EPI_ISL_18274515 | | | |
| see above | West of Scotland Specialist Virology Centre | West of Scotland Specialist Virology Centre | Lynne Ferguson, Imogen Johnston-Menzies, Rory Gunson |
| EPI_ISL_18277073 | Microbiology Department. Complexo Hospitalario Universitario de Vigo | Microbiology Department. Complexo Hospitalario Universitario de Vigo | Daviña C, Martinez L, Perez-Castro S |
| EPI_ISL_18277120, EPI_ISL_18277121, EPI_ISL_18277122, EPI_ISL_18277123, EPI_ISL_18277124, EPI_ISL_18277125, EPI_ISL_18277126, EPI_ISL_18277127, EPI_ISL_18277128, EPI_ISL_18277129, EPI_ISL_18277130, EPI_ISL_18277131 | | | |
| see above | West of Scotland Specialist Virology Centre | West of Scotland Specialist Virology Centre | Lynne Ferguson, Imogen Johnston-Menzies, Rory Gunson |
| EPI_ISL_18279071 | National Institute of Public Health | National Institute of Public Health | Chhorvann CHHEA, Darapheak CHAU, Sitha PRUM, Visal CHHE, Sokha DUL, Sengly MANH, Phally VY, Borann SAR, Chanthap LON, Jessica E. Manning, Sophana Chea, Savuth CHIN |
| EPI_ISL_18321021, EPI_ISL_18321022, EPI_ISL_18321023, EPI_ISL_18321024, EPI_ISL_18321025, EPI_ISL_18321026, EPI_ISL_18321027, EPI_ISL_18321028, EPI_ISL_18321029, EPI_ISL_18321030, EPI_ISL_18321031, EPI_ISL_18321032, EPI_ISL_18321033, EPI_ISL_18321056, EPI_ISL_18321057, EPI_ISL_18321058, EPI_ISL_18321059, EPI_ISL_18321060, EPI_ISL_18321061, EPI_ISL_18321062, EPI_ISL_18321063, EPI_ISL_18321064, EPI_ISL_18321065, EPI_ISL_18321066, EPI_ISL_18321067, EPI_ISL_18321068, EPI_ISL_18321069, EPI_ISL_18321070, EPI_ISL_18321071, EPI_ISL_18321072, EPI_ISL_18321073, EPI_ISL_18321074, EPI_ISL_18321075, EPI_ISL_18321076, EPI_ISL_18321077, EPI_ISL_18321102, EPI_ISL_18321103, EPI_ISL_18321104, EPI_ISL_18321105, EPI_ISL_18321106, EPI_ISL_18321107, EPI_ISL_18321108, EPI_ISL_18321109, EPI_ISL_18321110, EPI_ISL_18321111, EPI_ISL_18321112, EPI_ISL_18321113, EPI_ISL_18321114, EPI_ISL_18321115, EPI_ISL_18321116, EPI_ISL_18321117, EPI_ISL_18321118, EPI_ISL_18321119, EPI_ISL_18321120, EPI_ISL_18321121, EPI_ISL_18321122, EPI_ISL_18321123, EPI_ISL_18321124, EPI_ISL_18321125, EPI_ISL_18321126, EPI_ISL_18321127, EPI_ISL_18321128, EPI_ISL_18321129, EPI_ISL_18321130, EPI_ISL_18321132, EPI_ISL_18321133, EPI_ISL_18321134, EPI_ISL_18321135, EPI_ISL_18321136, EPI_ISL_18321137, EPI_ISL_18321138, EPI_ISL_18321139, EPI_ISL_18321140, EPI_ISL_18321141, EPI_ISL_18321142, EPI_ISL_18321143, EPI_ISL_18321144, EPI_ISL_18321145, EPI_ISL_18321146, EPI_ISL_18321148, EPI_ISL_18321149, EPI_ISL_18321150, EPI_ISL_18321151, EPI_ISL_18321152, EPI_ISL_18321153, EPI_ISL_18321154, EPI_ISL_18321155, EPI_ISL_18321156, EPI_ISL_18321157, EPI_ISL_18321158 | | | |
| see above | National Virus Reference Laboratory | National Virus Reference Laboratory | Michael Carr, Charlene Bennett, Jonathan Dean, Daniel Hare, Cillian F De Gascun |
| EPI_ISL_18323795, EPI_ISL_18323797, EPI_ISL_18323798 | Servicio de Microbiología Complejo Hospitalario Universitario Nuestra Señora de Candelaria | Institute of Technology and Renewable Energy (ITER) | Julia,Alcoba-Florez;Rafaela,González-Montelongo;Diego,García-Martínez de Artola;Adrián,Muñoz-Barrera;Helena,Gil-Campesino;Oscar,Diez-Gil;Jose Miguel,Lorenzo-Salazar;Carlos,Flores |
| EPI_ISL_18329464, EPI_ISL_18329467, EPI_ISL_18329468, EPI_ISL_18329469, EPI_ISL_18329470, EPI_ISL_18329471, EPI_ISL_18329472, EPI_ISL_18329473, EPI_ISL_18329474, EPI_ISL_18329475, EPI_ISL_18329476, EPI_ISL_18329477, EPI_ISL_18329478, EPI_ISL_18329479, EPI_ISL_18329480, EPI_ISL_18329481, EPI_ISL_18329482, EPI_ISL_18329483, EPI_ISL_18329484, EPI_ISL_18329485, EPI_ISL_18329486, EPI_ISL_18329487, EPI_ISL_18329488, EPI_ISL_18329489, EPI_ISL_18329490, EPI_ISL_18329491 | | | |
| see above | Microbiology and Virology Depertment, Fondazione IRCCS Policlinico San Matteo, Pavia | Microbiology and Virology Depertment, Fondazione IRCCS Policlinico San Matteo, Pavia | Gugliemo Ferrari, Federica Giardina, Antonino Pitrolo, Fausto Baldanti |
| EPI_ISL_18334164, EPI_ISL_18334165, EPI_ISL_18334168, EPI_ISL_18334170, EPI_ISL_18334172, EPI_ISL_18334174, EPI_ISL_18334181, EPI_ISL_18334182, EPI_ISL_18334183, EPI_ISL_18334184, EPI_ISL_18334186, EPI_ISL_18334188, EPI_ISL_18334189, EPI_ISL_18334192, EPI_ISL_18334193, EPI_ISL_18334197, EPI_ISL_18334198, EPI_ISL_18334200, EPI_ISL_18334202, EPI_ISL_18334203, EPI_ISL_18334207, EPI_ISL_18334208, EPI_ISL_18334209, EPI_ISL_18334210, EPI_ISL_18334211, EPI_ISL_18334212, EPI_ISL_18334213, EPI_ISL_18334217, EPI_ISL_18334218, EPI_ISL_18334219, EPI_ISL_18334223, EPI_ISL_18334224, EPI_ISL_18334226, EPI_ISL_18334227, EPI_ISL_18334228, EPI_ISL_18334229, EPI_ISL_18334233, EPI_ISL_18334234, EPI_ISL_18334235, EPI_ISL_18334237 | | | |
| see above | Baylor College of Medicine, Division of Pediatric Tropical Medicine | Baylor College of Medicine, Division of Pediatric Tropical Medicine | Avadhanula,V., Agustinho,D.P., Menon,V.K., Chemaly,R.F., Shah,D.P., Qin,X., Surathu,A., Doddapaneni,H., Muzny,D.M., Metcalf,G.A., Gregeen,S.J., Gibbs,R.A., Petrosino,J.J., Sedlazeck,F.J. and Piedra,P.A. |
| EPI_ISL_1834174, EPI_ISL_1834175, EPI_ISL_1834176, EPI_ISL_1834177, EPI_ISL_1834178, EPI_ISL_1834179, EPI_ISL_1834180, EPI_ISL_1834181, EPI_ISL_1834182, EPI_ISL_1834183, EPI_ISL_1834184 | | | |
| see above | Respiratory Virus Unit, National Infection Service, Public Health England | National Infection Service, Public Health England | Zambon M, Talts T, Ellis J, Miah S, Platt S |
| EPI_ISL_18348028, EPI_ISL_18348029, EPI_ISL_18348030, EPI_ISL_18348031 | Republic of Macedonia, Institute of Public Health Laboratory of Virology and Molecular Diagnostics | Republic of Macedonia, Institute of Public Health Laboratory of Virology and Molecular Diagnostics | Teodora Buzharova, Golubinka Boshevska, Elizabeta Jancheska |
| EPI_ISL_18351213, EPI_ISL_18351214 | Edendale Hospital | National Institute for Communicable Diseases of the National Health Laboratory Service | Everatt J, Kekana D, Mahlangu B, Stock N, Ntozini B, Ntuli N, Mnguni A, Nzimande A, Ismail A, Bhiman JN, Wolter N |
| EPI_ISL_18351215 | Rahima Moosa Mother and Child Hospital | National Institute for Communicable Diseases of the National Health Laboratory Service | Everatt J, Kekana D, Mahlangu B, Stock N, Ntozini B, Ntuli N, Mnguni A, Nzimande A, Ismail A, Bhiman JN, Wolter N |
| EPI_ISL_18351216 | Edendale Hospital | National Institute for Communicable Diseases of the National Health Laboratory Service | Everatt J, Kekana D, Mahlangu B, Stock N, Ntozini B, Ntuli N, Mnguni A, Nzimande A, Ismail A, Bhiman JN, Wolter N |
| EPI_ISL_18351219 | National Institute for Communicable Diseases of the National Health Laboratory Service | National Institute for Communicable Diseases of the National Health Laboratory Service | Everatt J, Kekana D, Mahlangu B, Stock N, Ntozini B, Ntuli N, Mnguni A, Nzimande A, Ismail A, Bhiman JN, Wolter N |
| EPI_ISL_18371219, EPI_ISL_18371220, EPI_ISL_18371221, EPI_ISL_18371222, EPI_ISL_18371223, EPI_ISL_18371224, EPI_ISL_18371225, EPI_ISL_18371226, EPI_ISL_18371227, EPI_ISL_18371228 | Virology Laboratory, International Centre for Diarrhoeal Disease Research, Bangladesh (ICDDR,B) | Genome Centre, International Centre for Diarrhoeal Disease Research, Bangladesh (ICDDR,B) | Mohammad Jubair, Tasnim Jabin, Md. Mobarok Hossain, Shahriar Islam, Shovan Basak Moon, Mustafizur Rahman |
| EPI_ISL_18387877, EPI_ISL_18387884, EPI_ISL_18387889, EPI_ISL_18387898, EPI_ISL_18387902, EPI_ISL_18387907, EPI_ISL_18387910, EPI_ISL_18387935, EPI_ISL_18387936, EPI_ISL_18387940, EPI_ISL_18387942, EPI_ISL_18387945, EPI_ISL_18387947, EPI_ISL_18387954 | | | |
| see above | The Brotman Baty Institute for Precision Medicine | The Brotman Baty Institute for Precision Medicine | Frazar,C.D., Lee,J., Ryke,E., Gamboa,L., McDermot,E., Kolar,T., Han,P.D., Sibley,T.R., Reinhart,D., Grindstaff,S., Truong,M., Babu,T.M., Feldstein,L.R., Saydah,S., Briggs-Hagen,M., Casto,A., Ehmen,B., Englund,J.A., Fortmann,S.P., Kuntz,J.L., Lockwood,T., Midgley,C.M., Mularski,R.A., Ogilvie,T., Reich,S., Schmidt,M.A., Smith,N., Starita,L., Stone,J., Vandermeer,M., Weil,A.A., Wolf,C.R., Chu,H.Y. and Naleway,A.L. |
| EPI_ISL_18446811, EPI_ISL_18446812, EPI_ISL_18446814, EPI_ISL_18446815, EPI_ISL_18446816, EPI_ISL_18446817, EPI_ISL_18446818, EPI_ISL_18446819, EPI_ISL_18446820, EPI_ISL_18446821, EPI_ISL_18446822, EPI_ISL_18446823, EPI_ISL_18446824, EPI_ISL_18446825, EPI_ISL_18446826, EPI_ISL_18446827, EPI_ISL_18446828, EPI_ISL_18446829, EPI_ISL_18446830, EPI_ISL_18446832, EPI_ISL_18446833, EPI_ISL_18446834, EPI_ISL_18446835, EPI_ISL_18446837, EPI_ISL_18446838, EPI_ISL_18446839, EPI_ISL_18446840, EPI_ISL_18446842, EPI_ISL_18446843, EPI_ISL_18446844, EPI_ISL_18446845, EPI_ISL_18446846, EPI_ISL_18446847, EPI_ISL_18446848 | | | |
| see above | Vittore Buzzi Children Hospital, Pediatric Department | Laboratory of Infectious Diseases, Department of Biomedical and Clinical Sciences, University of Milan | Alessia Lai, Annalisa Bergna, Valentina Fabiano, Carla della Ventura, Giulia Fumagalli, Alessandra Mari, Martina Loiodice, Gian Vincenzo Zuccotti, Gianguglielmo Zehender |
| EPI_ISL_18447451, EPI_ISL_18447452, EPI_ISL_18447453, EPI_ISL_18447454, EPI_ISL_18447455, EPI_ISL_18447456, EPI_ISL_18447457, EPI_ISL_18447458, EPI_ISL_18447459, EPI_ISL_18447460, EPI_ISL_18447461, EPI_ISL_18447462, EPI_ISL_18447463, EPI_ISL_18447464, EPI_ISL_18447465, EPI_ISL_18447466, EPI_ISL_18447467, EPI_ISL_18447468, EPI_ISL_18447469, EPI_ISL_18447470, EPI_ISL_18447471, EPI_ISL_18447472, EPI_ISL_18447473, EPI_ISL_18447474, EPI_ISL_18447475, EPI_ISL_18447476, EPI_ISL_18447478, EPI_ISL_18447479, EPI_ISL_18447480, EPI_ISL_18447481, EPI_ISL_18447482, EPI_ISL_18447483, EPI_ISL_18447484, EPI_ISL_18447485, EPI_ISL_18447486, EPI_ISL_18447487, EPI_ISL_18447488, EPI_ISL_18447489, EPI_ISL_18447490, EPI_ISL_18447491, EPI_ISL_18447492, EPI_ISL_18447493, EPI_ISL_18447494, EPI_ISL_18447495, EPI_ISL_18447498 | | | |
| see above | National Centre of Infectious and Parasitic Diseases National Laboratory "Influenza and ARD" | National Centre of Infectious and Parasitic Diseases National Laboratory "Influenza and ARD" | Ivelina Trifonova; Neli Korsun; Iveta Madzharova |
| EPI_ISL_18482716, EPI_ISL_18482721, EPI_ISL_18482722, EPI_ISL_18482725, EPI_ISL_18482726, EPI_ISL_18482727, EPI_ISL_18482732, EPI_ISL_18482733, EPI_ISL_18482737, EPI_ISL_18482738, EPI_ISL_18482739, EPI_ISL_18482742, EPI_ISL_18482743, EPI_ISL_18482745, EPI_ISL_18482747, EPI_ISL_18482750, EPI_ISL_18482754, EPI_ISL_18482755, EPI_ISL_18482758, EPI_ISL_18482763, EPI_ISL_18482764, EPI_ISL_18482768, EPI_ISL_18482770, EPI_ISL_18482774, EPI_ISL_18482783, EPI_ISL_18482789, EPI_ISL_18482790, EPI_ISL_18482794, EPI_ISL_18482796, EPI_ISL_18482798, EPI_ISL_18482803, EPI_ISL_18482806, EPI_ISL_18482810, EPI_ISL_18482812, EPI_ISL_18482818 | | | |
| see above | Beijing Children's Hospital, Capital Medical University, Laboratory of Infection and Virology | Beijing Children's Hospital, Capital Medical University, Laboratory of Infection and Virology | Li,F., Zhu,Y. and Chen,X. |
| EPI_ISL_18507279, EPI_ISL_18507280, EPI_ISL_18507284, EPI_ISL_18507290, EPI_ISL_18507299, EPI_ISL_18507300 | Victorian Infectious Diseases Reference Laboratory, Translational Diagnostics | Victorian Infectious Diseases Reference Laboratory, Translational Diagnostics | Steinig JOACHIM,E. |
| EPI_ISL_18509909, EPI_ISL_18509929, EPI_ISL_18509930, EPI_ISL_18509931, EPI_ISL_18509932, EPI_ISL_18509933, EPI_ISL_18510013, EPI_ISL_18510135, EPI_ISL_18510136 | Institut Pasteur de Dakar | Institut Pasteur de Dakar | Jallow,Mamadou Malado; Diagne,Moussa Moise; Sankhe,Safietou; Diop,Seynabou Mbaye Ba Souna; Mendy,Marie Pedapa; Sy,Sara; Ndiaye,Ndiende Koba; Kiory, Davy Evrard; Goudiaby,Deborah; Dia,Ndongo |
| EPI_ISL_18510438 | Microbiology Service, A Coruña University Hospital Complex (CHUAC) | Microbiology Service, A Coruña University Hospital Complex (CHUAC) | Ana Fernandez Gonzalez, Pablo Aja-Macaya, Maria Jose Muiño Andrade, Iria Sendon Sanvicente, Soraya Rumbo-Feal, Juan A. Vallejo, Jorge Arca-Suarez, German Bou |
| EPI_ISL_18522760, EPI_ISL_18522761, EPI_ISL_18522762 | Servicio Microbiología Hospital La Paz | Servicio Microbiología Hospital La Paz | Fernando Lázaro, Iván Bloise, Pablo Prieto Casado, Francisco López Rodrigo, Jesús Mingorance Cruz, Elie Dahdouh |
| EPI_ISL_18537466, EPI_ISL_18537467, EPI_ISL_18537471, EPI_ISL_18537473 | RELAB - BIOGROUP - Plateau technique St Denis | National Reference Center for Viruses of Respiratory Infections, Institut Pasteur, Paris | Léa Avon, Marion Barbet, Marine Bernard, Emma Bezot, Angela Brisebarre, Océane Dehan, Flora Donati, Vanessa Guimaraes, Banujaa Jeyarajah, Florian Préjean, Yannis Rahou, Sylvie van der Werf, Frédéric Lemoine, Samar Berreira Ibraim, Jérôme Bourret, Kévin Da Silva, Maud Vanpeene, Vincent Enouf, Marie-Anne Rameix-Welti, Yanis CHAIB |
| EPI_ISL_18537475 | RELAB - Laboratoire Biomer | National Reference Center for Viruses of Respiratory Infections, Institut Pasteur, Paris | Léa Avon, Marion Barbet, Marine Bernard, Emma Bezot, Angela Brisebarre, Océane Dehan, Flora Donati, Vanessa Guimaraes, Banujaa Jeyarajah, Florian Préjean, Yannis Rahou, Sylvie van der Werf, Frédéric Lemoine, Samar Berreira Ibraim, Jérôme Bourret, Kévin Da Silva, Maud Vanpeene, Vincent Enouf, Marie-Anne Rameix-Welti, Alexandra JACQUES |
| EPI_ISL_18537476, EPI_ISL_18537477, EPI_ISL_18537481, | RELAB - BIOGROUP - Plateau technique St Denis | National Reference Center for Viruses of | Léa Avon, Marion Barbet, Marine Bernard, Emma Bezot, Angela Brisebarre, Océane Dehan, Flora Donati, Vanessa Guimaraes, Banujaa Jeyarajah, Florian Préjean, Yannis Rahou, Sylvie van der Werf, Frédéric Lemoine, Samar Berreira |

| | | | |
|---|---|---|---|
| EPI_ISL_18537485 | | Respiratory Infections, Institut Pasteur, Paris | Ibraim, Jérôme Bourret, Kévin Da Silva, Maud Vanpeene, Vincent Enouf, Marie-Anne Rameix-Welti, Yanis CHAIB |
| EPI_ISL_18537486 | Sentinelles Ile-de-France | National Reference Center for Viruses of Respiratory Infections, Institut Pasteur, Paris | Léa Avon, Marion Barbet, Marine Bernard, Emma Bezot, Angela Brisebarre, Océane Dehan, Flora Donati, Vanessa Guimaraes, Banujaa Jeyarajah, Florian Préjean, Yannis Rahou, Sylvie van der Werf, Frédéric Lemoine, Samar Berreira Ibraim, Jérôme Bourret, Kévin Da Silva, Maud Vanpeene, Vincent Enouf, Marie-Anne Rameix-Welti, Patricia LUBELSKI |
| EPI_ISL_18537490, EPI_ISL_18537491, EPI_ISL_18537492, EPI_ISL_18537493 | RELAB - BIOGROUP - Plateau technique St Denis | National Reference Center for Viruses of Respiratory Infections, Institut Pasteur, Paris | Léa Avon, Marion Barbet, Marine Bernard, Emma Bezot, Angela Brisebarre, Océane Dehan, Flora Donati, Vanessa Guimaraes, Banujaa Jeyarajah, Florian Préjean, Yannis Rahou, Sylvie van der Werf, Frédéric Lemoine, Samar Berreira Ibraim, Jérôme Bourret, Kévin Da Silva, Maud Vanpeene, Vincent Enouf, Marie-Anne Rameix-Welti, Yanis CHAIB |
| EPI_ISL_18537498 | RELAB - Laboratoire Biomer | National Reference Center for Viruses of Respiratory Infections, Institut Pasteur, Paris | Léa Avon, Marion Barbet, Marine Bernard, Emma Bezot, Angela Brisebarre, Océane Dehan, Flora Donati, Vanessa Guimaraes, Banujaa Jeyarajah, Florian Préjean, Yannis Rahou, Sylvie van der Werf, Frédéric Lemoine, Samar Berreira Ibraim, Jérôme Bourret, Kévin Da Silva, Maud Vanpeene, Vincent Enouf, Marie-Anne Rameix-Welti, Alexandra JACQUES |
| EPI_ISL_18537500, EPI_ISL_18537501, EPI_ISL_18537502, EPI_ISL_18537509, EPI_ISL_18537513 | RELAB - BIOGROUP - Plateau technique St Denis | National Reference Center for Viruses of Respiratory Infections, Institut Pasteur, Paris | Léa Avon, Marion Barbet, Marine Bernard, Emma Bezot, Angela Brisebarre, Océane Dehan, Flora Donati, Vanessa Guimaraes, Banujaa Jeyarajah, Florian Préjean, Yannis Rahou, Sylvie van der Werf, Frédéric Lemoine, Samar Berreira Ibraim, Jérôme Bourret, Kévin Da Silva, Maud Vanpeene, Vincent Enouf, Marie-Anne Rameix-Welti, Yanis CHAIB |
| EPI_ISL_18559741, EPI_ISL_18559742, EPI_ISL_18559743, EPI_ISL_18559744, EPI_ISL_18559745 | Research Institute of Experimental and Clinical Medicine | WHO National Influenza Centre Russian Federation | Komissarova K.S., Yolshin N.D., Kurskaya O.G., Solomatina M.V., Saroyan T.A., Dubovitskiy N.A., Derko A.A., Sobolev I.A., Sharshov K.A., Danilenko D.M., Lioznov D.A. |
| EPI_ISL_18568046 | RELAB - Biogroup - Laborizon Bretagne | National Reference Center for Viruses of Respiratory Infections, Institut Pasteur, Paris | Léa Avon, Marion Barbet, Marine Bernard, Emma Bezot, Angela Brisebarre, Océane Dehan, Flora Donati, Vanessa Guimaraes, Banujaa Jeyarajah, Florian Préjean, Yannis Rahou, Sylvie van der Werf, Frédéric Lemoine, Samar Berreira Ibraim, Jérôme Bourret, Kévin Da Silva, Maud Vanpeene, Vincent Enouf, Marie-Anne Rameix-Welti, Jean-François COMES |
| EPI_ISL_18568047 | Sentinelles Ile-de-France | National Reference Center for Viruses of Respiratory Infections, Institut Pasteur, Paris | Léa Avon, Marion Barbet, Marine Bernard, Emma Bezot, Angela Brisebarre, Océane Dehan, Flora Donati, Vanessa Guimaraes, Banujaa Jeyarajah, Florian Préjean, Yannis Rahou, Sylvie van der Werf, Frédéric Lemoine, Samar Berreira Ibraim, Jérôme Bourret, Kévin Da Silva, Maud Vanpeene, Vincent Enouf, Marie-Anne Rameix-Welti, Marie Annick BURGESS |
| EPI_ISL_18568048 | Centre Hospitalier (CH) d'Argenteuil (Victor Dupouy) | National Reference Center for Viruses of Respiratory Infections, Institut Pasteur, Paris | Léa Avon, Marion Barbet, Marine Bernard, Emma Bezot, Angela Brisebarre, Océane Dehan, Flora Donati, Vanessa Guimaraes, Banujaa Jeyarajah, Florian Préjean, Yannis Rahou, Sylvie van der Werf, Frédéric Lemoine, Samar Berreira Ibraim, Jérôme Bourret, Kévin Da Silva, Maud Vanpeene, Vincent Enouf, Marie-Anne Rameix-Welti, L COURDAVAULT |
| EPI_ISL_18568049 | Hôpital Ambroise Paré - Service de Microbiologie et hygiène | National Reference Center for Viruses of Respiratory Infections, Institut Pasteur, Paris | Léa Avon, Marion Barbet, Marine Bernard, Emma Bezot, Angela Brisebarre, Océane Dehan, Flora Donati, Vanessa Guimaraes, Banujaa Jeyarajah, Florian Préjean, Yannis Rahou, Sylvie van der Werf, Frédéric Lemoine, Samar Berreira Ibraim, Jérôme Bourret, Kévin Da Silva, Maud Vanpeene, Vincent Enouf, Marie-Anne Rameix-Welti, Elyanne GAULT |
| EPI_ISL_18568050 | Sentinelles Ile-de-France | National Reference Center for Viruses of Respiratory Infections, Institut Pasteur, Paris | Léa Avon, Marion Barbet, Marine Bernard, Emma Bezot, Angela Brisebarre, Océane Dehan, Flora Donati, Vanessa Guimaraes, Banujaa Jeyarajah, Florian Préjean, Yannis Rahou, Sylvie van der Werf, Frédéric Lemoine, Samar Berreira Ibraim, Jérôme Bourret, Kévin Da Silva, Maud Vanpeene, Vincent Enouf, Marie-Anne Rameix-Welti, Hajer KANOUN-DRIRA |
| EPI_ISL_18568051 | Sentinelles Province | National Reference Center for Viruses of Respiratory Infections, Institut Pasteur, Paris | Léa Avon, Marion Barbet, Marine Bernard, Emma Bezot, Angela Brisebarre, Océane Dehan, Flora Donati, Vanessa Guimaraes, Banujaa Jeyarajah, Florian Préjean, Yannis Rahou, Sylvie van der Werf, Frédéric Lemoine, Samar Berreira Ibraim, Jérôme Bourret, Kévin Da Silva, Maud Vanpeene, Vincent Enouf, Marie-Anne Rameix-Welti, Frédéric LE MEUR |
| EPI_ISL_18568345, EPI_ISL_18568346, EPI_ISL_18569161, EPI_ISL_18569165, EPI_ISL_18569169, EPI_ISL_18569170, EPI_ISL_18569171, EPI_ISL_18569172, EPI_ISL_18591786, EPI_ISL_18591787, EPI_ISL_18591788, EPI_ISL_18591790, EPI_ISL_18591794, EPI_ISL_18591800, EPI_ISL_18591801, EPI_ISL_18591802, EPI_ISL_18591805, EPI_ISL_18591806, EPI_ISL_18591808, EPI_ISL_18591809, EPI_ISL_18591812, EPI_ISL_18591815 | | | |
| see above | Respiratory Virus Unit / Reference Microbiology Services / UK Health Security Agency | Reference Microbiology Services / UK Health Security Agency | Zambon M. Talts T. Kele B. Miah S |
| EPI_ISL_18592443, EPI_ISL_18592444, EPI_ISL_18592445, EPI_ISL_18592446, EPI_ISL_18592447, EPI_ISL_18592448, EPI_ISL_18592449, EPI_ISL_18592450, EPI_ISL_18592451, EPI_ISL_18592452, EPI_ISL_18592453, EPI_ISL_18592454, EPI_ISL_18592455, EPI_ISL_18592456, EPI_ISL_18592457, EPI_ISL_18592458 | | | |
| see above | HOSPITAL UNIVERSITARIO CENTRAL DE ASTURIAS | Laboratorio de Virología HUCA | Pérez-Martínez Z, Boga JA, Rojo S, González-Alba JM, Ochoa-Varela C, Rodríguez-Pérez M, Melón S, Alvarez-Argüelles ME |
| EPI_ISL_2156815, EPI_ISL_2156816, EPI_ISL_2156817, EPI_ISL_2156818, EPI_ISL_2156819, EPI_ISL_2156820 | Royal Children's Hospital | WHO Influenza Centre for Reference and Research on Influenza | Angela Todd, Yi-Mo Deng, Annette Alafaci, Naomi Komadina |
| EPI_ISL_2543763 | Royal Children's Hospital | WHO Collaborating Centre for Reference and Research on Influenza | Xiaomin Dong, Annette Alafaci,Yi-Mo Deng, Ammar Aziz, Naomi Komadina |
| EPI_ISL_2543848 | Monash Medical Centre | WHO Collaborating Centre for Reference and Research on Influenza | Xiaomin Dong, Michelle Francis, Tony Korman, Yi-Mo Deng, Ammar Aziz, Naomi Komadina |
| EPI_ISL_2543849, EPI_ISL_2543850, EPI_ISL_2543851, EPI_ISL_2543852, EPI_ISL_2543853 | Royal Children's Hospital | WHO Collaborating Centre for Reference and Research on Influenza | Xiaomin Dong, Annette Alafaci,Yi-Mo Deng, Ammar Aziz, Naomi Komadina |
| EPI_ISL_2543854, EPI_ISL_2543855, EPI_ISL_2543856, EPI_ISL_2543857, EPI_ISL_2543858, EPI_ISL_2543859 | Institut Pasteur de Cote d'Ivoire | WHO Collaborating Centre for Reference and Research on Influenza | Xiaomin Dong, Herve Kadjo, Yi-Mo Deng, Ammar Aziz, Naomi Komadina |
| EPI_ISL_2543860, EPI_ISL_2543861, EPI_ISL_2543862, EPI_ISL_2543863, EPI_ISL_2543864, EPI_ISL_2543865, EPI_ISL_2543866, EPI_ISL_2543867, EPI_ISL_2543868, EPI_ISL_2543869, EPI_ISL_2543870, EPI_ISL_2543871, EPI_ISL_2543872, EPI_ISL_2543873, EPI_ISL_2543874, EPI_ISL_2543875, EPI_ISL_2543876, EPI_ISL_2543877, EPI_ISL_2543878, EPI_ISL_2543879, EPI_ISL_2543880, EPI_ISL_2543881, EPI_ISL_2543882, EPI_ISL_2543883, EPI_ISL_2543884, EPI_ISL_2543885, EPI_ISL_2543886, EPI_ISL_2543887, EPI_ISL_2543888, EPI_ISL_2543889 | | | |
| see above | National Center for Communicable Diseases | WHO Collaborating Centre for Reference and Research on Influenza | Xiaomin Dong, Darmaa Bardach, Yi-Mo Deng, Ammar Aziz, Naomi Komadina |
| EPI_ISL_2543890 | NIC, National Institute of Health | WHO Collaborating Centre for Reference and Research on Influenza | Xiaomin Dong, Pilailuk Akkapaiboon Okada, Yi-Mo Deng, Ammar Aziz, Naomi Komadina |
| EPI_ISL_2544066, EPI_ISL_2544067 | KEMRI Wellcome Trust Research Programme | KEMRI Wellcome Trust Research Programme | Agoti,C.N., Otieno,J.R., Munywoki,P.K., Mwihuri,A.G., Cane,P.A., Nokes,D.J., Kellam,P. and Cotten,M.L. |
| EPI_ISL_2544090, EPI_ISL_2544091, EPI_ISL_2544092, EPI_ISL_2544093, EPI_ISL_2544094 | J. Craig Venter Institute | J. Craig Venter Institute | Shabman,R., Das,S.R., Puri,V., Fedorova,N., Amedeo,P., Williams,M., Shrivastava,S. and Halasa,N. |
| EPI_ISL_2544100, EPI_ISL_2544101, EPI_ISL_2544102, EPI_ISL_2544103, EPI_ISL_2544104 | Emerging Viral Infections, Oxford University Clinical Research Unit | Emerging Viral Infections, Oxford University Clinical Research Unit | Do,L.A.H., Wilm,A., van Doorn,H.R., Lam,H.M., Sukumaran,R., Tran,A.T., Nguyen,B.H., Tran,T.T.L., Tran,Q.H., Vo,Q.B., Tran Dac,N.A., Trinh,H.N., Nguyen,t.T.H., Le Binh,B.T., Le,K., Nguyen,M.T., Thai,Q.T., Vo,T.V., Ngo,N.Q.M., Dang,t.K.H., Cao,N.H., Tran,T.V., Ho,L.V., Farrar,J., de Jong,M.D., Chen,S., Nagaranjan,N., Bryant,J.E. and Hibberd,M.L. |
| EPI_ISL_2544106 | Medical Microbiology, University Medical Center Utrecht | Medical Microbiology, University Medical Center Utrecht | Tan,L., Viveen,M.C., Lemey,P. and Coenjaerts,F.E. |
| EPI_ISL_2544107 | J. Craig Venter Institute | J. Craig Venter Institute | Lorenzi,H., Town,C., Halpin,R., Bera,J., Ransier,A., Fedorova,N., Stockwell,T., Amedeo,P., Appalla,L., Bishop,B., Edworthy,P., Gupta,N., Hoover,J., Katzel,D., Li,K., Schobel,S., Shrivastava,S., Thovarai,V., Wang,S., Rebuffo-Scheer,C., Fan,J., He,J., Kehl,S.C., Lederboer,N., Jurgens,L.A., Bose,M.E., Beck,E.T., Kumar,S., Neumann-Haefelin,D., Wentworth,D.E. and Henrickson,K.J. |
| EPI_ISL_2544108 | J. Craig Venter Institute | J. Craig Venter Institute | Lorenzi,H., Town,C., Halpin,R., Bera,J., Ransier,A., Fedorova,N., Stockwell,T., Amedeo,P., Appalla,L., Bishop,B., Edworthy,P., Gupta,N., Hoover,J., Katzel,D., Li,K., Schobel,S., Shrivastava,S., Thovarai,V., Wang,S., Rebuffo-Scheer,C., Fan,J., He,J., Kehl,S.C., Lederboer,N., Jurgens,L.A., Bose,M.E., Beck,E.T., Kumar,S., Gerna,G., Wentworth,D.E. and Henrickson,K.J. |
| EPI_ISL_2544109, EPI_ISL_2544110, EPI_ISL_2544111, EPI_ISL_2544112, EPI_ISL_2544113, EPI_ISL_2544114, EPI_ISL_2544115 | Medical Microbiology, University Medical Center Utrecht | Medical Microbiology, University Medical Center Utrecht | Tan,L., Viveen,M.C., Lemey,P. and Coenjaerts,F.E. |
| EPI_ISL_2544121, EPI_ISL_2544122, EPI_ISL_2544123, EPI_ISL_2544124, EPI_ISL_2544125, EPI_ISL_2544126 | Reference Microbiology, Public Health England National Infection Services | Reference Microbiology, Public Health England National Infection Services | Valappil,M., Talts,T., Ellis,J., Sails,A., Eltringham,G., Waugh,S., Gould,K., Harrison,I., Pebody,R. and Zambon,M. |
| EPI_ISL_2544137, EPI_ISL_2544138 | J. Craig Venter Institute | J. Craig Venter Institute | Lorenzi,H., Town,C., Halpin,R., Bera,J., Ransier,A., Fedorova,N., Stockwell,T., Amedeo,P., Appalla,L., Bishop,B., Edworthy,P., Gupta,N., Hoover,J., Katzel,D., Li,K., Schobel,S., Shrivastava,S., Thovarai,V., Wang,S., Rebuffo-Scheer,C., Fan,J., He,J., Kehl,S.C., Lederboer,N., Jurgens,L.A., Bose,M.E., Beck,E.T., Kumar,S., Noyola,D.E., Wentworth,D.E. and Henrickson,K.J. |
| EPI_ISL_2544141, EPI_ISL_2544142, EPI_ISL_2544143, EPI_ISL_2544144, EPI_ISL_2544145, EPI_ISL_2544146 | J. Craig Venter Institute | J. Craig Venter Institute | Das,S.R., Halpin,R.A., Shilts,M., Puri,V., Akopov,A., Fedorova,N., Stockwell,T., Amedeo,P., Bishop,B., Katzel,D., Schobel,S., Shrivastava,S. and Hartert,T. |
| EPI_ISL_2544147 | Lab Medicine, UW | Lab Medicine, UW | Greninger,A.L., Makhsous,N., Kuypers,J.M., Shean,R.C. and Jerome,K.R. |
| EPI_ISL_2544148, EPI_ISL_2544149, EPI_ISL_2544150 | Pediatrics, UT Southwestern Medical Center | Pediatrics, UT Southwestern Medical Center | Levitz,R., Gao,Y., Dozmorov,I., Song,R., Wakeland,E.K. and Kahn,J.S. |
| EPI_ISL_2544151 | MedImmune | MedImmune | Cheng,X., Park,H. and Jin,H. |
| EPI_ISL_2544152 | St. Louis University Medical Center | Viral Vaccine Research | R. Belshe, Karron,R.A., Buonagurio,D.A., Georgiu,A.F., Whitehead,S.S., Adamus,J.E., Clements-Mann,M.L., Harris,D.O., Randolph,V.B., Udem,S.A., Murphy,B.R. and Sidhu,M.S. |
| EPI_ISL_2544153, EPI_ISL_2544154, EPI_ISL_2544155, EPI_ISL_2544156 | J. Craig Venter Institute | J. Craig Venter Institute | Das,S.R., Halpin,R.A., Puri,V., Akopov,A., Fedorova,N., Stockwell,T., Amedeo,P., Bishop,B., Katzel,D., Schobel,S., Shrivastava,S., Wentworth,D.E. and Caserta,M. |
| EPI_ISL_2544157, EPI_ISL_2544158, EPI_ISL_2544159, EPI_ISL_2544160, EPI_ISL_2544161, EPI_ISL_2544162 | J. Craig Venter Institute | J. Craig Venter Institute | Das,S.R., Halpin,R.A., Puri,V., Akopov,A., Fedorova,N., Stockwell,T., Amedeo,P., Bishop,B., Katzel,D., Schobel,S., Shrivastava,S., Hall,C.B., Tesini,B.L., Schnabel,K.C., Walsh,E.E. and Caserta,M. |
| EPI_ISL_2544163, EPI_ISL_2544164, EPI_ISL_2544165 | J. Craig Venter Institute | J. Craig Venter Institute | Shabman,R., Fedorova,N., Puri,V., Shrivastava,S., Amedeo,P., Isom,R., Hu,L., Pickett,B., Novotny,M., Durbin,A., Rocchi,I., Williams,T., Hall,C.B., Tesini,B.L., Schnabel,K.C., Walsh,E.E. and Caserta,M. |
| EPI_ISL_2544171, EPI_ISL_2544172 | Pediatrics - Infectious Diseases, Medical College of Wisconsin | Pediatrics - Infectious Diseases, Medical College of Wisconsin | Rebuffo-Scheer,C., Bose,M.E., He,J., Khaja,S., Ulatowski,M., Beck,E.T., Fan,J., Kumar,S., Nelson,M.I. and Henrickson,K.J. |
| EPI_ISL_2544173, EPI_ISL_2544174, EPI_ISL_2544175, EPI_ISL_2544176, EPI_ISL_2544177, EPI_ISL_2544178 | J. Craig Venter Institute | J. Craig Venter Institute | Shabman,R., Das,S.R., Shilts,M., Fedorova,N., Puri,V., Shrivastava,S., Amedeo,P., Williams,M., Barratt,K., Mitchell,J. and Jennings,L. |
| EPI_ISL_2544180, EPI_ISL_2544181 | J. Craig Venter Institute | J. Craig Venter Institute | Lorenzi,H., Town,C., Halpin,R., Bera,J., Ransier,A., Fedorova,N., Stockwell,T., Amedeo,P., Appalla,L., Bishop,B., Edworthy,P., Gupta,N., Hoover,J., Katzel,D., Li,K., Schobel,S., Shrivastava,S., Thovarai,V., Wang,S., Rebuffo-Scheer,C., Fan,J., He,J., Kehl,S.C., Lederboer,N., Jurgens,L.A., Bose,M.E., Beck,E.T., Kumar,S., Videla,C., Wentworth,D.E. and Henrickson,K.J. |
| EPI_ISL_2544184 | Virology Laboratory, Dr. Ricardo Gutierrez Children Hospital | Virology Laboratory, Dr. Ricardo Gutierrez Children Hospital | Goya,S., Valinotto,L.E., Tittarelli,E., Rojo,G.L., Greninger,A., Luso,S., Natale,M., Mistchenko,A.S. and Viegas,M. |
| EPI_ISL_2544185 | Virology Laboratory, Dr. Ricardo Gutierrez Children Hospital | Virology Laboratory, Dr. Ricardo Gutierrez Children Hospital | Goya,S., Valinotto,L.E., Tittarelli,E., Rojo,G.L., Greninger,A., Zaiat,J., Marti,M., Mistchenko,A.S. and Viegas,M. |
| EPI_ISL_2544186 | Virology Laboratory, Dr. Ricardo Gutierrez Children Hospital | Virology Laboratory, Dr. Ricardo Gutierrez Children Hospital | Goya,S., Rojo,G.L., Valinotto,L.E., Mistchenko,A.S. and Viegas,M. |
| EPI_ISL_2544188, EPI_ISL_2544189 | Microbiology, Institute of Biological Sciences, Universeity of Sao Paulo | Microbiology, Institute of Biological Sciences, Universeity of Sao Paulo | Di Paola,N., Cunha,M.P., Oliveira,D.B.L., Durigon,E., Durigon,G.S. and Zanotto,P.M.A. |
| EPI_ISL_2544190, EPI_ISL_2544191, EPI_ISL_2544192 | J. Craig Venter Institute | J. Craig Venter Institute | Wentworth,D.E., Halpin,R.A., Bera,J., Lin,X., Fedorova,N., Tsitrin,T., McLellan,M., Stockwell,T., Amedeo,P., Bishop,B., Gupta,N., Hoover,J., Katzel,D., Schobel,S., Shrivastava,S., Garcia,J., Laguna-Torres,V.A., Leguia,M., Benavides,J.G. and |

| | | | |
|---|---|---|---|
| | | | Halsey,E. |
| EPI_ISL_2558779, EPI_ISL_2558785, EPI_ISL_2558786, EPI_ISL_2558799 | Kazuya Shirato National Institute of Infectious Diseases, Virology III | Kazuya Shirato National Institute of Infectious Diseases, Virology III | Shirato,K., Sato,K., Dapat,I., Nao,N., Omiya,S., Matsuyama,S., Takeda,M. and Nishimura,H. |
| EPI_ISL_2558800 | J. Craig Venter Institute | J. Craig Venter Institute | Shabman,R., Fedorova,N., Puri,V., Shrivastava,S., Amedeo,P., Isom,R., Hu,L., Pickett,B., Novotny,M., Durbin,A., Rocchi,I., Williams,T., Hall,C.B., Tesini,B.L., Schnabel,K.C., Walsh,E.E. and Caserta,M. |
| EPI_ISL_2558884 | Kazuya Shirato National Institute of Infectious Diseases, Virology III | Kazuya Shirato National Institute of Infectious Diseases, Virology III | Shirato,K., Sato,K., Dapat,I., Nao,N., Omiya,S., Matsuyama,S., Takeda,M. and Nishimura,H. |
| EPI_ISL_2558885 | Pediatrics - Infectious Diseases, Medical College of Wisconsin | Pediatrics - Infectious Diseases, Medical College of Wisconsin | Rebuffo-Scheer,C., Bose,M.E., He,J., Khaja,S., Ulatowski,M., Beck,E.T., Fan,J., Kumar,S., Nelson,M.I. and Henrickson,K.J. |
| EPI_ISL_2558886, EPI_ISL_2558887, EPI_ISL_2558888, EPI_ISL_2558895, EPI_ISL_2558896, EPI_ISL_2558897, EPI_ISL_2558898, EPI_ISL_2558899, EPI_ISL_2558900, EPI_ISL_2558901, EPI_ISL_2558902, EPI_ISL_2558903, EPI_ISL_2558904, EPI_ISL_2558905, EPI_ISL_2558907, EPI_ISL_2558908 | | | |
| see above | Kazuya Shirato National Institute of Infectious Diseases, Virology III | Kazuya Shirato National Institute of Infectious Diseases, Virology III | Shirato,K., Sato,K., Dapat,I., Nao,N., Omiya,S., Matsuyama,S., Takeda,M. and Nishimura,H. |
| EPI_ISL_2558919 | J. Craig Venter Institute | J. Craig Venter Institute | Shabman,R., Fedorova,N., Puri,V., Shrivastava,S., Amedeo,P., Isom,R., Hu,L., Pickett,B., Novotny,M., Durbin,A., Rocchi,I., Williams,T., Hall,C.B., Tesini,B.L., Schnabel,K.C., Walsh,E.E. and Caserta,M. |
| EPI_ISL_2558924, EPI_ISL_2558925, EPI_ISL_2558926 | Kazuya Shirato National Institute of Infectious Diseases, Virology III | Kazuya Shirato National Institute of Infectious Diseases, Virology III | Shirato,K., Sato,K., Dapat,I., Nao,N., Omiya,S., Matsuyama,S., Takeda,M. and Nishimura,H. |
| EPI_ISL_2558927 | Lab Medicine, UW | Lab Medicine, UW | Greninger,A.L., Makhsous,N., Kuypers,J.M., Shean,R.C. and Jerome,K.R. |
| EPI_ISL_2558934, EPI_ISL_2558935, EPI_ISL_2558936 | J. Craig Venter Institute | J. Craig Venter Institute | Shabman,R., Fedorova,N., Puri,V., Shrivastava,S., Amedeo,P., Isom,R., Hu,L., Pickett,B., Novotny,M., Durbin,A., Rocchi,I., Williams,T., Hall,C.B., Tesini,B.L., Schnabel,K.C., Walsh,E.E. and Caserta,M. |
| EPI_ISL_2558971 | Lab Medicine, UW | Lab Medicine, UW | Greninger,A.L., Makhsous,N., Kuypers,J.M., Shean,R.C. and Jerome,K.R. |
| EPI_ISL_2558973, EPI_ISL_2558974, EPI_ISL_2558975, EPI_ISL_2558976 | J. Craig Venter Institute | J. Craig Venter Institute | Shabman,R., Fedorova,N., Puri,V., Shrivastava,S., Amedeo,P., Isom,R., Hu,L., Pickett,B., Novotny,M., Durbin,A., Rocchi,I., Williams,T., Hall,C.B., Tesini,B.L., Schnabel,K.C., Walsh,E.E. and Caserta,M. |
| EPI_ISL_2558985 | Koichi Hashimoto Fukushima Medical University, Department of Pediatrics, School of Medicine | Koichi Hashimoto Fukushima Medical University, Department of Pediatrics, School of Medicine | Fukuyama,Y., Takeshita,F., Norito,S., Hashimoto,K. and Hosoya,M. |
| EPI_ISL_2558987 | Department of Infectious Diseases and Pathobiology, Institute of Virology and Immunology (IVI) | Department of Infectious Diseases and Pathobiology, Institute of Virology and Immunology (IVI) | Thao,T.T.N., Labroussaa,F., Ebert,N., Stalder,H., Dijkman,R., Jores,J., Thiel,V., Bittel,P., Suter-Riniker,F. and Kelly,J. |
| EPI_ISL_2559009, EPI_ISL_2559010, EPI_ISL_2559011 | J. Craig Venter Institute | J. Craig Venter Institute | Shabman,R., Fedorova,N., Puri,V., Shrivastava,S., Amedeo,P., Isom,R., Hu,L., Pickett,B., Novotny,M., Durbin,A., Rocchi,I., Williams,T., Hall,C.B., Tesini,B.L., Schnabel,K.C., Walsh,E.E. and Caserta,M. |
| EPI_ISL_2559012 | Center for Infectious Diseases, School of Public Health, University of Texas Health Science Center | Center for Infectious Diseases, School of Public Health, University of Texas Health Science Center | Bahl,J., Hixson,J., Kim,D.-K., Qiu,X., Piedra,P.A., Piedra,F.-A., Avadhanula,V. and Machado,A.A. |
| EPI_ISL_2559013, EPI_ISL_2559014, EPI_ISL_2559015, EPI_ISL_2559016, EPI_ISL_2559017, EPI_ISL_2559018, EPI_ISL_2559019, EPI_ISL_2559069, EPI_ISL_2559070, EPI_ISL_2559071 | J. Craig Venter Institute | J. Craig Venter Institute | Shabman,R., Fedorova,N., Puri,V., Shrivastava,S., Amedeo,P., Isom,R., Hu,L., Pickett,B., Novotny,M., Durbin,A., Rocchi,I., Williams,T., Hall,C.B., Tesini,B.L., Schnabel,K.C., Walsh,E.E. and Caserta,M. |
| EPI_ISL_2559073 | Kazuya Shirato National Institute of Infectious Diseases, Virology III | Kazuya Shirato National Institute of Infectious Diseases, Virology III | Shirato,K., Sato,K., Dapat,I., Nao,N., Omiya,S., Matsuyama,S., Takeda,M. and Nishimura,H. |
| EPI_ISL_2559080 | Pediatrics - Infectious Diseases, Medical College of Wisconsin | Pediatrics - Infectious Diseases, Medical College of Wisconsin | Rebuffo-Scheer,C., Bose,M.E., He,J., Khaja,S., Ulatowski,M., Beck,E.T., Fan,J., Kumar,S., Nelson,M.I. and Henrickson,K.J. |
| EPI_ISL_2559081, EPI_ISL_2560801, EPI_ISL_2560802 | Kazuya Shirato National Institute of Infectious Diseases, Virology III | Kazuya Shirato National Institute of Infectious Diseases, Virology III | Shirato,K., Sato,K., Dapat,I., Nao,N., Omiya,S., Matsuyama,S., Takeda,M. and Nishimura,H. |
| EPI_ISL_2560803 | Lab Medicine, UW | Lab Medicine, UW | Greninger,A.L., Makhsous,N., Kuypers,J.M., Shean,R.C. and Jerome,K.R. |
| EPI_ISL_2560804 | Virology, University of Washington | Virology, University of Washington | Greninger,A.L., Shean,R.C. and Makhsous,N. |
| EPI_ISL_2560805 | Laboratory Medicine, UW Virology | Laboratory Medicine, UW Virology | Greninger,A.L., Tait,A. and Makhsous,N. |
| EPI_ISL_2560806 | Lab Medicine, UW | Lab Medicine, UW | Greninger,A.L., Makhsous,N., Kuypers,J.M., Shean,R.C. and Jerome,K.R. |
| EPI_ISL_2560807 | Kazuya Shirato National Institute of Infectious Diseases, Virology III | Kazuya Shirato National Institute of Infectious Diseases, Virology III | Shirato,K., Sato,K., Dapat,I., Nao,N., Omiya,S., Matsuyama,S., Takeda,M. and Nishimura,H. |
| EPI_ISL_2560808, EPI_ISL_2560809, EPI_ISL_2560815, EPI_ISL_2560837 | Lab Medicine, UW | Lab Medicine, UW | Greninger,A.L., Makhsous,N., Kuypers,J.M., Shean,R.C. and Jerome,K.R. |
| EPI_ISL_2560838 | Kazuya Shirato National Institute of Infectious Diseases, Virology III | Kazuya Shirato National Institute of Infectious Diseases, Virology III | Shirato,K., Sato,K., Dapat,I., Nao,N., Omiya,S., Matsuyama,S., Takeda,M. and Nishimura,H. |
| EPI_ISL_2560839 | Lab Medicine, UW | Lab Medicine, UW | Greninger,A.L., Makhsous,N., Kuypers,J.M., Shean,R.C. and Jerome,K.R. |
| EPI_ISL_2561008, EPI_ISL_2561009, EPI_ISL_2561010 | Kazuya Shirato National Institute of Infectious Diseases, Virology III | Kazuya Shirato National Institute of Infectious Diseases, Virology III | Shirato,K., Sato,K., Dapat,I., Nao,N., Omiya,S., Matsuyama,S., Takeda,M. and Nishimura,H. |
| EPI_ISL_2561401 | Lab Medicine, UW | Lab Medicine, UW | Greninger,A.L., Makhsous,N., Kuypers,J.M., Shean,R.C. and Jerome,K.R. |
| EPI_ISL_2561453 | Maximum Containment Laboratory, National Institue of Virology | Maximum Containment Laboratory, National Institue of Virology | Yadav,P.D. |
| EPI_ISL_2575367 | Suguru Takeuchi Nagoya University Graduate School of Medicine, Pediatrics | Suguru Takeuchi Nagoya University Graduate School of Medicine, Pediatrics | Takeuchi,S., Kawada,J. and Ito,Y. |
| EPI_ISL_2575417 | Kazuya Shirato National Institute of Infectious Diseases, Virology III | Kazuya Shirato National Institute of Infectious Diseases, Virology III | Shirato,K., Sato,K., Dapat,I., Nao,N., Omiya,S., Matsuyama,S., Takeda,M. and Nishimura,H. |
| EPI_ISL_2575423 | J. Craig Venter Institute | J. Craig Venter Institute | Tan,G., Pickett,B., Fedorova,N., Amedeo,P., Hu,L., Christensen,J., Miller,J., Durbin,A., Williams,T., Arumemi,F., Cadiz,C., Alanis,R., Balmseda,A., Wiliams,T., Schiller,A., Patel,M., Kubale,J. and Gordon,A. |
| EPI_ISL_2575424 | Marie Bashir Institute for Infectious Diseases and Biosecurity & Sydney Medical School, The University of Sydney, Westmead Institute for Medical Research | Marie Bashir Institute for Infectious Diseases and Biosecurity & Sydney Medical School, The University of Sydney, Westmead Institute for Medical Research | Eden,J.-S., Kok,J., Dwyer,D.E., Fernandez,M., Carter,I. and Holmes,E.C. |
| EPI_ISL_2575425 | Virus Research Group, Beijing Pediatric Research Institute, Beijing Children's Hospital, Capital Medical University | Virus Research Group, Beijing Pediatric Research Institute, Beijing Children's Hospital, Capital Medical University | Xu,L. and Xie,Z. |
| EPI_ISL_2575427, EPI_ISL_2575428, EPI_ISL_2575429, EPI_ISL_2575430 | J. Craig Venter Institute | J. Craig Venter Institute | Tan,G., Pickett,B., Fedorova,N., Amedeo,P., Hu,L., Christensen,J., Miller,J., Durbin,A., Williams,T., Arumemi,F., Cadiz,C., Alanis,R., Balmseda,A., Wiliams,T., Schiller,A., Patel,M., Kubale,J. and Gordon,A. |
| EPI_ISL_2575431 | Reference Microbiology, Public Health England National Infection Services | Reference Microbiology, Public Health England National Infection Services | Valappil,M., Talts,T., Ellis,J., Sails,A., Eltringham,G., Waugh,S., Gould,K., Harrison,I., Pebody,R. and Zambon,M. |
| EPI_ISL_2575432 | Marie Bashir Institute for Infectious Diseases and Biosecurity & Sydney Medical School, The University of Sydney, Westmead Institute for Medical Research | Marie Bashir Institute for Infectious Diseases and Biosecurity & Sydney Medical School, The University of Sydney, Westmead Institute for Medical Research | Eden,J.-S., Kok,J., Dwyer,D.E., Fernandez,M., Carter,I. and Holmes,E.C. |
| EPI_ISL_2575463 | J. Craig Venter Institute | J. Craig Venter Institute | Tan,G., Pickett,B., Fedorova,N., Amedeo,P., Hu,L., Christensen,J., Miller,J., Durbin,A., Williams,T., Arumemi,F., Cadiz,C., Alanis,R., Balmseda,A., Wiliams,T., Schiller,A., Patel,M., Kubale,J. and Gordon,A. |
| EPI_ISL_2575465 | Pediatrics, University of New Mexico | Pediatrics, University of New Mexico | Dinwiddle,D. |
| EPI_ISL_2575471 | Michiko Okamoto Graduate School of Medicine, Tohoku University, Virology | Michiko Okamoto Graduate School of Medicine, Tohoku University, Virology | Okamoto,M. and Oshitani,H. |
| EPI_ISL_2575474, EPI_ISL_2575475, EPI_ISL_2575476, EPI_ISL_2575478, EPI_ISL_2575479, EPI_ISL_2575480, EPI_ISL_2575481, EPI_ISL_2575482, EPI_ISL_2575483, EPI_ISL_2575484, EPI_ISL_2575485, EPI_ISL_2575486, EPI_ISL_2575487, EPI_ISL_2575488, EPI_ISL_2575489 | | | |
| see above | Epidemiology and Demography Department, KEMRI Wellcome Trust Research Collaborative Programme | Epidemiology and Demography Department, KEMRI Wellcome Trust Research Collaborative Programme | Agoti,C.N., Phan,M.V.T., Munywoki,P.K., Githinji,G., Medley,G.F., Cane,P.A., Kellam,K., Cotten,M. and Nokes,D.J. |
| EPI_ISL_2575490 | Marie Bashir Institute for Infectious Diseases and Biosecurity & Sydney Medical School, The University of Sydney, Westmead Institute for Medical Research | Marie Bashir Institute for Infectious Diseases and Biosecurity & Sydney Medical School, The University of Sydney, Westmead Institute for Medical Research | Eden,J.-S., Kok,J., Dwyer,D.E., Fernandez,M., Carter,I. and Holmes,E.C. |
| EPI_ISL_2575491 | J. Craig Venter Institute | J. Craig Venter Institute | Tan,G., Pickett,B., Fedorova,N., Amedeo,P., Hu,L., Christensen,J., Miller,J., Durbin,A., Williams,T., Arumemi,F., Cadiz,C., Alanis,R., Balmseda,A., Wiliams,T., Schiller,A., Patel,M., Kubale,J. and Gordon,A. |
| EPI_ISL_2575492 | Reference Microbiology, Public Health England National Infection Services | Reference Microbiology, Public Health England National Infection Services | Valappil,M., Talts,T., Ellis,J., Sails,A., Eltringham,G., Waugh,S., Gould,K., Harrison,I., Pebody,R. and Zambon,M. |
| EPI_ISL_2575493 | J. Craig Venter Institute | J. Craig Venter Institute | Tan,G., Pickett,B., Fedorova,N., Amedeo,P., Isom,R., Hu,L., Christensen,J., Miller,J., Novotny,M., Durbin,A., Rocchi,I., Williams,T., Arumemi,F. and Das,S. |
| EPI_ISL_2575494, EPI_ISL_2575495 | Marie Bashir Institute for Infectious Diseases and | Marie Bashir Institute for Infectious Diseases and | Eden,J.-S., Kok,J., Dwyer,D.E., Fernandez,M., Carter,I. and Holmes,E.C. |

| | | | |
|---|---|---|---|
| | Biosecurity & Sydney Medical School, The University of Sydney, Westmead Institute for Medical Research | Biosecurity & Sydney Medical School, The University of Sydney, Westmead Institute for Medical Research | |
| EPI_ISL_2575496, EPI_ISL_2575497 | Pediatrics, University of New Mexico | Pediatrics, University of New Mexico | Dinwiddle,D. |
| EPI_ISL_2575498 | J. Craig Venter Institute | J. Craig Venter Institute | Tan,G., Pickett,B., Fedorova,N., Amedeo,P., Hu,L., Christensen,J., Miller,J., Durbin,A., Williams,T., Arumemi,F., Cadiz,C., Alanis,R., Balmseda,A., Wiliams,T., Schiller,A., Patel,M., Kubale,J. and Gordon,A. |
| EPI_ISL_2575499 | Department of Experimental Modeling and Infectious Diseases Pathogenesis, Federal Research Center of Fundamental and Translational Medicine | Department of Experimental Modeling and Infectious Diseases Pathogenesis, Federal Research Center of Fundamental and Translational Medicine | Kurskaya,O.G., Sobolev,I.A., Sharshov,K.A., Alexeev,A.Y., Murashkina,T.A., Kabilov,M.R., Alikina,T.Y. and Shestopalov,A.M. |
| EPI_ISL_2575500 | Department of Experimental Modeling and Infectious Diseases Pathogenesis, Federal Research Center of Fundamental and Translational Medicine | Department of Experimental Modeling and Infectious Diseases Pathogenesis, Federal Research Center of Fundamental and Translational Medicine | Sobolev,I.A., Kurskaya,O.G., Sharshov,K.A., Alexeev,A.Y., Murashkina,T.A., Kabilov,M.R., Alikina,T.Y. and Shestopalov,A.M. |
| EPI_ISL_2575505, EPI_ISL_2575506 | Marie Bashir Institute for Infectious Diseases and Biosecurity & Sydney Medical School, The University of Sydney, Westmead Institute for Medical Research | Marie Bashir Institute for Infectious Diseases and Biosecurity & Sydney Medical School, The University of Sydney, Westmead Institute for Medical Research | Eden,J.-S., Kok,J., Dwyer,D.E., Fernandez,M., Carter,I. and Holmes,E.C. |
| EPI_ISL_2575507 | J. Craig Venter Institute | J. Craig Venter Institute | Tan,G., Pickett,B., Fedorova,N., Amedeo,P., Hu,L., Christensen,J., Miller,J., Durbin,A., Williams,T., Arumemi,F., Cadiz,C., Alanis,R., Balmseda,A., Wiliams,T., Schiller,A., Patel,M., Kubale,J. and Gordon,A. |
| EPI_ISL_2575508, EPI_ISL_2575509, EPI_ISL_2575510, EPI_ISL_2575511 | Marie Bashir Institute for Infectious Diseases and Biosecurity & Sydney Medical School, The University of Sydney, Westmead Institute for Medical Research | Marie Bashir Institute for Infectious Diseases and Biosecurity & Sydney Medical School, The University of Sydney, Westmead Institute for Medical Research | Eden,J.-S., Kok,J., Dwyer,D.E., Fernandez,M., Carter,I. and Holmes,E.C. |
| EPI_ISL_2575512 | Michiko Okamoto Graduate School of Medicine, Tohoku University, Virology | Michiko Okamoto Graduate School of Medicine, Tohoku University, Virology | Okamoto,M. and Oshitani,H. |
| EPI_ISL_2575513 | Marie Bashir Institute for Infectious Diseases and Biosecurity & Sydney Medical School, The University of Sydney, Westmead Institute for Medical Research | Marie Bashir Institute for Infectious Diseases and Biosecurity & Sydney Medical School, The University of Sydney, Westmead Institute for Medical Research | Eden,J.-S., Kok,J., Dwyer,D.E., Fernandez,M., Carter,I. and Holmes,E.C. |
| EPI_ISL_2575515, EPI_ISL_2575516, EPI_ISL_2575517 | Reference Microbiology, Public Health England National Infection Services | Reference Microbiology, Public Health England National Infection Services | Valappil,M., Talts,T., Ellis,J., Sails,A., Eltringham,G., Waugh,S., Gould,K., Harrison,I., Pebody,R. and Zambon,M. |
| EPI_ISL_2575518, EPI_ISL_2575522, EPI_ISL_2575523, EPI_ISL_2575524 | Pediatrics, University of New Mexico | Pediatrics, University of New Mexico | Dinwiddle,D. |
| EPI_ISL_2575531 | Center for Infectious Diseases, School of Public Health, University of Texas Health Science Center | Center for Infectious Diseases, School of Public Health, University of Texas Health Science Center | Bahl,J., Hixson,J., Kim,D.-K., Qiu,X., Piedra,P.A., Piedra,F.-A., Avadhanula,V. and Machado,A.A. |
| EPI_ISL_2575532 | Epidemiology and Demography Department, KEMRI Wellcome Trust Research Collaborative Programme | Epidemiology and Demography Department, KEMRI Wellcome Trust Research Collaborative Programme | Agoti,C.N., Phan,M.V.T., Munywoki,P.K., Githinji,G., Medley,G.F., Cane,P.A., Kellam,K., Cotten,M. and Nokes,D.J. |
| EPI_ISL_2575536, EPI_ISL_2575538 | Marie Bashir Institute for Infectious Diseases and Biosecurity & Sydney Medical School, The University of Sydney, Westmead Institute for Medical Research | Marie Bashir Institute for Infectious Diseases and Biosecurity & Sydney Medical School, The University of Sydney, Westmead Institute for Medical Research | Eden,J.-S., Kok,J., Dwyer,D.E., Fernandez,M., Carter,I. and Holmes,E.C. |
| EPI_ISL_2575546, EPI_ISL_2575547 | Department of Experimental Modeling and Infectious Diseases Pathogenesis, Federal Research Center of Fundamental and Translational Medicine | Department of Experimental Modeling and Infectious Diseases Pathogenesis, Federal Research Center of Fundamental and Translational Medicine | Kurskaya,O.G., Sobolev,I.A., Sharshov,K.A., Alexeev,A.Y., Murashkina,T.A., Kabilov,M.R., Alikina,T.Y. and Shestopalov,A.M. |
| EPI_ISL_2575548 | J. Craig Venter Institute | J. Craig Venter Institute | Tan,G., Pickett,B., Fedorova,N., Amedeo,P., Hu,L., Christensen,J., Miller,J., Durbin,A., Williams,T., Arumemi,F., Cadiz,C., Alanis,R., Balmseda,A., Wiliams,T., Schiller,A., Patel,M., Kubale,J. and Gordon,A. |
| EPI_ISL_2575549 | Marie Bashir Institute for Infectious Diseases and Biosecurity & Sydney Medical School, The University of Sydney, Westmead Institute for Medical Research | Marie Bashir Institute for Infectious Diseases and Biosecurity & Sydney Medical School, The University of Sydney, Westmead Institute for Medical Research | Eden,J.-S., Kok,J., Dwyer,D.E., Fernandez,M., Carter,I. and Holmes,E.C. |
| EPI_ISL_2575564 | Reference Microbiology, Public Health England National Infection Services | Reference Microbiology, Public Health England National Infection Services | Valappil,M., Talts,T., Ellis,J., Sails,A., Eltringham,G., Waugh,S., Gould,K., Harrison,I., Pebody,R. and Zambon,M. |
| EPI_ISL_2575565 | Epidemiology and Demography Department, KEMRI Wellcome Trust Research Collaborative Programme | Epidemiology and Demography Department, KEMRI Wellcome Trust Research Collaborative Programme | Agoti,C.N., Phan,M.V.T., Munywoki,P.K., Githinji,G., Medley,G.F., Cane,P.A., Kellam,K., Cotten,M. and Nokes,D.J. |
| EPI_ISL_2575566 | J. Craig Venter Institute | J. Craig Venter Institute | Tan,G., Pickett,B., Fedorova,N., Amedeo,P., Isom,R., Hu,L., Christensen,J., Miller,J., Novotny,M., Durbin,A., Rocchi,I., Williams,T., Arumemi,F. and Das,S. |
| EPI_ISL_2575567 | J. Craig Venter Institute | J. Craig Venter Institute | Tan,G., Pickett,B., Fedorova,N., Amedeo,P., Hu,L., Christensen,J., Miller,J., Durbin,A., Williams,T., Arumemi,F., Cadiz,C., Alanis,R., Balmseda,A., Wiliams,T., Schiller,A., Patel,M., Kubale,J. and Gordon,A. |
| EPI_ISL_2575568, EPI_ISL_2575569 | Epidemiology and Demography Department, KEMRI Wellcome Trust Research Collaborative Programme | Epidemiology and Demography Department, KEMRI Wellcome Trust Research Collaborative Programme | Agoti,C.N., Phan,M.V.T., Munywoki,P.K., Githinji,G., Medley,G.F., Cane,P.A., Kellam,K., Cotten,M. and Nokes,D.J. |
| EPI_ISL_2575570 | J. Craig Venter Institute | J. Craig Venter Institute | Tan,G., Pickett,B., Fedorova,N., Amedeo,P., Hu,L., Christensen,J., Miller,J., Durbin,A., Williams,T., Arumemi,F., Cadiz,C., Alanis,R., Balmseda,A., Wiliams,T., Schiller,A., Patel,M., Kubale,J. and Gordon,A. |
| EPI_ISL_2575571 | Department of Experimental Modeling and Pathogenesis of Infectious Diseases, Federal Research Center of Fundamental and Translational Medicine | Department of Experimental Modeling and Pathogenesis of Infectious Diseases, Federal Research Center of Fundamental and Translational Medicine | Dubovitskiy,N.A., Sobolev,I.A., Kurskaya,O.G., Sharshov,K.A., Anoshina,A.V., Leonova,N.V., Murashkina,T.A., Solomatina,M.V., Derko,A.A., Saroyan,T.A., Kabilov,M.R., Alikina,T.Y. and Shestopalov,A.M. |
| EPI_ISL_2575572, EPI_ISL_2575573, EPI_ISL_2575574, EPI_ISL_2575575, EPI_ISL_2575576, EPI_ISL_2575577, EPI_ISL_2575578, EPI_ISL_2575579 | Epidemiology and Demography Department, KEMRI Wellcome Trust Research Collaborative Programme | Epidemiology and Demography Department, KEMRI Wellcome Trust Research Collaborative Programme | Agoti,C.N., Phan,M.V.T., Munywoki,P.K., Githinji,G., Medley,G.F., Cane,P.A., Kellam,K., Cotten,M. and Nokes,D.J. |
| EPI_ISL_2575580, EPI_ISL_2575581, EPI_ISL_2575582, EPI_ISL_2575583, EPI_ISL_2575584, EPI_ISL_2575585, EPI_ISL_2575586, EPI_ISL_2575587 | J. Craig Venter Institute | J. Craig Venter Institute | Tan,G., Pickett,B., Fedorova,N., Amedeo,P., Hu,L., Christensen,J., Miller,J., Durbin,A., Williams,T., Arumemi,F., Cadiz,C., Alanis,R., Balmseda,A., Wiliams,T., Schiller,A., Patel,M., Kubale,J. and Gordon,A. |
| EPI_ISL_2575588, EPI_ISL_2575589, EPI_ISL_2575590 | Marie Bashir Institute for Infectious Diseases and Biosecurity & Sydney Medical School, The University of Sydney, Westmead Institute for Medical Research | Marie Bashir Institute for Infectious Diseases and Biosecurity & Sydney Medical School, The University of Sydney, Westmead Institute for Medical Research | Eden,J.-S., Kok,J., Dwyer,D.E., Fernandez,M., Carter,I. and Holmes,E.C. |
| EPI_ISL_2575591 | Epidemiology and Demography Department, KEMRI Wellcome Trust Research Collaborative Programme | Epidemiology and Demography Department, KEMRI Wellcome Trust Research Collaborative Programme | Agoti,C.N., Phan,M.V.T., Munywoki,P.K., Githinji,G., Medley,G.F., Cane,P.A., Kellam,K., Cotten,M. and Nokes,D.J. |
| EPI_ISL_2575592 | Reference Microbiology, Public Health England National Infection Services | Reference Microbiology, Public Health England National Infection Services | Valappil,M., Talts,T., Ellis,J., Sails,A., Eltringham,G., Waugh,S., Gould,K., Harrison,I., Pebody,R. and Zambon,M. |
| EPI_ISL_2575593 | Kazuya Shirato National Institute of Infectious Diseases, Virology III | Kazuya Shirato National Institute of Infectious Diseases, Virology III | Shirato,K., Sato,K., Dapat,I., Nao,N., Omiya,S., Matsuyama,S., Takeda,M. and Nishimura,H. |
| EPI_ISL_2575594, EPI_ISL_2575595, EPI_ISL_2575596 | J. Craig Venter Institute | J. Craig Venter Institute | Tan,G., Pickett,B., Fedorova,N., Amedeo,P., Hu,L., Christensen,J., Miller,J., Durbin,A., Williams,T., Arumemi,F., Cadiz,C., Alanis,R., Balmseda,A., Wiliams,T., Schiller,A., Patel,M., Kubale,J. and Gordon,A. |
| EPI_ISL_2575597, EPI_ISL_2575598, EPI_ISL_2575599, EPI_ISL_2575600 | Marie Bashir Institute for Infectious Diseases and Biosecurity & Sydney Medical School, The University of Sydney, Westmead Institute for Medical Research | Marie Bashir Institute for Infectious Diseases and Biosecurity & Sydney Medical School, The University of Sydney, Westmead Institute for Medical Research | Eden,J.-S., Kok,J., Dwyer,D.E., Fernandez,M., Carter,I. and Holmes,E.C. |
| EPI_ISL_2575601 | J. Craig Venter Institute | J. Craig Venter Institute | Tan,G., Pickett,B., Fedorova,N., Amedeo,P., Isom,R., Hu,L., Christensen,J., Miller,J., Novotny,M., Durbin,A., Rocchi,I., Williams,T., Arumemi,F. and Das,S. |
| EPI_ISL_2575602, EPI_ISL_2575603, EPI_ISL_2575604, EPI_ISL_2575605 | Marie Bashir Institute for Infectious Diseases and Biosecurity & Sydney Medical School, The University of Sydney, Westmead Institute for Medical Research | Marie Bashir Institute for Infectious Diseases and Biosecurity & Sydney Medical School, The University of Sydney, Westmead Institute for Medical Research | Eden,J.-S., Kok,J., Dwyer,D.E., Fernandez,M., Carter,I. and Holmes,E.C. |
| EPI_ISL_2575606 | Department of Experimental Modeling and Pathogenesis of Infectious Diseases, Federal Research Center of Fundamental and Translational Medicine | Department of Experimental Modeling and Pathogenesis of Infectious Diseases, Federal Research Center of Fundamental and Translational Medicine | Dubovitskiy,N.A., Sobolev,I.A., Kurskaya,O.G., Sharshov,K.A., Anoshina,A.V., Leonova,N.V., Murashkina,T.A., Solomatina,M.V., Derko,A.A., Saroyan,T.A., Kabilov,M.R., Alikina,T.Y. and Shestopalov,A.M. |
| EPI_ISL_2575607, EPI_ISL_2575608, EPI_ISL_2575609, EPI_ISL_2575610, EPI_ISL_2575611, EPI_ISL_2575612, EPI_ISL_2575613, EPI_ISL_2575614 | Marie Bashir Institute for Infectious Diseases and Biosecurity & Sydney Medical School, The University of Sydney, Westmead Institute for Medical Research | Marie Bashir Institute for Infectious Diseases and Biosecurity & Sydney Medical School, The University of Sydney, Westmead Institute for Medical Research | Eden,J.-S., Kok,J., Dwyer,D.E., Fernandez,M., Carter,I. and Holmes,E.C. |
| EPI_ISL_2575615 | Department of Experimental Modeling and | Department of Experimental Modeling and | Dubovitskiy,N.A., Sobolev,I.A., Kurskaya,O.G., Sharshov,K.A., Anoshina,A.V., Leonova,N.V., Murashkina,T.A., Solomatina,M.V., Derko,A.A., Saroyan,T.A., Kabilov,M.R., Alikina,T.Y. and Shestopalov,A.M. |

| | | | |
|---|---|---|---|
| | Pathogenesis of Infectious Diseases, Federal Research Center of Fundamental and Translational Medicine | Pathogenesis of Infectious Diseases, Federal Research Center of Fundamental and Translational Medicine | |
| EPI_ISL_2575617 | J. Craig Venter Institute | J. Craig Venter Institute | Tan,G., Pickett,B., Fedorova,N., Amedeo,P., Isom,R., Hu,L., Christensen,J., Miller,J., Novotny,M., Durbin,A., Rocchi,I., Williams,T., Arumemi,F. and Das,S. |
| EPI_ISL_2575618, EPI_ISL_2575619, EPI_ISL_2575620 | J. Craig Venter Institute | J. Craig Venter Institute | Tan,G., Pickett,B., Fedorova,N., Amedeo,P., Hu,L., Christensen,J., Miller,J., Durbin,A., Williams,T., Arumemi,F., Cadiz,C., Alanis,R., Balmseda,A., Wiliams,T., Schiller,A., Patel,M., Kubale,J. and Gordon,A. |
| EPI_ISL_2575621, EPI_ISL_2575623, EPI_ISL_2577154, EPI_ISL_2577155 | Marie Bashir Institute for Infectious Diseases and Biosecurity & Sydney Medical School, The University of Sydney, Westmead Institute for Medical Research | Marie Bashir Institute for Infectious Diseases and Biosecurity & Sydney Medical School, The University of Sydney, Westmead Institute for Medical Research | Eden,J.-S., Kok,J., Dwyer,D.E., Fernandez,M., Carter,I. and Holmes,E.C. |
| EPI_ISL_2577156 | Reference Microbiology, Public Health England National Infection Services | Reference Microbiology, Public Health England National Infection Services | Valappil,M., Talts,T., Ellis,J., Sails,A., Eltringham,G., Waugh,S., Gould,K., Harrison,I., Pebody,R. and Zambon,M. |
| EPI_ISL_2577266, EPI_ISL_2577267, EPI_ISL_2577268, EPI_ISL_2577269, EPI_ISL_2577278, EPI_ISL_2577279, EPI_ISL_2577280, EPI_ISL_2577281, EPI_ISL_2577282, EPI_ISL_2577283, EPI_ISL_2577284 | | | |
| see above | Epidemiology and Demography Department, KEMRI Wellcome Trust Research Collaborative Programme | Epidemiology and Demography Department, KEMRI Wellcome Trust Research Collaborative Programme | Agoti,C.N., Phan,M.V.T., Munywoki,P.K., Githinji,G., Medley,G.F., Cane,P.A., Kellam,K., Cotten,M. and Nokes,D.J. |
| EPI_ISL_2577285 | J. Craig Venter Institute | J. Craig Venter Institute | Tan,G., Pickett,B., Fedorova,N., Amedeo,P., Isom,R., Hu,L., Christensen,J., Miller,J., Novotny,M., Durbin,A., Rocchi,I., Williams,T., Arumemi,F. and Das,S. |
| EPI_ISL_2577286, EPI_ISL_2577287, EPI_ISL_2577288 | Reference Microbiology, Public Health England National Infection Services | Reference Microbiology, Public Health England National Infection Services | Valappil,M., Talts,T., Ellis,J., Sails,A., Eltringham,G., Waugh,S., Gould,K., Harrison,I., Pebody,R. and Zambon,M. |
| EPI_ISL_2577289 | J. Craig Venter Institute | J. Craig Venter Institute | Tan,G., Pickett,B., Fedorova,N., Amedeo,P., Isom,R., Hu,L., Christensen,J., Miller,J., Novotny,M., Durbin,A., Rocchi,I., Williams,T., Arumemi,F. and Das,S. |
| EPI_ISL_2577291 | Marie Bashir Institute for Infectious Diseases and Biosecurity & Sydney Medical School, The University of Sydney, Westmead Institute for Medical Research | Marie Bashir Institute for Infectious Diseases and Biosecurity & Sydney Medical School, The University of Sydney, Westmead Institute for Medical Research | Eden,J.-S., Kok,J., Dwyer,D.E., Fernandez,M., Carter,I. and Holmes,E.C. |
| EPI_ISL_2577292, EPI_ISL_2577293, EPI_ISL_2577294, EPI_ISL_2577295, EPI_ISL_2577296, EPI_ISL_2577297, EPI_ISL_2577298, EPI_ISL_2577299 | Epidemiology and Demography Department, KEMRI Wellcome Trust Research Collaborative Programme | Epidemiology and Demography Department, KEMRI Wellcome Trust Research Collaborative Programme | Agoti,C.N., Phan,M.V.T., Munywoki,P.K., Githinji,G., Medley,G.F., Cane,P.A., Kellam,K., Cotten,M. and Nokes,D.J. |
| EPI_ISL_2577300 | Marie Bashir Institute for Infectious Diseases and Biosecurity & Sydney Medical School, The University of Sydney, Westmead Institute for Medical Research | Marie Bashir Institute for Infectious Diseases and Biosecurity & Sydney Medical School, The University of Sydney, Westmead Institute for Medical Research | Eden,J.-S., Kok,J., Dwyer,D.E., Fernandez,M., Carter,I. and Holmes,E.C. |
| EPI_ISL_2577301 | Pediatrics - Infectious Diseases, Medical College of Wisconsin | Pediatrics - Infectious Diseases, Medical College of Wisconsin | Rebuffo-Scheer,C., Bose,M.E., He,J., Khaja,S., Ulatowski,M., Beck,E.T., Fan,J., Kumar,S., Nelson,M.I. and Henrickson,K.J. |
| EPI_ISL_2577302, EPI_ISL_2577303, EPI_ISL_2577304, EPI_ISL_2577305, EPI_ISL_2577306, EPI_ISL_2577307 | Epidemiology and Demography Department, KEMRI Wellcome Trust Research Collaborative Programme | Epidemiology and Demography Department, KEMRI Wellcome Trust Research Collaborative Programme | Agoti,C.N., Phan,M.V.T., Munywoki,P.K., Githinji,G., Medley,G.F., Cane,P.A., Kellam,K., Cotten,M. and Nokes,D.J. |
| EPI_ISL_2577308 | Reference Microbiology, Public Health England National Infection Services | Reference Microbiology, Public Health England National Infection Services | Valappil,M., Talts,T., Ellis,J., Sails,A., Eltringham,G., Waugh,S., Gould,K., Harrison,I., Pebody,R. and Zambon,M. |
| EPI_ISL_2577309, EPI_ISL_2577310 | Michiko Okamoto Graduate School of Medicine, Tohoku University, Virology | Michiko Okamoto Graduate School of Medicine, Tohoku University, Virology | Okamoto,M. and Oshitani,H. |
| EPI_ISL_2577311, EPI_ISL_2577312 | Marie Bashir Institute for Infectious Diseases and Biosecurity & Sydney Medical School, The University of Sydney, Westmead Institute for Medical Research | Marie Bashir Institute for Infectious Diseases and Biosecurity & Sydney Medical School, The University of Sydney, Westmead Institute for Medical Research | Eden,J.-S., Kok,J., Dwyer,D.E., Fernandez,M., Carter,I. and Holmes,E.C. |
| EPI_ISL_2577323 | Epidemiology and Demography Department, KEMRI Wellcome Trust Research Collaborative Programme | Epidemiology and Demography Department, KEMRI Wellcome Trust Research Collaborative Programme | Agoti,C.N., Phan,M.V.T., Munywoki,P.K., Githinji,G., Medley,G.F., Cane,P.A., Kellam,K., Cotten,M. and Nokes,D.J. |
| EPI_ISL_2577324, EPI_ISL_2577325 | Marie Bashir Institute for Infectious Diseases and Biosecurity & Sydney Medical School, The University of Sydney, Westmead Institute for Medical Research | Marie Bashir Institute for Infectious Diseases and Biosecurity & Sydney Medical School, The University of Sydney, Westmead Institute for Medical Research | Eden,J.-S., Kok,J., Dwyer,D.E., Fernandez,M., Carter,I. and Holmes,E.C. |
| EPI_ISL_2577326 | Reference Microbiology, Public Health England National Infection Services | Reference Microbiology, Public Health England National Infection Services | Valappil,M., Talts,T., Ellis,J., Sails,A., Eltringham,G., Waugh,S., Gould,K., Harrison,I., Pebody,R. and Zambon,M. |
| EPI_ISL_2577327, EPI_ISL_2577328, EPI_ISL_2577329, EPI_ISL_2577330 | Marie Bashir Institute for Infectious Diseases and Biosecurity & Sydney Medical School, The University of Sydney, Westmead Institute for Medical Research | Marie Bashir Institute for Infectious Diseases and Biosecurity & Sydney Medical School, The University of Sydney, Westmead Institute for Medical Research | Eden,J.-S., Kok,J., Dwyer,D.E., Fernandez,M., Carter,I. and Holmes,E.C. |
| EPI_ISL_2577331 | Michiko Okamoto Graduate School of Medicine, Tohoku University, Virology | Michiko Okamoto Graduate School of Medicine, Tohoku University, Virology | Okamoto,M. and Oshitani,H. |
| EPI_ISL_2577334 | J. Craig Venter Institute | J. Craig Venter Institute | Tan,G., Pickett,B., Fedorova,N., Amedeo,P., Isom,R., Hu,L., Christensen,J., Miller,J., Novotny,M., Durbin,A., Rocchi,I., Williams,T., Arumemi,F. and Das,S. |
| EPI_ISL_2577335, EPI_ISL_2577336, EPI_ISL_2577337, EPI_ISL_2577338, EPI_ISL_2577339, EPI_ISL_2577382 | Marie Bashir Institute for Infectious Diseases and Biosecurity & Sydney Medical School, The University of Sydney, Westmead Institute for Medical Research | Marie Bashir Institute for Infectious Diseases and Biosecurity & Sydney Medical School, The University of Sydney, Westmead Institute for Medical Research | Eden,J.-S., Kok,J., Dwyer,D.E., Fernandez,M., Carter,I. and Holmes,E.C. |
| EPI_ISL_2577383 | J. Craig Venter Institute | J. Craig Venter Institute | Tan,G., Pickett,B., Fedorova,N., Amedeo,P., Isom,R., Hu,L., Christensen,J., Miller,J., Novotny,M., Durbin,A., Rocchi,I., Williams,T., Arumemi,F. and Das,S. |
| EPI_ISL_2577384 | Michiko Okamoto Graduate School of Medicine, Tohoku University, Virology | Michiko Okamoto Graduate School of Medicine, Tohoku University, Virology | Okamoto,M. and Oshitani,H. |
| EPI_ISL_2577385 | Reference Microbiology, Public Health England National Infection Services | Reference Microbiology, Public Health England National Infection Services | Valappil,M., Talts,T., Ellis,J., Sails,A., Eltringham,G., Waugh,S., Gould,K., Harrison,I., Pebody,R. and Zambon,M. |
| EPI_ISL_2577386, EPI_ISL_2577387 | Marie Bashir Institute for Infectious Diseases and Biosecurity & Sydney Medical School, The University of Sydney, Westmead Institute for Medical Research | Marie Bashir Institute for Infectious Diseases and Biosecurity & Sydney Medical School, The University of Sydney, Westmead Institute for Medical Research | Eden,J.-S., Kok,J., Dwyer,D.E., Fernandez,M., Carter,I. and Holmes,E.C. |
| EPI_ISL_2577388, EPI_ISL_2577389, EPI_ISL_2577390, EPI_ISL_2577391, EPI_ISL_2577392, EPI_ISL_2577393 | Epidemiology and Demography Department, KEMRI Wellcome Trust Research Collaborative Programme | Epidemiology and Demography Department, KEMRI Wellcome Trust Research Collaborative Programme | Agoti,C.N., Phan,M.V.T., Munywoki,P.K., Githinji,G., Medley,G.F., Cane,P.A., Kellam,K., Cotten,M. and Nokes,D.J. |
| EPI_ISL_2577394, EPI_ISL_2577395, EPI_ISL_2577396 | Reference Microbiology, Public Health England National Infection Services | Reference Microbiology, Public Health England National Infection Services | Valappil,M., Talts,T., Ellis,J., Sails,A., Eltringham,G., Waugh,S., Gould,K., Harrison,I., Pebody,R. and Zambon,M. |
| EPI_ISL_2577397, EPI_ISL_2577398, EPI_ISL_2577399 | J. Craig Venter Institute | J. Craig Venter Institute | Tan,G., Pickett,B., Fedorova,N., Amedeo,P., Hu,L., Christensen,J., Miller,J., Durbin,A., Williams,T., Arumemi,F., Cadiz,C., Alanis,R., Balmseda,A., Wiliams,T., Schiller,A., Patel,M., Kubale,J. and Gordon,A. |
| EPI_ISL_2577400, EPI_ISL_2577401, EPI_ISL_2577402 | Marie Bashir Institute for Infectious Diseases and Biosecurity & Sydney Medical School, The University of Sydney, Westmead Institute for Medical Research | Marie Bashir Institute for Infectious Diseases and Biosecurity & Sydney Medical School, The University of Sydney, Westmead Institute for Medical Research | Eden,J.-S., Kok,J., Dwyer,D.E., Fernandez,M., Carter,I. and Holmes,E.C. |
| EPI_ISL_2577403, EPI_ISL_2577404, EPI_ISL_2577405 | Epidemiology and Demography Department, KEMRI Wellcome Trust Research Collaborative Programme | Epidemiology and Demography Department, KEMRI Wellcome Trust Research Collaborative Programme | Agoti,C.N., Phan,M.V.T., Munywoki,P.K., Githinji,G., Medley,G.F., Cane,P.A., Kellam,K., Cotten,M. and Nokes,D.J. |
| EPI_ISL_2577406 | J. Craig Venter Institute | J. Craig Venter Institute | Tan,G., Pickett,B., Fedorova,N., Amedeo,P., Isom,R., Hu,L., Christensen,J., Miller,J., Novotny,M., Durbin,A., Rocchi,I., Williams,T., Arumemi,F. and Das,S. |
| EPI_ISL_2577407 | Marie Bashir Institute for Infectious Diseases and Biosecurity & Sydney Medical School, The University of Sydney, Westmead Institute for Medical Research | Marie Bashir Institute for Infectious Diseases and Biosecurity & Sydney Medical School, The University of Sydney, Westmead Institute for Medical Research | Eden,J.-S., Kok,J., Dwyer,D.E., Fernandez,M., Carter,I. and Holmes,E.C. |
| EPI_ISL_2577408 | Reference Microbiology, Public Health England National Infection Services | Reference Microbiology, Public Health England National Infection Services | Valappil,M., Talts,T., Ellis,J., Sails,A., Eltringham,G., Waugh,S., Gould,K., Harrison,I., Pebody,R. and Zambon,M. |
| EPI_ISL_2577409, EPI_ISL_2577410 | Marie Bashir Institute for Infectious Diseases and Biosecurity & Sydney Medical School, The University of Sydney, Westmead Institute for Medical Research | Marie Bashir Institute for Infectious Diseases and Biosecurity & Sydney Medical School, The University of Sydney, Westmead Institute for Medical Research | Eden,J.-S., Kok,J., Dwyer,D.E., Fernandez,M., Carter,I. and Holmes,E.C. |
| EPI_ISL_2577411, EPI_ISL_2577412, EPI_ISL_2577413 | J. Craig Venter Institute | J. Craig Venter Institute | Tan,G., Pickett,B., Fedorova,N., Amedeo,P., Hu,L., Christensen,J., Miller,J., Durbin,A., Williams,T., Arumemi,F., Cadiz,C., Alanis,R., Balmseda,A., Wiliams,T., Schiller,A., Patel,M., Kubale,J. and Gordon,A. |
| EPI_ISL_2577414, EPI_ISL_2577415 | Marie Bashir Institute for Infectious Diseases and | Marie Bashir Institute for Infectious Diseases and | Eden,J.-S., Kok,J., Dwyer,D.E., Fernandez,M., Carter,I. and Holmes,E.C. |

| | | | |
|---|---|---|---|
| | Biosecurity & Sydney Medical School, The University of Sydney, Westmead Institute for Medical Research | Biosecurity & Sydney Medical School, The University of Sydney, Westmead Institute for Medical Research | |
| EPI_ISL_2577416, EPI_ISL_2577417, EPI_ISL_2577418, EPI_ISL_2577419 | Epidemiology and Demography Department, KEMRI Wellcome Trust Research Collaborative Programme | Epidemiology and Demography Department, KEMRI Wellcome Trust Research Collaborative Programme | Agoti,C.N., Phan,M.V.T., Munywoki,P.K., Githinji,G., Medley,G.F., Cane,P.A., Kellam,K., Cotten,M. and Nokes,D.J. |
| EPI_ISL_2577420 | Marie Bashir Institute for Infectious Diseases and Biosecurity & Sydney Medical School, The University of Sydney, Westmead Institute for Medical Research | Marie Bashir Institute for Infectious Diseases and Biosecurity & Sydney Medical School, The University of Sydney, Westmead Institute for Medical Research | Eden,J.-S., Kok,J., Dwyer,D.E., Fernandez,M., Carter,I. and Holmes,E.C. |
| EPI_ISL_2577421, EPI_ISL_2577422 | Reference Microbiology, Public Health England National Infection Services | Reference Microbiology, Public Health England National Infection Services | Valappil,M., Talts,T., Ellis,J., Sails,A., Eltringham,G., Waugh,S., Gould,K., Harrison,I., Pebody,R. and Zambon,M. |
| EPI_ISL_2577423, EPI_ISL_2577424 | Marie Bashir Institute for Infectious Diseases and Biosecurity & Sydney Medical School, The University of Sydney, Westmead Institute for Medical Research | Marie Bashir Institute for Infectious Diseases and Biosecurity & Sydney Medical School, The University of Sydney, Westmead Institute for Medical Research | Eden,J.-S., Kok,J., Dwyer,D.E., Fernandez,M., Carter,I. and Holmes,E.C. |
| EPI_ISL_2577425 | J. Craig Venter Institute | J. Craig Venter Institute | Tan,G., Pickett,B., Fedorova,N., Amedeo,P., Hu,L., Christensen,J., Miller,J., Durbin,A., Williams,T., Arumemi,F., Cadiz,C., Alanis,R., Balmseda,A., Wiliams,T., Schiller,A., Patel,M., Kubale,J. and Gordon,A. |
| EPI_ISL_2577426 | Reference Microbiology, Public Health England National Infection Services | Reference Microbiology, Public Health England National Infection Services | Valappil,M., Talts,T., Ellis,J., Sails,A., Eltringham,G., Waugh,S., Gould,K., Harrison,I., Pebody,R. and Zambon,M. |
| EPI_ISL_2577427, EPI_ISL_2577428 | Marie Bashir Institute for Infectious Diseases and Biosecurity & Sydney Medical School, The University of Sydney, Westmead Institute for Medical Research | Marie Bashir Institute for Infectious Diseases and Biosecurity & Sydney Medical School, The University of Sydney, Westmead Institute for Medical Research | Eden,J.-S., Kok,J., Dwyer,D.E., Fernandez,M., Carter,I. and Holmes,E.C. |
| EPI_ISL_2577429 | J. Craig Venter Institute | J. Craig Venter Institute | Tan,G., Pickett,B., Fedorova,N., Amedeo,P., Hu,L., Christensen,J., Miller,J., Durbin,A., Williams,T., Arumemi,F., Cadiz,C., Alanis,R., Balmseda,A., Wiliams,T., Schiller,A., Patel,M., Kubale,J. and Gordon,A. |
| EPI_ISL_2577430 | Marie Bashir Institute for Infectious Diseases and Biosecurity & Sydney Medical School, The University of Sydney, Westmead Institute for Medical Research | Marie Bashir Institute for Infectious Diseases and Biosecurity & Sydney Medical School, The University of Sydney, Westmead Institute for Medical Research | Eden,J.-S., Kok,J., Dwyer,D.E., Fernandez,M., Carter,I. and Holmes,E.C. |
| EPI_ISL_2577431, EPI_ISL_2577432, EPI_ISL_2577433, EPI_ISL_2577434 | Epidemiology and Demography Department, KEMRI Wellcome Trust Research Collaborative Programme | Epidemiology and Demography Department, KEMRI Wellcome Trust Research Collaborative Programme | Agoti,C.N., Phan,M.V.T., Munywoki,P.K., Githinji,G., Medley,G.F., Cane,P.A., Kellam,K., Cotten,M. and Nokes,D.J. |
| EPI_ISL_2577435 | Marie Bashir Institute for Infectious Diseases and Biosecurity & Sydney Medical School, The University of Sydney, Westmead Institute for Medical Research | Marie Bashir Institute for Infectious Diseases and Biosecurity & Sydney Medical School, The University of Sydney, Westmead Institute for Medical Research | Eden,J.-S., Kok,J., Dwyer,D.E., Fernandez,M., Carter,I. and Holmes,E.C. |
| EPI_ISL_2577436, EPI_ISL_2577437 | J. Craig Venter Institute | J. Craig Venter Institute | Tan,G., Pickett,B., Fedorova,N., Amedeo,P., Isom,R., Hu,L., Christensen,J., Miller,J., Novotny,M., Durbin,A., Rocchi,I., Williams,T., Arumemi,F. and Das,S. |
| EPI_ISL_2577438, EPI_ISL_2577439, EPI_ISL_2577440 | Marie Bashir Institute for Infectious Diseases and Biosecurity & Sydney Medical School, The University of Sydney, Westmead Institute for Medical Research | Marie Bashir Institute for Infectious Diseases and Biosecurity & Sydney Medical School, The University of Sydney, Westmead Institute for Medical Research | Eden,J.-S., Kok,J., Dwyer,D.E., Fernandez,M., Carter,I. and Holmes,E.C. |
| EPI_ISL_2577441, EPI_ISL_2577442, EPI_ISL_2577443 | J. Craig Venter Institute | J. Craig Venter Institute | Tan,G., Pickett,B., Fedorova,N., Amedeo,P., Hu,L., Christensen,J., Miller,J., Durbin,A., Williams,T., Arumemi,F., Cadiz,C., Alanis,R., Balmseda,A., Wiliams,T., Schiller,A., Patel,M., Kubale,J. and Gordon,A. |
| EPI_ISL_2577444 | Marie Bashir Institute for Infectious Diseases and Biosecurity & Sydney Medical School, The University of Sydney, Westmead Institute for Medical Research | Marie Bashir Institute for Infectious Diseases and Biosecurity & Sydney Medical School, The University of Sydney, Westmead Institute for Medical Research | Eden,J.-S., Kok,J., Dwyer,D.E., Fernandez,M., Carter,I. and Holmes,E.C. |
| EPI_ISL_2577471, EPI_ISL_2577472, EPI_ISL_2577473, EPI_ISL_2577474, EPI_ISL_2577475, EPI_ISL_2577476, EPI_ISL_2577477, EPI_ISL_2577478, EPI_ISL_2577479, EPI_ISL_2577480 | Epidemiology and Demography Department, KEMRI Wellcome Trust Research Collaborative Programme | Epidemiology and Demography Department, KEMRI Wellcome Trust Research Collaborative Programme | Agoti,C.N., Phan,M.V.T., Munywoki,P.K., Githinji,G., Medley,G.F., Cane,P.A., Kellam,K., Cotten,M. and Nokes,D.J. |
| EPI_ISL_2577483, EPI_ISL_2577484 | Marie Bashir Institute for Infectious Diseases and Biosecurity & Sydney Medical School, The University of Sydney, Westmead Institute for Medical Research | Marie Bashir Institute for Infectious Diseases and Biosecurity & Sydney Medical School, The University of Sydney, Westmead Institute for Medical Research | Eden,J.-S., Kok,J., Dwyer,D.E., Fernandez,M., Carter,I. and Holmes,E.C. |
| EPI_ISL_2577662 | Reference Microbiology, Public Health England National Infection Services | Reference Microbiology, Public Health England National Infection Services | Valappil,M., Talts,T., Ellis,J., Sails,A., Eltringham,G., Waugh,S., Gould,K., Harrison,I., Pebody,R. and Zambon,M. |
| EPI_ISL_2577698, EPI_ISL_2577709, EPI_ISL_2577719, EPI_ISL_2577730, EPI_ISL_2577736, EPI_ISL_2577749, EPI_ISL_2577754 | J. Craig Venter Institute | J. Craig Venter Institute | Tan,G., Pickett,B., Fedorova,N., Amedeo,P., Isom,R., Hu,L., Christensen,J., Miller,J., Novotny,M., Durbin,A., Rocchi,I., Williams,T., Arumemi,F. and Das,S. |
| EPI_ISL_2577788 | Central Laboratory, Guangzhou Women and Children's Medical Center | Central Laboratory, Guangzhou Women and Children's Medical Center | Xie,J.H., Zhu,B., Zhong,J.Y., Chen,Y. and Zhang,Y.Y. |
| EPI_ISL_2578154, EPI_ISL_2578157 | Michiko Okamoto Graduate School of Medicine, Tohoku University, Virology | Michiko Okamoto Graduate School of Medicine, Tohoku University, Virology | Okamoto,M. and Oshitani,H. |
| EPI_ISL_2578161 | Marie Bashir Institute for Infectious Diseases and Biosecurity & Sydney Medical School, The University of Sydney, Westmead Institute for Medical Research | Marie Bashir Institute for Infectious Diseases and Biosecurity & Sydney Medical School, The University of Sydney, Westmead Institute for Medical Research | Eden,J.-S., Kok,J., Dwyer,D.E., Fernandez,M., Carter,I. and Holmes,E.C. |
| EPI_ISL_2578167 | Michiko Okamoto Graduate School of Medicine, Tohoku University, Virology | Michiko Okamoto Graduate School of Medicine, Tohoku University, Virology | Okamoto,M. and Oshitani,H. |
| EPI_ISL_2578239 | Marie Bashir Institute for Infectious Diseases and Biosecurity & Sydney Medical School, The University of Sydney, Westmead Institute for Medical Research | Marie Bashir Institute for Infectious Diseases and Biosecurity & Sydney Medical School, The University of Sydney, Westmead Institute for Medical Research | Eden,J.-S., Kok,J., Dwyer,D.E., Fernandez,M., Carter,I. and Holmes,E.C. |
| EPI_ISL_2578352, EPI_ISL_2578361, EPI_ISL_2578362, EPI_ISL_2578365 | J. Craig Venter Institute | J. Craig Venter Institute | Tan,G., Pickett,B., Fedorova,N., Amedeo,P., Hu,L., Christensen,J., Miller,J., Durbin,A., Williams,T., Arumemi,F., Cadiz,C., Alanis,R., Balmseda,A., Wiliams,T., Schiller,A., Patel,M., Kubale,J. and Gordon,A. |
| EPI_ISL_2578381 | Marie Bashir Institute for Infectious Diseases and Biosecurity & Sydney Medical School, The University of Sydney, Westmead Institute for Medical Research | Marie Bashir Institute for Infectious Diseases and Biosecurity & Sydney Medical School, The University of Sydney, Westmead Institute for Medical Research | Eden,J.-S., Kok,J., Dwyer,D.E., Fernandez,M., Carter,I. and Holmes,E.C. |
| EPI_ISL_2584483, EPI_ISL_2584484, EPI_ISL_2584485, EPI_ISL_2584486 | J. Craig Venter Institute | J. Craig Venter Institute | Shabman,R., Das,S.R., Puri,V., Fedorova,N., Amedeo,P., Williams,M., Shrivastava,S. and Halasa,N. |
| EPI_ISL_2584489, EPI_ISL_2584491, EPI_ISL_2584492, EPI_ISL_2584494, EPI_ISL_2584496, EPI_ISL_2584497, EPI_ISL_2584498, EPI_ISL_2584499, EPI_ISL_2584500, EPI_ISL_2584504 | Epidemiology and Demography, KEMRI-Wellcome Trust | Epidemiology and Demography, KEMRI-Wellcome Trust | Kamau,E., Otieno,J.R., Murunga,N., Nyiro,J.U., Oketch,J.W., Ngoi,J.M., de Laurent,Z.R., Mwema,A., Agoti,C.N. and Nokes,D.J. |
| EPI_ISL_2584505 | Medical Microbiology, University Medical Center Utrecht | Medical Microbiology, University Medical Center Utrecht | Tan,L., Viveen,M.C., Lemey,P. and Coenjaerts,F.E. |
| EPI_ISL_2584506 | Medical Microbiology, University Medical Center Utrecht | Medical Microbiology, University Medical Center Utrecht | Tan,L., Lemey,P., Viveen,M. and Coenjaerts,F. |
| EPI_ISL_2584511, EPI_ISL_2584512, EPI_ISL_2584513, EPI_ISL_2584514, EPI_ISL_2584515, EPI_ISL_2584516 | J. Craig Venter Institute | J. Craig Venter Institute | Shabman,R., Das,S.R., Shilts,M., Fedorova,N., Puri,V., Shrivastava,S., Amedeo,P., Williams,M., Barratt,K., Mitchell,J. and Jennings,L. |
| EPI_ISL_2584518 | J. Craig Venter Institute | J. Craig Venter Institute | Wentworth,D.E., Halpin,R.A., Bera,J., Lin,X., Fedorova,N., Tsitrin,T., McLellan,M., Stockwell,T., Amedeo,P., Bishop,B., Gupta,N., Hoover,J., Katzel,D., Schobel,S., Shrivastava,S., Garcia,J., Laguna-Torres,V.A., Leguia,M., Benavides,J.G. and Halsey,E. |
| EPI_ISL_2584558, EPI_ISL_2584560, EPI_ISL_2584562, EPI_ISL_2584565, EPI_ISL_2584566, EPI_ISL_2584568, EPI_ISL_2584569, EPI_ISL_2584570 | J. Craig Venter Institute | J. Craig Venter Institute | Das,S.R., Halpin,R.A., Shilts,M., Puri,V., Akopov,A., Fedorova,N., Stockwell,T., Amedeo,P., Bishop,B., Katzel,D., Schobel,S., Shrivastava,S. and Hartert,T. |
| EPI_ISL_2584571, EPI_ISL_2584572 | J. Craig Venter Institute | J. Craig Venter Institute | Wentworth,D.E., Halpin,R.A., Bera,J., Lin,X., Fedorova,N., Tsitrin,T., McLellan,M., Stockwell,T., Amedeo,P., Bishop,B., Gupta,N., Hoover,J., Katzel,D., Schobel,S., Shrivastava,S., Garcia,J., Laguna-Torres,V.A., Leguia,M., Benavides,J.G. and Halsey,E. |

| | | | |
|---|---|---|---|
| EPI_ISL_2584579, EPI_ISL_2584582 | Medicine, University of Washington, 300 9th Ave, Harborview Research & Training Building | Medicine, University of Washington, 300 9th Ave, Harborview Research & Training Building | Chu,H., Scott,E. and Roychoudhury,P. |
| EPI_ISL_2584591, EPI_ISL_2584592, EPI_ISL_2584593, EPI_ISL_2584594 | Epidemiology and Demography, KEMRI-Wellcome Trust | Epidemiology and Demography, KEMRI-Wellcome Trust | Kamau,E., Otieno,J.R., Murunga,N., Nyiro,J.U., Oketch,J.W., Ngoi,J.M., de Laurent,Z.R., Mwema,A., Agoti,C.N. and Nokes,D.J. |
| EPI_ISL_2584595 | J. Craig Venter Institute | J. Craig Venter Institute | Das,S.R., Halpin,R.A., Shilts,M., Puri,V., Akopov,A., Fedorova,N., Stockwell,T., Amedeo,P., Bishop,B., Katzel,D., Schobel,S., Shrivastava,S. and Hartert,T. |
| EPI_ISL_2584596 | J. Craig Venter Institute | J. Craig Venter Institute | Das,S.R., Halpin,R.A., Puri,V., Akopov,A., Fedorova,N., Stockwell,T., Amedeo,P., Bishop,B., Katzel,D., Schobel,S., Shrivastava,S., Hall,C.B., Tesini,B.L., Schnabel,K.C., Walsh,E.E. and Caserta,M. |
| EPI_ISL_2584597 | Pediatrics, UT Southwestern Medical Center | Pediatrics, UT Southwestern Medical Center | Levitz,R., Gao,Y., Dozmorov,I., Song,R., Wakeland,E.K. and Kahn,J.S. |
| EPI_ISL_2584598 | Medical Microbiology, University Medical Center Utrecht | Medical Microbiology, University Medical Center Utrecht | Tan,L., Lemey,P., Viveen,M. and Coenjaerts,F.E.J. |
| EPI_ISL_2584599, EPI_ISL_2584600 | J. Craig Venter Institute | J. Craig Venter Institute | Das,S.R., Halpin,R.A., Puri,V., Akopov,A., Fedorova,N., Stockwell,T., Amedeo,P., Bishop,B., Katzel,D., Schobel,S., Shrivastava,S., Hall,C.B., Tesini,B.L., Schnabel,K.C., Walsh,E.E. and Caserta,M. |
| EPI_ISL_2584601 | J. Craig Venter Institute | J. Craig Venter Institute | Das,S.R., Halpin,R.A., Puri,V., Akopov,A., Fedorova,N., Stockwell,T., Amedeo,P., Bishop,B., Katzel,D., Schobel,S., Shrivastava,S., Wentworth,D.E. and Caserta,M. |
| EPI_ISL_2584602 | Medical Microbiology, University Medical Center Utrecht | Medical Microbiology, University Medical Center Utrecht | Tan,L., Lemey,P., Viveen,M. and Coenjaerts,F.E.J. |
| EPI_ISL_2584604 | J. Craig Venter Institute | J. Craig Venter Institute | Das,S.R., Halpin,R.A., Puri,V., Akopov,A., Fedorova,N., Tsitrin,T., Stockwell,T., Amedeo,P., Bishop,B., Gupta,N., Hoover,J., Katzel,D., Schobel,S., Shrivastava,S., Wentworth,D.E. and Caserta,M. |
| EPI_ISL_2584605, EPI_ISL_2584606, EPI_ISL_2584610, EPI_ISL_2584611, EPI_ISL_2584613, EPI_ISL_2584614 | J. Craig Venter Institute | J. Craig Venter Institute | Das,S.R., Halpin,R.A., Puri,V., Akopov,A., Fedorova,N., Stockwell,T., Amedeo,P., Bishop,B., Katzel,D., Schobel,S., Shrivastava,S., Hall,C.B., Tesini,B.L., Schnabel,K.C., Walsh,E.E. and Caserta,M. |
| EPI_ISL_2584615 | J. Craig Venter Institute | J. Craig Venter Institute | Das,S.R., Halpin,R.A., Puri,V., Akopov,A., Fedorova,N., Stockwell,T., Amedeo,P., Bishop,B., Katzel,D., Schobel,S., Shrivastava,S., Wentworth,D.E. and Caserta,M. |
| EPI_ISL_2584617 | Pediatrics and Microbiology, University of Texas Southwestern Medical Center | Pediatrics and Microbiology, University of Texas Southwestern Medical Center | Levitz,R., Wattier,R., Phillips,P., Solomon,A., Lawler,J., Lazar,I. and Kahn,J.S. |
| EPI_ISL_2584618 | J. Craig Venter Institute | J. Craig Venter Institute | Das,S.R., Halpin,R.A., Puri,V., Akopov,A., Fedorova,N., Stockwell,T., Amedeo,P., Bishop,B., Katzel,D., Schobel,S., Shrivastava,S., Wentworth,D.E. and Caserta,M. |
| EPI_ISL_2584619 | J. Craig Venter Institute | J. Craig Venter Institute | Das,S.R., Halpin,R.A., Puri,V., Akopov,A., Fedorova,N., Stockwell,T., Amedeo,P., Bishop,B., Katzel,D., Schobel,S., Shrivastava,S., Hall,C.B., Tesini,B.L., Schnabel,K.C., Walsh,E.E. and Caserta,M. |
| EPI_ISL_2584620 | J. Craig Venter Institute | J. Craig Venter Institute | Das,S.R., Halpin,R.A., Puri,V., Akopov,A., Fedorova,N., Stockwell,T., Amedeo,P., Bishop,B., Katzel,D., Schobel,S., Shrivastava,S., Wentworth,D.E. and Caserta,M. |
| EPI_ISL_2584621 | Biologia Molecular y Validacion de Tecnicas, Instituto de Diagnostico y Referencia Epidemiologicos (InDRE) | Biologia Molecular y Validacion de Tecnicas, Instituto de Diagnostico y Referencia Epidemiologicos (InDRE) | Ortiz-Alcantara,J.M., Garces-Ayala,F., Perez-Agueros,S.I., Hernandez-Moreno,A.L., Munoz-Medina,J.E., Monroy-Munoz,I.E., Santos Coy-Arechavaleta,A., Meza-Chavez,A., Angeles-Martinez,J., Anguiano-Hernandez,Y.-M., Martinez-Miguel,B., Santacruz-Tinoco,C.E., Gonzalez-Ibarra,J., Alvarado-Yaah,J.E., Gonzalez-Bonilla,C.R., Diaz-Quinonez,J.A. and Ramirez-Gonzalez,J.E. |
| EPI_ISL_2584622 | Medical Microbiology, University Medical Center Utrecht | Medical Microbiology, University Medical Center Utrecht | Tan,L., Viveen,M.C., Lemey,P. and Coenjaerts,F.E. |
| EPI_ISL_2584623 | J. Craig Venter Institute | J. Craig Venter Institute | Wentworth,D.E., Halpin,R.A., Bera,J., Lin,X., Fedorova,N., Tsitrin,T., McLellan,M., Stockwell,T., Amedeo,P., Bishop,B., Gupta,N., Hoover,J., Katzel,D., Schobel,S., Shrivastava,S., Garcia,J., Laguna-Torres,V.A., Leguia,M., Benavides,J.G. and Halsey,E. |
| EPI_ISL_2584624 | Medical Microbiology, University Medical Center Utrecht | Medical Microbiology, University Medical Center Utrecht | Tan,L., Viveen,M.C., Lemey,P. and Coenjaerts,F.E. |
| EPI_ISL_2584626 | KEMRI Wellcome Trust Research Programme | KEMRI Wellcome Trust Research Programme | Agoti,C.N., Otieno,J.R., Munywoki,P.K., Mwihuri,A.G., Cane,P.A., Nokes,D.J., Kellam,P. and Cotten,M.L. |
| EPI_ISL_2584627 | Virology, Public Health Institution of Turkey | Virology, Public Health Institution of Turkey | Bayrakdar,F. |
| EPI_ISL_2584628 | J. Craig Venter Institute | J. Craig Venter Institute | Shabman,R., Das,S.R., Shilts,M., Fedorova,N., Puri,V., Shrivastava,S., Amedeo,P., Williams,M., Barratt,K., Mitchell,J. and Jennings,L. |
| EPI_ISL_2584630 | J. Craig Venter Institute | J. Craig Venter Institute | Wentworth,D.E., Halpin,R.A., Bera,J., Lin,X., Fedorova,N., Tsitrin,T., McLellan,M., Stockwell,T., Amedeo,P., Bishop,B., Gupta,N., Hoover,J., Katzel,D., Schobel,S., Shrivastava,S., Garcia,J., Laguna-Torres,V.A., Leguia,M., Benavides,J.G. and Halsey,E. |
| EPI_ISL_2584631, EPI_ISL_2584632 | Medical Microbiology, University Medical Center Utrecht | Medical Microbiology, University Medical Center Utrecht | Tan,L., Viveen,M.C., Lemey,P. and Coenjaerts,F.E. |
| EPI_ISL_2584633 | J. Craig Venter Institute | J. Craig Venter Institute | Shabman,R., Das,S.R., Shilts,M., Fedorova,N., Puri,V., Shrivastava,S., Amedeo,P., Williams,M., Barratt,K., Mitchell,J. and Jennings,L. |
| EPI_ISL_2584634 | Medical Microbiology, University Medical Center Utrecht | Medical Microbiology, University Medical Center Utrecht | Tan,L., Lemey,P., Viveen,M. and Coenjaerts,F.E.J. |
| EPI_ISL_2584635 | J. Craig Venter Institute | J. Craig Venter Institute | Das,S.R., Halpin,R.A., Shilts,M., Puri,V., Akopov,A., Fedorova,N., Stockwell,T., Amedeo,P., Bishop,B., Katzel,D., Schobel,S., Shrivastava,S. and Hartert,T. |
| EPI_ISL_2584636, EPI_ISL_2584637 | J. Craig Venter Institute | J. Craig Venter Institute | Shabman,R., Das,S.R., Shilts,M., Fedorova,N., Puri,V., Shrivastava,S., Amedeo,P., Williams,M., Barratt,K., Mitchell,J. and Jennings,L. |
| EPI_ISL_2584638 | J. Craig Venter Institute | J. Craig Venter Institute | Shabman,R., Das,S.R., Puri,V., Fedorova,N., Amedeo,P., Williams,M., Shrivastava,S. and Halasa,N. |
| EPI_ISL_2584639 | Biologia Molecular y Validacion de Tecnicas, Instituto de Diagnostico y Referencia Epidemiologicos (InDRE) | Biologia Molecular y Validacion de Tecnicas, Instituto de Diagnostico y Referencia Epidemiologicos (InDRE) | Munoz-Medina,J.E., Monroy-Munoz,I.E., Santos Coy-Arechavaleta,A., Meza-Chavez,A., Angeles-Martinez,J., Anguiano-Hernandez,Y.M., Santacruz-Tinoco,C.E., Gonzalez-Ibarra,J., Martinez-Miguel,B., Alvarado-Yaah,J.E., Palomec-Nava,I.D., Ortiz-Alcantara,J.M., Garces-Ayala,F., Ramirez-Gonzalez,J.E., Diaz-Quinonez,J.A. and Gonzalez-Bonilla,C.R. |
| EPI_ISL_2584640, EPI_ISL_2584641 | J. Craig Venter Institute | J. Craig Venter Institute | Shabman,R., Das,S.R., Puri,V., Fedorova,N., Amedeo,P., Williams,M., Shrivastava,S. and Halasa,N. |
| EPI_ISL_2584642 | Medical Microbiology, University Medical Center Utrecht | Medical Microbiology, University Medical Center Utrecht | Tan,L., Viveen,M.C., Lemey,P. and Coenjaerts,F.E. |
| EPI_ISL_2584644, EPI_ISL_2584646, EPI_ISL_2584648, EPI_ISL_2584650 | J. Craig Venter Institute | J. Craig Venter Institute | Das,S.R., Halpin,R.A., Shilts,M., Puri,V., Akopov,A., Fedorova,N., Stockwell,T., Amedeo,P., Bishop,B., Katzel,D., Schobel,S., Shrivastava,S. and Hartert,T. |
| EPI_ISL_2584652 | J. Craig Venter Institute | J. Craig Venter Institute | Das,S.R., Halpin,R.A., Puri,V., Akopov,A., Fedorova,N., Tsitrin,T., Stockwell,T., Amedeo,P., Bishop,B., Gupta,N., Hoover,J., Katzel,D., Schobel,S., Shrivastava,S., Wentworth,D.E. and Caserta,M. |
| EPI_ISL_2584653, EPI_ISL_2584654 | Epidemiology and Demography, KEMRI-Wellcome Trust | Epidemiology and Demography, KEMRI-Wellcome Trust | Kamau,E., Otieno,J.R., Murunga,N., Nyiro,J.U., Oketch,J.W., Ngoi,J.M., de Laurent,Z.R., Mwema,A., Agoti,C.N. and Nokes,D.J. |
| EPI_ISL_2584656 | J. Craig Venter Institute | J. Craig Venter Institute | Shabman,R., Das,S.R., Shilts,M., Fedorova,N., Puri,V., Shrivastava,S., Amedeo,P., Williams,M., Barratt,K., Mitchell,J. and Jennings,L. |
| EPI_ISL_2584657 | Epidemiology and Demography, KEMRI-Wellcome Trust | Epidemiology and Demography, KEMRI-Wellcome Trust | Kamau,E., Otieno,J.R., Murunga,N., Nyiro,J.U., Oketch,J.W., Ngoi,J.M., de Laurent,Z.R., Mwema,A., Agoti,C.N. and Nokes,D.J. |
| EPI_ISL_2584658 | J. Craig Venter Institute | J. Craig Venter Institute | Shabman,R., Das,S.R., Shilts,M., Fedorova,N., Puri,V., Shrivastava,S., Amedeo,P., Williams,M., Barratt,K., Mitchell,J. and Jennings,L. |
| EPI_ISL_2584659, EPI_ISL_2584660 | J. Craig Venter Institute | J. Craig Venter Institute | Wentworth,D.E., Halpin,R.A., Bera,J., Lin,X., Fedorova,N., Tsitrin,T., McLellan,M., Stockwell,T., Amedeo,P., Bishop,B., Gupta,N., Hoover,J., Katzel,D., Schobel,S., Shrivastava,S., Garcia,J., Laguna-Torres,V.A., Leguia,M., Benavides,J.G. and Halsey,E. |
| EPI_ISL_2584661 | Epidemiology and Demography, KEMRI-Wellcome Trust | Epidemiology and Demography, KEMRI-Wellcome Trust | Kamau,E., Otieno,J.R., Murunga,N., Nyiro,J.U., Oketch,J.W., Ngoi,J.M., de Laurent,Z.R., Mwema,A., Agoti,C.N. and Nokes,D.J. |
| EPI_ISL_2584662 | J. Craig Venter Institute | J. Craig Venter Institute | Shabman,R., Das,S.R., Shilts,M., Fedorova,N., Puri,V., Shrivastava,S., Amedeo,P., Williams,M., Barratt,K., Mitchell,J. and Jennings,L. |
| EPI_ISL_2584664, EPI_ISL_2584665, EPI_ISL_2584666, EPI_ISL_2584667, EPI_ISL_2584668, EPI_ISL_2584669, EPI_ISL_2584670 | Epidemiology and Demography, KEMRI-Wellcome Trust | Epidemiology and Demography, KEMRI-Wellcome Trust | Kamau,E., Otieno,J.R., Murunga,N., Nyiro,J.U., Oketch,J.W., Ngoi,J.M., de Laurent,Z.R., Mwema,A., Agoti,C.N. and Nokes,D.J. |
| EPI_ISL_2584671 | J. Craig Venter Institute | J. Craig Venter Institute | Shabman,R., Das,S.R., Shilts,M., Fedorova,N., Puri,V., Shrivastava,S., Amedeo,P., Williams,M., Barratt,K., Mitchell,J. and Jennings,L. |
| EPI_ISL_2584673, EPI_ISL_2584674 | Epidemiology and Demography, KEMRI-Wellcome Trust | Epidemiology and Demography, KEMRI-Wellcome Trust | Kamau,E., Otieno,J.R., Murunga,N., Nyiro,J.U., Oketch,J.W., Ngoi,J.M., de Laurent,Z.R., Mwema,A., Agoti,C.N. and Nokes,D.J. |
| EPI_ISL_2584675 | Medical Microbiology, University Medical Center Utrecht | Medical Microbiology, University Medical Center Utrecht | Tan,L., Lemey,P., Viveen,M. and Coenjaerts,F. |
| EPI_ISL_2584676, EPI_ISL_2584677 | Epidemiology and Demography, KEMRI-Wellcome Trust | Epidemiology and Demography, KEMRI-Wellcome Trust | Kamau,E., Otieno,J.R., Murunga,N., Nyiro,J.U., Oketch,J.W., Ngoi,J.M., de Laurent,Z.R., Mwema,A., Agoti,C.N. and Nokes,D.J. |
| EPI_ISL_2584678 | J. Craig Venter Institute | J. Craig Venter Institute | Das,S.R., Halpin,R.A., Puri,V., Akopov,A., Fedorova,N., Stockwell,T., Amedeo,P., Bishop,B., Katzel,D., Schobel,S., Shrivastava,S., Wentworth,D.E. and Caserta,M. |
| EPI_ISL_2584680 | Epidemiology and Demography, KEMRI-Wellcome Trust | Epidemiology and Demography, KEMRI-Wellcome Trust | Kamau,E., Otieno,J.R., Murunga,N., Nyiro,J.U., Oketch,J.W., Ngoi,J.M., de Laurent,Z.R., Mwema,A., Agoti,C.N. and Nokes,D.J. |
| EPI_ISL_2584681 | J. Craig Venter Institute | J. Craig Venter Institute | Das,S.R., Halpin,R.A., Puri,V., Akopov,A., Fedorova,N., Stockwell,T., Amedeo,P., Bishop,B., Katzel,D., Schobel,S., Shrivastava,S., Hall,C.B., Tesini,B.L., Schnabel,K.C., Walsh,E.E. and Caserta,M. |
| EPI_ISL_2584682 | J. Craig Venter Institute | J. Craig Venter Institute | Das,S.R., Halpin,R.A., Puri,V., Akopov,A., Fedorova,N., Tsitrin,T., Stockwell,T., Amedeo,P., Bishop,B., Gupta,N., Hoover,J., Katzel,D., Schobel,S., Shrivastava,S., Wentworth,D.E. and Caserta,M. |
| EPI_ISL_2584683, EPI_ISL_2584684, EPI_ISL_2584685 | J. Craig Venter Institute | J. Craig Venter Institute | Das,S.R., Halpin,R.A., Puri,V., Akopov,A., Fedorova,N., Stockwell,T., Amedeo,P., Bishop,B., Katzel,D., Schobel,S., Shrivastava,S., Hall,C.B., Tesini,B.L., Schnabel,K.C., Walsh,E.E. and Caserta,M. |
| EPI_ISL_2584686 | J. Craig Venter Institute | J. Craig Venter Institute | Shabman,R., Das,S.R., Puri,V., Fedorova,N., Amedeo,P., Williams,M., Shrivastava,S. and Halasa,N. |
| EPI_ISL_2584688, EPI_ISL_2584690, EPI_ISL_2584691 | Epidemiology and Demography, KEMRI-Wellcome Trust | Epidemiology and Demography, KEMRI-Wellcome Trust | Kamau,E., Otieno,J.R., Murunga,N., Nyiro,J.U., Oketch,J.W., Ngoi,J.M., de Laurent,Z.R., Mwema,A., Agoti,C.N. and Nokes,D.J. |
| EPI_ISL_2584692 | Medicine, University of Washington, 300 9th Ave, Harborview Research & Training Building | Medicine, University of Washington, 300 9th Ave, Harborview Research & Training Building | Chu,H., Scott,E. and Roychoudhury,P. |
| EPI_ISL_2584693 | Epidemiology and Demography, KEMRI-Wellcome | Epidemiology and Demography, KEMRI-Wellcome | Kamau,E., Otieno,J.R., Murunga,N., Nyiro,J.U., Oketch,J.W., Ngoi,J.M., de Laurent,Z.R., Mwema,A., Agoti,C.N. and Nokes,D.J. |

| | | | |
|---|---|---|---|
| | Trust | Trust | |
| EPI_ISL_2584696 | J. Craig Venter Institute | J. Craig Venter Institute | Das,S.R., Halpin,R.A., Puri,V., Akopov,A., Fedorova,N., Tsitrin,T., Stockwell,T., Amedeo,P., Bishop,B., Gupta,N., Hoover,J., Katzel,D., Schobel,S., Shrivastava,S., Wentworth,D.E. and Caserta,M. |
| EPI_ISL_2584697 | J. Craig Venter Institute | J. Craig Venter Institute | Das,S.R., Halpin,R.A., Puri,V., Akopov,A., Fedorova,N., Stockwell,T., Amedeo,P., Bishop,B., Katzel,D., Schobel,S., Shrivastava,S., Wentworth,D.E. and Caserta,M. |
| EPI_ISL_2584698 | J. Craig Venter Institute | J. Craig Venter Institute | Das,S.R., Halpin,R.A., Puri,V., Akopov,A., Fedorova,N., Stockwell,T., Amedeo,P., Bishop,B., Katzel,D., Schobel,S., Shrivastava,S., Hall,C.B., Tesini,B.L., Schnabel,K.C., Walsh,E.E. and Caserta,M. |
| EPI_ISL_2584699 | Epidemiology and Demography, KEMRI-Wellcome Trust | Epidemiology and Demography, KEMRI-Wellcome Trust | Kamau,E., Otieno,J.R., Murunga,N., Nyiro,J.U., Oketch,J.W., Ngoi,J.M., de Laurent,Z.R., Mwema,A., Agoti,C.N. and Nokes,D.J. |
| EPI_ISL_2584703, EPI_ISL_2584705 | J. Craig Venter Institute | J. Craig Venter Institute | Das,S.R., Halpin,R.A., Puri,V., Akopov,A., Fedorova,N., Stockwell,T., Amedeo,P., Bishop,B., Katzel,D., Schobel,S., Shrivastava,S., Hall,C.B., Tesini,B.L., Schnabel,K.C., Walsh,E.E. and Caserta,M. |
| EPI_ISL_2584706, EPI_ISL_2584708, EPI_ISL_2584711, EPI_ISL_2584715, EPI_ISL_2584717, EPI_ISL_2584718, EPI_ISL_2584719, EPI_ISL_2584720, EPI_ISL_2584722, EPI_ISL_2584723, EPI_ISL_2584724, EPI_ISL_2584725 | | | |
| see above | Epidemiology and Demography, KEMRI-Wellcome Trust | Epidemiology and Demography, KEMRI-Wellcome Trust | Kamau,E., Otieno,J.R., Murunga,N., Nyiro,J.U., Oketch,J.W., Ngoi,J.M., de Laurent,Z.R., Mwema,A., Agoti,C.N. and Nokes,D.J. |
| EPI_ISL_2584726 | KEMRI Wellcome Trust Research Programme | KEMRI Wellcome Trust Research Programme | Agoti,C.N., Otieno,J.R., Munywoki,P.K., Mwihuri,A.G., Cane,P.A., Nokes,D.J., Kellam,P. and Cotten,M.L. |
| EPI_ISL_2584727 | J. Craig Venter Institute | J. Craig Venter Institute | Das,S., Halpin,R.A., Bera,J., Fedorova,N., Tsitrin,T., Stockwell,T., Amedeo,P., Bishop,B., Gupta,N., Hoover,J., Katzel,D., Schobel,S., Shrivastava,S., Hartert,T., Moore,M., Chappell,J., Larkin,E., Wentworth,D.E. and Anderson,L.J. |
| EPI_ISL_2584728 | J. Craig Venter Institute | J. Craig Venter Institute | Wentworth,D.E., Halpin,R.A., Bera,J., Lin,X., Fedorova,N., Tsitrin,T., McLellan,M., Stockwell,T., Amedeo,P., Bishop,B., Gupta,N., Hoover,J., Katzel,D., Schobel,S., Shrivastava,S., Garcia,J., Laguna-Torres,V.A., Leguia,M., Benavides,J.G. and Halsey,E. |
| EPI_ISL_2584729 | J. Craig Venter Institute | J. Craig Venter Institute | Das,S.R., Halpin,R.A., Shilts,M., Puri,V., Akopov,A., Fedorova,N., Stockwell,T., Amedeo,P., Bishop,B., Katzel,D., Schobel,S., Shrivastava,S. and Hartert,T. |
| EPI_ISL_2584730, EPI_ISL_2584731 | Medical Microbiology, University Medical Center Utrecht | Medical Microbiology, University Medical Center Utrecht | Tan,L., Viveen,M.C., Lemey,P. and Coenjaerts,F.E. |
| EPI_ISL_2584732 | J. Craig Venter Institute | J. Craig Venter Institute | Shabman,R., Das,S.R., Shilts,M., Fedorova,N., Puri,V., Shrivastava,S., Amedeo,P., Williams,M., Barratt,K., Mitchell,J. and Jennings,L. |
| EPI_ISL_2584733 | J. Craig Venter Institute | J. Craig Venter Institute | Shabman,R., Das,S.R., Shilts,M., Fedorova,N., Puri,V., Shrivastava,S., Amedeo,P., Hu,L., Durbin,A., Rocchi,I., Williams,T. and Hartert,T. |
| EPI_ISL_2584734 | J. Craig Venter Institute | J. Craig Venter Institute | Shabman,R., Das,S.R., Puri,V., Fedorova,N., Amedeo,P., Williams,M., Shrivastava,S. and Halasa,N. |
| EPI_ISL_2584735, EPI_ISL_2584736 | J. Craig Venter Institute | J. Craig Venter Institute | Das,S.R., Halpin,R.A., Shilts,M., Puri,V., Akopov,A., Fedorova,N., Stockwell,T., Amedeo,P., Bishop,B., Katzel,D., Schobel,S., Shrivastava,S. and Hartert,T. |
| EPI_ISL_2584737 | J. Craig Venter Institute | J. Craig Venter Institute | Shabman,R., Das,S.R., Shilts,M., Fedorova,N., Puri,V., Shrivastava,S., Amedeo,P., Williams,M., Barratt,K., Mitchell,J. and Jennings,L. |
| EPI_ISL_2584738 | Laboratory Medicine, UW Virology | Laboratory Medicine, UW Virology | Lin,M.J., Tait,A. and Greninger,A.L. |
| EPI_ISL_2584739 | J. Craig Venter Institute | J. Craig Venter Institute | Das,S.R., Halpin,R.A., Shilts,M., Puri,V., Akopov,A., Fedorova,N., Stockwell,T., Amedeo,P., Bishop,B., Katzel,D., Schobel,S., Shrivastava,S. and Hartert,T. |
| EPI_ISL_2584740 | Medical Microbiology, University Medical Center Utrecht | Medical Microbiology, University Medical Center Utrecht | Tan,L., Viveen,M.C., Lemey,P. and Coenjaerts,F.E. |
| EPI_ISL_2584741 | J. Craig Venter Institute | J. Craig Venter Institute | Lorenzi,H., Town,C., Halpin,R., Bera,J., Ransier,A., Fedorova,N., Stockwell,T., Amedeo,P., Appalla,L., Bishop,B., Edworthy,P., Gupta,N., Hoover,J., Katzel,D., Li,K., Schobel,S., Shrivastava,S., Thovarai,V., Wang,S., Rebuffo-Scheer,C., Fan,J., He,J., Kehl,S.C., Lederboer,N., Jurgens,L.A., Bose,M.E., Beck,E.T., Kumar,S., Wentworth,D.E. and Henrickson,K.J. |
| EPI_ISL_2584742 | J. Craig Venter Institute | J. Craig Venter Institute | Shabman,R., Das,S.R., Shilts,M., Fedorova,N., Puri,V., Shrivastava,S., Amedeo,P., Williams,M., Barratt,K., Mitchell,J. and Jennings,L. |
| EPI_ISL_2584745 | J. Craig Venter Institute | J. Craig Venter Institute | Das,S.R., Halpin,R.A., Puri,V., Akopov,A., Fedorova,N., Stockwell,T., Amedeo,P., Bishop,B., Katzel,D., Schobel,S., Shrivastava,S., Wentworth,D.E. and Caserta,M. |
| EPI_ISL_2584748, EPI_ISL_2584749 | J. Craig Venter Institute | J. Craig Venter Institute | Das,S.R., Halpin,R.A., Puri,V., Akopov,A., Fedorova,N., Stockwell,T., Amedeo,P., Bishop,B., Katzel,D., Schobel,S., Shrivastava,S., Hall,C.B., Tesini,B.L., Schnabel,K.C., Walsh,E.E. and Caserta,M. |
| EPI_ISL_2584750, EPI_ISL_2584751 | J. Craig Venter Institute | J. Craig Venter Institute | Das,S.R., Halpin,R.A., Puri,V., Akopov,A., Fedorova,N., Stockwell,T., Amedeo,P., Bishop,B., Katzel,D., Schobel,S., Shrivastava,S., Wentworth,D.E. and Caserta,M. |
| EPI_ISL_2584752 | J. Craig Venter Institute | J. Craig Venter Institute | Das,S.R., Halpin,R.A., Puri,V., Akopov,A., Fedorova,N., Stockwell,T., Amedeo,P., Bishop,B., Katzel,D., Schobel,S., Shrivastava,S., Hall,C.B., Tesini,B.L., Schnabel,K.C., Walsh,E.E. and Caserta,M. |
| EPI_ISL_2584753 | J. Craig Venter Institute | J. Craig Venter Institute | Das,S.R., Halpin,R.A., Puri,V., Akopov,A., Fedorova,N., Tsitrin,T., Stockwell,T., Amedeo,P., Bishop,B., Gupta,N., Hoover,J., Katzel,D., Schobel,S., Shrivastava,S., Wentworth,D.E. and Caserta,M. |
| EPI_ISL_2584754, EPI_ISL_2584755, EPI_ISL_2584756 | J. Craig Venter Institute | J. Craig Venter Institute | Das,S.R., Halpin,R.A., Puri,V., Akopov,A., Fedorova,N., Stockwell,T., Amedeo,P., Bishop,B., Katzel,D., Schobel,S., Shrivastava,S., Wentworth,D.E. and Caserta,M. |
| EPI_ISL_2584760 | J. Craig Venter Institute | J. Craig Venter Institute | Das,S.R., Halpin,R.A., Puri,V., Akopov,A., Fedorova,N., Tsitrin,T., Stockwell,T., Amedeo,P., Bishop,B., Gupta,N., Hoover,J., Katzel,D., Schobel,S., Shrivastava,S., Wentworth,D.E. and Caserta,M. |
| EPI_ISL_2584762 | J. Craig Venter Institute | J. Craig Venter Institute | Das,S.R., Halpin,R.A., Puri,V., Akopov,A., Fedorova,N., Stockwell,T., Amedeo,P., Bishop,B., Katzel,D., Schobel,S., Shrivastava,S., Hall,C.B., Tesini,B.L., Schnabel,K.C., Walsh,E.E. and Caserta,M. |
| EPI_ISL_2584764 | J. Craig Venter Institute | J. Craig Venter Institute | Das,S.R., Halpin,R.A., Puri,V., Akopov,A., Fedorova,N., Stockwell,T., Amedeo,P., Bishop,B., Katzel,D., Schobel,S., Shrivastava,S., Wentworth,D.E. and Caserta,M. |
| EPI_ISL_2584765 | J. Craig Venter Institute | J. Craig Venter Institute | Das,S.R., Halpin,R.A., Puri,V., Akopov,A., Fedorova,N., Stockwell,T., Amedeo,P., Bishop,B., Katzel,D., Schobel,S., Shrivastava,S., Hall,C.B., Tesini,B.L., Schnabel,K.C., Walsh,E.E. and Caserta,M. |
| EPI_ISL_2584766 | J. Craig Venter Institute | J. Craig Venter Institute | Das,S.R., Halpin,R.A., Puri,V., Akopov,A., Fedorova,N., Stockwell,T., Amedeo,P., Bishop,B., Katzel,D., Schobel,S., Shrivastava,S., Wentworth,D.E. and Caserta,M. |
| EPI_ISL_2584768, EPI_ISL_2584770, EPI_ISL_2584771 | J. Craig Venter Institute | J. Craig Venter Institute | Das,S.R., Halpin,R.A., Puri,V., Akopov,A., Fedorova,N., Stockwell,T., Amedeo,P., Bishop,B., Katzel,D., Schobel,S., Shrivastava,S., Hall,C.B., Tesini,B.L., Schnabel,K.C., Walsh,E.E. and Caserta,M. |
| EPI_ISL_2584773, EPI_ISL_2584774 | Epidemiology and Demography, KEMRI-Wellcome Trust | Epidemiology and Demography, KEMRI-Wellcome Trust | Kamau,E., Otieno,J.R., Murunga,N., Nyiro,J.U., Oketch,J.W., Ngoi,J.M., de Laurent,Z.R., Mwema,A., Agoti,C.N. and Nokes,D.J. |
| EPI_ISL_2584812 | J. Craig Venter Institute | J. Craig Venter Institute | Shabman,R., Das,S.R., Puri,V., Fedorova,N., Amedeo,P., Williams,M., Shrivastava,S. and Halasa,N. |
| EPI_ISL_2584814 | J. Craig Venter Institute | J. Craig Venter Institute | Das,S.R., Halpin,R.A., Puri,V., Akopov,A., Fedorova,N., Tsitrin,T., Stockwell,T., Amedeo,P., Bishop,B., Gupta,N., Hoover,J., Katzel,D., Schobel,S., Shrivastava,S., Wentworth,D.E. and Caserta,M. |
| EPI_ISL_2584815 | J. Craig Venter Institute | J. Craig Venter Institute | Das,S., Halpin,R.A., Bera,J., Fedorova,N., Tsitrin,T., Stockwell,T., Amedeo,P., Bishop,B., Gupta,N., Hoover,J., Katzel,D., Schobel,S., Shrivastava,S., Hartert,T., Moore,M., Chappell,J., Larkin,E., Wentworth,D.E. and Anderson,L.J. |
| EPI_ISL_2584817 | J. Craig Venter Institute | J. Craig Venter Institute | Das,S.R., Halpin,R.A., Puri,V., Akopov,A., Fedorova,N., Stockwell,T., Amedeo,P., Bishop,B., Katzel,D., Schobel,S., Shrivastava,S., Wentworth,D.E. and Caserta,M. |
| EPI_ISL_2584818 | J. Craig Venter Institute | J. Craig Venter Institute | Das,S.R., Halpin,R.A., Puri,V., Akopov,A., Fedorova,N., Tsitrin,T., Stockwell,T., Amedeo,P., Bishop,B., Gupta,N., Hoover,J., Katzel,D., Schobel,S., Shrivastava,S., Wentworth,D.E. and Caserta,M. |
| EPI_ISL_2584819, EPI_ISL_2584820 | J. Craig Venter Institute | J. Craig Venter Institute | Das,S.R., Halpin,R.A., Puri,V., Akopov,A., Fedorova,N., Stockwell,T., Amedeo,P., Bishop,B., Katzel,D., Schobel,S., Shrivastava,S., Hall,C.B., Tesini,B.L., Schnabel,K.C., Walsh,E.E. and Caserta,M. |
| EPI_ISL_2584824, EPI_ISL_2584847 | KEMRI Wellcome Trust Research Programme | KEMRI Wellcome Trust Research Programme | Agoti,C.N., Otieno,J.R., Munywoki,P.K., Mwihuri,A.G., Cane,P.A., Nokes,D.J., Kellam,P. and Cotten,M.L. |
| EPI_ISL_2584849 | J. Craig Venter Institute | J. Craig Venter Institute | Wentworth,D.E., Halpin,R.A., Bera,J., Lin,X., Fedorova,N., Tsitrin,T., McLellan,M., Stockwell,T., Amedeo,P., Bishop,B., Gupta,N., Hoover,J., Katzel,D., Schobel,S., Shrivastava,S., Garcia,J., Laguna-Torres,V.A., Leguia,M., Benavides,J.G. and Halsey,E. |
| EPI_ISL_2584850 | J. Craig Venter Institute | J. Craig Venter Institute | Das,S.R., Halpin,R.A., Puri,V., Akopov,A., Fedorova,N., Tsitrin,T., Stockwell,T., Amedeo,P., Bishop,B., Gupta,N., Hoover,J., Katzel,D., Schobel,S., Shrivastava,S., Wentworth,D.E. and Caserta,M. |
| EPI_ISL_2584852 | Medical Microbiology, University Medical Center Utrecht | Medical Microbiology, University Medical Center Utrecht | Tan,L., Viveen,M.C., Lemey,P. and Coenjaerts,F.E. |
| EPI_ISL_2584853 | J. Craig Venter Institute | J. Craig Venter Institute | Shabman,R., Das,S.R., Shilts,M., Fedorova,N., Puri,V., Shrivastava,S., Amedeo,P., Williams,M., Barratt,K., Mitchell,J. and Jennings,L. |
| EPI_ISL_2584855 | J. Craig Venter Institute | J. Craig Venter Institute | Das,S., Halpin,R.A., Bera,J., Puri,V., Fedorova,N., Tsitrin,T., Stockwell,T., Amedeo,P., Bishop,B., Katzel,D., Schobel,S., Shrivastava,S., Hartert,T., Moore,M., Chappell,J., Larkin,E., Wentworth,D.E. and Anderson,L.J. |
| EPI_ISL_2584856 | Epidemiology and Demography, KEMRI-Wellcome Trust | Epidemiology and Demography, KEMRI-Wellcome Trust | Kamau,E., Otieno,J.R., Murunga,N., Nyiro,J.U., Oketch,J.W., Ngoi,J.M., de Laurent,Z.R., Mwema,A., Agoti,C.N. and Nokes,D.J. |
| EPI_ISL_2584859 | KEMRI Wellcome Trust Research Programme | KEMRI Wellcome Trust Research Programme | Agoti,C.N., Otieno,J.R., Munywoki,P.K., Mwihuri,A.G., Cane,P.A., Nokes,D.J., Kellam,P. and Cotten,M.L. |
| EPI_ISL_2584860 | J. Craig Venter Institute | J. Craig Venter Institute | Shabman,R., Das,S.R., Shilts,M., Fedorova,N., Puri,V., Shrivastava,S., Amedeo,P., Williams,M., Barratt,K., Mitchell,J. and Jennings,L. |
| EPI_ISL_2584861 | J. Craig Venter Institute | J. Craig Venter Institute | Wentworth,D.E., Halpin,R.A., Bera,J., Lin,X., Fedorova,N., Tsitrin,T., McLellan,M., Stockwell,T., Amedeo,P., Bishop,B., Gupta,N., Hoover,J., Katzel,D., Schobel,S., Shrivastava,S., Garcia,J., Laguna-Torres,V.A., Leguia,M., Benavides,J.G. and Halsey,E. |
| EPI_ISL_2584862 | Michiko Okamoto Tohoku University Graduate School of Medicine, Virology | Michiko Okamoto Tohoku University Graduate School of Medicine, Virology | Okamoto,M., Malasaor,R. and Oshitani,H. |
| EPI_ISL_2584863 | J. Craig Venter Institute | J. Craig Venter Institute | Das,S.R., Halpin,R.A., Shilts,M., Puri,V., Akopov,A., Fedorova,N., Stockwell,T., Amedeo,P., Bishop,B., Katzel,D., Schobel,S., Shrivastava,S. and Hartert,T. |
| EPI_ISL_2584865 | J. Craig Venter Institute | J. Craig Venter Institute | Shabman,R., Das,S.R., Puri,V., Fedorova,N., Amedeo,P., Williams,M., Shrivastava,S. and Halasa,N. |
| EPI_ISL_2584866, EPI_ISL_2584867, EPI_ISL_2584868, EPI_ISL_2584869, EPI_ISL_2584871, EPI_ISL_2584872 | Epidemiology and Demography, KEMRI-Wellcome Trust | Epidemiology and Demography, KEMRI-Wellcome Trust | Kamau,E., Otieno,J.R., Murunga,N., Nyiro,J.U., Oketch,J.W., Ngoi,J.M., de Laurent,Z.R., Mwema,A., Agoti,C.N. and Nokes,D.J. |
| EPI_ISL_2584873, EPI_ISL_2584874, EPI_ISL_2584875 | J. Craig Venter Institute | J. Craig Venter Institute | Wentworth,D.E., Halpin,R.A., Bera,J., Lin,X., Fedorova,N., Tsitrin,T., McLellan,M., Stockwell,T., Amedeo,P., Bishop,B., Gupta,N., Hoover,J., Katzel,D., Schobel,S., Shrivastava,S., Garcia,J., Laguna-Torres,V.A., Leguia,M., Benavides,J.G. and Halsey,E. |
| EPI_ISL_2584879 | J. Craig Venter Institute | J. Craig Venter Institute | Das,S., Halpin,R.A., Bera,J., Puri,V., Fedorova,N., Tsitrin,T., Stockwell,T., Amedeo,P., Bishop,B., Katzel,D., Schobel,S., Shrivastava,S., Hartert,T., Moore,M., Chappell,J., Larkin,E., Wentworth,D.E. and Anderson,L.J. |
| EPI_ISL_2584880 | J. Craig Venter Institute | J. Craig Venter Institute | Das,S.R., Halpin,R.A., Shilts,M., Puri,V., Akopov,A., Fedorova,N., Stockwell,T., Amedeo,P., Bishop,B., Katzel,D., Schobel,S., Shrivastava,S. and Hartert,T. |
| EPI_ISL_2584881, EPI_ISL_2584882, EPI_ISL_2584883, EPI_ISL_2584884, EPI_ISL_2584885, EPI_ISL_2584886, EPI_ISL_2584887, EPI_ISL_2584888, EPI_ISL_2584889 | Epidemiology and Demography, KEMRI-Wellcome Trust | Epidemiology and Demography, KEMRI-Wellcome Trust | Kamau,E., Otieno,J.R., Murunga,N., Nyiro,J.U., Oketch,J.W., Ngoi,J.M., de Laurent,Z.R., Mwema,A., Agoti,C.N. and Nokes,D.J. |
| EPI_ISL_2584890 | Medical Microbiology, University Medical Center Utrecht | Medical Microbiology, University Medical Center Utrecht | Tan,L., Lemey,P., Viveen,M. and Coenjaerts,F.E.J. |
| EPI_ISL_2584891, EPI_ISL_2584893 | J. Craig Venter Institute | J. Craig Venter Institute | Shabman,R., Das,S.R., Shilts,M., Fedorova,N., Puri,V., Shrivastava,S., Amedeo,P., Williams,M., Barratt,K., Mitchell,J. and Jennings,L. |
| EPI_ISL_2584894 | J. Craig Venter Institute | J. Craig Venter Institute | Wentworth,D.E., Halpin,R.A., Bera,J., Lin,X., Fedorova,N., Tsitrin,T., McLellan,M., Stockwell,T., Amedeo,P., Bishop,B., Gupta,N., Hoover,J., Katzel,D., Schobel,S., Shrivastava,S., Garcia,J., Laguna-Torres,V.A., Leguia,M., Benavides,J.G. and Halsey,E. |
| EPI_ISL_2584896 | Medicine, University of Washington, 300 9th Ave, | Medicine, University of Washington, 300 9th Ave, | Chu,H., Scott,E. and Roychoudhury,P. |

| | | | |
|---|---|---|---|
| | Harborview Research & Training Building | Harborview Research & Training Building | |
| EPI_ISL_2584897 | J. Craig Venter Institute | J. Craig Venter Institute | Das,S., Halpin,R.A., Bera,J., Fedorova,N., Tsitrin,T., Stockwell,T., Amedeo,P., Bishop,B., Gupta,N., Hoover,J., Katzel,D., Schobel,S., Shrivastava,S., Hartert,T., Moore,M., Chappell,J., Larkin,E., Wentworth,D.E. and Anderson,L.J. |
| EPI_ISL_2584898, EPI_ISL_2584899, EPI_ISL_2584901, EPI_ISL_2584902, EPI_ISL_2584904, EPI_ISL_2584905, EPI_ISL_2584906, EPI_ISL_2584907, EPI_ISL_2584909 | J. Craig Venter Institute | J. Craig Venter Institute | Das,S.R., Halpin,R.A., Shilts,M., Puri,V., Akopov,A., Fedorova,N., Stockwell,T., Amedeo,P., Bishop,B., Katzel,D., Schobel,S., Shrivastava,S. and Hartert,T. |
| EPI_ISL_2584912, EPI_ISL_2584913, EPI_ISL_2584914, EPI_ISL_2584915, EPI_ISL_2584916, EPI_ISL_2584917, EPI_ISL_2584918, EPI_ISL_2584919, EPI_ISL_2584920, EPI_ISL_2584921, EPI_ISL_2584922, EPI_ISL_2584923, EPI_ISL_2584924, EPI_ISL_2584925, EPI_ISL_2584926, EPI_ISL_2584927, EPI_ISL_2584928, EPI_ISL_2584929, EPI_ISL_2584930, EPI_ISL_2584931, EPI_ISL_2584932, EPI_ISL_2584933, EPI_ISL_2584934, EPI_ISL_2584935 | | | |
| see above | Epidemiology and Demography, KEMRI-Wellcome Trust | Epidemiology and Demography, KEMRI-Wellcome Trust | Kamau,E., Otieno,J.R., Murunga,N., Nyiro,J.U., Oketch,J.W., Ngoi,J.M., de Laurent,Z.R., Mwema,A., Agoti,C.N. and Nokes,D.J. |
| EPI_ISL_2584937 | Medical Microbiology, University Medical Center Utrecht | Medical Microbiology, University Medical Center Utrecht | Tan,L., Viveen,M.C., Lemey,P. and Coenjaerts,F.E. |
| EPI_ISL_2584938, EPI_ISL_2584939, EPI_ISL_2584940 | J. Craig Venter Institute | J. Craig Venter Institute | Shabman,R., Das,S.R., Shilts,M., Fedorova,N., Puri,V., Shrivastava,S., Amedeo,P., Williams,M., Barratt,K., Mitchell,J. and Jennings,L. |
| EPI_ISL_2584941 | Medicine, University of Washington, 300 9th Ave, Harborview Research & Training Building | Medicine, University of Washington, 300 9th Ave, Harborview Research & Training Building | Chu,H., Scott,E. and Roychoudhury,P. |
| EPI_ISL_2584944, EPI_ISL_2584945, EPI_ISL_2584946, EPI_ISL_2584947, EPI_ISL_2584948, EPI_ISL_2584950, EPI_ISL_2584952, EPI_ISL_2584953 | J. Craig Venter Institute | J. Craig Venter Institute | Das,S.R., Halpin,R.A., Shilts,M., Puri,V., Akopov,A., Fedorova,N., Stockwell,T., Amedeo,P., Bishop,B., Katzel,D., Schobel,S., Shrivastava,S. and Hartert,T. |
| EPI_ISL_2584955, EPI_ISL_2584956, EPI_ISL_2584958, EPI_ISL_2584959, EPI_ISL_2584960, EPI_ISL_2584961, EPI_ISL_2584963, EPI_ISL_2584964, EPI_ISL_2584965, EPI_ISL_2584967, EPI_ISL_2584968, EPI_ISL_2584969, EPI_ISL_2584970, EPI_ISL_2584972, EPI_ISL_2584973, EPI_ISL_2584975, EPI_ISL_2584976, EPI_ISL_2584977, EPI_ISL_2584978, EPI_ISL_2584980, EPI_ISL_2584981, EPI_ISL_2584982, EPI_ISL_2584983, EPI_ISL_2584985 | | | |
| see above | Epidemiology and Demography, KEMRI-Wellcome Trust | Epidemiology and Demography, KEMRI-Wellcome Trust | Kamau,E., Otieno,J.R., Murunga,N., Nyiro,J.U., Oketch,J.W., Ngoi,J.M., de Laurent,Z.R., Mwema,A., Agoti,C.N. and Nokes,D.J. |
| EPI_ISL_2584986, EPI_ISL_2584987 | Medical Microbiology, University Medical Center Utrecht | Medical Microbiology, University Medical Center Utrecht | Tan,L., Viveen,M.C., Lemey,P. and Coenjaerts,F.E. |
| EPI_ISL_2584988, EPI_ISL_2584989, EPI_ISL_2584990, EPI_ISL_2584991 | J. Craig Venter Institute | J. Craig Venter Institute | Shabman,R., Das,S.R., Shilts,M., Fedorova,N., Puri,V., Shrivastava,S., Amedeo,P., Williams,M., Barratt,K., Mitchell,J. and Jennings,L. |
| EPI_ISL_2584992, EPI_ISL_2584993 | J. Craig Venter Institute | J. Craig Venter Institute | Wentworth,D.E., Halpin,R.A., Bera,J., Lin,X., Fedorova,N., Tsitrin,T., McLellan,M., Stockwell,T., Amedeo,P., Bishop,B., Gupta,N., Hoover,J., Katzel,D., Schobel,S., Shrivastava,S., Garcia,J., Laguna-Torres,V.A., Leguia,M., Benavides,J.G. and Halsey,E. |
| EPI_ISL_2584994 | Division of Biosafety Evaluation and Control, Korea National Institute of Health | Division of Biosafety Evaluation and Control, Korea National Institute of Health | Yun,M.-R., Lee,W.-J., Kim,A.-R., Lee,H.S., Kim,K., Kim,S.S., Kim,Y.-J. and Kim,D.-W. |
| EPI_ISL_2584995, EPI_ISL_2584996, EPI_ISL_2584997, EPI_ISL_2584998, EPI_ISL_2584999, EPI_ISL_2585000, EPI_ISL_2585001, EPI_ISL_2585002, EPI_ISL_2585003, EPI_ISL_2585004, EPI_ISL_2585005, EPI_ISL_2585006, EPI_ISL_2585007, EPI_ISL_2585009, EPI_ISL_2585010, EPI_ISL_2585012, EPI_ISL_2585013, EPI_ISL_2585014, EPI_ISL_2585015 | | | |
| see above | J. Craig Venter Institute | J. Craig Venter Institute | Das,S.R., Halpin,R.A., Shilts,M., Puri,V., Akopov,A., Fedorova,N., Stockwell,T., Amedeo,P., Bishop,B., Katzel,D., Schobel,S., Shrivastava,S. and Hartert,T. |
| EPI_ISL_2585016 | J. Craig Venter Institute | J. Craig Venter Institute | Shabman,R., Das,S.R., Puri,V., Fedorova,N., Amedeo,P., Williams,M., Shrivastava,S. and Halasa,N. |
| EPI_ISL_2585017, EPI_ISL_2585018, EPI_ISL_2585019, EPI_ISL_2585020, EPI_ISL_2585021, EPI_ISL_2585022, EPI_ISL_2585023, EPI_ISL_2585025, EPI_ISL_2585026, EPI_ISL_2585027, EPI_ISL_2585028, EPI_ISL_2585029, EPI_ISL_2585031, EPI_ISL_2585032, EPI_ISL_2585033, EPI_ISL_2585036, EPI_ISL_2585037, EPI_ISL_2585039, EPI_ISL_2585040, EPI_ISL_2585041, EPI_ISL_2585042, EPI_ISL_2585043, EPI_ISL_2585044, EPI_ISL_2585045, EPI_ISL_2585047, EPI_ISL_2585049, EPI_ISL_2585050, EPI_ISL_2585051, EPI_ISL_2585052, EPI_ISL_2585053, EPI_ISL_2585054, EPI_ISL_2585056, EPI_ISL_2585057, EPI_ISL_2585058, EPI_ISL_2585059, EPI_ISL_2585060, EPI_ISL_2585061 | | | |
| see above | Epidemiology and Demography, KEMRI-Wellcome Trust | Epidemiology and Demography, KEMRI-Wellcome Trust | Kamau,E., Otieno,J.R., Murunga,N., Nyiro,J.U., Oketch,J.W., Ngoi,J.M., de Laurent,Z.R., Mwema,A., Agoti,C.N. and Nokes,D.J. |
| EPI_ISL_2585063 | Medical Microbiology, University Medical Center Utrecht | Medical Microbiology, University Medical Center Utrecht | Tan,L., Viveen,M.C., Lemey,P. and Coenjaerts,F.E. |
| EPI_ISL_2585064, EPI_ISL_2585065, EPI_ISL_2585066, EPI_ISL_2585067, EPI_ISL_2585068, EPI_ISL_2585070, EPI_ISL_2585072 | J. Craig Venter Institute | J. Craig Venter Institute | Shabman,R., Das,S.R., Shilts,M., Fedorova,N., Puri,V., Shrivastava,S., Amedeo,P., Williams,M., Barratt,K., Mitchell,J. and Jennings,L. |
| EPI_ISL_2585074 | J. Craig Venter Institute | J. Craig Venter Institute | Lorenzi,H., Town,C., Halpin,R., Bera,J., Ransier,A., Fedorova,N., Stockwell,T., Amedeo,P., Appalla,L., Bishop,B., Edworthy,P., Gupta,N., Hoover,J., Katzel,D., Li,K., Schobel,S., Shrivastava,S., Thovarai,V., Wang,S., Rebuffo-Scheer,C., Fan,J., He,J., Kehl,S.C., Lederboer,N., Jurgens,L.A., Bose,M.E., Beck,E.T., Kumar,S., Wentworth,D.E. and Henrickson,K.J. |
| EPI_ISL_2585075 | J. Craig Venter Institute | J. Craig Venter Institute | Das,S.R., Halpin,R.A., Shilts,M., Puri,V., Akopov,A., Fedorova,N., Stockwell,T., Amedeo,P., Bishop,B., Katzel,D., Schobel,S., Shrivastava,S. and Hartert,T. |
| EPI_ISL_2585077 | J. Craig Venter Institute | J. Craig Venter Institute | Shabman,R., Das,S.R., Shilts,M., Fedorova,N., Puri,V., Shrivastava,S., Amedeo,P., Hu,L., Durbin,A., Rocchi,I., Williams,T. and Hartert,T. |
| EPI_ISL_2585078, EPI_ISL_2585079, EPI_ISL_2585080, EPI_ISL_2585081, EPI_ISL_2585082, EPI_ISL_2585083, EPI_ISL_2585084, EPI_ISL_2585085, EPI_ISL_2585086, EPI_ISL_2585087, EPI_ISL_2585091, EPI_ISL_2585093, EPI_ISL_2585094 | | | |
| see above | J. Craig Venter Institute | J. Craig Venter Institute | Das,S.R., Halpin,R.A., Shilts,M., Puri,V., Akopov,A., Fedorova,N., Stockwell,T., Amedeo,P., Bishop,B., Katzel,D., Schobel,S., Shrivastava,S. and Hartert,T. |
| EPI_ISL_2585095 | J. Craig Venter Institute | J. Craig Venter Institute | Shabman,R., Das,S.R., Puri,V., Fedorova,N., Amedeo,P., Williams,M., Shrivastava,S. and Halasa,N. |
| EPI_ISL_2585096, EPI_ISL_2585098, EPI_ISL_2585099, EPI_ISL_2585100, EPI_ISL_2585101, EPI_ISL_2585102, EPI_ISL_2585103, EPI_ISL_2585104, EPI_ISL_2585105, EPI_ISL_2585106, EPI_ISL_2585107, EPI_ISL_2585108, EPI_ISL_2585110, EPI_ISL_2585111, EPI_ISL_2585112, EPI_ISL_2585113, EPI_ISL_2585114, EPI_ISL_2585115, EPI_ISL_2585116, EPI_ISL_2585117, EPI_ISL_2585118, EPI_ISL_2585119, EPI_ISL_2585120, EPI_ISL_2585121, EPI_ISL_2585123, EPI_ISL_2585124, EPI_ISL_2585125, EPI_ISL_2585126, EPI_ISL_2585127, EPI_ISL_2585128, EPI_ISL_2585130, EPI_ISL_2585131, EPI_ISL_2585132, EPI_ISL_2585133, EPI_ISL_2585134, EPI_ISL_2585135, EPI_ISL_2585136, EPI_ISL_2585137, EPI_ISL_2585139, EPI_ISL_2585140 | | | |
| see above | Epidemiology and Demography, KEMRI-Wellcome Trust | Epidemiology and Demography, KEMRI-Wellcome Trust | Kamau,E., Otieno,J.R., Murunga,N., Nyiro,J.U., Oketch,J.W., Ngoi,J.M., de Laurent,Z.R., Mwema,A., Agoti,C.N. and Nokes,D.J. |
| EPI_ISL_2585142 | Medical Microbiology, University Medical Center Utrecht | Medical Microbiology, University Medical Center Utrecht | Tan,L., Viveen,M.C., Lemey,P. and Coenjaerts,F.E. |
| EPI_ISL_2585143 | J. Craig Venter Institute | J. Craig Venter Institute | Shabman,R., Das,S.R., Shilts,M., Fedorova,N., Puri,V., Shrivastava,S., Amedeo,P., Williams,M., Barratt,K., Mitchell,J. and Jennings,L. |
| EPI_ISL_2585144 | J. Craig Venter Institute | J. Craig Venter Institute | Wentworth,D.E., Halpin,R.A., Bera,J., Lin,X., Fedorova,N., Tsitrin,T., McLellan,M., Stockwell,T., Amedeo,P., Bishop,B., Gupta,N., Hoover,J., Katzel,D., Schobel,S., Shrivastava,S., Garcia,J., Laguna-Torres,V.A., Leguia,M., Benavides,J.G. and Halsey,E. |
| EPI_ISL_2585146 | Taylor G.S., Virology laboratory, Bristol Public Health Laboratory | Taylor G.S., Virology laboratory, Bristol Public Health Laboratory | Taylor,G.S. |
| EPI_ISL_2585147, EPI_ISL_2585148 | Medicine, University of Washington, 300 9th Ave, Harborview Research & Training Building | Medicine, University of Washington, 300 9th Ave, Harborview Research & Training Building | Chu,H., Scott,E. and Roychoudhury,P. |
| EPI_ISL_2585149, EPI_ISL_2585150 | J. Craig Venter Institute | J. Craig Venter Institute | Das,S.R., Halpin,R.A., Shilts,M., Puri,V., Akopov,A., Fedorova,N., Stockwell,T., Amedeo,P., Bishop,B., Katzel,D., Schobel,S., Shrivastava,S. and Hartert,T. |
| EPI_ISL_2585153, EPI_ISL_2585154, EPI_ISL_2585155 | J. Craig Venter Institute | J. Craig Venter Institute | Shabman,R., Das,S.R., Puri,V., Fedorova,N., Amedeo,P., Williams,M., Shrivastava,S. and Halasa,N. |
| EPI_ISL_2585156, EPI_ISL_2585157, EPI_ISL_2585158, EPI_ISL_2585160, EPI_ISL_2585161, EPI_ISL_2585162, EPI_ISL_2585163, EPI_ISL_2585164, EPI_ISL_2585165, EPI_ISL_2585167, EPI_ISL_2585168, EPI_ISL_2585169, EPI_ISL_2585171, EPI_ISL_2585172, EPI_ISL_2585173, EPI_ISL_2585174, EPI_ISL_2585175, EPI_ISL_2585176, EPI_ISL_2585177, EPI_ISL_2585178, EPI_ISL_2585179, EPI_ISL_2585180, EPI_ISL_2585181, EPI_ISL_2585182, EPI_ISL_2585183, EPI_ISL_2585184, EPI_ISL_2585185, EPI_ISL_2585186, EPI_ISL_2585187, EPI_ISL_2585188 | | | |
| see above | Epidemiology and Demography, KEMRI-Wellcome Trust | Epidemiology and Demography, KEMRI-Wellcome Trust | Kamau,E., Otieno,J.R., Murunga,N., Nyiro,J.U., Oketch,J.W., Ngoi,J.M., de Laurent,Z.R., Mwema,A., Agoti,C.N. and Nokes,D.J. |
| EPI_ISL_2585189, EPI_ISL_2585190 | Medical Microbiology, University Medical Center Utrecht | Medical Microbiology, University Medical Center Utrecht | Tan,L., Viveen,M.C., Lemey,P. and Coenjaerts,F.E. |
| EPI_ISL_2585191, EPI_ISL_2585192, EPI_ISL_2585193, EPI_ISL_2585195 | J. Craig Venter Institute | J. Craig Venter Institute | Shabman,R., Das,S.R., Shilts,M., Fedorova,N., Puri,V., Shrivastava,S., Amedeo,P., Williams,M., Barratt,K., Mitchell,J. and Jennings,L. |
| EPI_ISL_2585196 | Medicine, University of Washington, 300 9th Ave, Harborview Research & Training Building | Medicine, University of Washington, 300 9th Ave, Harborview Research & Training Building | Chu,H., Scott,E. and Roychoudhury,P. |
| EPI_ISL_2585199, EPI_ISL_2585200 | J. Craig Venter Institute | J. Craig Venter Institute | Das,S.R., Halpin,R.A., Shilts,M., Puri,V., Akopov,A., Fedorova,N., Stockwell,T., Amedeo,P., Bishop,B., Katzel,D., Schobel,S., Shrivastava,S. and Hartert,T. |
| EPI_ISL_2588651 | Virology Laboratory, Dr. Ricardo Gutierrez Children Hospital | Virology Laboratory, Dr. Ricardo Gutierrez Children Hospital | Goya,S., Rojo,G.L., Valinotto,L.E., Mistchenko,A.S. and Viegas,M. |
| EPI_ISL_2588654 | Virology Laboratory, Dr. Ricardo Gutierrez Children Hospital | Virology Laboratory, Dr. Ricardo Gutierrez Children Hospital | Goya,S., Valinotto,L.E., Tittarelli,E., Rojo,G.L., Greninger,A., Luso,S., Natale,M., Mistchenko,A.S. and Viegas,M. |
| EPI_ISL_2588655 | Virology Laboratory, Dr. Ricardo Gutierrez Children Hospital | Virology Laboratory, Dr. Ricardo Gutierrez Children Hospital | Goya,S., Rojo,G.L., Valinotto,L.E., Mistchenko,A.S. and Viegas,M. |
| EPI_ISL_2588656, EPI_ISL_2588657, EPI_ISL_2588658 | J. Craig Venter Institute | J. Craig Venter Institute | Lorenzi,H., Town,C., Halpin,R., Bera,J., Ransier,A., Fedorova,N., Stockwell,T., Amedeo,P., Appalla,L., Bishop,B., Edworthy,P., Gupta,N., Hoover,J., Katzel,D., Li,K., Schobel,S., Shrivastava,S., Thovarai,V., Wang,S., Rebuffo-Scheer,C., Fan,J., He,J., Kehl,S.C., Lederboer,N., Jurgens,L.A., Bose,M.E., Beck,E.T., Kumar,S., Gerna,G., Wentworth,D.E. and Henrickson,K.J. |
| EPI_ISL_2588659 | Key Laboratory of Emergency Detection for Public Health of Zhejiang Province, Zhejiang Provincial Centre for Disease Control and Prevention | Key Laboratory of Emergency Detection for Public Health of Zhejiang Province, Zhejiang Provincial Centre for Disease Control and Prevention | Li,C.-X., Li,W., Zhou,J., Zhang,B., Feng,Y., Xu,C.-P., Lu,Y.-Y., Holmes,E.C. and Shi,M. |
| EPI_ISL_2588660 | Medical Microbiology, University Medical Center Utrecht | Medical Microbiology, University Medical Center Utrecht | Tan,L., Viveen,M.C., Lemey,P. and Coenjaerts,F.E. |
| EPI_ISL_2588661 | J. Craig Venter Institute | J. Craig Venter Institute | Lorenzi,H., Town,C., Halpin,R., Bera,J., Ransier,A., Fedorova,N., Stockwell,T., Amedeo,P., Appalla,L., Bishop,B., Edworthy,P., Gupta,N., Hoover,J., Katzel,D., Schobel,S., Shrivastava,S., Thovarai,V., Wang,S., Rebuffo-Scheer,C., Fan,J., He,J., Kehl,S.C., Lederboer,N., Jurgens,L.A., Bose,M.E., Beck,E.T., Kumar,S., Gerna,G., Wentworth,D.E. and Henrickson,K.J. |

| | | | |
|---|---|---|---|
| EPI_ISL_2588662 | Medical Microbiology, University Medical Center Utrecht | Medical Microbiology, University Medical Center Utrecht | Tan,L., Viveen,M.C., Lemey,P. and Coenjaerts,F.E. |
| EPI_ISL_2588663 | Microbiology, Institute of Biological Sciences, Universeity of Sao Paulo | Microbiology, Institute of Biological Sciences, Universeity of Sao Paulo | Di Paola,N., Cunha,M.P., Oliveira,D.B.L., Durigon,E., Durigon,G.S. and Zanotto,P.M.A. |
| EPI_ISL_2588664 | Key Laboratory of Emergency Detection for Public Health of Zhejiang Province, Zhejiang Provincial Centre for Disease Control and Prevention | Key Laboratory of Emergency Detection for Public Health of Zhejiang Province, Zhejiang Provincial Centre for Disease Control and Prevention | Li,C.-X., Li,W., Zhou,J., Zhang,B., Feng,Y., Xu,C.-P., Lu,Y.-Y., Holmes,E.C. and Shi,M. |
| EPI_ISL_2588666 | Virology Laboratory, Dr. Ricardo Gutierrez Children Hospital | Virology Laboratory, Dr. Ricardo Gutierrez Children Hospital | Goya,S., Valinotto,L.E., Tittarelli,E., Rojo,G.L., Greninger,A., Luso,S., Natale,M., Mistchenko,A.S. and Viegas,M. |
| EPI_ISL_2588667 | Mami Nagashima Tokyo Metropolitan Institute of Public Health, Microbiology | Mami Nagashima Tokyo Metropolitan Institute of Public Health, Microbiology | Hasegawa,M., Okazaki,T., Sakamoto,T., Murata,R., Nagashima,M., Shinkai,T. and Sadamasu,K. |
| EPI_ISL_2588668 | J. Craig Venter Institute | J. Craig Venter Institute | Lorenzi,H., Town,C., Halpin,R., Bera,J., Ransier,A., Fedorova,N., Stockwell,T., Amedeo,P., Appalla,L., Bishop,B., Edworthy,P., Gupta,N., Hoover,J., Katzel,D., Li,K., Schobel,S., Shrivastava,S., Thovarai,V., Wang,S., Rebuffo-Scheer,C., Fan,J., He,J., Kehl,S.C., Lederboer,N., Jurgens,L.A., Bose,M.E., Beck,E.T., Kumar,S., Gerna,G., Wentworth,D.E. and Henrickson,K.J. |
| EPI_ISL_2588669 | Key Laboratory of Emergency Detection for Public Health of Zhejiang Province, Zhejiang Provincial Centre for Disease Control and Prevention | Key Laboratory of Emergency Detection for Public Health of Zhejiang Province, Zhejiang Provincial Centre for Disease Control and Prevention | Li,C.-X., Li,W., Zhou,J., Zhang,B., Feng,Y., Xu,C.-P., Lu,Y.-Y., Holmes,E.C. and Shi,M. |
| EPI_ISL_2588684 | J. Craig Venter Institute | J. Craig Venter Institute | Lorenzi,H., Town,C., Halpin,R., Bera,J., Ransier,A., Fedorova,N., Stockwell,T., Amedeo,P., Appalla,L., Bishop,B., Edworthy,P., Gupta,N., Hoover,J., Katzel,D., Li,K., Schobel,S., Shrivastava,S., Thovarai,V., Wang,S., Rebuffo-Scheer,C., Fan,J., He,J., Kehl,S.C., Lederboer,N., Jurgens,L.A., Bose,M.E., Beck,E.T., Kumar,S., Gerna,G., Wentworth,D.E. and Henrickson,K.J. |
| EPI_ISL_2588685 | J. Craig Venter Institute | J. Craig Venter Institute | Lorenzi,H., Town,C., Halpin,R., Bera,J., Ransier,A., Fedorova,N., Stockwell,T., Amedeo,P., Appalla,L., Bishop,B., Edworthy,P., Gupta,N., Hoover,J., Katzel,D., Li,K., Schobel,S., Shrivastava,S., Thovarai,V., Wang,S., Rebuffo-Scheer,C., Fan,J., He,J., Kehl,S.C., Lederboer,N., Jurgens,L.A., Bose,M.E., Beck,E.T., Kumar,S., Videla,C., Wentworth,D.E. and Henrickson,K.J. |
| EPI_ISL_2588688 | J. Craig Venter Institute | J. Craig Venter Institute | Lorenzi,H., Town,C., Halpin,R., Bera,J., Ransier,A., Fedorova,N., Stockwell,T., Amedeo,P., Appalla,L., Bishop,B., Edworthy,P., Gupta,N., Hoover,J., Katzel,D., Schobel,S., Shrivastava,S., Thovarai,V., Wang,S., Rebuffo-Scheer,C., Fan,J., He,J., Kehl,S.C., Lederboer,N., Jurgens,L.A., Bose,M.E., Beck,E.T., Kumar,S., Neumann-Haefelin,D., Wentworth,D.E. and Henrickson,K.J. |
| EPI_ISL_2588720 | Virology Laboratory, Dr. Ricardo Gutierrez Children Hospital | Virology Laboratory, Dr. Ricardo Gutierrez Children Hospital | Goya,S., Valinotto,L.E., Tittarelli,E., Rojo,G.L., Greninger,A., Luso,S., Natale,M., Mistchenko,A.S. and Viegas,M. |
| EPI_ISL_2588721, EPI_ISL_2588722 | Virology Laboratory, Dr. Ricardo Gutierrez Children Hospital | Virology Laboratory, Dr. Ricardo Gutierrez Children Hospital | Goya,S., Valinotto,L.E., Tittarelli,E., Rojo,G.L., Greninger,A., Zaiat,J., Marti,M., Mistchenko,A.S. and Viegas,M. |
| EPI_ISL_2588782 | J. Craig Venter Institute | J. Craig Venter Institute | Lorenzi,H., Town,C., Halpin,R., Bera,J., Ransier,A., Fedorova,N., Stockwell,T., Amedeo,P., Appalla,L., Bishop,B., Edworthy,P., Gupta,N., Hoover,J., Katzel,D., Li,K., Schobel,S., Shrivastava,S., Thovarai,V., Wang,S., Rebuffo-Scheer,C., Fan,J., He,J., Kehl,S.C., Lederboer,N., Jurgens,L.A., Bose,M.E., Beck,E.T., Kumar,S., Gerna,G., Wentworth,D.E. and Henrickson,K.J. |
| EPI_ISL_2588785 | Mami Nagashima Tokyo Metropolitan Institute of Public Health, Microbiology | Mami Nagashima Tokyo Metropolitan Institute of Public Health, Microbiology | Hasegawa,M., Okazaki,T., Sakamoto,T., Murata,R., Nagashima,M., Shinkai,T. and Sadamasu,K. |
| EPI_ISL_2839185, EPI_ISL_2839366, EPI_ISL_2839368, EPI_ISL_2839374 | Centre for Infectious Diseases and Microbiology Laboratory Services | Centre for Infectious Diseases and Microbiology Laboratory Services | "John-Sebastian Eden, Jen Kok, Dominic Dwyer, Edward Holmes, Philip Britton, Alison Kesson, Elena Cutmore, Rachel Tulloch, Bethany Horsburgh" |
| EPI_ISL_2839376 | PathWest Laboratory Medicine WA Microbial Surveillance Unit | PathWest Laboratory Medicine WA Microbial Surveillance Unit | "Chisha Sikazwe, Avram Levy, David Smith, Chris Blyth, Alice Michie, Cara Minney-Smith, David Speers" |
| EPI_ISL_2839416, EPI_ISL_2839417, EPI_ISL_2839442, EPI_ISL_2839449 | Centre for Infectious Diseases and Microbiology Laboratory Services | Centre for Infectious Diseases and Microbiology Laboratory Services | "John-Sebastian Eden, Jen Kok, Dominic Dwyer, Edward Holmes, Philip Britton, Alison Kesson, Elena Cutmore, Rachel Tulloch, Bethany Horsburgh" |
| EPI_ISL_2839450 | Departments of Clinical Microbiology and Infectious Diseases | Centre for Infectious Diseases and Microbiology Laboratory Services | "John-Sebastian Eden, Jen Kok, Dominic Dwyer, Edward Holmes, Philip Britton, Alison Kesson, Elena Cutmore, Rachel Tulloch, Bethany Horsburgh" |
| EPI_ISL_2839452 | Centre for Infectious Diseases and Microbiology Laboratory Services | Centre for Infectious Diseases and Microbiology Laboratory Services | "John-Sebastian Eden, Jen Kok, Dominic Dwyer, Edward Holmes, Philip Britton, Alison Kesson, Elena Cutmore, Rachel Tulloch, Bethany Horsburgh" |
| EPI_ISL_2839453 | PathWest Laboratory Medicine WA Microbial Surveillance Unit | PathWest Laboratory Medicine WA Microbial Surveillance Unit | "Chisha Sikazwe, Avram Levy, David Smith, Chris Blyth, Alice Michie, Cara Minney-Smith, David Speers" |
| EPI_ISL_2839454 | Centre for Infectious Diseases and Microbiology Laboratory Services | Centre for Infectious Diseases and Microbiology Laboratory Services | "John-Sebastian Eden, Jen Kok, Dominic Dwyer, Edward Holmes, Philip Britton, Alison Kesson, Elena Cutmore, Rachel Tulloch, Bethany Horsburgh" |
| EPI_ISL_2839455, EPI_ISL_2839456 | Departments of Clinical Microbiology and Infectious Diseases | Centre for Infectious Diseases and Microbiology Laboratory Services | "John-Sebastian Eden, Jen Kok, Dominic Dwyer, Edward Holmes, Philip Britton, Alison Kesson, Elena Cutmore, Rachel Tulloch, Bethany Horsburgh" |
| EPI_ISL_2839457 | Centre for Infectious Diseases and Microbiology Laboratory Services | Centre for Infectious Diseases and Microbiology Laboratory Services | "John-Sebastian Eden, Jen Kok, Dominic Dwyer, Edward Holmes, Philip Britton, Alison Kesson, Elena Cutmore, Rachel Tulloch, Bethany Horsburgh" |
| EPI_ISL_2989828 | Royal Children's Hospital | WHO Collaborating Centre for Reference and Research on Influenza | Xiaomin Dong, Annette Alafaci,Yi-Mo Deng, Ammar Aziz, Naomi Komadina |
| EPI_ISL_412460, EPI_ISL_412461, EPI_ISL_412863 | Virology Laboratory, Ricardo Gutiérrez Children's Hospital | Virology Laboratory, Ricardo Gutiérrez Children's Hospital / Vanderbilt University Medical Center | Goya, Stephanie; Lucion, Maria Florencia; Juarez, Maria del Valle; Shilts, Meghan; Gentile, Angela; Mistchenko, Alicia S.; Das, Suman & Viegas, Mariana |
| EPI_ISL_412864 | Virology Laboratory, Ricardo Gutiérrez Children's Hospital. | Virology Laboratory, Ricardo Gutiérrez Children's Hospital / Vanderbilt University Medical Center. | Goya, Stephanie; Lucion, Maria Florencia; Juarez, Maria del Valle; Shilts, Meghan; Gentile, Angela; Mistchenko, Alicia S.; Das, Suman & Viegas, Mariana |
| EPI_ISL_412867, EPI_ISL_412868 | Respiratory Virus Unit, Microbiology Services Colindale, Public Health England | Microbiology Services Colindale, Public Health England | Zambon M |
| EPI_ISL_413293, EPI_ISL_413352 | Institut Pasteur de Madagascar | Institut Pasteur de Madagascar | Jean-Michel HERAUD |
| EPI_ISL_4569432 | VIC, Victorian Infectious Diseases Reference Laboratory | WHO Collaborating Centre for Reference and Research on Influenza | Xiaomin Dong, Annette Alafaci,Yi-Mo Deng, Ammar Aziz, Naomi Komadina |
| EPI_ISL_4848359 | Respiratory Virus Unit, National Infection Service, Public Health England | Public Health England, Respiratory Virus Unit, National Infection Service | Zambon M, Talts T, Ellis J, Miah S, Platt S |
| EPI_ISL_5522625, EPI_ISL_5522626, EPI_ISL_5522627, EPI_ISL_5522628, EPI_ISL_5522629 | Institut Pasteur de Côte d'Ivoire | WHO Collaborating Centre for Reference and Research on Influenza | Xiaomin Dong, Herve Kadjo, Yi-Mo Deng, Ammar Aziz, Naomi Komadina |
| EPI_ISL_6174132, EPI_ISL_6174133, EPI_ISL_6174134, EPI_ISL_6174137 | Cote d'ivoire, Institut Pasteur de Cote d'ivoire | WHO Collaborating Centre for Reference and Research on Influenza | Xiaomin Dong, Herve Kadjo, Yi-Mo Deng, Ammar Aziz, Naomi Komadina |
| EPI_ISL_6174138, EPI_ISL_6208721 | Egypt, Central Public Health Laboratory (CPHL) | WHO Collaborating Centre for Reference and Research on Influenza | Xiaomin Dong,Yi-Mo Deng, Amel Naguib, Naomi Komadina |
| EPI_ISL_6268437, EPI_ISL_6268455, EPI_ISL_6268456, EPI_ISL_6268457, EPI_ISL_6268594, EPI_ISL_6268595, EPI_ISL_6268596, EPI_ISL_6268597, EPI_ISL_6268598, EPI_ISL_6268599, EPI_ISL_6268606, EPI_ISL_6268607, EPI_ISL_6268611, EPI_ISL_6268612, EPI_ISL_6268617, EPI_ISL_6268618, EPI_ISL_6268620, EPI_ISL_6268621, EPI_ISL_6268622, EPI_ISL_6268623, EPI_ISL_6268625, EPI_ISL_6268626, EPI_ISL_6268627, EPI_ISL_6268628, EPI_ISL_6314463, EPI_ISL_6314464, EPI_ISL_6314465, EPI_ISL_6314466, EPI_ISL_6314467, EPI_ISL_6314468, EPI_ISL_6314469, EPI_ISL_6314470, EPI_ISL_6314471, EPI_ISL_6314585, EPI_ISL_6314640, EPI_ISL_6314641, EPI_ISL_6314642, EPI_ISL_6314658, EPI_ISL_6314659, EPI_ISL_6314660, EPI_ISL_6314661, EPI_ISL_6314662, EPI_ISL_6314663 | | | |
| see above | Department of Respiratory Medicine, Wuhan Children's Hospital | CAS Key Laboratory of Special Pathogens and Biosafety, Chinese Academy of Sciences | Decheng Wang, Yi Yan, Ying Li, Xiaoxia Lu, Di Liu |
| EPI_ISL_6494786, EPI_ISL_6494789, EPI_ISL_6494790, EPI_ISL_6494791, EPI_ISL_6494795, EPI_ISL_6494797, EPI_ISL_6494798, EPI_ISL_6494799, EPI_ISL_6494801, EPI_ISL_6494802, EPI_ISL_6494803, EPI_ISL_6494804, EPI_ISL_6494810, EPI_ISL_6494812, EPI_ISL_6494813, EPI_ISL_6494814, EPI_ISL_6494817, EPI_ISL_6494825, EPI_ISL_6494826, EPI_ISL_6494827, EPI_ISL_6494896, EPI_ISL_6494897, EPI_ISL_6494898, EPI_ISL_6494899, EPI_ISL_6494900, EPI_ISL_6494901, EPI_ISL_6494902, EPI_ISL_6494903, EPI_ISL_6494904, EPI_ISL_6494905, EPI_ISL_6494906, EPI_ISL_6494907, EPI_ISL_6494908, EPI_ISL_6494910, EPI_ISL_6494912, EPI_ISL_6494913, EPI_ISL_6494914, EPI_ISL_6494915, EPI_ISL_6494916, EPI_ISL_6494917, EPI_ISL_6494918, EPI_ISL_6494919, EPI_ISL_6494920, EPI_ISL_6494921, EPI_ISL_6494922, EPI_ISL_6494923, EPI_ISL_6494924, EPI_ISL_6494925, EPI_ISL_6494926, EPI_ISL_6494927, EPI_ISL_6494928, EPI_ISL_6494929, EPI_ISL_6494930, EPI_ISL_6494931, EPI_ISL_6494932, EPI_ISL_6494933, EPI_ISL_6494934, EPI_ISL_6494936, EPI_ISL_6494937, EPI_ISL_6494938, EPI_ISL_6494939, EPI_ISL_6494940, EPI_ISL_6494941, EPI_ISL_6494942, EPI_ISL_6494943, EPI_ISL_6494944, EPI_ISL_6494945, EPI_ISL_6494946, EPI_ISL_6494947, EPI_ISL_6494948, EPI_ISL_6494949, EPI_ISL_6494950, EPI_ISL_6494951, EPI_ISL_6494952, EPI_ISL_6494956, EPI_ISL_6494959, EPI_ISL_6494961, EPI_ISL_6494964, EPI_ISL_6494968, EPI_ISL_6494969, EPI_ISL_6494972, EPI_ISL_6494978, EPI_ISL_6494982, EPI_ISL_6494986, EPI_ISL_6494987, EPI_ISL_6494989, EPI_ISL_6494990, EPI_ISL_6494994, EPI_ISL_6494999, EPI_ISL_6495000, EPI_ISL_6495002, EPI_ISL_6495005, EPI_ISL_6495006, EPI_ISL_732343, EPI_ISL_732349, EPI_ISL_732350, EPI_ISL_732351, EPI_ISL_732352, EPI_ISL_732353, EPI_ISL_732354, EPI_ISL_732355, EPI_ISL_732356, EPI_ISL_732358, EPI_ISL_732362, EPI_ISL_732363, EPI_ISL_732364, EPI_ISL_732365, EPI_ISL_732366, EPI_ISL_732367, EPI_ISL_732371, EPI_ISL_732373 | | | |
| see above | Respiratory Virus Unit, National Infection Service, Public Health England | National Infection Service, Public Health England | Zambon M, Talts T, Ellis J, Miah S, Platt S |
| EPI_ISL_9003920 | National Institute for Communicable Diseases of the National Health Laboratory Service | National Institute for Communicable Diseases of the National Health Laboratory Service | Amoako DG, Everatt J, Mohale T, Ntuli N, Mahlangu B, Mnguni A, Ismail A, Bhiman JN, Wolter N |
